# N-Acetylglucosamine Facilitates Coordinated Myoblast Flow, Forming the Foundation for Efficient Myogenesis

**DOI:** 10.1101/2024.08.25.609576

**Authors:** Masahiko S. Satoh, Ann Rancourt, Guillaume St-Pierre, Elizabeth Bouchard, Maude Fillion, Kana Hagiwara, Kazuki Nakajima, Sachiko Sato

**Affiliations:** Laboratory of DNA Damage Responses and Bioimaging, Research Centre of CHU de Quebec, Faculty of Medicine, Laval University, 2705 Boulevard Laurier, Quebec, G1V 4G2, Canada; Glycobiology and Bioimaging Laboratory of Research Center for Infectious Diseases and Axe of Infectious and Immunological Diseases, Research Centre of CHU de Quebec, Faculty of Medicine, Laval University, 2705 Boulevard Laurier, Quebec, G1V 4G2, Canada; Glycoanalytical Chemistry Laboratory, Institute for Glyco-core Research (iGCORE), Gifu University, Japan

## Abstract

Skeletal muscle comprises 30-40% of a mammal’s body mass, maintaining its integrity through efficient muscle fiber regeneration, which involves myoblast differentiation into myotubes. Previously, we reported that N-acetylglucosamine (GlcNAc) promotes myogenesis in C2C12 cells, although the underlying mechanisms were unclear. UDP-GlcNAc, the activated form of GlcNAc, is critical for the biosynthesis of highly branched (N-acetyllactosamine-rich) N-linked oligosaccharides, which are recognized by galectin-3 (Gal-3), facilitating dynamic cell-cell and cell-matrix interactions. In this study, we used primary myoblasts from wild-type and Gal-3 null (Gal-3KO) mice, observing myotube formation through long-term live-cell imaging and single-cell tracking. We found that GlcNAc enhances myoblast fusion in a dose-dependent manner, and the addition of Gal-3 with GlcNAc leads to the formation of larger myotubes. Gal-3KO myoblasts exhibited a reduced capacity for myotube formation, a deficiency that was rectified by supplementing with GlcNAc and Gal-3. Our results highlight the critical role of Gal-3 interaction with oligosaccharides whose synthesis was promoted by GlcNAc in facilitating myotube formation. Single-cell tracking revealed that GlcNAc and Gal-3 increase myoblast motility, creating a faster-coordinated cell flow—a directed movement of myoblasts, along which myotubes form through cell fusion. Interestingly, myoblasts contributing to myotube formation were pre-positioned along the eventual shape of the myotubes before the establishment of the coordinated flow. These myoblasts moved along the flow, paused, and even moved against the flow, suggesting that both flow and initial positioning play roles in aligning myoblasts into the shape of a myotube. Overall, our findings demonstrate that GlcNAc, in conjunction with Gal-3, enhances myotube formation by fostering an environment conducive to myoblast positioning, establishing coordinated flow, and facilitating fusion. This suggests potential therapeutic applications of GlcNAc in muscle repair and muscle disorders.

## Introduction

Skeletal muscle, which comprises 30-40% of the total body weight in mammals, is subject to damage from various types of contractions, including repeated eccentric contractions. Efficient regeneration of damaged muscle is critical for mammals to maintain mobility and survival, with myogenesis playing a pivotal role in the regeneration and repair of skeletal muscles (1–3).

Skeletal muscle consists of long multinucleated cells called myofibers which are surrounded by an extracellular matrix known as the basal lamina composed of laminin, collagens, and proteoglycans (4, 5). Upon muscle damage, quiescent muscle stem cells located between the basal lamina and myofibers exit quiescence, proliferate, and differentiate into myogenic lineage cells, known as myoblasts while some of them undergo asymmetric cell divisions for self-renewal (4–6). Most of these cells are destined to become fusion-competent, undergoing fusion to form myotubes. Subsequently, further fusion of myotubes leads to the formation of newly regenerated myofibers. It is suggested that the initial regeneration of a damaged myofiber occurs primarily through muscle stem cells residing on the myofiber itself rather than from stem cells derived from other myofibers (7, 8). Recent studies indicate that within 30 hours after injury, rapid proliferation of myoblasts occurs, with a doubling time as short as 8-10 hours (7, 8). Following 2-3 days post-injury (DPI), massive myoblast proliferation takes place, and the produced cells are randomly distributed. Extensive fusion between myoblasts occurs at 3.5-4.5 DPI, and the enlargement of the myotube continues from 5 to 14 DPI. This indicates that rapid proliferation and fusion of myoblasts, rather than solely relying on sourcing myoblasts from muscle stem cells, may facilitate efficient muscle repair (7, 8).

Studying muscle regeneration *in vivo* in a time-lapsed manner remains challenging and an *in vitro* cell culture system for myotube formation has been used for the investigations of molecules involved in myogenesis(9–13). The fusion process of primary myoblasts isolated from skeletal muscle or established cultured immortal myoblast lines, such as C2C12, is induced by changing the growth medium, which contains serum and basic fibroblast growth factor, to a medium containing a lower concentration of serum (14). Although various factors can influence fusion efficiency and the speed of myotube formation, this culture system enables the identification of key factors such as myomaker and myomerge, which are critical for myoblast fusion (1, 9–13). Previously, we reported that the addition of N-acetylglucosamine (GlcNAc) and galectin-3 (Gal-3) to C2C12 cells enhances myotube formation (15). Additionally, GlcNAc has been shown to mitigate the muscle dystrophy progression in a mouse model of Duchenne mascular dystrophy (15). Thus, GlcNAc and Gal-3 play a role in the process of myotube formation, although the mechanisms of this enhancement are not fully understood.

GlcNAc, an endogenous precursor, is endocytosed by cells and converted into its active form, UDP-GlcNAc, a substrate for various N-acetylglucosaminyltransferases. These oligosaccharide-processing enzymes, including mannosyl glycoprotein acetylglucosaminyltransferases (MGATs), are involved in the biosynthesis of Asn (N)-linked oligosaccharides in the secretory pathway(16, 17). N-linked oligosaccharides play a critical role in regulating cell-cell and cell-matrix interactions, as well as signal transduction, mediated by oligosaccharide-recognition proteins, including galectins(18–24). Gal-3 is one of the most abundant proteins, with its expression being regulated during myogenesis (25). Upon binding to its oligosaccharide ligands, Gal-3 oligomerizes, influencing cell-laminin and cell-cell interactions(18–24, 26–30). Specifically, Gal-3 shows a strong binding preference for N-linked glycans carrying at least three N-acetyllactosamine (Galactose-GlcNAc, hereafter referred to as lactosamine) residues (26, 31–33), whose biosynthesis is highly dependent on the intracellular concentration of UDP-GlcNAc, due to MGAT5’s low affinity for UDP-GlcNAc (Km > 10 mM)(19, 34–37). Therefore, the usage of GlcNAc in this context is crucial as it can enhance UDP-GlcNAc levels, subsequently influencing the availability of glycan ligands for Gal-3 binding. Previous studies suggest that GlcNAc supplementation increases intracellular UDP-GlcNAc levels (19, 35–37).

In our current work, we employed live cell imaging (38–40) to accurately monitor the process of myoblast fusion and myotube formation using primary wild-type and Gal-3 knockout (Gal-3KO) myoblasts. Simultaneous monitoring of myoblasts under different treatment conditions, alongside controls, enabled an accurate assessment of the myogenesis process. Each video-recorded myoblast was individually tracked to reveal the progression of myotube formation, including myoblasts that did not form myotubes, providing a comprehensive overview of each myoblast’s contribution to myogenesis. We found that myoblast fusion occurred in a GlcNAc dose-dependent manner and that Gal-3 further enhanced fusion in wild-type myoblasts. Conversely, Gal-3KO myoblasts showed a reduced ability to form myotubes, a deficiency compensated by GlcNAc and Gal-3. Furthermore, we observed that GlcNAc promoted the synthesis of N-linked oligosaccharides rich in lactosamine. These results suggest that GlcNAc-optimized biosynthesis of lactosamine-rich oligosaccharides in conjunction with Gal-3, plays a critical role in myoblast fusion. Furthermore, the addition of GlcNAc and/or Gal-3 results in increased cell motility, which could relate to the formation of a faster-coordinated flow of myoblasts. This flow, represented by the movement of myoblasts in a certain direction, results in the formation of myotubes along the direction of flow. Myoblasts involved in myotube formation were aligned before the flow was fully established, and the cells aligned moved to follow the flow, resist it, or oppose the flow, suggesting that such movement could contribute to generating a myotube. These results also suggest that myotube formation could occur by the alignment of myoblasts along the eventual shape of myotube. Thus, GlcNAc and Gal-3 create an environment that facilitates these processes, resulting in efficient myotube formation. Creating such an environment can thus be used to promote the repair of damaged muscles caused by contractions and genetic disorders.

## Results

### Monitoring Myotube Formation through Single-Cell Tracking

During the differentiation process of primary myoblasts in culture, we observed significant variability in the rate and efficiency of myotube formation, influenced by factors including cell passage number. Typically, myotube formation varied from 3 to 5 days. To facilitate quantitative analysis of myotube formation, independent of these variable factors, we employed a live cell imaging-assisted single-cell lineage tracking system (Fig. 1)(38–40). Myoblasts were cultured under various conditions, including a control setup. We continuously monitored these myoblasts until the formation of mature myotubes, using differential interference contrast (DIC) imaging(38–40)(Fig. 1). This approach allowed us to normalize variations in myotube formation timing across experiments. Myoblasts captured in the videos were identified using grayscale image segmentation, assigned unique identifiers, and tracked until myotube formation occurred (38–40). This single-cell tracking also included myoblasts that did not form myotubes, providing a comprehensive view of myoblast behavior during myogenesis. The tracking data was used for bioinformatics analysis to generate maps depicting each myoblast’s fusion into myotubes by software utilized for this study has been made available on GitHub (DOI 10.5281/zenodo.12988726), and results using this software have been previously reported(38, 39). Representative still images of myoblast-to-myoblast, myoblast-to-myotube, and myotube-to-myotube fusion are shown in Fig. 1B, with corresponding videos provided as Supplementary Video 1, 2, and 3. To ensure objective analysis, we initially selected 100 centrally located myoblasts at 0 minutes and tracked each one. If a myoblast moved out of the image frame, it was excluded, and another myoblast close to the center of the imaging area was selected and subsequently tracked. If the tracked myoblast contributed to the formation of a myotube, other myoblasts involved in the formation of the myotube were identified by retrospectively tracking their fusion partners, thereby completing the cell lineage fusion map.

**Fig. 1.**
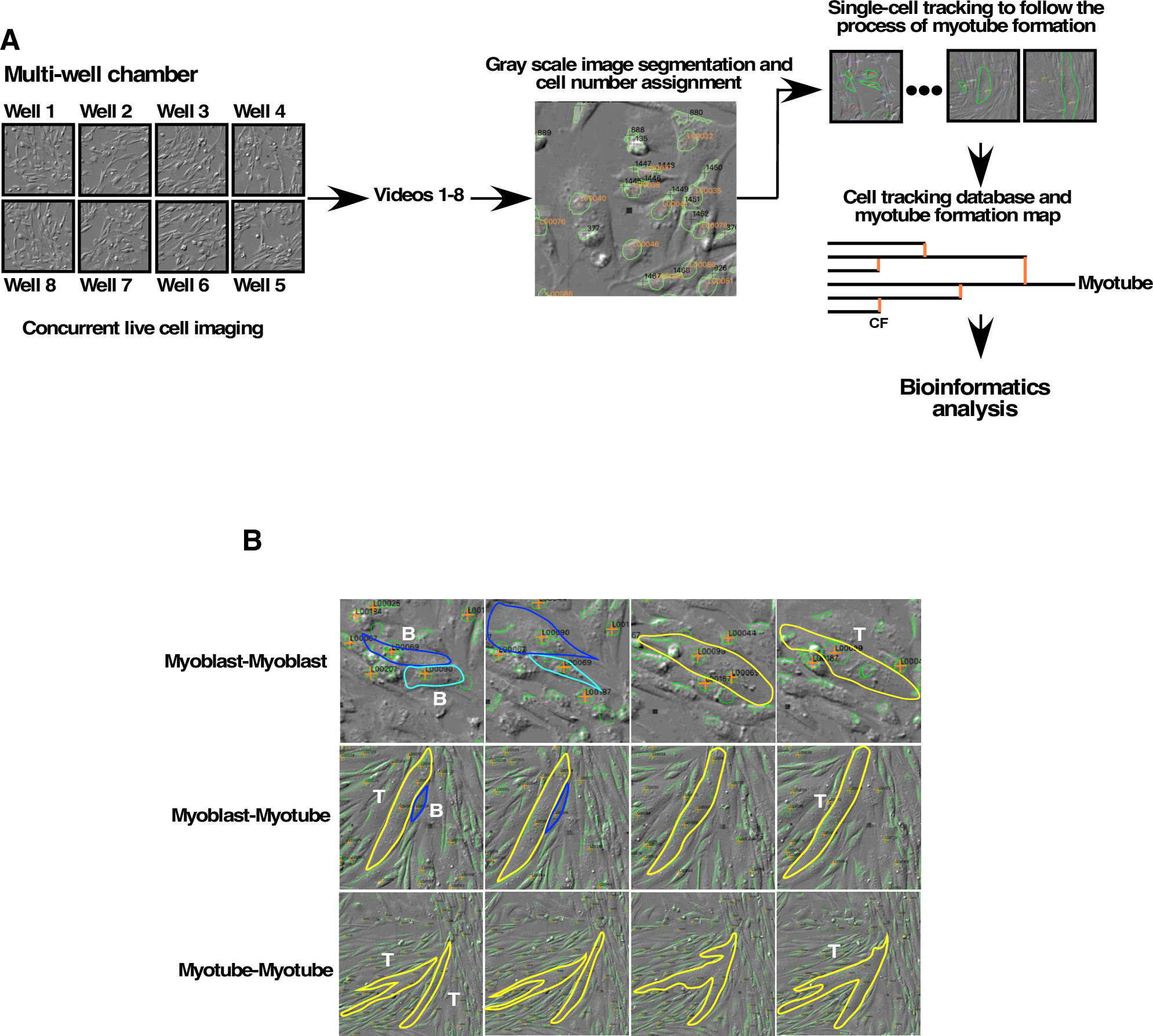
System for Investigating Myotube Formation Using Long-Term Live Cell Imaging, Single-Cell Tracking, and Bioinformatics Analysis. **A.** Primary mouse myoblasts were cultured in multi-well chamber slides. Areas of interest were monitored using differential interference contrast (DIC) imaging. Treatments included Gal-3, GlcNAc, and a combination of both, with one well serving as a control. Up to 8 live cell imaging videos were generated in real-time, subjected to image segmentation, identification, and tracking of myoblasts. A database was created to record myoblast positions at each time point and the occurrences of fusion events. Maps illustrating myotube formation processes were generated for bioinformatics analysis. **B.** Still images of representative myoblast-myoblast, myoblast-myotube, and myotube-myotube fusions are displayed. Myoblasts are marked with blue and light blue circles and myotubes with yellow circles. Corresponding videos are available as Supplementary Videos 1, 2, and 3.

### GlcNAc Increases Dose-Response Fusion of Myoblasts

Using live-cell imaging and single-cell tracking, we analyzed the impact of various GlcNAc concentrations on myoblast fusion. Imaging commenced 2-3 hours after switching from growth to differentiation medium. Representative cases of myotube formation for Control, and 1 mM, 5 mM, and 10 mM GlcNAc treatments are shown in Fig. 2 A-D. Comprehensive cell-lineage fusion maps and tracking videos are included in Supplementary Data 1-4 and Supplementary Videos 4-7. The fusion map for the Control (Fig. 2A) shows six myoblasts formed a myotube, with fusion events marked in orange. The single-cell lineage analysis approach allowed us to quantify fusion events at the single-cell level and plot them against GlcNAc concentration (Fig. 2E), revealing a dose-dependent increase in cell fusion events and the formation of larger myotubes arising from primary myoblasts, consistent with previous findings using C2C12 cells (15).

**Fig. 2.**
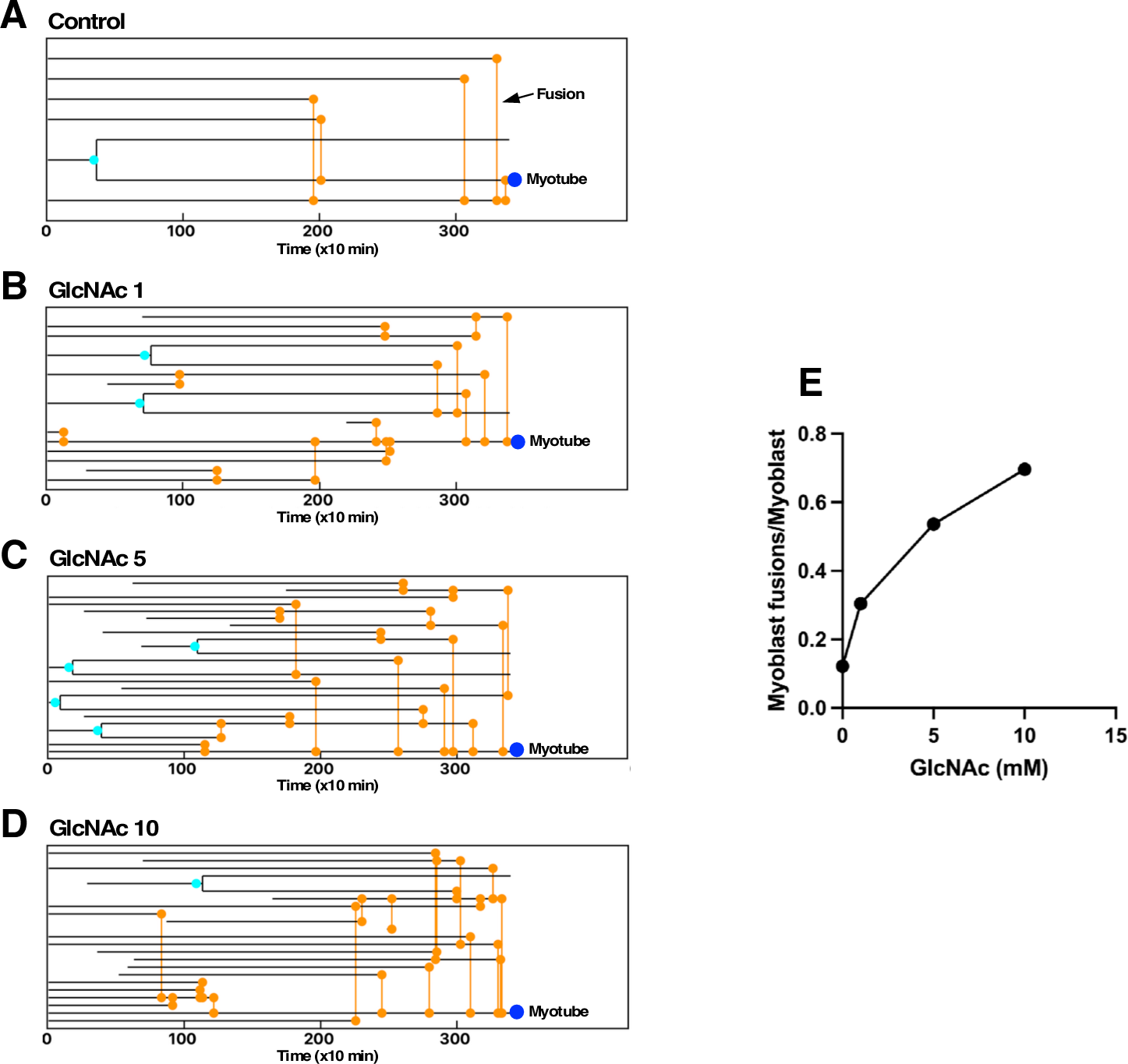
GlcNAc Dose-Dependent Promotion of Myotube Formation. Maps depicting myoblast fusion and myotube formation for Control (**A**), 1 mM GlcNAc (**B**), 5 mM GlcNAc (**C**), and 10 mM GlcNAc (**D**) are presented. Black lines represent myoblasts or myotubes. Orange circles and lines indicate fusion events and blue circles represent resulting myotubes. Some myoblasts enter the imaging frame after time 0, appearing as black lines post-time 0. **E.** A dose-dependent increase in fusion events is charted, based on counts of fusion events per treatment, expressed as the number of fusions per myoblast tracked. Primary wild-type myoblasts were used. Representative results from one of two independent experiments are shown.

### Uptake of GlcNAc by differentiating myoblasts

GlcNAc enhances myotube formation, implying the uptake of GlcNAc impacts the process of N-linked glycosylation. To determine if GlcNAc was assimilated by myoblasts, lysates from myoblasts exposed to GlcNAc for 24 hrs were analyzed for the intracellular concentration of UDP-GlcNAc. The analysis showed a significant increase in UDP-N-acetylhexosaminase (UDP-HexNAc that included UDP-GlcNAc) levels (Fig. 3A), a precursor and critical regulator of the synthesis of lactosamine-rich N-linked oligosaccharide, while other monosaccharide metabolites like CMP-Neu5Ac, GDP-Man, and UDP-GlcA remained unchanged (Figs. 3B-D). These results demonstrate that GlcNAc is incorporated and converted to UDP-GlcNAc in myoblasts.

**Fig. 3.**
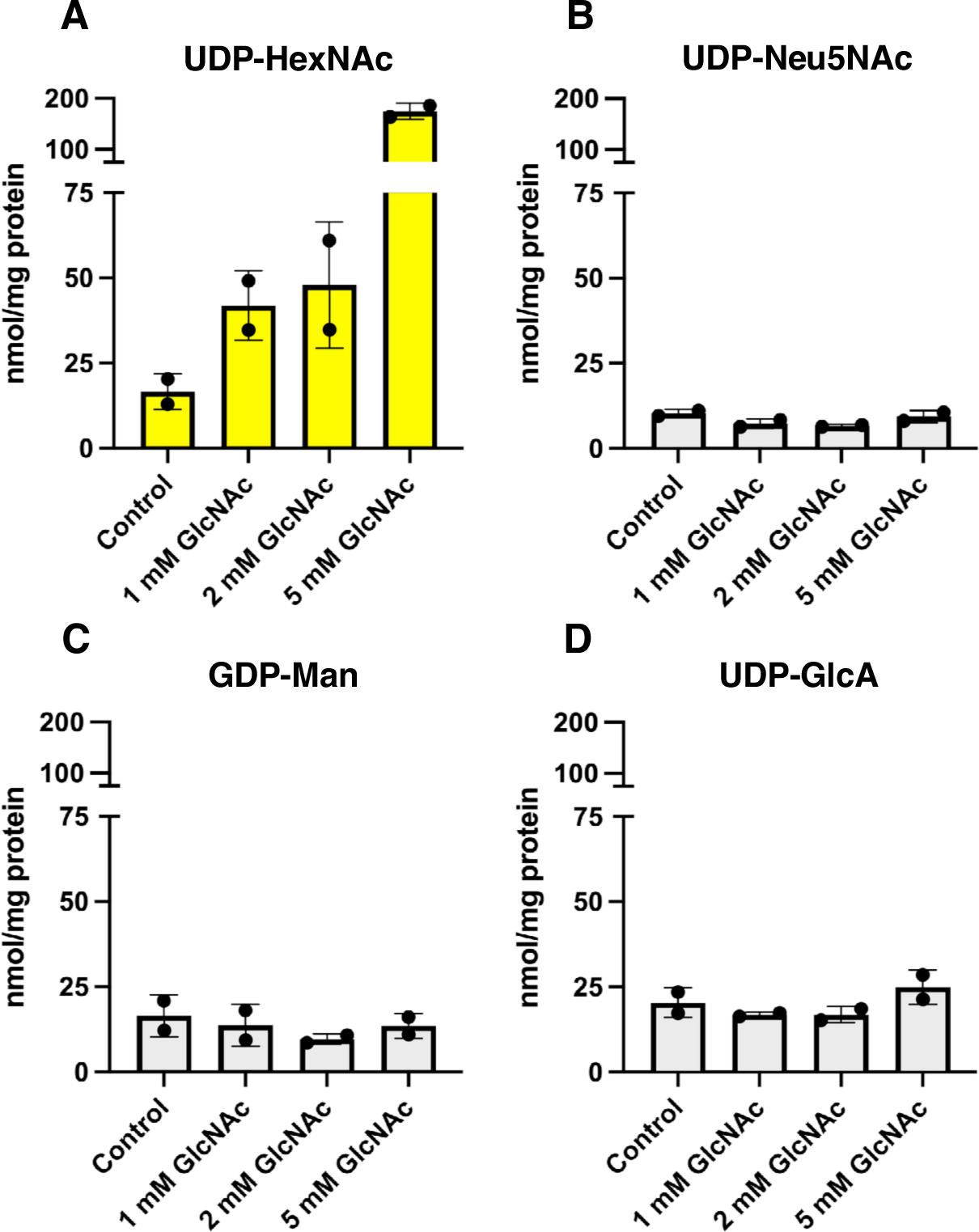
Assimilation of GlcNAc by Myoblasts. **A-D.** Intracellular levels of monosaccharide nucleotides were measured using mass spectrometry. Due to the indistinguishability of UDP-GlcNAc and UDP-N-acetylgalactosamine by the mass spectrometer, they are collectively reported as UDP-N-acetylhexosamine (UDP-HexNAc; **A**). Levels of cytidine monophosphate N-acetylneuraminic acid (CMP-Neu5Ac; **B**), guanosine diphosphate mannose (GDP-Man; **C**), and uridine diphosphate glucuronic acid (UDP-GlcA; **D**) are also shown. Data represented with ranges indicated.

### Effect of N-linked Oligosaccharide Synthesis on Myogenesis

To investigate the role of lactosamine-rich N-linked oligosaccharides in myotube formation, we used kifunensine, which inhibits N-linked oligosaccharide processing in the Golgi apparatus, thereby reducing the synthesis of these glycans (41). Western blotting with L-phytohemagglutinin (L-PHA), which binds to these N-linked oligosaccharides(35), revealed that GlcNAc increased the presence of proteins (75-100 KDa) modified with these N-linked oligosaccharides (Fig. 4A and B). In contrast, the level of Concanavalin A (Con A)-binding high-mannose-rich glycans, which are precursors of complex type N-linked oligosaccharides, did not change with GlcNAc treatment (Supplement Fig. 1). Kifunensine, an inhibitor of endoplasmic reticulum α-Mannosidase-I, inhibited the synthesis of lactosamine-rich N-glycan (Fig. 4A-C) while increasing the accumulation of processing intermediate oligosaccharides, rich in mannose, as detected by Con A binding (Supplementary Fig. 1). Furthermore, indirect immunofluorescence staining using myosin heavy chain (MHC), a marker for myotubes and DAPI showed that kifunensine reduced myotube formation (Fig. 4D and E), suggesting that the presence of lactosamine-rich N-linked oligosaccharides whose synthesis is promoted by GlcNAc is crucial for effective myotube formation.

**Fig. 4.**
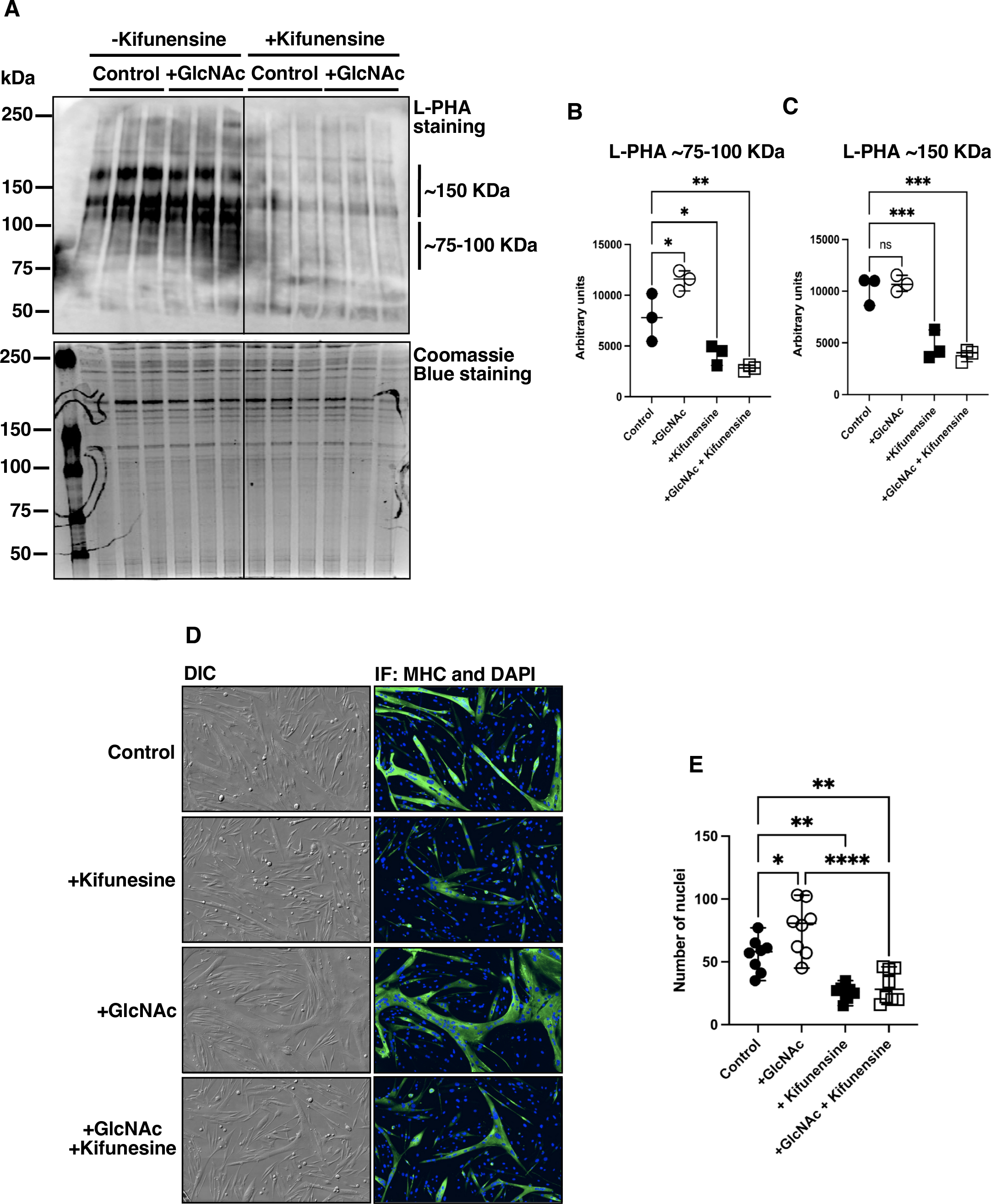
Inhibition of Lactosamine-rich N-linked Oligosaccharide Synthesis and Myotube by Kifunensine. **A.** Primary wild-type myoblasts were cultured in differentiation mediums with 5 mM GlcNAc and/or 40 μM kifunensine for 24 hours. Myoblasts were then harvested and analyzed by Western blotting with L-phytohemagglutinin (L-PHA), which binds to N-linked oligosaccharides. Staining with L-PHA and Coomassie brilliant blue is shown. The intensity of bands corresponding to proteins in the 75-100 kDa (**B**) and around 150 kDa (**C**) range are quantified. Representative results from one of two independent experiments are shown. **D.** Primary wild-type myoblasts were cultured in a differentiation medium with 2 mM GlcNAc and/or 40 μM kifunensine for approximately 66 hours until myotube formation. Myoblasts and myotubes were fixed, and myosin heavy chain (MHC, green) was visualized using indirect immunofluorescence (IF). Nuclei were stained with DAPI (blue). **E.** Nuclei within myotubes were counted; images (2560 x 2560 pixels) were divided into 4 frames to count nuclei per myotube in each frame, with 8 frames analyzed per treatment. Representative results from one of two independent experiments are shown. **B, C,** and **E.** Statistical analysis was performed using one-way ANOVA (Tukey’s test), and significance is indicated as *P < 0.01, **P < 0.05, ***P < 0.001, and ****P < 0.0001, while “ns” indicates no significance. Data represent means ± standard errors from eight samples.

### Promotion of Myotube Formation by GlcNAc and Gal-3 in Wild-Type Myoblasts and the Impact of Gal-3 Knockout

One of the major oligosaccharide-binding proteins expressed in differentiating myoblast is galectin-3. Our previous study also suggests that galectin-3 promotes the myogenesis of C2C12 myoblasts (15). Indeed, the mouse primary myoblasts continuously released galectin-3 (Supplementary Fig. 2). To further explore the processes of myotube formation, we conducted analyses using myoblasts treated with 2 mM GlcNAc, Gal-3, and a combination of both. In these experiments with wild-type myoblasts, myotube formation was observed by 1,700 minutes after initiating differentiation, compared to the 3,200 minutes noted in Fig. 2 under control conditions. Due to this variability, we monitored the myoblasts in real time to precisely determine the duration required for live cell imaging. Representative results of myotube formation in the presence of Gal-3, and GlcNAc, and both are shown in Fig. 5A-D. Comprehensive maps and corresponding videos for single-cell tracking of Control, +Gal-3, +GlcNAc, and +Gal-3+GlcNAc treatments are available in Supplementary Data 5, 6, 7, and 8, and Supplementary Videos 8, 9, 10, and 11, respectively. The maps shown in Fig. 5A-D illustrate that myotube formation involved 11, 40, 53, and 62 myoblasts for Control, +Gal-3, +GlcNAc, and +Gal-3+GlcNAc, respectively, indicating enhanced myotube formation with +Gal-3 and +GlcNAc, with the most substantial effect observed with the combination of both.

**Fig. 5.**
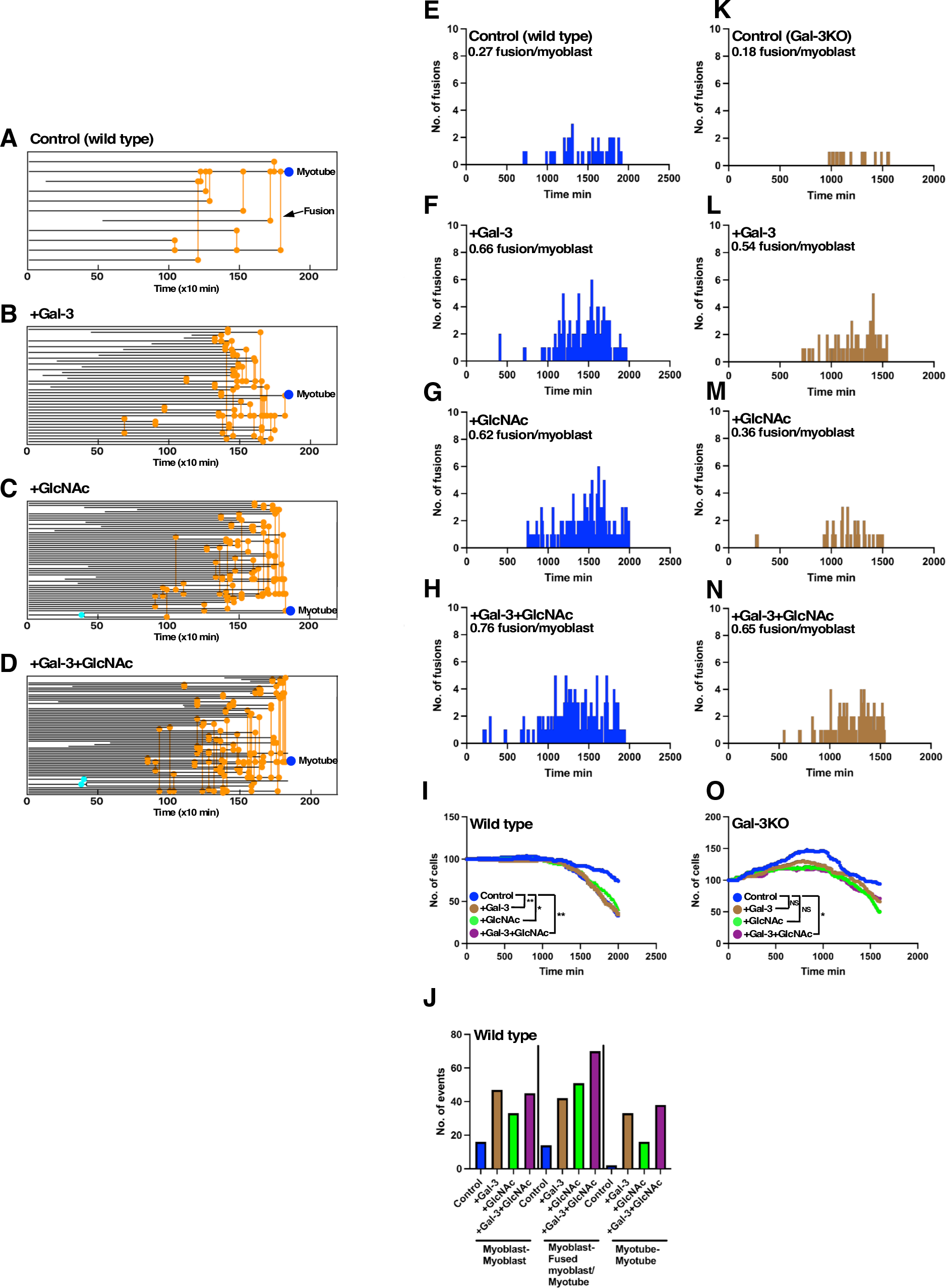
Effect of Gal-3 and GlcNAc on Myotube Formation. **A-D.** Maps display myoblast fusion and myotube formation for Control (**A**), 2 μM Gal-3 (**B**), 10 mM GlcNAc (**C**), and a combination of 2 μM Gal-3 with 10 mM GlcNAc (**D**). Black lines depict myoblasts or myotubes, orange circles, and lines indicate fusion events, and blue circles represent resulting myotubes. Some myoblasts that enter the imaging frame post-time 0 are shown as black lines. **E-H.** The number of fusion events per tracked myoblast for Control (**E**), +Gal-3 (**F**), +GlcNAc (**G**), and +Gal-3+GlcNAc (**H**) is plotted over time. **I.** The number of myoblasts at each time point was plotted. The differences between consecutive two-time points were determined and used for statistical analysis. **J.** The number of fusion events occurring between myoblasts, between a myoblast and a fused myoblast or myotube, and between myotubes was plotted. **K-N.** The number of fusion events for primary Gal-3KO myoblasts for Control (**K**), +Gal-3 (**L**), +GlcNAc (**M**), and +Gal-3+GlcNAc (**N**) is plotted over time. **O.** The number of myoblasts at each time point was plotted. The differences between consecutive two-time points (time 0 to 1000 min) were determined and used for statistical analysis. **I and O.** Statistical analyses were performed using one-way ANOVA (Tukey’s test), with significance indicated as *P < 0.01, **P < 0.05, and “ns” for no significance. Representative results from one of two independent experiments are shown.

Fig. 5E-H plots the number of fusions at each time point, peaking around 1,500 minutes for all conditions. However, no significant shift in the timing of myoblast fusion relative to the Control was observed, suggesting that +Gal-3, +GlcNAc, and +Gal-3+GlcNAc primarily increase the frequency of fusions at each time point compared to the Control. Statistical analysis of the cell fusion effects, shown in Fig. 5I, indicates that the number of myoblasts decreases following fusions (a fused myotube is counted as one), with statistically significant differences between Control and each of the treatments with Gal-3 and/or GlcNAc. Furthermore, the addition of Gal-3, GlcNAc, or both resulted in an increased number of fusion events between myoblasts, between a myoblast and a fused myoblast or myotube, and between myotubes (Fig. 5J and Supplementary Data 13), indicating that Gal-3 and GlcNAc also enhance the fusion process that occurs during the formation of mature myotubes.

Further analysis was conducted on myoblasts isolated from Gal-3 knockout (Gal-3KO) mice (Fig. 5K-N; comprehensive maps and videos are in Supplementary Data 9, 10, 11, and 12, and Supplementary Videos 12, 13, 14, and 15). Compared to the wild-type, the fusion competence of Control Gal-3KO myoblasts was notably reduced (Fig. 5E and K). However, the addition of Gal-3 or GlcNAc improved fusion efficiency by 2 to 3-fold (Fig. 5L and M), with the most significant enhancement observed with the combination of both (Fig. 5N). Interestingly, while Gal-3KO myoblasts continued to proliferate in the differentiation medium (Fig. 5O), the addition of Gal-3 and/or GlcNAc mitigated this increase, with a statistically significant effect noted with the combination of both. These results underscore that Gal-3 is essential for efficient myotube formation and that the interaction between Gal-3 and lactosamine-rich N-linked oligosaccharides, whose synthesis is promoted by GlcNAc facilitates this process.

### Impact of Gal-3 and GlcNAc on Myoblast Motility

To further investigate the effect of Gal-3 and GlcNAc on myotube formation, we focused on the motility of myoblasts, as changes were observed through visual inspection of live cell imaging videos (Fig. 6). Using single-cell tracking data, we calculated the moving distance of each myoblast from Time A to Time A+1 and determined the average motility at every time point. This analysis included all tracked myoblasts, regardless of whether they were involved in myotube formation or not (Fig. 6A), those specifically involved in myotube formation (Fig. 6B), and those that did not fuse (Fig. 6C). Fig. 6A shows that myoblast motility decreased over time after switching to the differentiation medium. However, statistically significant increases in motility were observed relative to Control with the addition of Gal-3, GlcNAc, or both, regardless of whether the myoblasts eventually formed myotubes, indicating a global effect of Gal-3 and GlcNAc on motility (Fig. 6A-C). In the case of Gal-3 knockout (Gal-3KO) myoblasts, the effects of Gal-3 and GlcNAc were reversed compared to wild-type myoblasts (Fig. 6D-E). Gal-3KO myoblasts exhibited higher motility compared to the wild-type control, and the addition of both Gal-3 and GlcNAc significantly reduced their motility. These findings suggest that Gal-3 and GlcNAc act as regulators to optimize motility, thereby favoring efficient myotube formation.

**Fig. 6.**
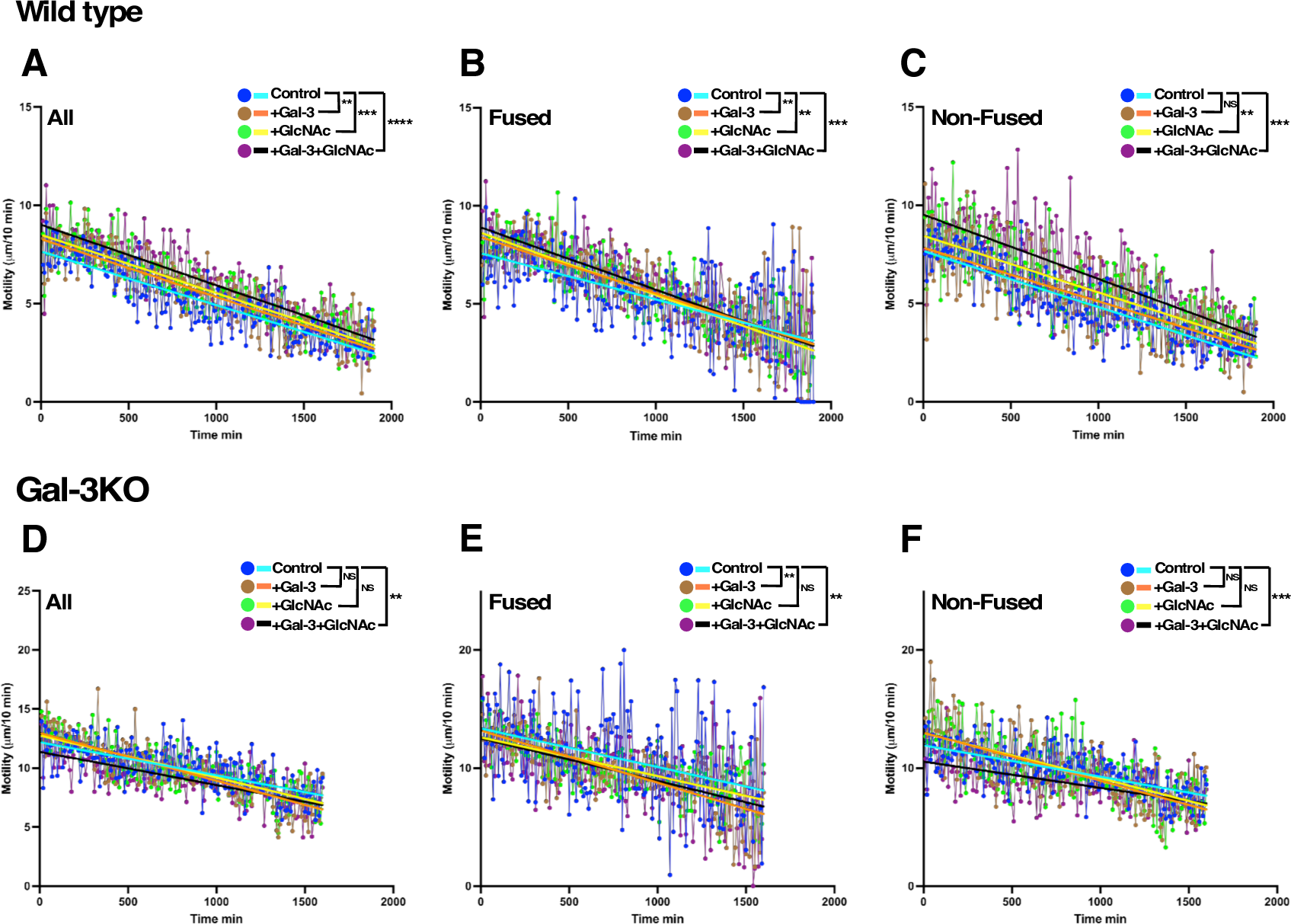
Motility Analysis of Myoblasts. **A-F.** The motility of primary wild-type (**A-C**) and Gal-3-KO (**D-F**) myoblasts was analyzed by determining the distance moved between consecutive time points. **A** and **D.** Average motilities were calculated for all recorded myoblasts in the single-cell tracking database, with results plotted at each time point. Additional analyses were conducted for myoblasts involved in myotube formation (**B** and **E**) and those that did not fuse during tracking (**C** and **F**). Statistical analysis was performed using one-way ANOVA (Tukey’s test), with significance denoted as **P < 0.05, ***P < 0.001, ****P < 0.0001, and “ns” indicating no significance. Simple linear regression lines are shown to illustrate trends.

### Formation of Coordinated Flow

Further insights into myoblast motility were gained by visually inspecting live cell imaging videos, which showed myoblasts beginning to move in a directed manner, creating what appeared to be a flow. In control, myoblasts (Fig. 7A), these flows, clearly visible at 1,000 minutes (Supplementary Video 8), were formed. With the addition of Gal-3 and/or GlcNAc, longer flows were observed (Fig. 7B, C, D, and Supplementary Videos 9, 10, 11). There was no specific directionality to the flows, suggesting they were determined by the coordinated behavior of the myoblasts. To quantify the speed of the flow, we identified myoblasts that moved for 300 minutes in a relatively straight line without pausing or changing direction. Fig. 7E-G displays the moving distances for Control, +Gal-3, +GlcNAc, and both. The length of the green lines represents the speed of movement. Quantitative results, expressed in μm per 10 min, are shown in Fig. 7I, indicating that faster-coordinated flows relative to the Control are generated by Gal-3 and/or GlcNAc. Similar flows were observed with Gal-3KO myoblasts, where Control cells (Fig. 7J) formed longer flows compared to those treated with +Gal-3, +GlcNAc, or both (Fig. 7K-L, and Supplementary Videos 12, 13, 14, and 15). Further analysis, identifying myoblasts moving for 200 minutes in a straight direction (Fig. 7N, O, P, and Q), revealed that Control Gal-3KO myoblasts exhibited faster flows than wild-type myoblasts, but the speed of flows was reduced with the addition of Gal-3, GlcNAc, and both (Fig. 7R). Notably, myoblasts lacking Gal-3 formed fast flows but failed to form myotubes, instead developing into thin, spindle-shaped cells (see Supplementary Video 12). Thus, it appears that creating movement of myoblasts at an optimal speed and its motility is crucial for efficient myotube formation.

**Fig. 7.**
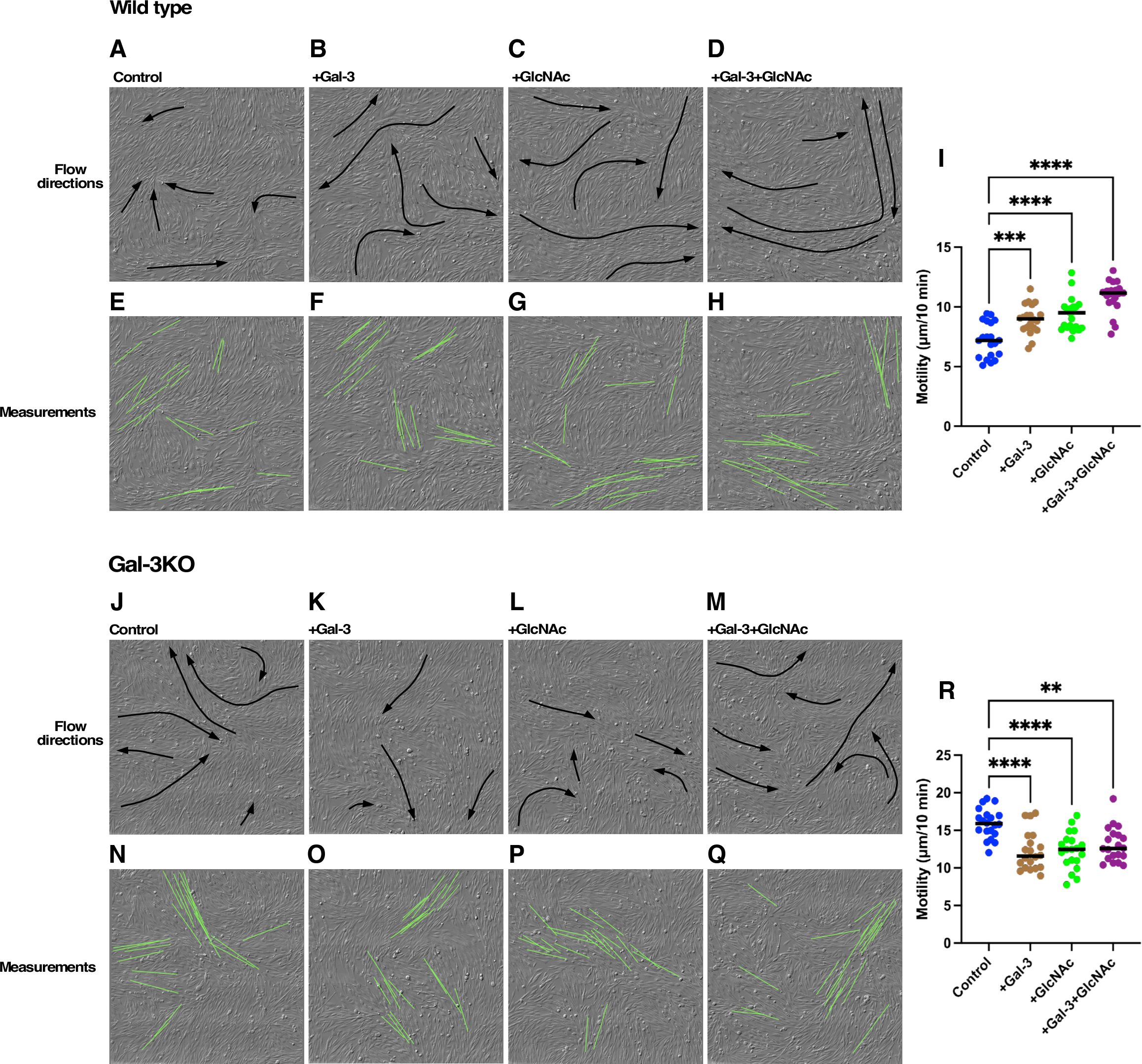
Formation of Coordinated Flow. Live cell imaging videos of wild-type myoblasts (**A-D**) and Gal-3KO myoblasts (**J-M**) were inspected to identify coordinated flows. These flows are outlined with black lines, and their directions are indicated by arrows at 1,000 minutes for wild-type and 1,100 minutes for Gal-3KO myoblasts. To measure the speed of coordinated flow for Control (**E** and **N**), +Gal-3 (**F** and **O**), +GlcNAc (**G** and **P**), and +Gal-3+GlcNAc (**H** and **Q**), myoblasts moving in a straight trajectory without pausing or changing direction between 700 to 1,000 minutes for wild-type and 900 to 1,100 minutes for Gal-3KO were analyzed. The start and end positions of these movements are represented by green lines. Motilities, expressed as µm/10 min, were plotted (**I** and **R**). Statistical analysis (n = 20) was performed using one-way ANOVA (Tukey’s test), with significance indicated by **P < 0.05, ***P < 0.001, and ****P < 0.0001.

### Alignment of Myoblasts Along the Eventual Shape of the Myotube

It is plausible that myoblasts with optimal motility and directional movement have a higher chance of encountering each other to form a myotube. To test this hypothesis, we selected the largest myotube for each condition (Control, +Gal-3, +GlcNAc, and +Gal-3+GlcNAc) among myotubes identified by single-cell tracking and selected all myoblasts that contributed to its formation (Fig. 8A, green outlines; in the case of Control and +Gal-3+GlcNAc, the second largest myotube was also analyzed, shown in orange outlines). Myoblasts that moved into the imaging frame after 0 minutes were excluded from the analysis. The positions of myoblasts were then marked in images from 0 to 2,000 minutes (Fig. 8A). Although we anticipated observing myoblasts moving along a coordinated flow to form a myotube upon encountering a fusion-competent myoblast or myotube, Fig. 8A reveals that myoblasts were already aligned along the eventual shape of the myotube by the 400 minutes before any clear coordinated flows had formed. For instance, myoblasts treated with GlcNAc were already outlining the shape of the myotube by time 400 min despite that no myotubes were formed. The direction and length of the flow, as depicted in Fig. 7A-D, could have influenced myotube formation, yet myoblasts positioned themselves in the shape of the myotube even before the flow solidified. Despite their alignment, these myoblasts were not static. Fig. 8B overlays the trajectory of myoblasts on their positions at time 0 min, indicating that myoblasts that formed the myotube were actively moving but remained along the eventual shape of the myotube. Supplementary Video 16 demonstrates the behavior of GlcNAc-treated myoblasts, their trajectories, and the resulting myotube formation. The video also illustrates the formation of coordinated flows and the dynamics of each myoblast within these flows. Interestingly, some myoblasts moved along the flow, paused, and then moved in opposition to the flow as if they were using the force of the flow to position themselves.

**Fig. 8.**
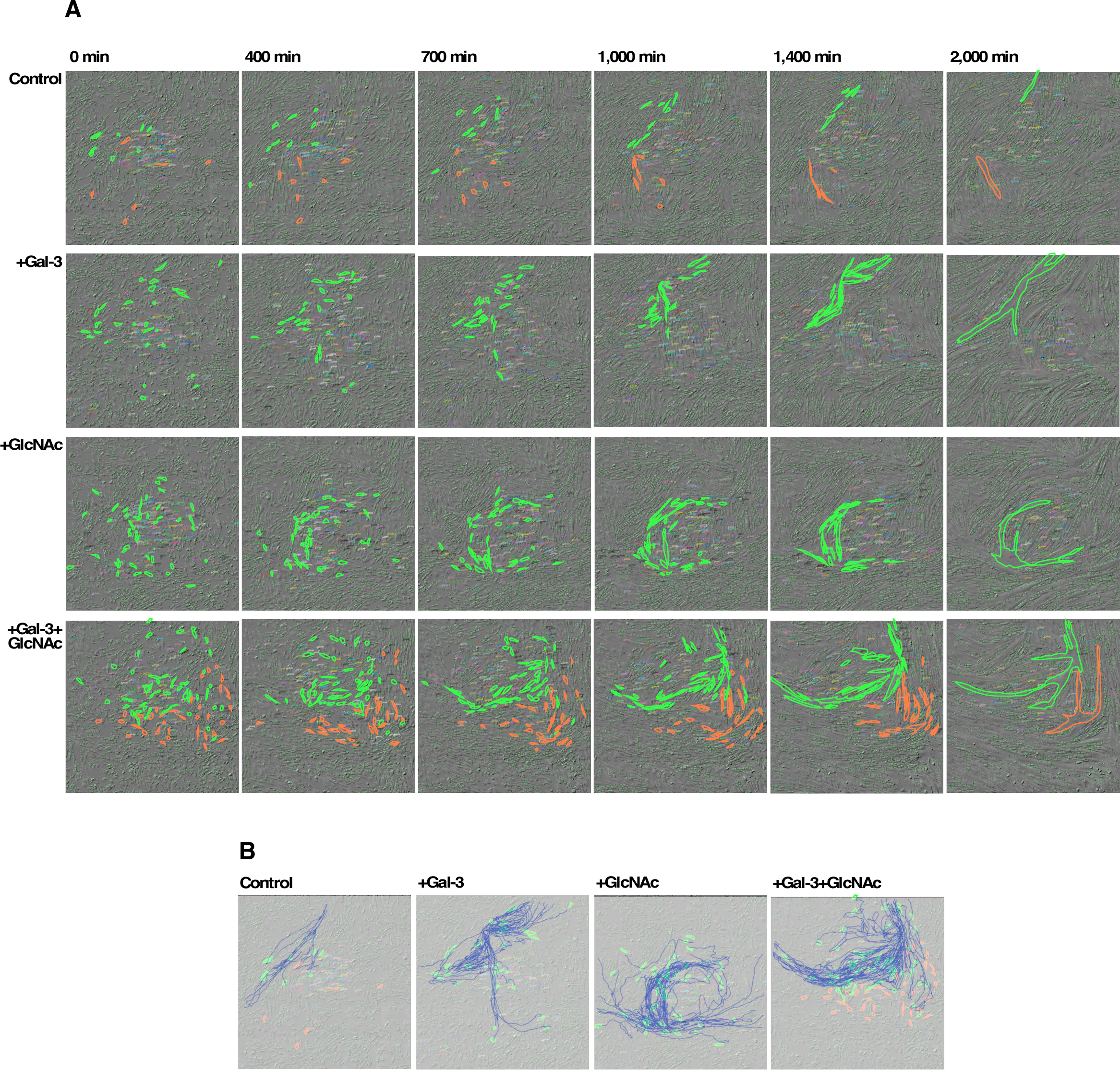
Depicting the Process That Individual Myoblasts Form a Myotube. **A.** The largest (outlined in green) and second largest (outlined in orange) myotubes identified by single-cell tracking in each treatment were selected, and the myoblasts that formed these myotubes were identified. Myoblasts entering the image frame from outside during the experiment were excluded. Each myoblast and the resulting myotube formations were outlined at every time point. **B.** The trajectories of myoblasts from 0 minutes until fusion were overlaid on the image from 0 minutes.

### Analysis of Individual Myoblast Trajectories

To gain further insight into myoblast behavior during myotube formation, we compiled a panel of trajectories for individual myoblasts treated with GlcNAc (Fig. 9A). The initial position at 0 minutes is indicated by a black circle and the point of fusion by a brown square. The time at which the coordinated flow was approximately formed and each myoblast positioned along the shape of the myotube is marked by a blue circle at 350 minutes. Moving patterns varied; some myoblasts moved straight, then paused, and then moved back toward their original position. In some instances, myoblasts moved in one direction and fused. No specific pattern of movement was discernible, nor was there a clear pattern in the moving distance from 0 to 350 minutes, or from 350 minutes to the point of fusion. Therefore, each myoblast’s trajectory was unique, yet all converged to form a myotube. We also calculated the difference in motility between myoblasts that formed myotubes and the speed of the coordinated flow (Fig. 9B). Results showed that the motility of myoblasts forming myotubes was slower than the speed of the coordinated flow, and this difference was more pronounced when myoblasts were treated with both Gal-3 and GlcNAc. This suggests that the movement of myoblasts along the flow, pausing, or moving against the flow, plays a role in enabling myoblasts to form mature myotubes. Therefore, Gal-3 and GlcNAc contribute to creating an environment conducive to myotube formation, and ultimately, to the formation of myofibers.

**Fig. 9.**
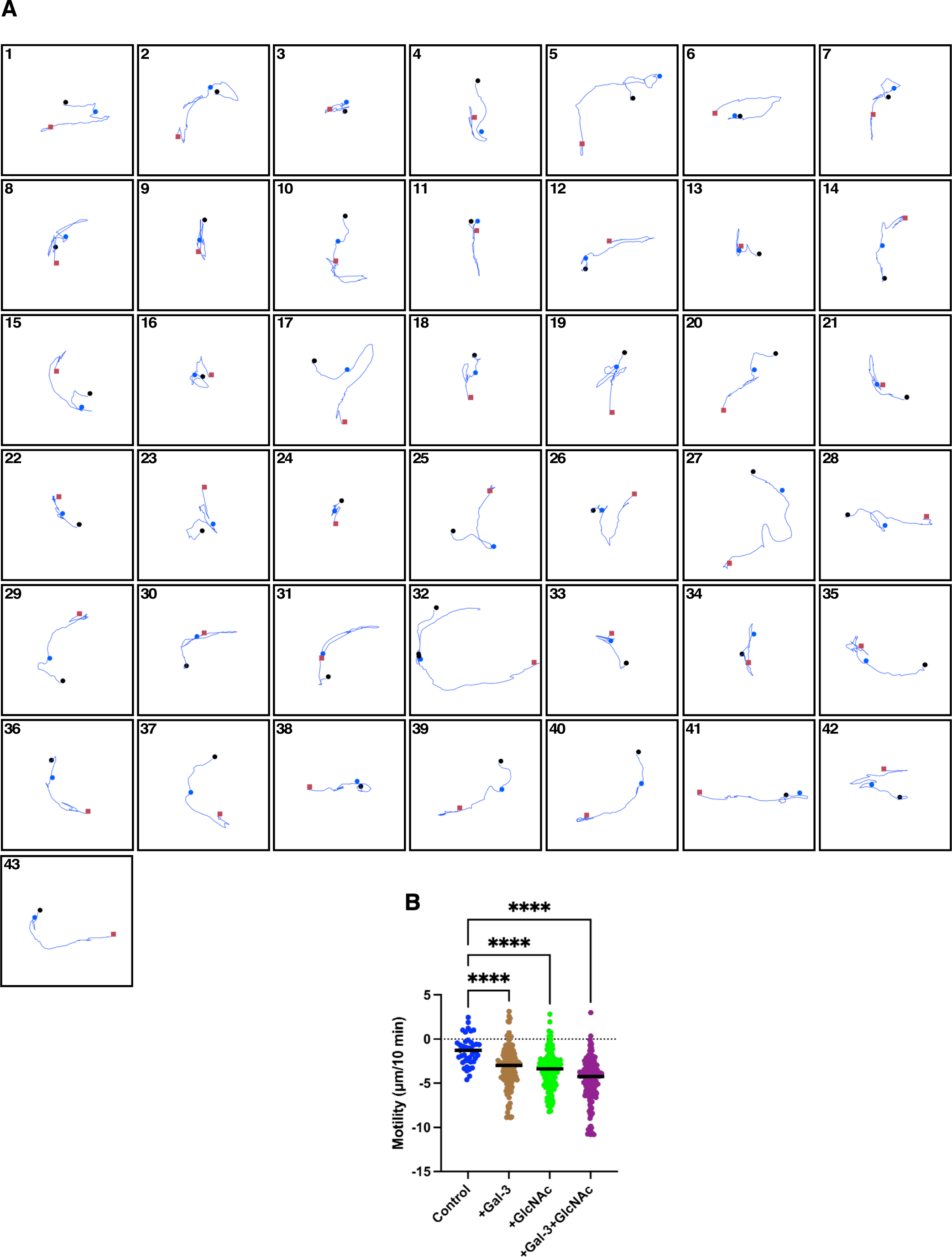
Trajectory Analysis of Individual Myoblasts. **A.** For the largest myotube identified by single-cell tracking during the GlcNAc treatment, the trajectories of 43 myoblasts that contributed to the formation of the myotube are depicted. The initial position at 0 minutes is marked with a black circle, while the fusion points are indicated by brown squares. The positions of myoblasts at 350 min, which coincide with the beginning of coordinated flow and alignment along the eventual shape of the myotube, are marked with blue circles. **B.** The motility of the myoblasts forming the myotube was calculated and compared to the speed of the coordinated flow (from Fig. 7). Differences in motility relative to the flow speed were determined. Statistical analysis was conducted using one-way ANOVA (Tukey’s test), with significance indicated by ****P < 0.0001. The sample sizes for Control, +Gal-3, +GlcNAc, and +Gal-3+GlcNAc were 47, 141, 145, and 177, respectively.

**Fig. 10.**
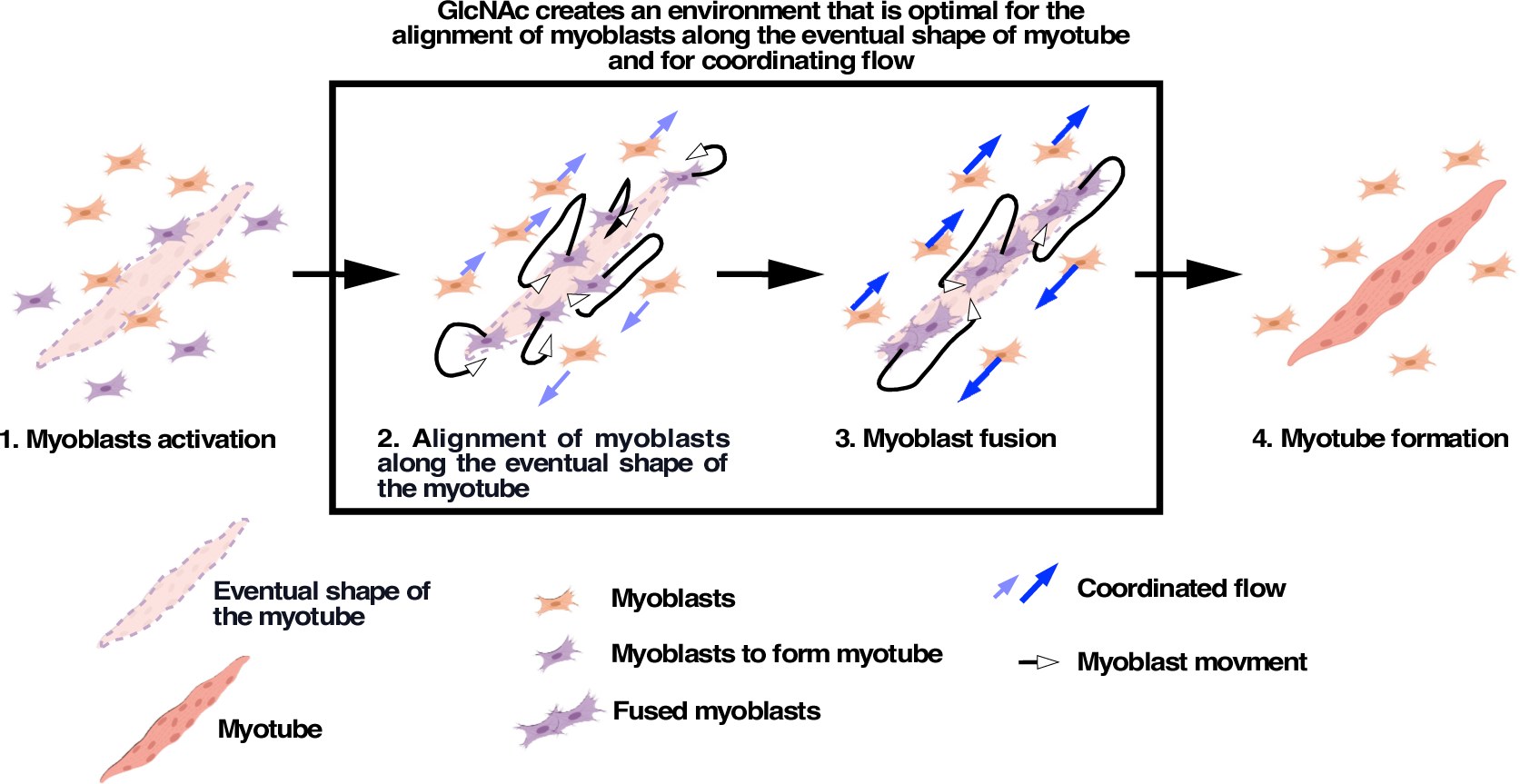
A Model for the Impact of GlcNAc and Gal-3 on Myogenesis. **[1]** Myoblasts are activated and become competent to form myotubes. **[2]** Among these myoblasts, some align along the eventual shape of the myotube (illustrated by a dotted outline), while others form coordinated flows along this shape, with aligned myoblasts still maintaining their movement. **[3]** Myoblasts involved in myotube formation move along the flow, pause, or move against the flow. These processes contribute to forming a mature myotube with correct alignment, resulting in the fusion of myoblasts. **[4]** Fused myoblasts eventually form a myotube. Gal-3 and GlcNAc create an environment optimal for myoblast alignment and coordinated flow.

## Discussion

In this study, we demonstrated that GlcNAc promotes myotube formation by increasing the fusion of primary mouse myoblasts in a dose-dependent manner. A decrease in the maturation of N-glycans by kifunensin diminished myogenesis, suggesting that lactosamine-rich N-glycans, whose synthesis is enhanced by GlcNAc, are essential for creating the optimal environment for myogenesis. It is well established in various biological contexts that the biological impact of GlcNAc is associated with the recognition of lactosamine-rich N-glycans by a glycan-binding lectin, galectin-3 (18–24). In this context, galectin-3 is recognized as a major oligosaccharide binding protein in differentiating myoblast (25), and our results reveal that myoblasts lacking galectin-3 expression exhibit impairment in myogenesis. Notably, supplementation of GlcNAc and galectin-3 improved the myogenesis of myoblast from Gal-3KO mice, demonstrating that the interaction between lactosamine-rich N-glycan and Gal-3 is critical for efficient myogenesis.

The interactions between myoblasts and myoblast-matrix interactions could influence the motility of myoblasts, thereby facilitating the myogenesis process. Gal-3 has the capacity to regulate cell-cell and cell-matrix interaction through its binding to lactosamine-rich oligosaccharides presented on the cell surface and cell-matrix proteins, laminin, and fibronectin (18–24, 26–30, 42, 43). Indeed, the motility of wild-type myoblasts was increased by Gal-3, GlcNAc, and both. In contrast, Gal-3KO myoblasts already exhibited higher motility than wild-type myoblasts, possibly due to the lack of Gal-3-mediated interactions, while the increased motility was reduced by treating Gal-3KO myoblasts with Gal-3 together with GlcNAc, suggesting that interactions mediated by Gal-3 do not simply increase or decrease the motility of myoblasts but rather optimize motility for myotube formation. Long-time single-cell tracking also showed that this motility does not result in random movement of myoblasts but instead creates coordinated flow, meaning a movement of myoblasts in a specific direction. We found that flows were formed during the process of myotube formation, and Gal-3 and GlcNAc promoted a faster flow relative to wild-type control myoblasts. Our findings reveal that the myotube is formed along the flow, suggesting the occurrence of the flow is an important process in myotube formation. However, despite the impaired efficiency in myogenesis exhibited by Gal-3KO myoblasts, they can still form even faster flows compared to wild-type myoblasts, suggesting that flow formation is likely related to the intrinsic nature of myoblasts themselves and that flow formation is not the only factor affecting the efficacy of myotube formation.

One possible scenario for the optimal speed of flow formation is that it maximizes the likelihood of encountering myoblasts with each other, allowing them to fuse. In this case, as encounters could stochastically occur, the formation of myotube might seem to occur in a random manner. However, single-cell tracking analysis reveals that myoblasts destined to form a myotube were already aligned along the eventual shape of the myotube even before a clear flow was established, indicating that the myotube is not formed in a stochastic manner. The process of myotube formation is illustrated in Fig. 11. In [1], myoblasts are activated to initiate the process of myogenesis. In [2], some of the myoblasts are aligned along the eventual shape of the myotube (light brown shape) in the early stage of the coordinated flow formation. In [3], a coordinated flow is generated, and while it might be expected that the aligned myoblasts to simply move along the flow, we observed that the aligned myoblasts not only moved along the flow, but also paused, or even opposed the flow as if using its force to shape themselves in preparation for fusion, ultimately leading to the formation of fused myoblasts. In [4], a myotube is formed. In this context, the interaction of Gal-3 with oligosaccharides plays a role in generating an environment conducive to alignment and the formation of an optimal flow speed, and GlcNAc contributes to the optimization of the biosynthesis of lactosamine-rich N-linked oligosaccharides, which provide binding sites for Gal-3. If such flow plays a critical role in the formation of tissue structure including muscles, GlcNAc could be the first reported natural factor involved in this process by influencing flow formation.

The mechanism of flow creation remains unclear. We utilized an ECM-coated 8-well chamber plastic slide, which required several weeks of storage at 4°C after drying for optimal differentiation, as we accidentally found that immediate use did not yield efficient myotube formation. This suggests that microscopic surface variations, created during storage, may guide myoblasts to form flows along an uneven surface. However, since this is an *in vitro* setup, it is uncertain whether similar flow formation and myoblast alignment occur *in vivo*, though it is plausible that *in vitro* events might reflect some characteristics observed *in vivo*. It has been suggested that muscle fiber regeneration *in vivo* is initiated by the activation followed by the asymmetric division of muscle stem cells, producing myoblasts that become fusion-competent and form myotubes(4, 6). Additionally, recent reports suggest that muscle stem cells, residing between a damaged myofiber and its basal lamina, rapidly proliferate (with an 8-10 hour doubling time) through a symmetric division, filling the residual basement membrane tube of the necrotic myofiber with myoblasts (7, 8). Then, first, density-dependent myoblast-myoblast fusion that occurs at 3.5-4.5 DPI establishes myotubes, followed by myoblast-myotube fusion and myotube-myotube fusion, further enlarging the fiber. Long-term live cell imaging of *in vitro* myogenesis with primary myoblasts showed that *in vitro* myotube/myofiber formation reflects some of the above-mentioned *in vivo* observations; under the differential condition, myoblasts begin to align using a coordinated flow, possibly playing a role in aligning myoblasts to fuse in the direction guided by the flow. Time-lapse observation showed that similar to *in vivo*, the fusion starts with myoblast-myoblast, followed by myoblast-myotube, and further enlargement of the myofiber through myotube-myotube fusion. Previous 2D *in vitro* myogenesis assay systems used in many studies to identify various factors involved in myogenesis (9–13) are generally limited to the observation of initial myoblast-myoblast fusion since the myogenesis is required to be stopped before the formation of myofibers because immunofluorescence staining process artificially removes larger myofibers. Thus, a live cell-single cell lineage tracking system may provide an alternative accessible approach for screening various factors to improve myogenesis.

In 2D *in vitro* culture, it is generally understood that myoblast movements are random (7). As shown in Fig. 9, when focusing solely on the movement of individual cells, the behavior appears random. However, when observed as a population, myoblasts undergoing differentiation exhibit coordinated flow from the early stages. Moreover, when tracking the movement of myoblasts that contribute to muscle fiber formation, these cells tend to migrate only in the vicinity where muscle fibers will eventually be formed. Occasionally, if a cell moves away from this region, it can be observed reversing direction, moving against the overall flow to return to the vicinity of fiber formation. Therefore, the comprehensive observation and analysis provided by long-term single-cell tracking have revealed that myoblasts committed to differentiation do not move randomly, even in a 2D *in vitro* culture system. In other words, what appears to be random movement at first glance is actually governed by a kind of grand design, whereby cells become aligned for fiber formation. This suggests that the collective movement of myoblasts contributing to muscle fibers is coordinated, and when this coordinated movement flow occurs at an optimal speed, muscle fibers are efficiently formed. *In vivo*, bi-directional movements have been observed, and it is possible that such movements also adhere to a grand design. In any case, this study suggests that the interaction between lactosamine-rich N-glycan and galectin-3 contributes to establishing the optimal speed of this coordinated flow to facilitate muscle fiber formation.

Given the critical need for rapid muscle repair in mammals for survival, it is plausible that the mechanisms for rapid muscle fiber regeneration have evolved. In this study, pre-distributing myoblasts along the shape of damaged muscle fibers may be essential to form proper myotubes and myofibers, although it is not yet known how specific myoblasts are aligned along the eventual shape of the myotube. Because these myoblasts can oppose the coordinated flow, their attachment to the matrix is likely tighter than that of other myoblasts. Previously, we reported that a subpopulation of pancreatic cancer cells expresses oligosaccharides with a higher content of α2-6 sialic acid structure, which is related to stemness, conferring resistance to the cancer cells against cell death and abnormal cellular events, including multipolar cell division (39). Such heterogeneity of cell surface glycosylation at a single cell level can be observed in any type of cell population; therefore, myoblast alignment may also be related to the expression levels of lactosamine-rich N-linked oligosaccharides on individual myoblasts. Thus, to further gain insight into the process of myogenesis, an individual myoblast level of investigation needs to be conducted.

In this study, we employed live-cell imaging and the single-cell tracking technique that we developed (38–40, 43). Because the fusion of myoblasts and myotube formation occurred within a short timeframe and fusion began within 3 to 5 days, continuous monitoring of myoblasts was essential to capture the moment of myoblast fusion. Additionally, single-cell tracking allows for precise quantitative analyses of the process. We believe that this technique will be useful for the study of myogenesis, and the software used for this study is available on GitHub (DOI 10.5281/zenodo.12988726).

## Materials and Methods

### Materials

GlcNAc (Ultimate glucosamine which is 100% USP grade N-acetylglucosamine) was procured from Wellesley Therapeutics, Ontario, Canada. Kifunensine and lectins (L-PHA amd Con A) were purchased from Sigma-Aldrich and Vector Laboratories, respectively. Anti-myosin heavy chain mouse monoclonal antibody (anti-MHC antibody; MF-20) and anti-mouse secondary antibody conjugated with Alexa 488 were obtained from the Developmental Studies Hybridoma Bank and Molecular Probes, respectively. Gal-3 and an anti-Gal-3 mouse monoclonal antibody (Mac-2) were purified as previously described (15, 27, 43).

### Isolation of Primary Mouse Myoblasts

Myoblasts were isolated from the hindlimb muscles of 2-month-old mice using the method described by Hindi et al. (44), with modifications. The muscles were finely dissected with scissors, minced, and enzymatically digested with collagenase II (3.5 mg/ml; Sigma-Aldrich) in Ham’s F-10 media containing 10% horse serum (Sigma-Aldrich), 1x penicillin/streptomycin, and 5 mM CaCl_2_ for 60 minutes at 37 °C. The enzymatic treatment was repeated with 0.43 mg/ml of collagenase II and subsequently with 2 mg/ml of collagenase II, each in the presence of 2.5 mM CaCl_2_ for another 60 minutes at 37 °C. The digested muscle was then passed through a 20 G needle, followed by a 40-μm cell strainer, and centrifuged to collect the cell pellet, which was resuspended in growth medium (GM) composed of Ham’s F-10, 10 ng/ml basic fibroblast growth factor (bFGF), 20% fetal bovine serum (FBS), and penicillin/streptomycin.

The harvested cells (5 x 10^6^ cells per dish) were incubated for 48 hours at 37 °C on a plate coated with extracellular matrix gel (ECM, Sigma-Aldrich) diluted 25-fold (approximately 400 µg/ml). The adherent cells were then trypsinized and plated on a gelatin-coated plate for 45 minutes at 37 °C. Detached cells were replated on an ECM-coated plate and cultured for a few more days. Myogenic progenitors/myoblasts were purified by consecutively replating on gelatin-coated plates until myoblast purity exceeded 99%. The myoblasts were maintained in GM on ECM-coated plates, and used up to 10 passages after establishment.

### Concurrent Long-Term Live Cell Imaging of Myoblasts

Primary mouse myoblasts were propagated in the GM within a humidified atmosphere containing 5% CO_2_. For imaging, 50 µL of cell suspension, containing approximately 65,000 myoblasts, was seeded into the center of each well of a µ-Slide 8 (ibidi), which was coated with ECM (25-fold dilution) and kept at 4°C for at least two weeks before use. The cells were allowed to adhere to the surface of µ-Slide 8 and after adding 500µl of GM, the cells were cultured for 24 hours. For concurrent long-term live cell imaging, the GM was replaced with a differentiation medium (DM) composed of DMEM high glucose, 2% horse serum, and penicillin-streptomycin. Following the medium change, the chamber slide was incubated in a CO_2_ incubator for 2 hours before being transferred to the microscope stage of an Olympus IX81 system. Live cell imaging was performed as previously described (38–40), employing near-infrared DIC imaging with a ×20 dry objective (UPlanSApo, 20×/0.75 NA, α/0.17/FN2G.5) coupled with a ×1.5 coupler (Quorum Technologies) to achieve a ×30 magnification equivalent. Images were captured in a 5 × 5 area scanning format, with each frame measuring 512 × 512 pixels (1.77 mm²) using the multi-dimensional acquisition mode in MetaMorph software (Quorum Technology Inc., WaveFX, v7.8.12.0), with an exposure time of 34 milliseconds and controlled via an XY piezo stage. Typically, 30 z-planes were acquired at 1 µm intervals, generating 25 multi-layered TIFF files of 512 × 512 pixels per well. These TIFF files were first selected in the focal plane and stitched to generate live cell imaging videos using in-house software as previously published (38–40). The cells were maintained on the microscope stage within an environmental chamber (Live Cell Instrument, Korea) under a humidified atmosphere with 7.5% CO_2_, and images were captured every 10 min. Concurrently, eight live cell imaging videos were generated in real-time using custom in-house software, which has been made available on GitHub (DOI 10.5281/zenodo.12988726). This setup allowed for continuous monitoring of myotube formation, and imaging continued until the formation of myotubes adequate for single-cell tracking analysis was observed.

### DIC Image Segmentation and Single-Cell Tracking

Grayscale DIC image segmentation and single-cell tracking were performed using custom in-house software available on GitHub (DOI 10.5281/zenodo.12988726). Each time point in the live cell imaging videos underwent grayscale image segmentation using a thresholding approach to define cell boundaries. This allowed for the identification of areas representing myoblasts, to which unique identifiers (cell numbers) were assigned. For objective analysis, 100 wild-type myoblasts and 50 Gal-3KO myoblasts were selected from the central areas of the images without adding any bias. Although the software features an automatic tracking function, myoblast fusion detection proved challenging and required manual tracking. A database was created to record the position of each myoblast at every time point, alongside events occurring in each cell. This database was utilized to generate maps depicting the fusion process of myoblasts and supported other bioinformatics analyses such as motility analysis and trajectory mapping.

### Western Blotting Using L-PHA, Con A and Anti-Gal-3 Mouse Monoclonal Antibody (Mac-2)

Myoblasts were plated at a density of 5.25 x 10^4^ cells/cm^2^ in 12-well or 24-well plates and incubated overnight in the GM. Differentiation was initiated by switching the medium to the DM. After 24 and 48 hours of differentiation, the culture medium was collected to analyze extracellular Gal-3 levels. Alternatively, cells were washed twice with calcium and magnesium-free PBS and lysed in 25 mM Tris-buffered saline containing 1% Triton X-100 and a protease inhibitor cocktail (Sigma-Aldrich) on ice for 15 minutes. The lysates were centrifuged at 10,000 rpm for 15 minutes to extract cellular proteins. Proteins from both the medium and lysates were subjected to SDS-PAGE (12% acrylamide for Gal-3 and 7.5% for L-PHA and Con A analyses and transferred on a PVDF membrane (Cyiva) for Western blot analysis. The filters were blocked with 5 % skimmed milk in 25 mM Tris-buffered saline with 0.02% Tween (for anti-Gal-3 antibody) or Chem Blot Blocking Buffer (for lectins, Azure Biosystems) and then incubated with 2 μg/ml biotinylated anti-Gal-3 antibody, 2 μg/ml L-PHA-biotin or Con A-biotin, followed by streptavidin-peroxidase (Biosynth). The blots were visualized on a digital imager (Vilber Fusion Fx) to quantify the levels of Gal-3, lactosamine-rich N-glycans which are detected with L-PHA and high-mannose-rich N-glycans with Con A. Band quantification was performed using ImageJ.

### Indirect Immunofluorescence

Myoblasts or myotubes were fixed with 3.7% paraformaldehyde for 15 minutes at room temperature, followed by three washes with PBS. Cells were then blocked with a Carbo-Free Blocking Solution (Vector Laboratories) for 1 hour at room temperature. After blocking, cells were incubated with MF-20 (anti-MHC antibody; diluted 1:5 in Carbo-Free Blocking Solution) for 60 minutes at room temperature. Following the primary antibody incubation, cells were washed three times with PBS and then incubated with an anti-mouse secondary antibody conjugated with Alexa 488 (diluted 1:1000) for 1 hour at room temperature. After the secondary antibody incubation, cells were washed three additional times with PBS, stained with DAPI, and washed three more times. Fluorescent imaging was performed with a 250-millisecond exposure using lasers with wavelengths of 403 nm for DAPI and 491 nm for Alexa Fluor 488.

### Analysis of Monosaccharide Metabolites in Myoblasts

Nucleotide sugars from myoblasts cultured in DM with or without GlcNAc for 24 hours were prepared and quantified by hydrophilic interaction chromatography-electrospray ionization-tandem mass spectrometry as reported previously (45, 46). Myoblasts (5.25 x 10^4^ cells/cm^2^) in 12-well were collected in ice-cold 70% ethanol (1.5 ml), and the supernatants after centrifugation were lyophilized. The freeze-dried samples were dissolved in 1 ml of 10 mM NH_4_HCO_3_ and purified on an Envi-Carb column as reported previously (45, 46). For measuring the protein concentrations, the precipitate of 70% ethanol extraction was dissolved in 2% SDS, followed by quantification of protein concentration using Pierce BCA Protein Assay Kit (#23227). Liquid chromatography-mass spectrometry (LC-MS) analysis was performed on an ion mobility time-of-flight mass spectrometer Synapt XS (Waters, MA) coupled with the Acquity UPLC system (Waters, MA). Chromatography was performed on an Acquity UPLC BEH-amido column (2.1 mm i.d. x 150 mm, 1.7 µm; Waters) (45, 47) The mobile phases were as follows: A, 20 mM ammonium acetate containing 90 % acetonitrile: B, 20 mM ammonium acetate. Portions of cellular extracts were analyzed by MS/MS analysis. For quantification, we calculated the peak area of extracted ion chromatograms of the major product ions. MS/MS transition for nucleotide sugars was defined as follows: UDP-HexNAc, *m/z* 606.1, UDP-Hex, *m/z* 565.1, CMP-NeuAc, *m/z* 613.1, UDP-GlcA, *m/z* 579.1, GDP-Glc and GDP-Man, *m/z* 604.1, and GDP-Fuc, *m/z* 588.1. The nucleotide sugar levels were normalized as pmol/mg proteins.

## Statistical Analysis

Statistical analyses were performed using Prism 10.

## Data availability

All data are included in the article and/or supporting information. The software was deposited to GitHub; DOI 10.5281/zenodo.12988726. Any questions regarding data availability can be directed to the corresponding author.

## Supporting information

This article contains supporting information.

## Acknowledgments

We thank Willem Wassenaar (Wellesley Therapeutics, Ontario) for providing GlcNAc, and the Bioimaging Platform of the Research Centre for Infectious Diseases, Axe of Infectious and Inflammatory Diseases at the CHU de Quebec Research Centre for their invaluable technical support with the microscopes. This research received financial support from the Canadian Foundation for Innovation, the Canadian Institutes for Health Research, GlycoNET, the Network of Centers of Excellence of Canada, Defeat Duchenne Canada, and the joint research program of the J-GlycoNet cooperative network, which is accredited by the Minister of Education, Culture, Sports, Science and Technology, MEXT, Japan, as a Joint Usage/Research Center as well as JSPS Core-toCore Program No. JPJSCCA202000007.

## Author contributions

M.S.S. and S.S. designed the study, conducted experiments, and contributed to manuscript writing. G.S.-P., E.B., and M.F. carried out experiments related to Western blotting, while K.H. and K.N. focused on mass spectrometry experiments. A.R. conducted experiments, analyzed single-cell tracking data, and contributed to manuscript editing. All authors have reviewed and approved the final version of the manuscript.

## Legends for Supplementary Figures, Data and Videos

**Supplementary Fig. 1. Impact of kifunensin on glycosylation of differentiating myoblasts**

Primary wild-type myoblasts were cultured in differentiation mediums with or without 5 mM GlcNAc and/or 40 μM kifunensine for 24 hours. Myoblasts were then harvested and analyzed by Western blotting using Con A, which binds to high-mannose N-linked oligosaccharides. Staining with Con A is shown.

**Supplementary Fig. 2. Effect of GlcNAc on Galectin-3 release**

**A.** Primary wild-type myoblasts were cultured in a differentiation medium with or without 5 mM GlcNAc for 24 and 48 hours. Gal-3 released into the medium was quantified by Western blotting using an anti-Gal-3 antibody, Mac-2. The quantity of Gal-3 (arbitrary units) at 24 hours (**B**) and 48 hours (**C**) is plotted. Data represent means ± standard errors from three samples, with no statistically significant differences observed.

**Supplementary Data 1. Maps depicting the process of myoblast fusion in non-treated wild-type mouse myoblasts used in the GlcNAc dose-response experiments (Control)**

Maps were generated using data from the single-cell tracking of control myoblasts for the GlcNAc dose-response experiments. In these maps, the following symbols are used to represent specific events: Light blue circle: Mitosis, Pink square: Cell death, Orange circle and vertical line: Cell fusion, Black vertical line: Bipolar cell division, and Gray square: Movement out of the field of view.

**Supplementary Data 2. Maps depicting the process of myoblast fusion in wild-type mouse myoblasts treated with 1 mM GlcNAc (GlcNAc 1)**

Maps were generated using data from the single-cell tracking of myoblasts treated with 1 mM GlcNAc. In these maps, the following symbols are used to represent specific events: Light blue circle: Mitosis, Pink square: Cell death, Orange circle and vertical line: Cell fusion, Black vertical line: Bipolar cell division, and Gray square: Movement out of the field of view.

**Supplementary Data 3. Maps depicting the process of myoblast fusion in wild-type mouse myoblasts treated with 5 mM GlcNAc (GlcNAc 5)**

Maps were generated using data from the single-cell tracking of myoblasts treated with 5 mM GlcNAc. In these maps, the following symbols are used to represent specific events: Light blue circle: Mitosis, Pink square: Cell death, Orange circle and vertical line: Cell fusion, Black vertical line: Bipolar cell division, and Gray square: Movement out of the field of view.

**Supplementary Data 4. Maps depicting the process of myoblast fusion in wild-type mouse myoblasts treated with 10 mM GlcNAc (GlcNAc 10)**

Maps were generated using data from the single-cell tracking of myoblasts treated with 10 mM GlcNAc. In these maps, the following symbols are used to represent specific events: Light blue circle: Mitosis, Pink square: Cell death, Orange circle and vertical line: Cell fusion, Black vertical line: Bipolar cell division, and Gray square: Movement out of the field of view.

**Supplementary Data 5. Maps depicting the process of myoblast fusion in non-treated wild-type mouse myoblasts used in the Gal-3 and/or GlcNAc addition experiments (Control)**

Maps were generated using data from the single-cell tracking of control myoblasts for the Gal-3 and/or GlcNAc addition experiments. In these maps, the following symbols are used to represent specific events: Light blue circle: Mitosis, Pink square: Cell death, Orange circle and vertical line: Cell fusion, Black vertical line: Bipolar cell division, and Gray square: Movement out of the field of view.

**Supplementary Data 6. Maps depicting the process of myoblast fusion in wild-type mouse myoblasts treated with 2 μM Gal-3 (+Gal-3)**

Maps were generated using data from the single-cell tracking of myoblasts treated with 2 μM Gal-3. In these maps, the following symbols are used to represent specific events: Light blue circle: Mitosis, Pink square: Cell death, Orange circle and vertical line: Cell fusion, Black vertical line: Bipolar cell division, and Gray square: Movement out of the field of view.

**Supplementary Data 7. Maps depicting the process of myoblast fusion in wild-type mouse myoblasts treated with 10 mM GlcNAc (+GlcNAc)**

Maps were generated using data from the single-cell tracking of myoblasts treated with 10 mM GlcNAc. In these maps, the following symbols are used to represent specific events: Light blue circle: Mitosis, Pink square: Cell death, Orange circle and vertical line: Cell fusion, Black vertical line: Bipolar cell division, and Gray square: Movement out of the field of view.

**Supplementary Data 8. Maps depicting the process of myoblast fusion in wild-type mouse myoblasts treated with 2 μM Gal-3 and 10 mM GlcNAc (+Gal-3+GlcNAc)**

Maps were generated using data from the single-cell tracking of myoblasts treated with 2 μM Gal-3 and 10 mM GlcNAc. In these maps, the following symbols are used to represent specific events: Light blue circle: Mitosis, Pink square: Cell death, Orange circle and vertical line: Cell fusion, Black vertical line: Bipolar cell division, and Gray square: Movement out of the field of view.

**Supplementary Data 9. Maps depicting the process of myoblast fusion in non-treated Gal-3KO mouse myoblasts used in the Gal-3 and/or GlcNAc addition experiments (Control)**

Maps were generated using data from the single-cell tracking of non-treated Gal-3KO myoblasts for the Gal-3 and/or GlcNAc addition experiments. In these maps, the following symbols are used to represent specific events: Light blue circle: Mitosis, Pink square: Cell death, Orange circle and vertical line: Cell fusion, Black vertical line: Bipolar cell division, and Gray square: Movement out of the field of view.

**Supplementary Data 10. Maps depicting the process of myoblast fusion in Gal-3KO mouse myoblasts treated with 2 μM Gal-3 (+Gal-3)**

Maps were generated using data from the single-cell tracking of Gal-3KO myoblasts treated with 2 μM Gal-3. In these maps, the following symbols are used to represent specific events: Light blue circle: Mitosis, Pink square: Cell death, Orange circle and vertical line: Cell fusion, Black vertical line: Bipolar cell division, and Gray square: Movement out of the field of view.

**Supplementary Data 11. Maps depicting the process of myoblast fusion in Gal-3KO mouse myoblasts treated with 10 mM GlcNAc (+GlcNAc)**

Maps were generated using data from the single-cell tracking of Gal-3KO myoblasts treated with 10 mM GlcNAc. In these maps, the following symbols are used to represent specific events: Light blue circle: Mitosis, Pink square: Cell death, Orange circle and vertical line: Cell fusion, Black vertical line: Bipolar cell division, and Gray square: Movement out of the field of view.

**Supplementary Data 12. Maps depicting the process of myoblast fusion in Gal-3KO mouse myoblasts treated with 2 μM Gal-3 and 10 mM GlcNAc (+Gal-3+GlcNAc)**

Maps were generated using data from the single-cell tracking of Gal-3KO myoblasts treated with 2 μM Gal-3 and 10 mM GlcNAc. In these maps, the following symbols are used to represent specific events: Light blue circle: Mitosis, Pink square: Cell death, Orange circle and vertical line: Cell fusion, Black vertical line: Bipolar cell division, and Gray square: Movement out of the field of view.

**Supplementary Data 13. Analysis of Fusion Patterns**

The fusion patterns of wild-type myoblasts and myotubes were analyzed. The abbreviations used are as follows: B-B represents a fusion between myoblasts; B-T represents a fusion between a myoblast and a fused myoblast that has not yet formed a myotube or has formed a myotube; and T-T represents a fusion between myotubes. The number following each abbreviation (B-B, B-T, T-T) indicates the number of those specific fusion events. The numbers in the table represent the types of fusion events and the number of nuclei in the fused myoblast or myotube. For example, in the Control condition, Column A, 164 (2) indicates that at time point 164 (1,640 minutes from the start of single-cell tracking), two mononuclear myoblasts fused, resulting in the formation of a myoblast with two nuclei. Another example is Column A in the Control condition, where 127 (2) is followed by 194 (3). This indicates that at time point 194, a myoblast fused with a binuclear myoblast, resulting in the formation of a myoblast or myotube with three nuclei. Arrows indicate fusion events involving a myoblast or myotube with more than two nuclei. For instance, in the Control condition, Column H, Row 7, a myotube with 11 nuclei fused with a myoblast or myotube with 3 nuclei at time point 196, resulting in a myotube with 11 nuclei. “E” indicates that the myoblast or myotube reached the end of the single-cell tracking period.

**Supplementary Video 1. Myoblast-Myoblast Fusion**

This video showcases the fusion process between myoblasts, which are highlighted with orange circles. Scale bar: 25 µm.

**Supplementary Video 2. Myoblast-Myotube Fusion**

This video showcases the fusion process between a myoblast and a myotube, which are highlighted with orange circles. Scale bar: 25 µm.

**Supplementary Video 3. Myotube-Myotube Fusion**

This video showcases the fusion process between myotubes, which are highlighted with orange circles. Scale bar: 25 µm.

**Supplementary Video 4. Single-cell tracking of non-treated wild-type mouse myoblasts used in the GlcNAc dose-response experiments (Control)**

A video for single-cell tracking of non-treated wild type myoblasts used for GlcNAc dose-response experiments is shown (0–3690 min). A unique color was assigned to each cell lineage. The total number of tracked cells was 135. Scale bar: 50 µm.

**Supplementary Video 5. Single-cell tracking of wild-type mouse myoblasts treated with 1 mM GlcNAc dose-response experiments (GlcNAc 1)**

A video for single-cell tracking of wild type myoblasts treated with 1 mM GlcNAc is shown (0–3690 min). A unique color was assigned to each cell lineage. The total number of tracked cells was 149.

Scale bar: 50 µm.

**Supplementary Video 6. Single-cell tracking of wild-type mouse myoblasts treated with 5 mM GlcNAc dose-response experiments (GlcNAc 5)**

A video for single-cell tracking of wild type myoblasts treated with 5 mM GlcNAc is shown (0–3690 min). A unique color was assigned to each cell lineage. The total number of tracked cells was 191. Scale bar: 50 µm.

**Supplementary Video 7. Single-cell tracking of wild-type mouse myoblasts treated with 10 mM GlcNAc dose-response experiments (GlcNAc 10)**

A video for single-cell tracking of wild type myoblasts treated with 10 mM GlcNAc is shown (0–3690 min). A unique color was assigned to each cell lineage. The total number of tracked cells was 178.Scale bar: 50 µm.

**Supplementary Video 8. Single-cell tracking of non-treated wild-type mouse myoblasts used in the Gal-3 and/or GlcNAc addition experiments (Control)**

A video for single-cell tracking of non-treated wild type myoblasts used in the Gal-3 and GlcNAc addition experiments is shown (0–1990 min). A unique color was assigned to each cell lineage. The total number of tracked cells was 136. Scale bar: 50 µm.

**Supplementary Video 9. Single-cell tracking of wild-type mouse myoblasts treated with 2 μM Gal-3 (+Gal-3)**

A video for single-cell tracking of wild type myoblasts treated with 2 μM Gal-3 is shown (0–1990 min). A unique color was assigned to each cell lineage. The total number of tracked cells was 201. Scale bar: 50 µm.

**Supplementary Video 10. Single-cell tracking of wild-type mouse myoblasts treated with 10 mM GlcNAc (+GlcNAc)**

A video for single-cell tracking of wild type myoblasts treated with 10 mM GlcNAc is shown (0–1990 min). A unique color was assigned to each cell lineage. The total number of tracked cells was 237. Scale bar: 50 µm.

**Supplementary Video 11. Single-cell tracking of wild-type mouse myoblasts treated with 2 μM Gal-3 and 10 mM GlcNAc (+Gal-3+GlcNAc)**

A video for single-cell tracking of wild type myoblasts treated with 2 μM Gal-3 and 10 mM GlcNAc is shown (0–1990 min). A unique color was assigned to each cell lineage. The total number of tracked cells was 269. Scale bar: 50 µm.

**Supplementary Video 12. Single-cell tracking of non-treated Gal-3KO mouse myoblasts used in the Gal-3 and/or GlcNAc addition experiments (Control)**

A video for single-cell tracking of non-treated Gal-3KO myoblasts used in the Gal-3 and GlcNAc addition experiments is shown (0–1590 min). A unique color was assigned to each cell lineage. The total number of tracked cells was 133. Scale bar: 50 µm.

**Supplementary Video 13. Single-cell tracking of Gal-3KO mouse myoblasts treated with 2 μM Gal-3 (+Gal-3)**

A video for single-cell tracking of Gal-3KO myoblasts treated with 2 μM Gal-3 is shown (0–1590 min). A unique color was assigned to each cell lineage. The total number of tracked cells was 175. Scale bar: 50 µm.

**Supplementary Video 14. Single-cell tracking of Gal-3KO mouse myoblasts treated with 10 mM GlcNAc2 (+GlcNAc)**

A video for single-cell tracking of Gal-3KO myoblasts treated with 10 mM GlcNAc is shown (0–1590 min). A unique color was assigned to each cell lineage. The total number of tracked cells was 110. Scale bar: 50 µm.

**Supplementary Video 15. Single-cell tracking of Gal-3KO mouse myoblasts treated with 2 μM and 10 mM GlcNAc (+Gal-3+GlcNAc)**

A video for single-cell tracking of Gal-3KO myoblasts treated with 2 μM Gal-3 and 10 mM GlcNAc is shown (0–1590 min). A unique color was assigned to each cell lineage. The total number of tracked cells was 205. Scale bar: 50 µm.

**Supplementary Video 16. Process of Myotube Formation**

This video displays the formation of a myotube (0 to 1960 min). A myotube formed in the presence of 10 mM GlcNAc was selected for demonstration (refer to Fig. 9A). The myoblasts contributing to this myotube are outlined in green, while fused myoblasts or myotubes are outlined in yellow. Scale bar: 50 µm.

