## Supplementary figures and images for "N-Acetylglucosamine Facilitates Coordinated Myoblast Flow, Forming the Foundation for Efficient Myogenesis"

### Supplemental figures

**Sup. Fig. 1**

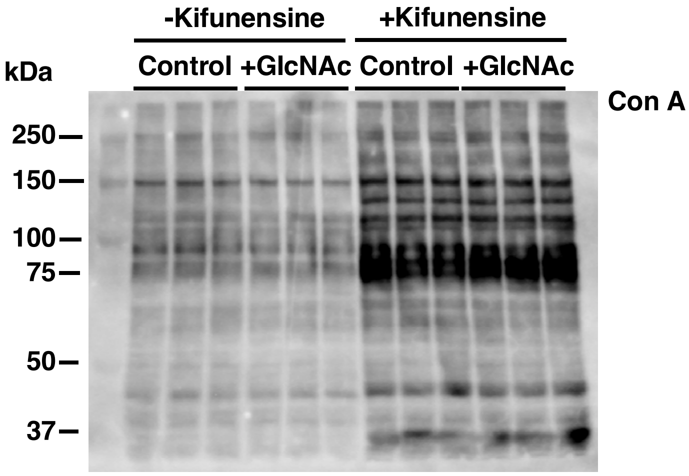


Sato et al.

**Sup.Fig. 2**

A

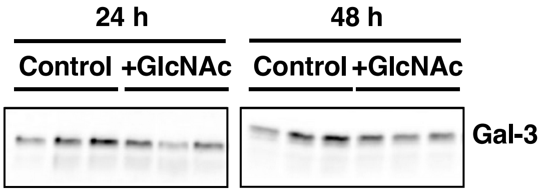


B

C

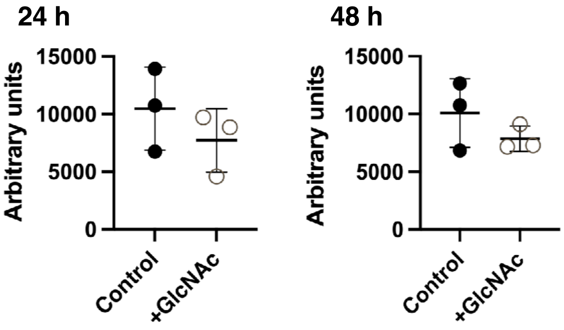


Sato et al.
