## Supplemental data 1 for "N-Acetylglucosamine Facilitates Coordinated Myoblast Flow, Forming the Foundation for Efficient Myogenesis"

Analysis: Wild type-Dose, Treat.: Control, Cell: Myoblasts

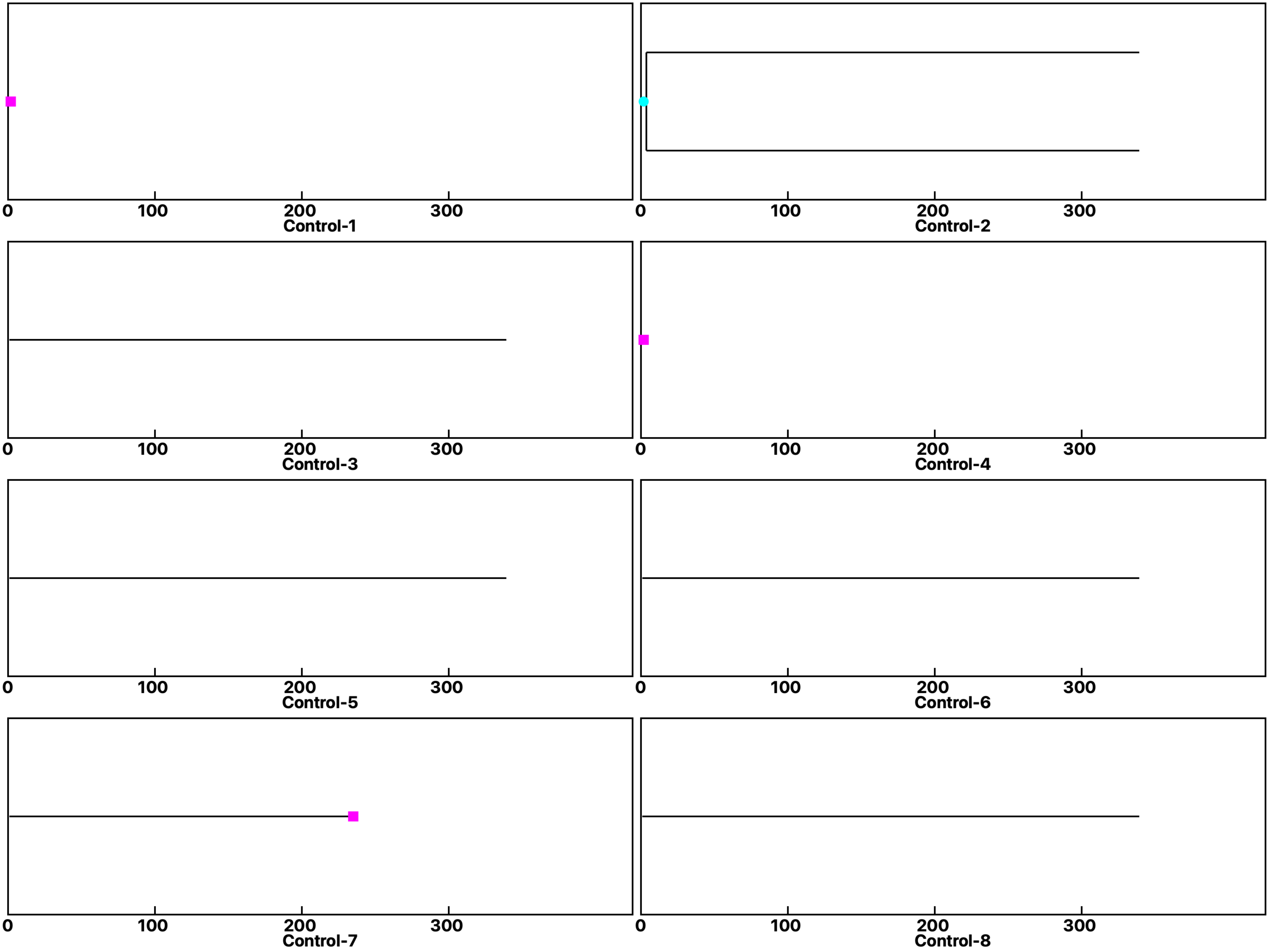

Analysis: Wild type-Dose, Treat.: Control, Cell: Myoblasts

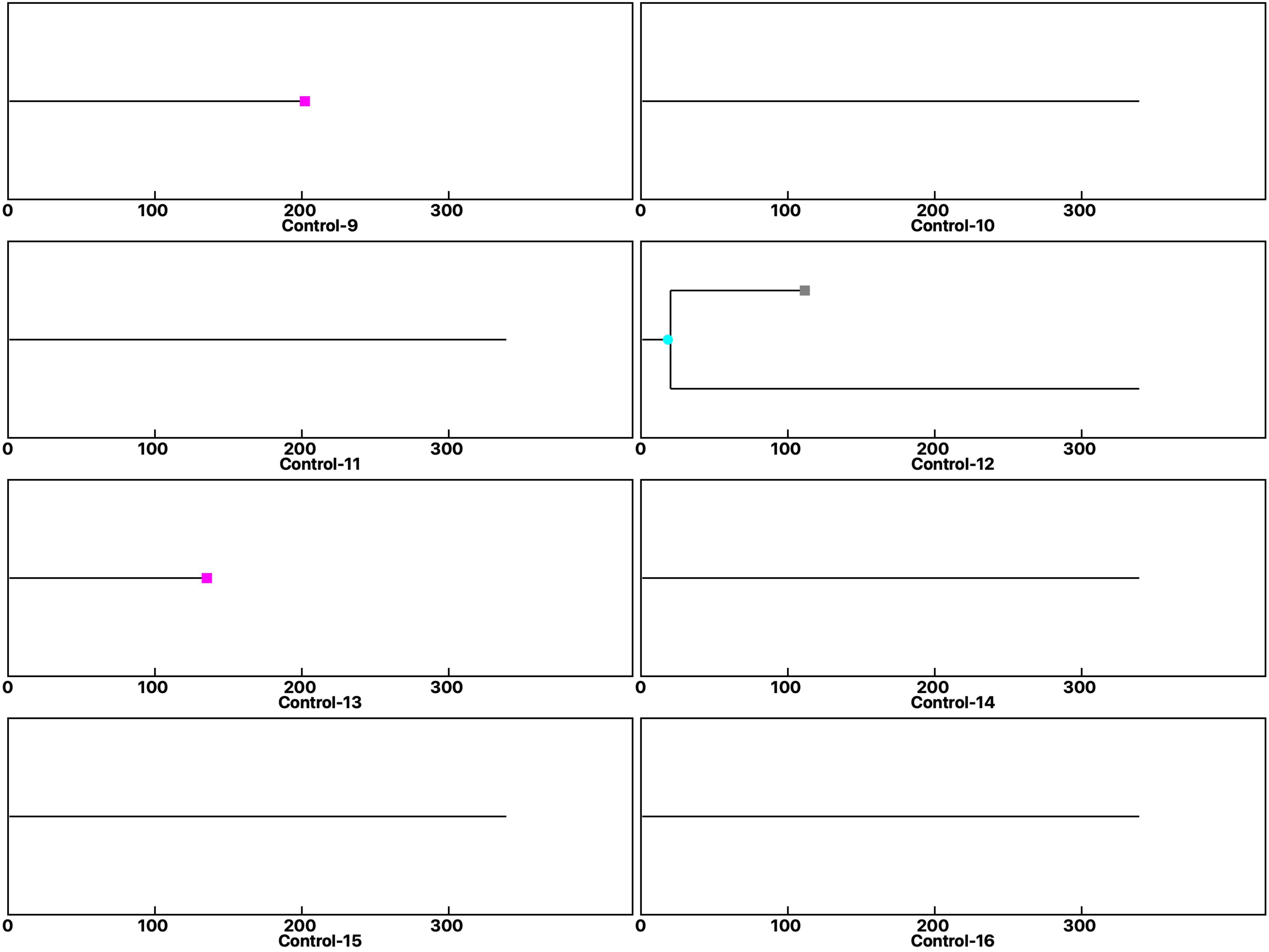

Analysis: Wild type-Dose, Treat.: Control, Cell: Myoblasts

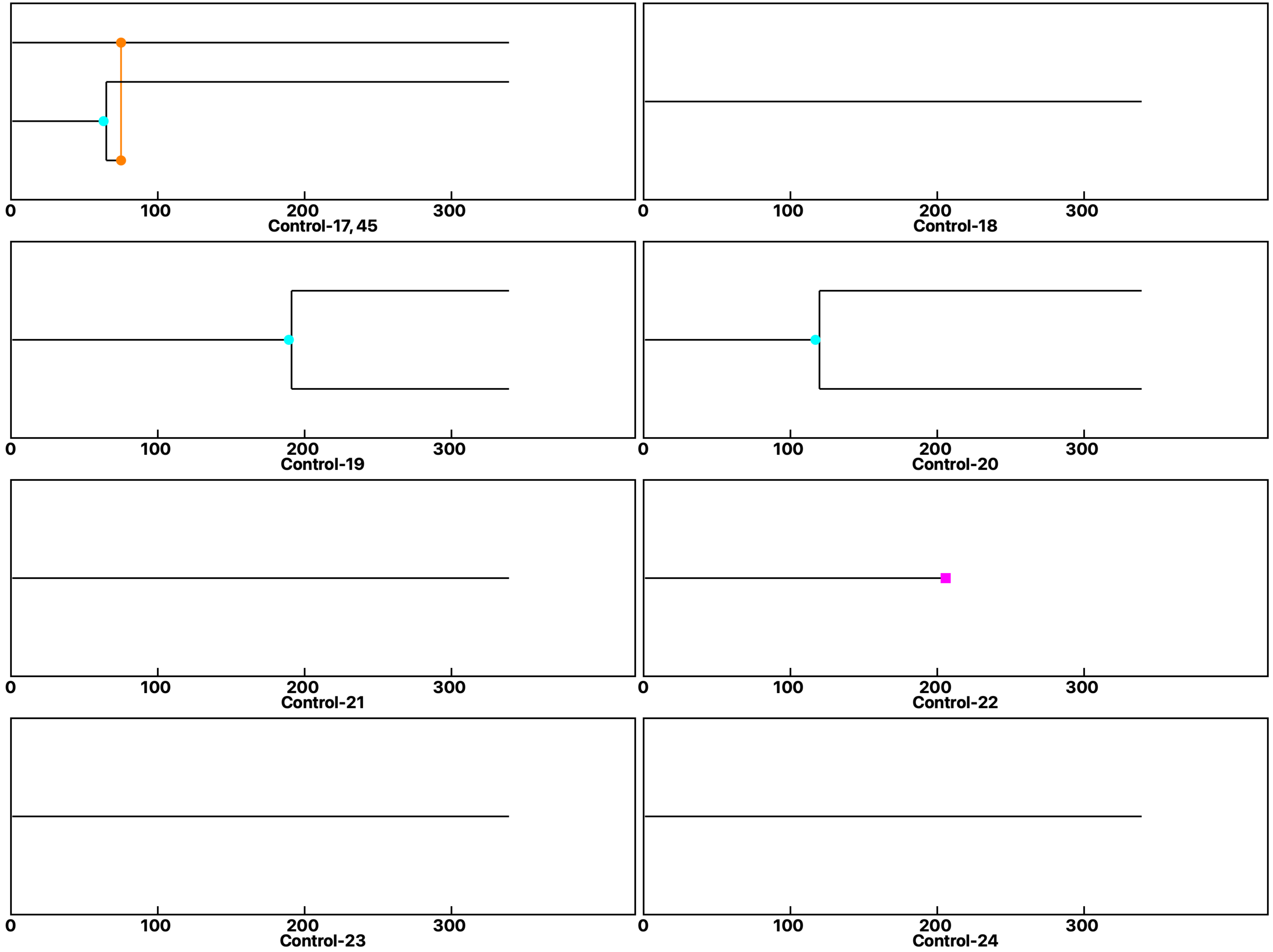

**Analysis: Wild type-Dose, Treat.: Control, Cell: Myoblasts**

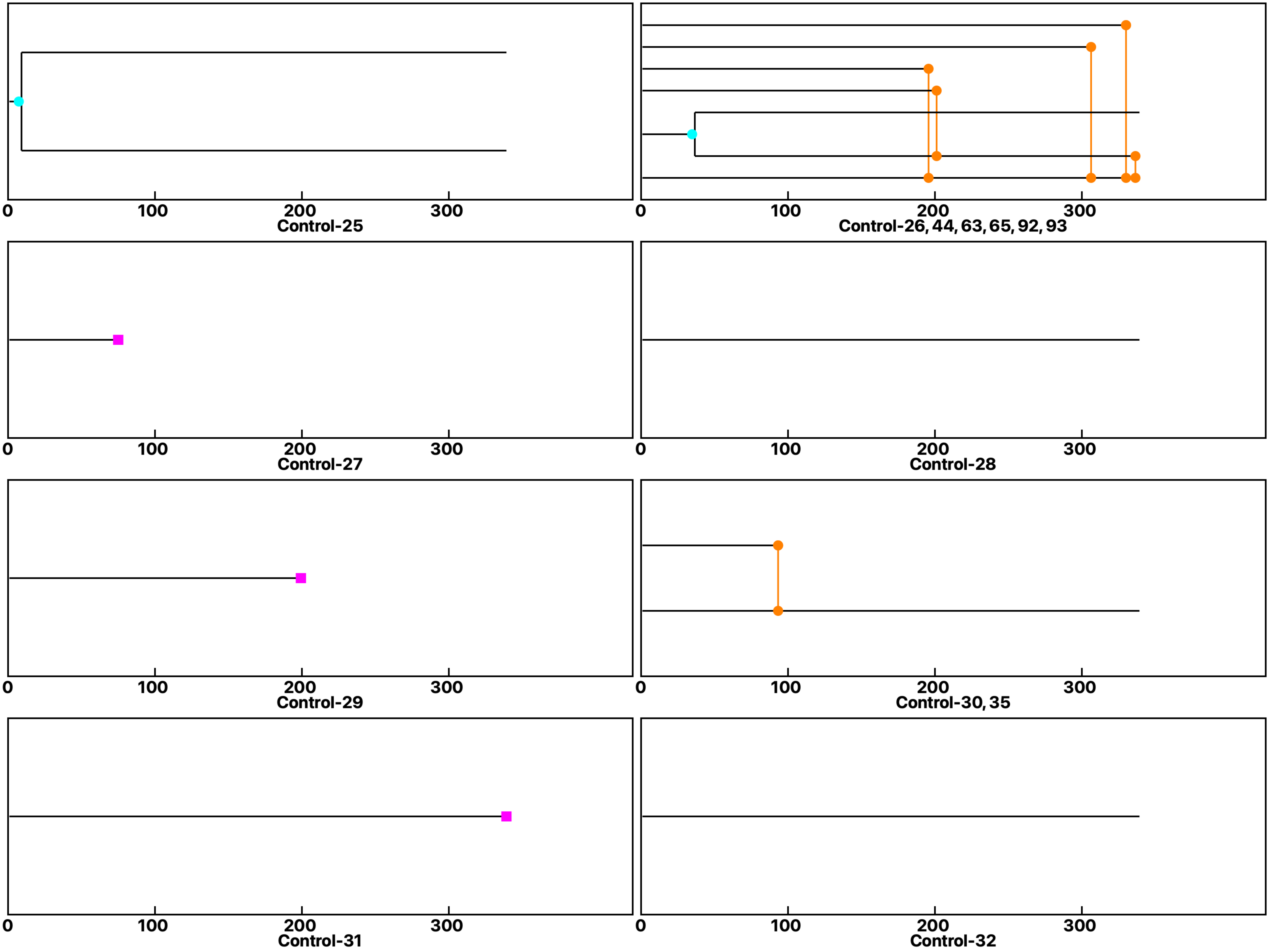

Analysis: Wild type-Dose, Treat.: Control, Cell: Myoblasts

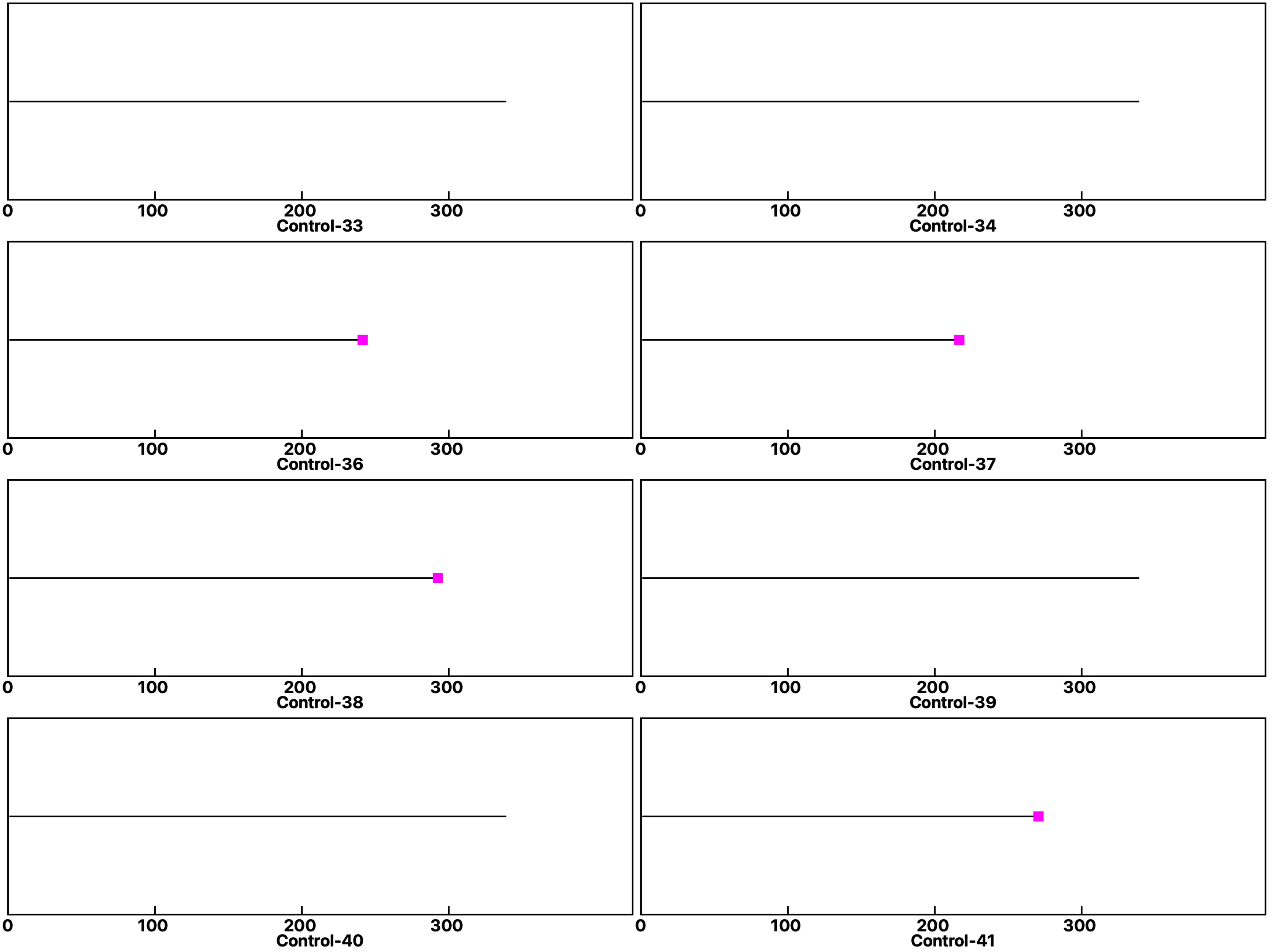

Analysis: Wild type-Dose, Treat.: Control, Cell: Myoblasts

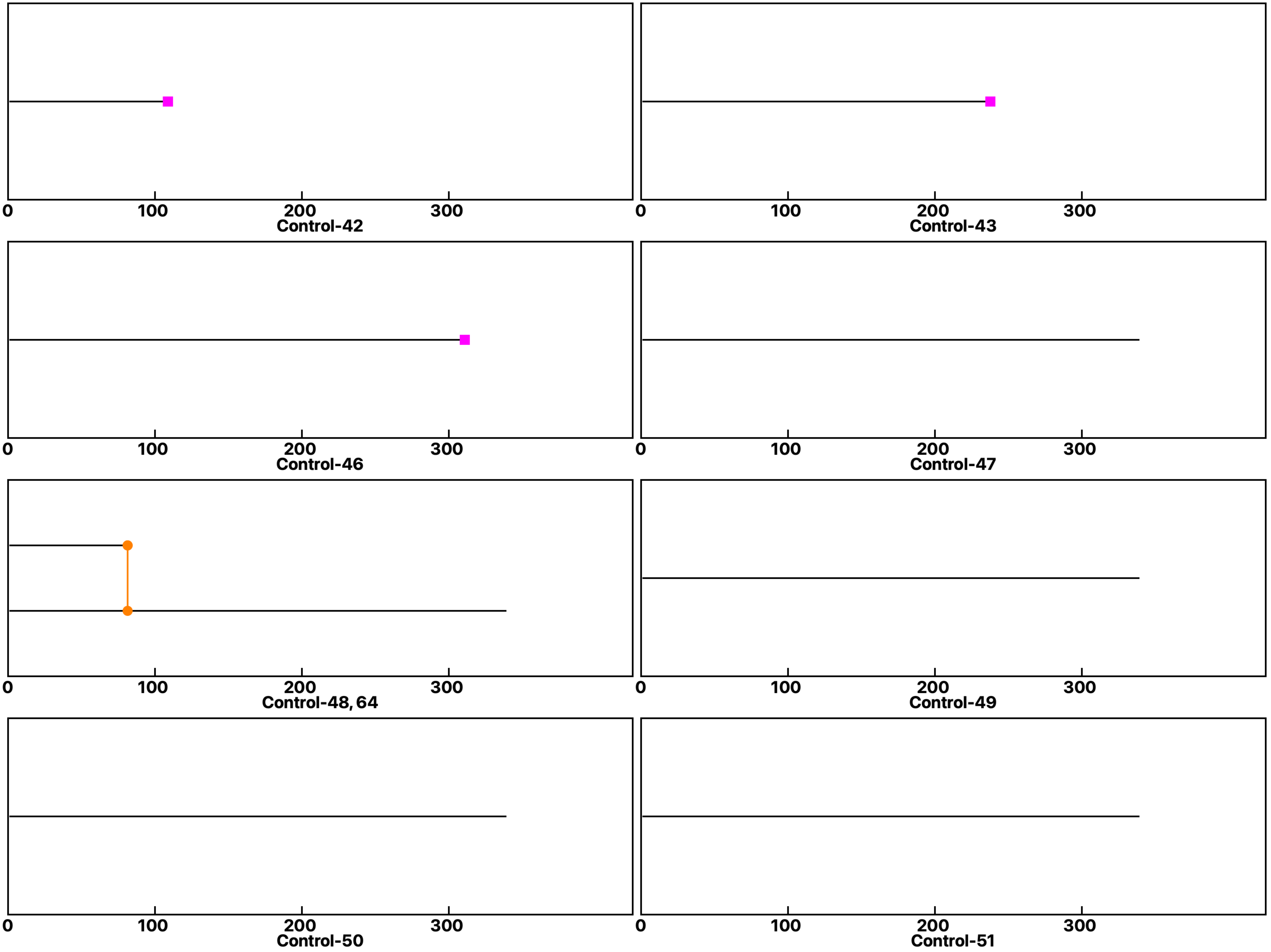

**Analysis: Wild type-Dose, Treat.: Control, Cell: Myoblasts**

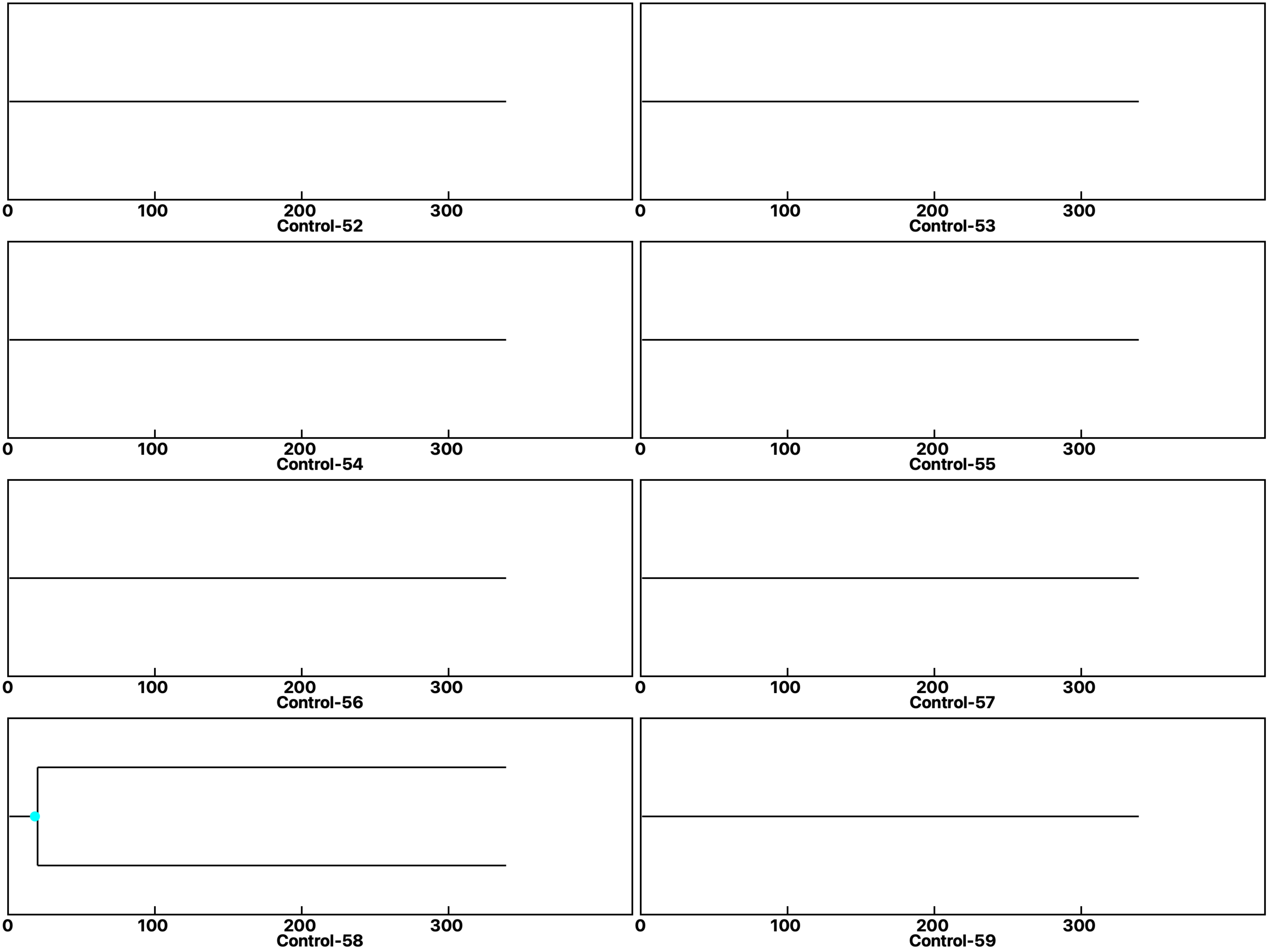

Analysis: Wild type-Dose, Treat.: Control, Cell: Myoblasts

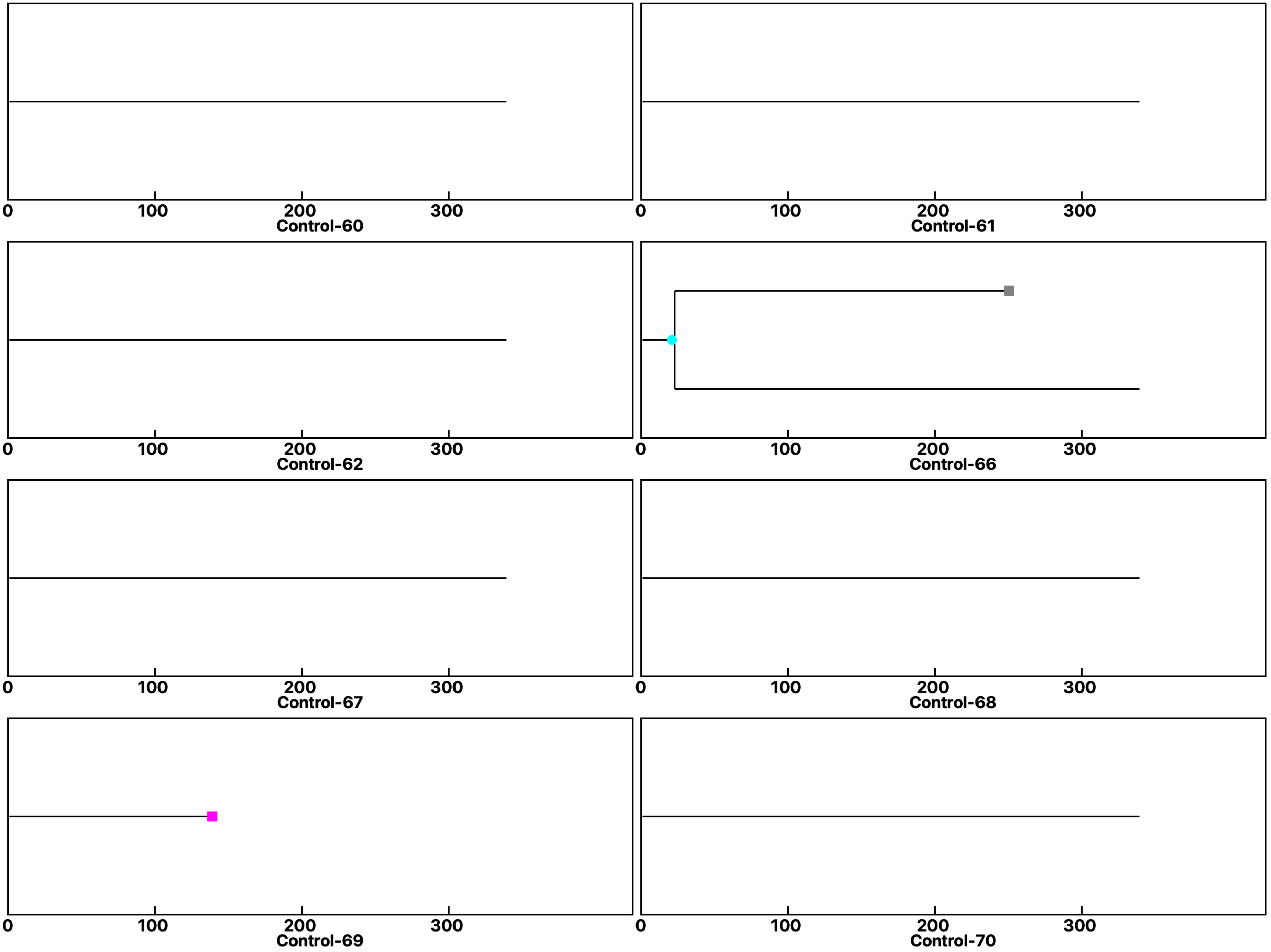

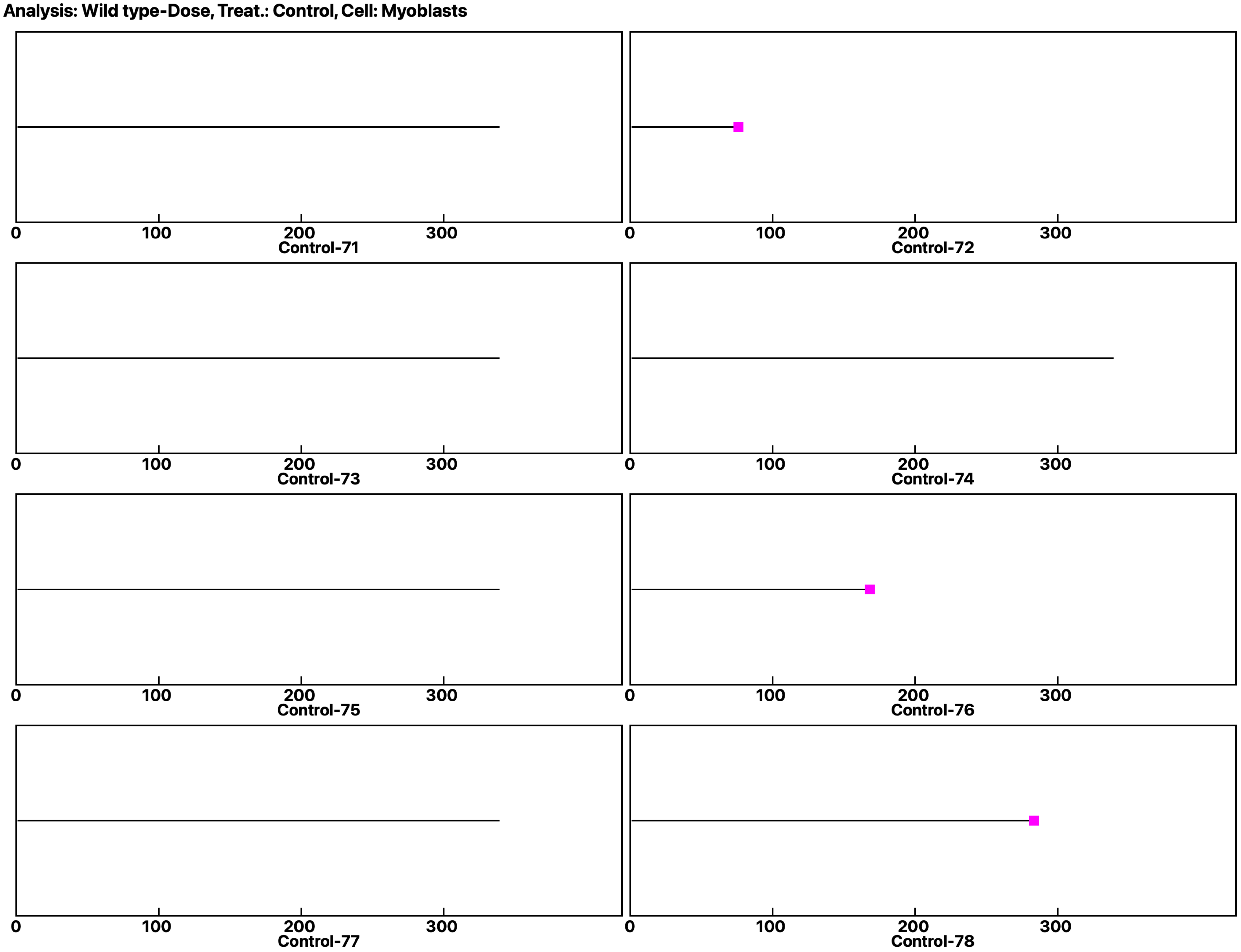
Analysis: Wild type-Dose, Treat.: Control, Cell: Myoblasts
0
100
200
300
Control-71
Control-72
Control-73
Control-74
Control-75
Control-76
Control-77
Control-78

Analysis: Wild type-Dose, Treat.: Control, Cell: Myoblasts

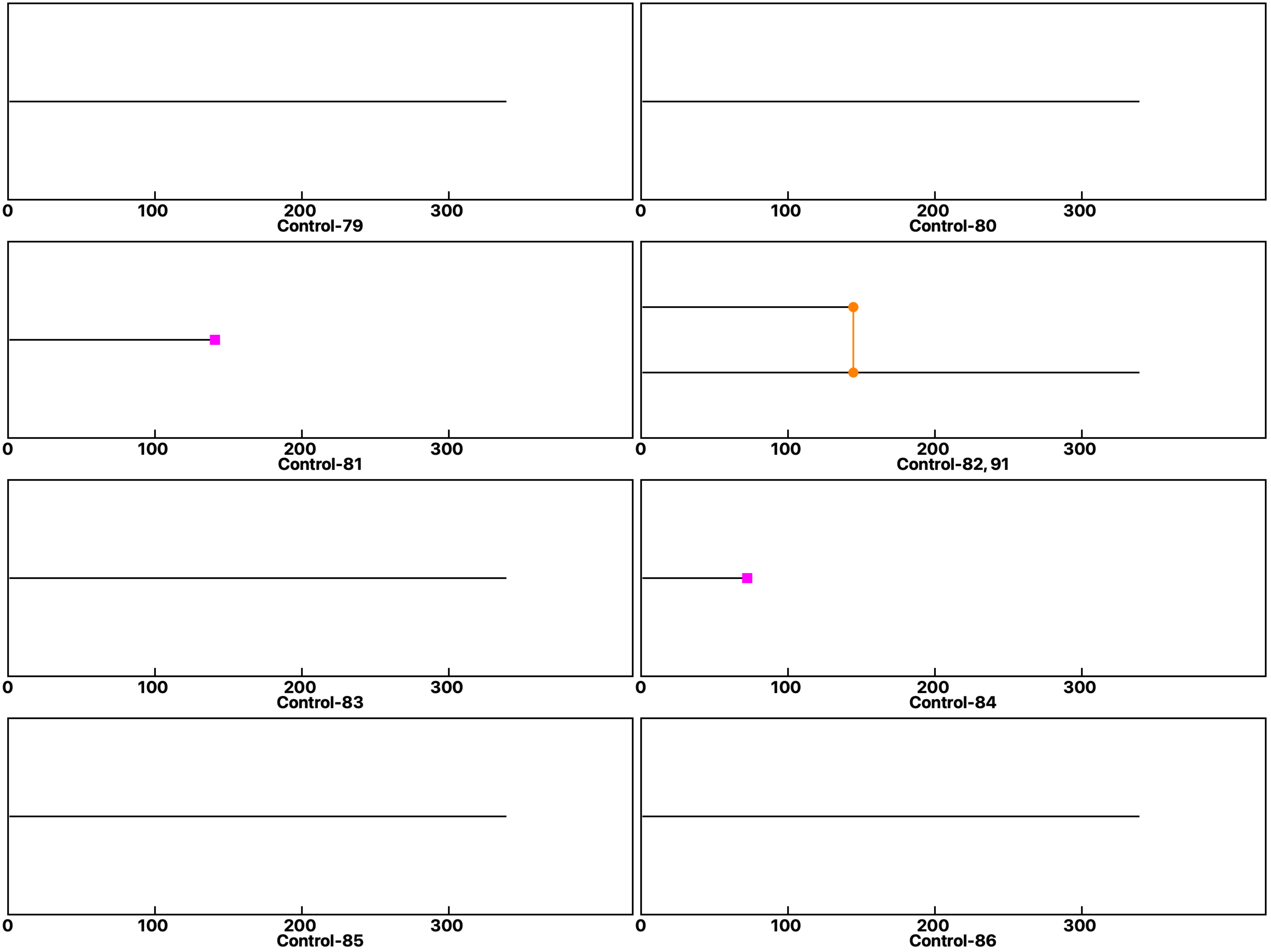

Analysis: Wild type-Dose, Treat.: Control, Cell: Myoblasts

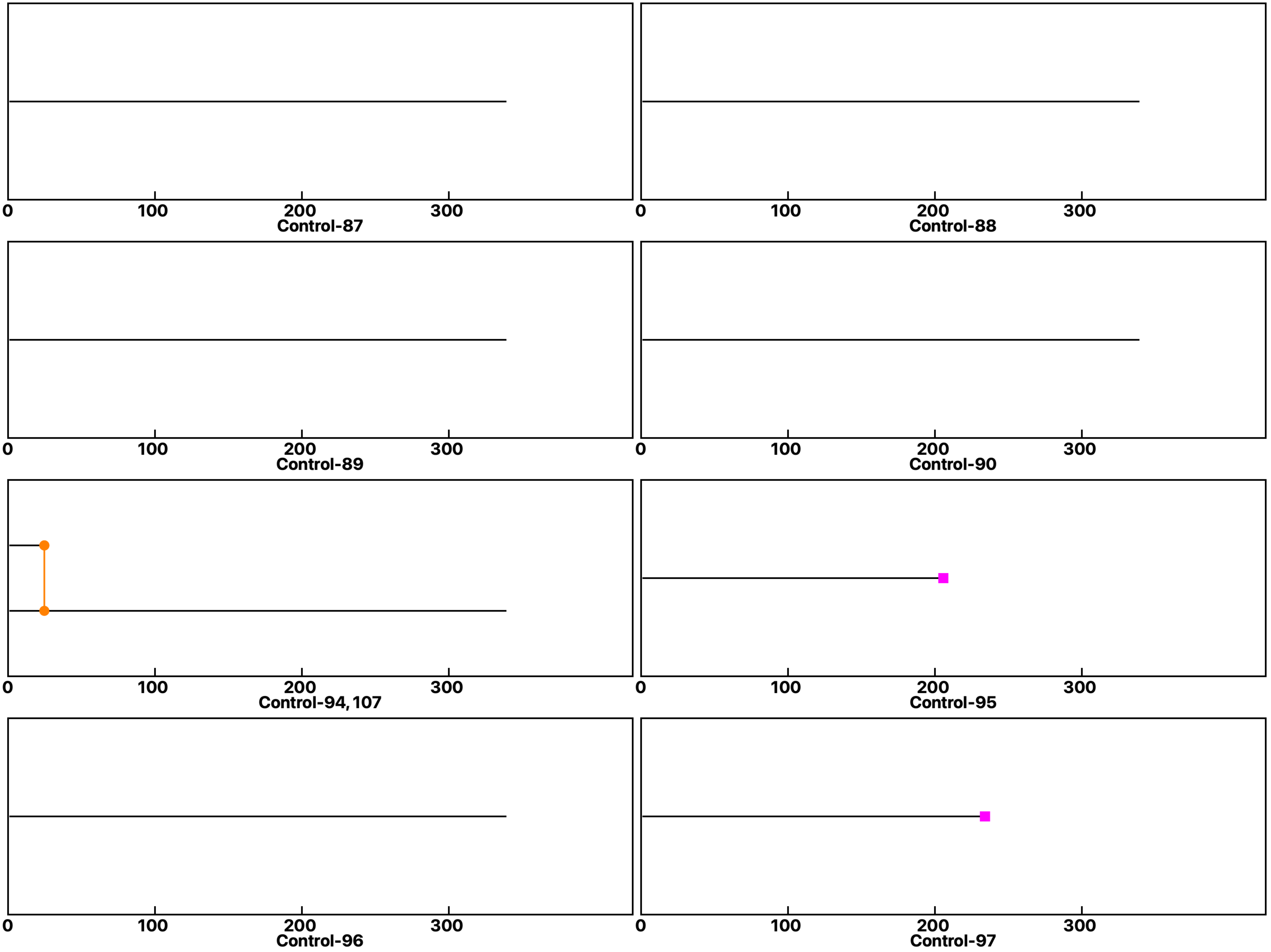

Analysis: Wild type-Dose, Treat.: Control, Cell: Myoblasts

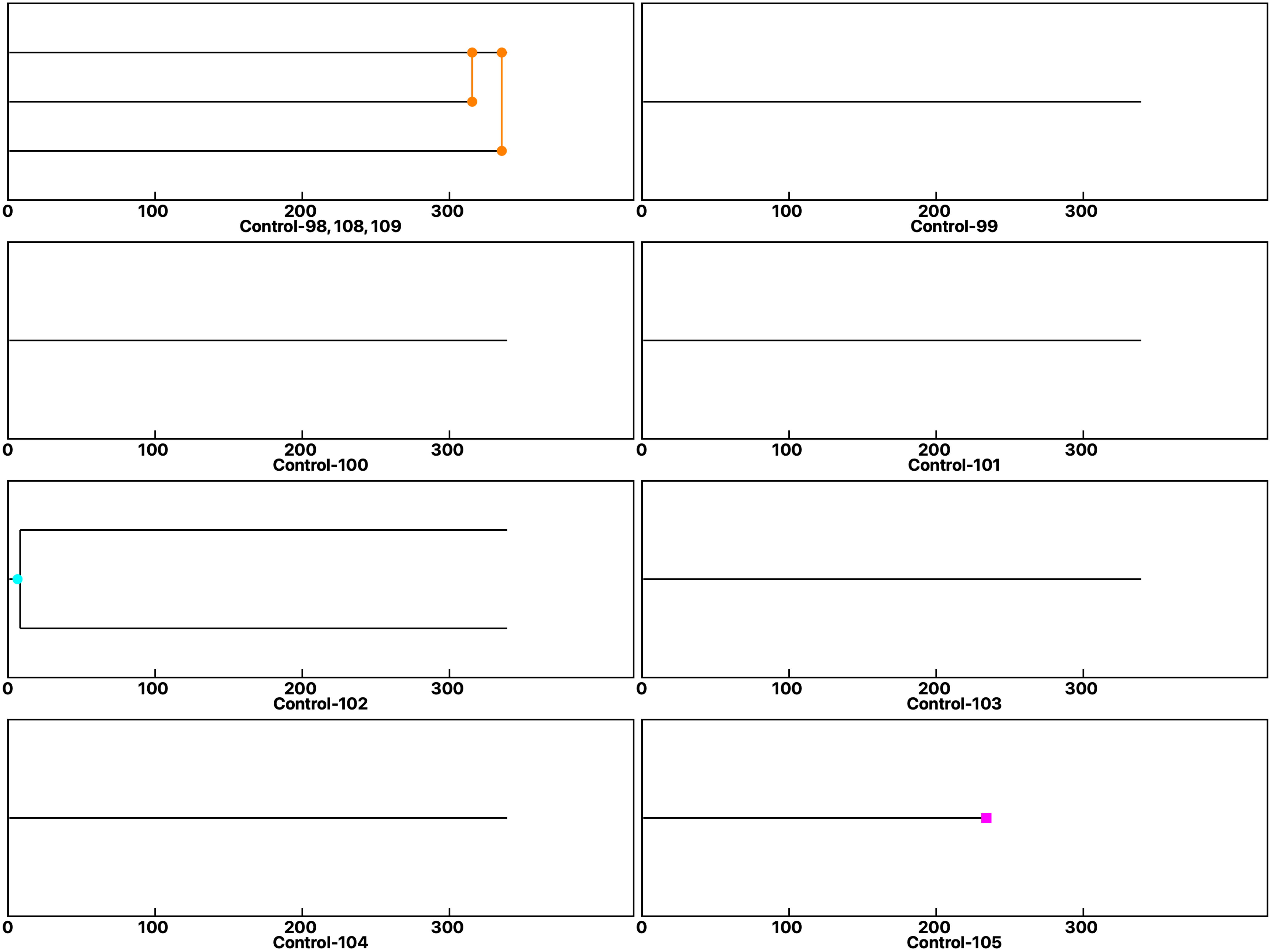

**Analysis: Wild type-Dose, Treat.: Control, Cell: Myoblasts**

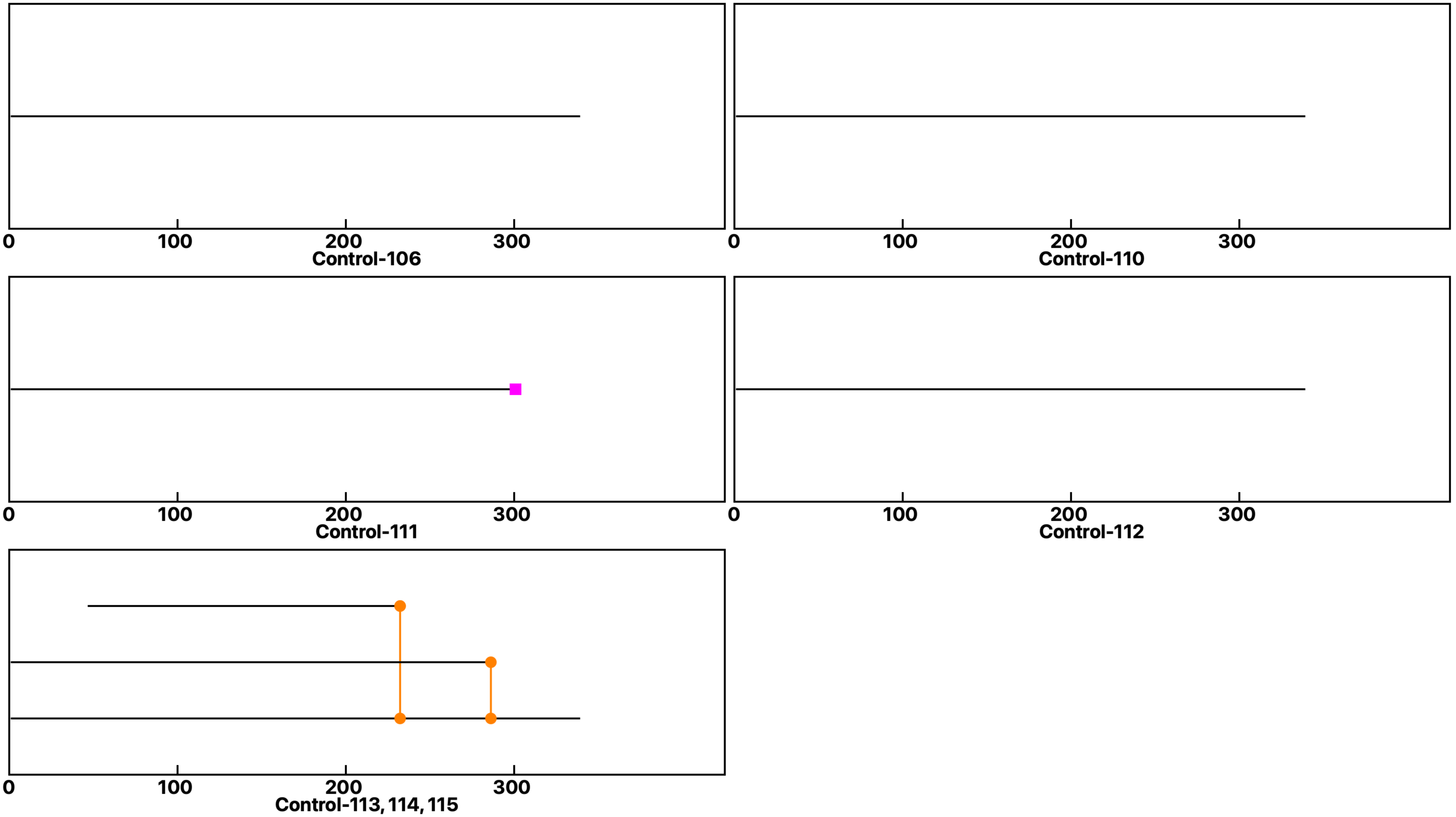
