## Supplemental data 2 for "N-Acetylglucosamine Facilitates Coordinated Myoblast Flow, Forming the Foundation for Efficient Myogenesis"

**Analysis: Wild type-Dose, Treat.: GlcNAc 1, Cell: Myoblasts**

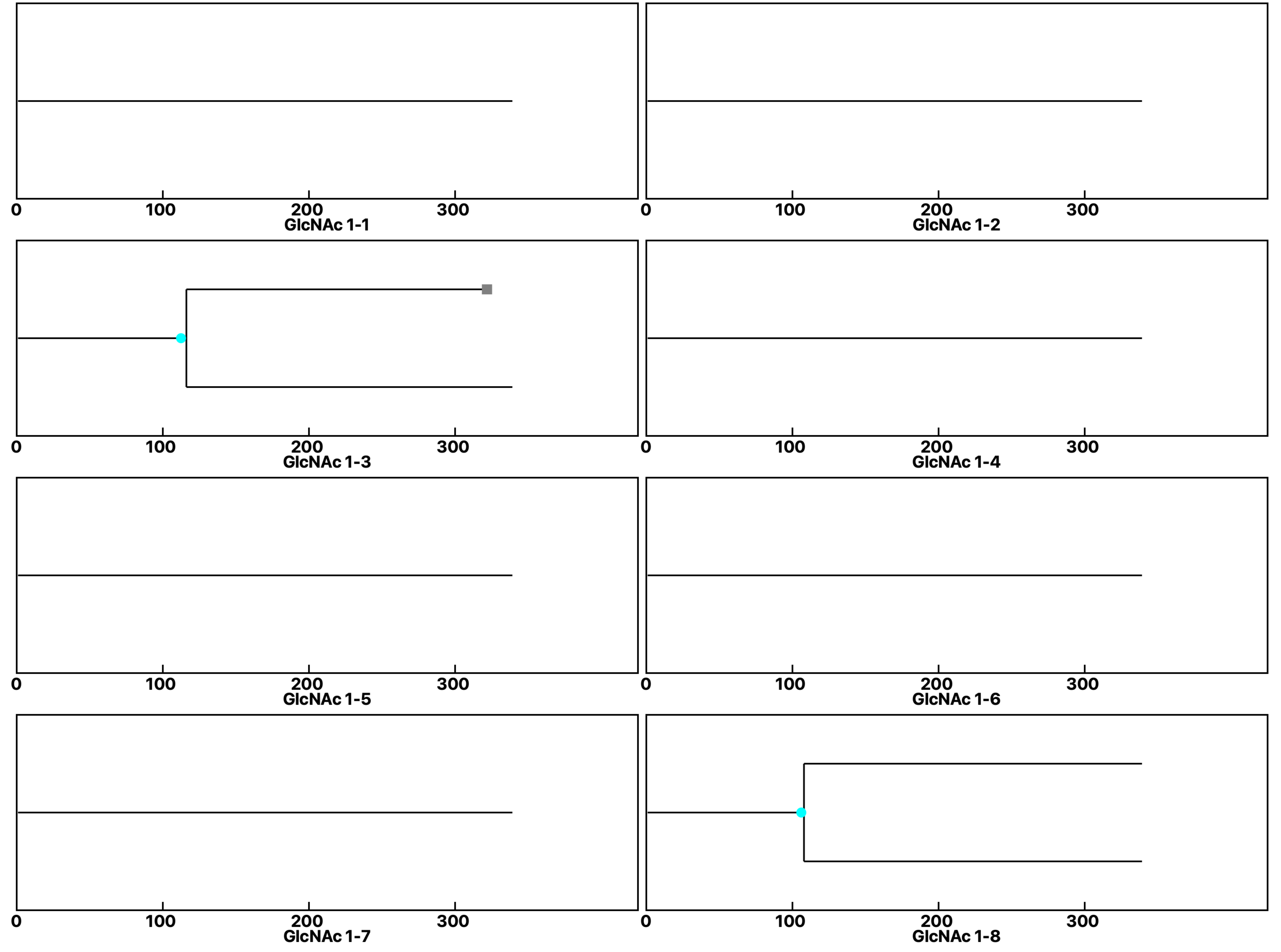

**Analysis: Wild type-Dose, Treat.: GlcNAc 1, Cell: Myoblasts**

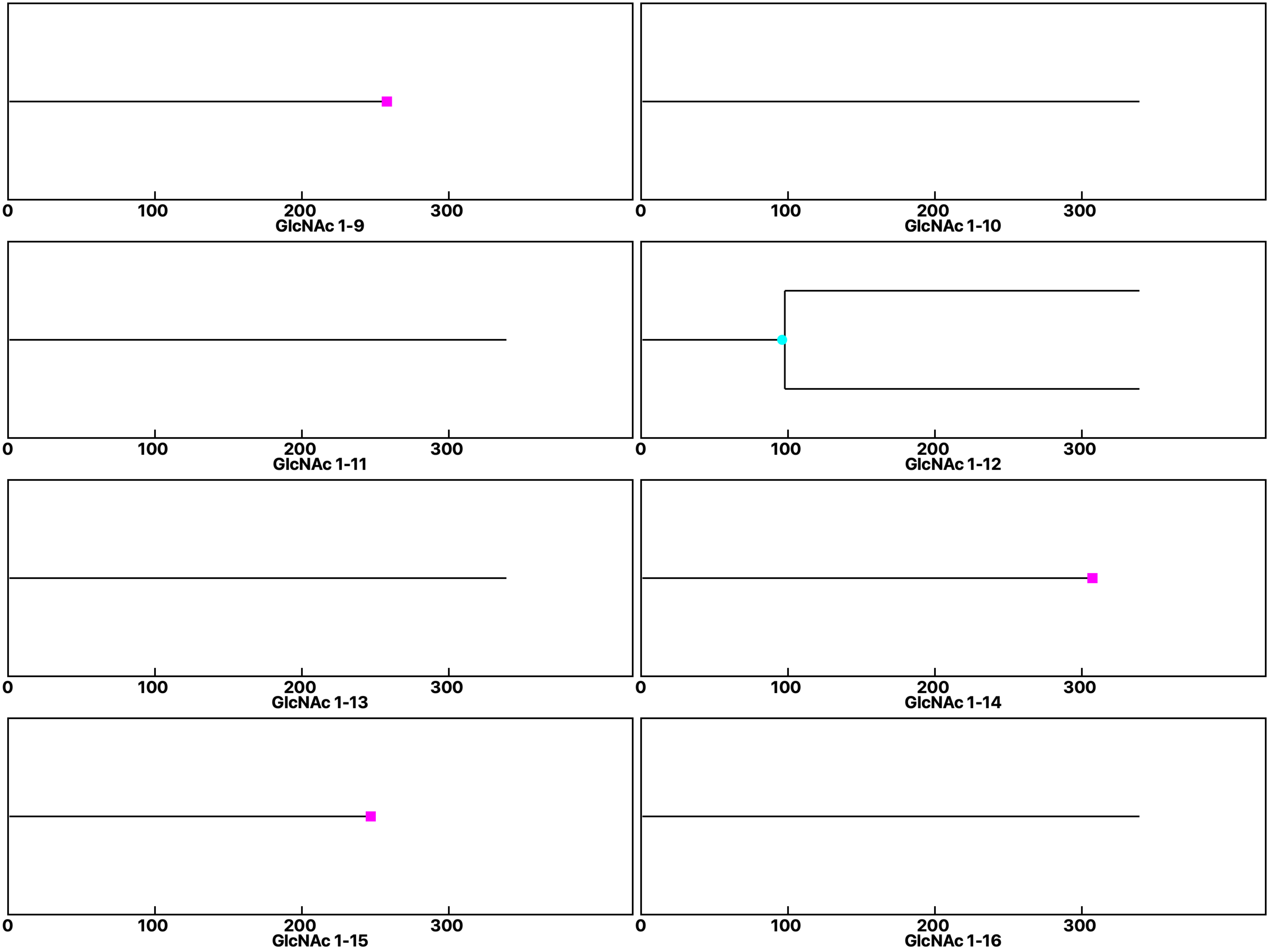

Analysis: Wild type-Dose, Treat.: GlcNAc 1, Cell: Myoblasts

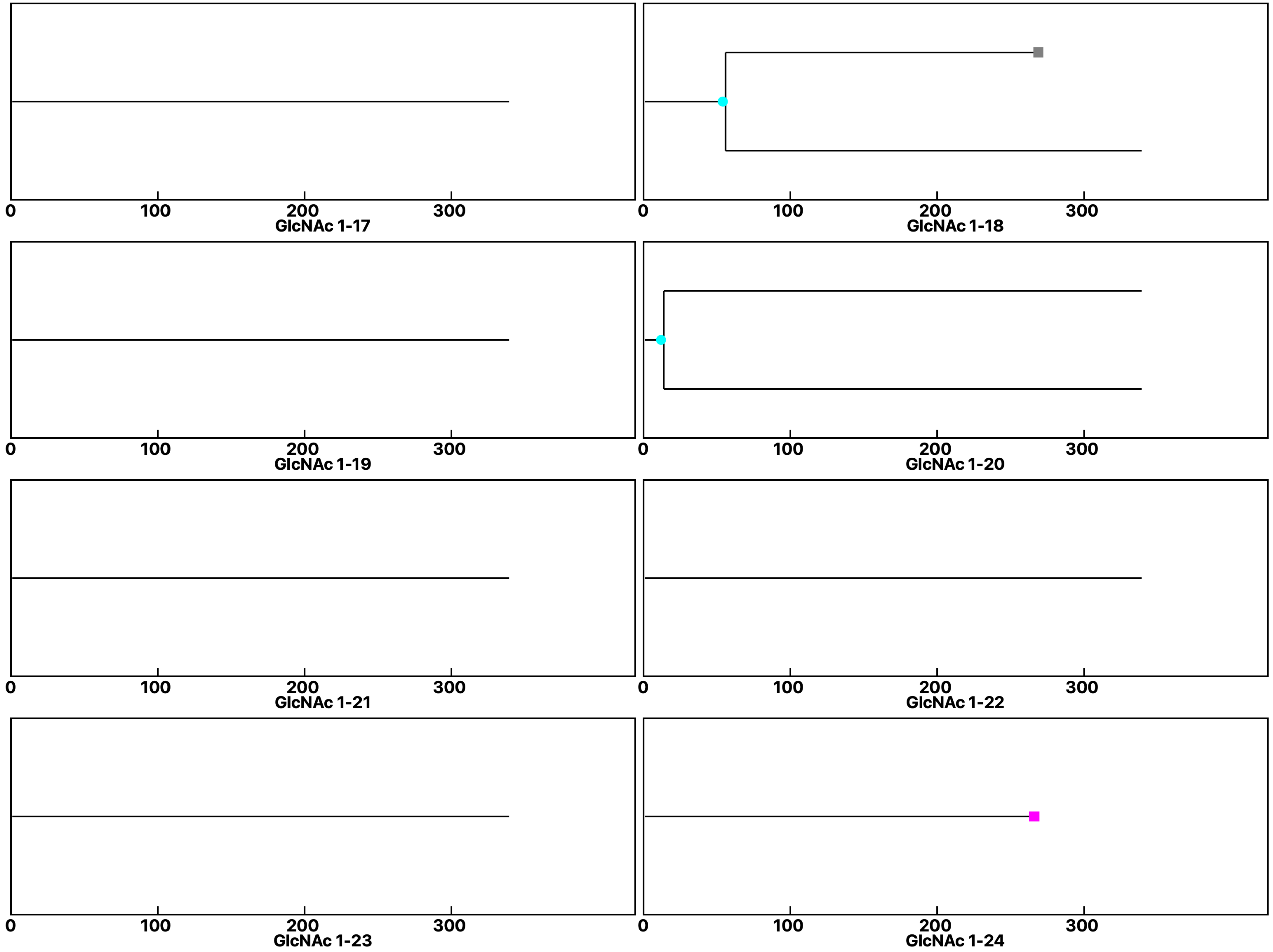

**Analysis: Wild type-Dose, Treat.: GlcNAc 1, Cell: Myoblasts**

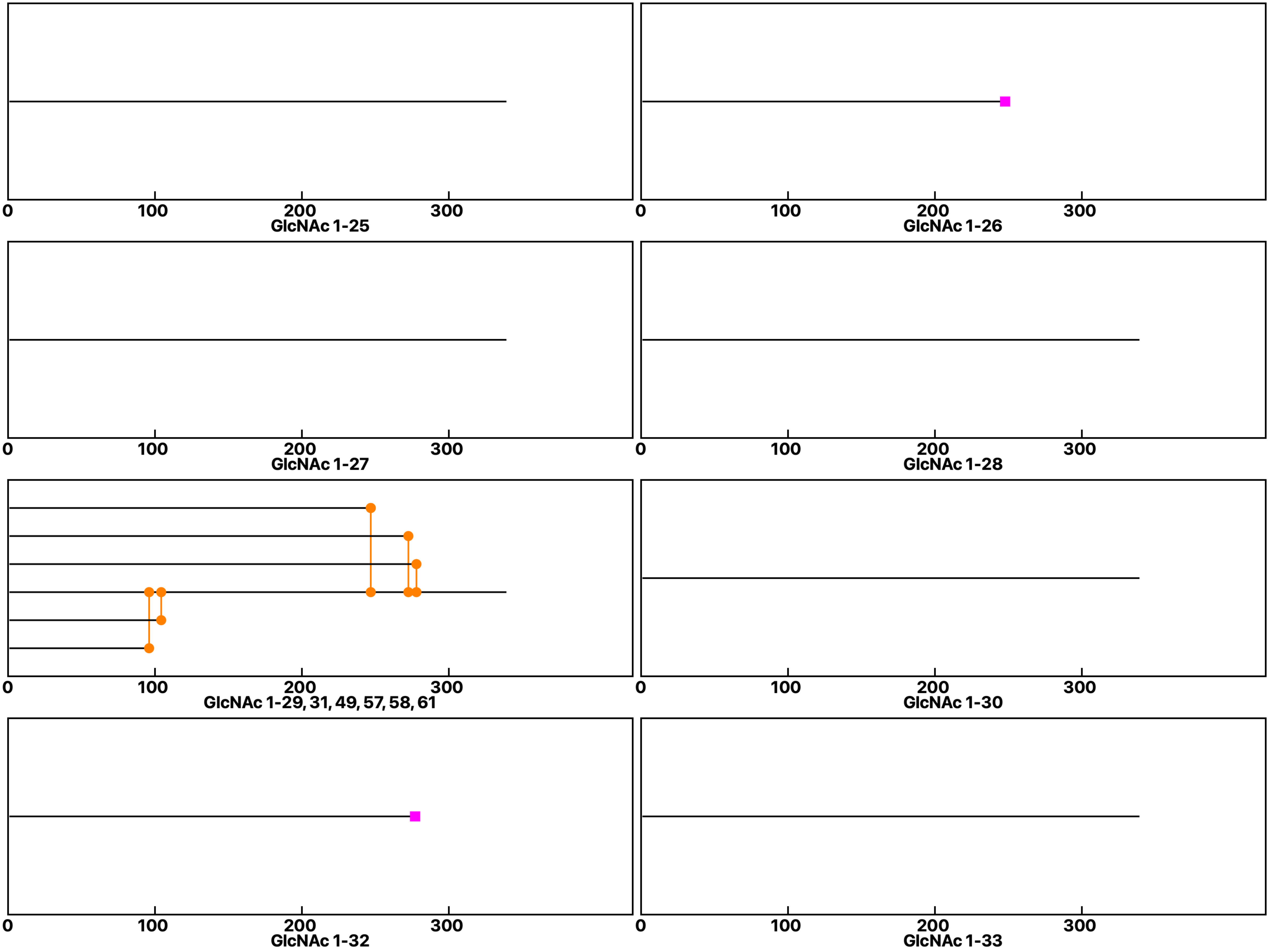

### Analysis: Wild type-Dose, Treat.: GlcNAc 1, Cell: Myoblasts

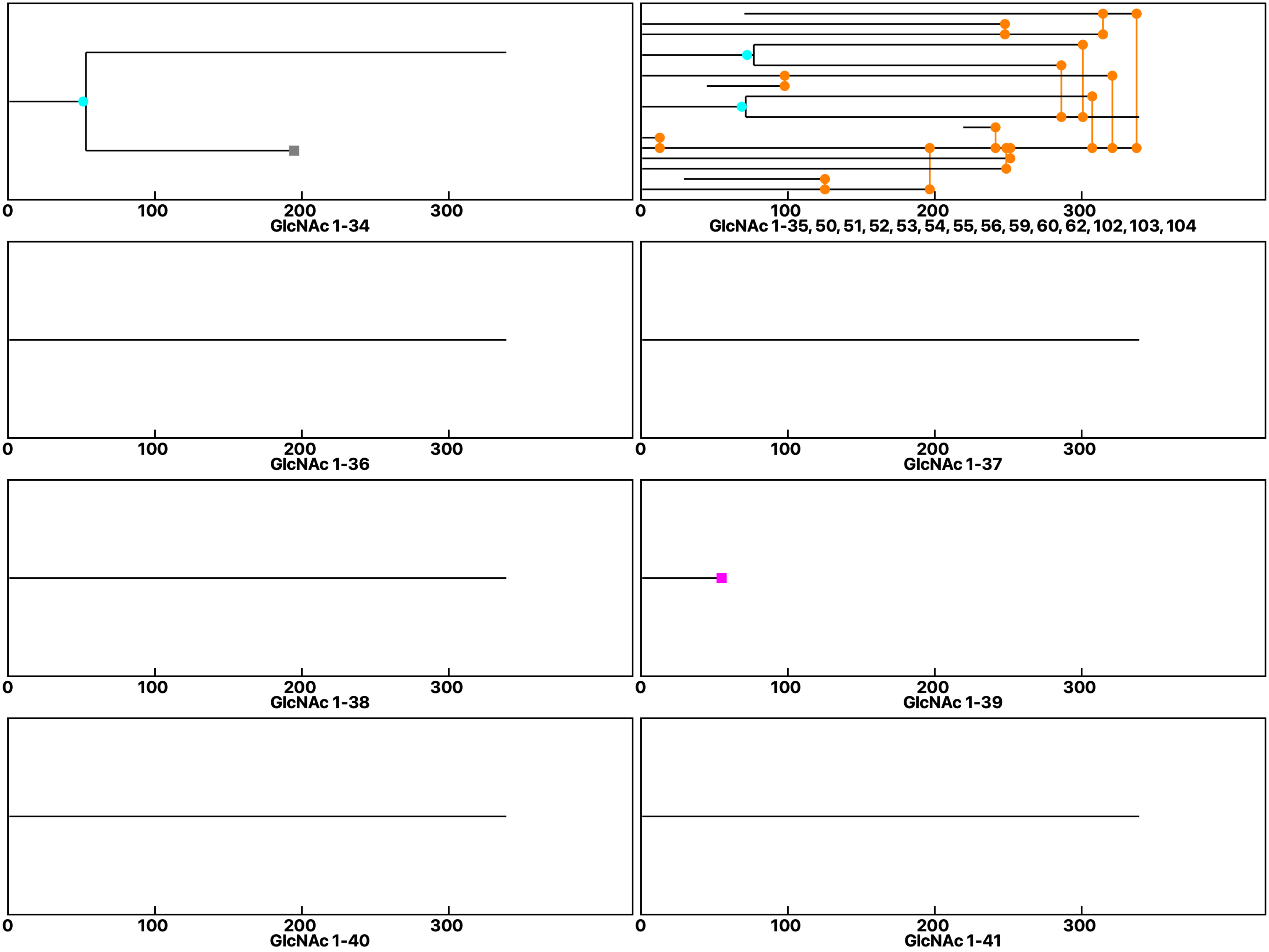

**Analysis: Wild type-Dose, Treat.: GlcNAc 1, Cell: Myoblasts**

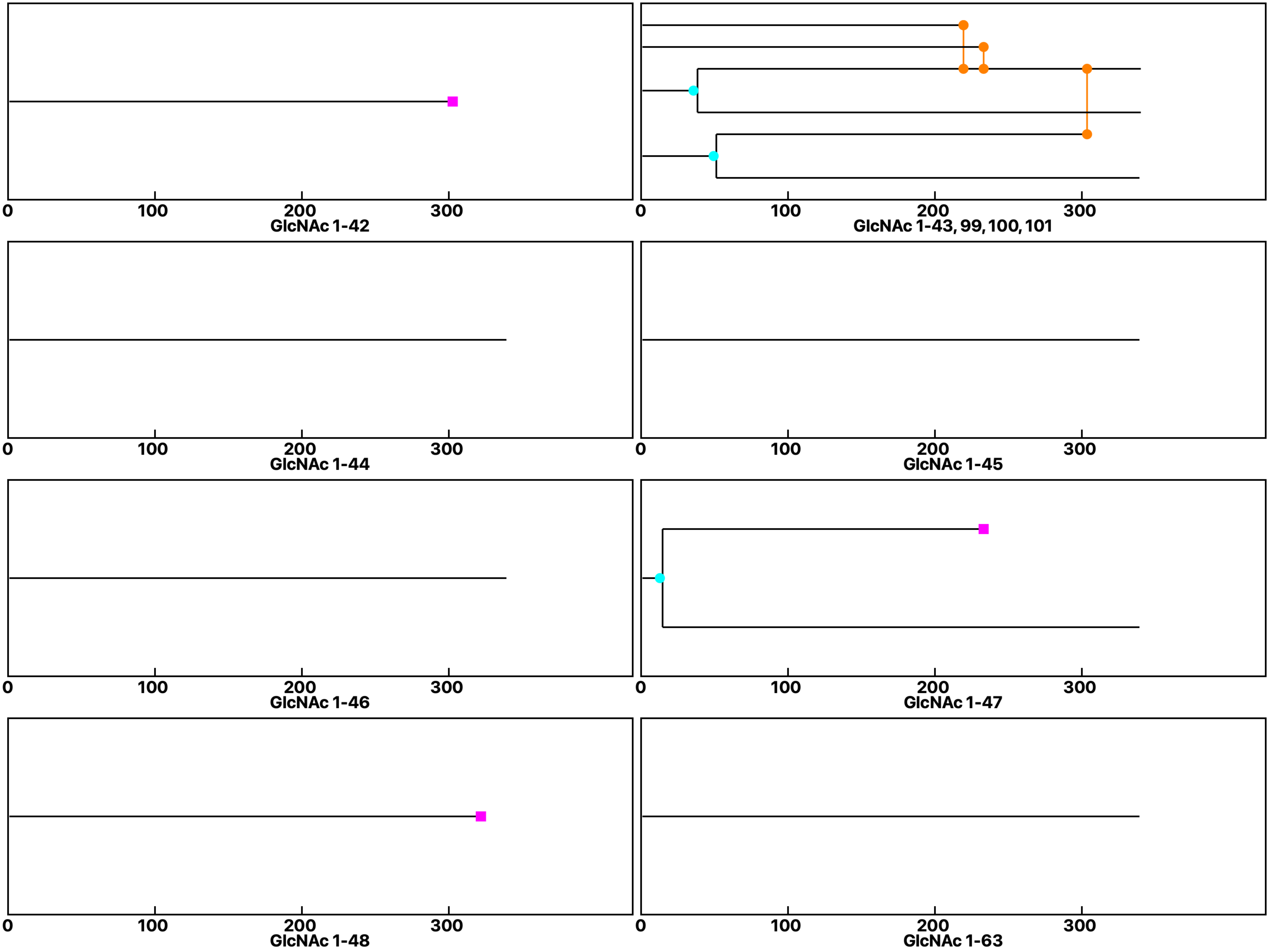

**Analysis: Wild type-Dose, Treat.: GlcNAc 1, Cell: Myoblasts**

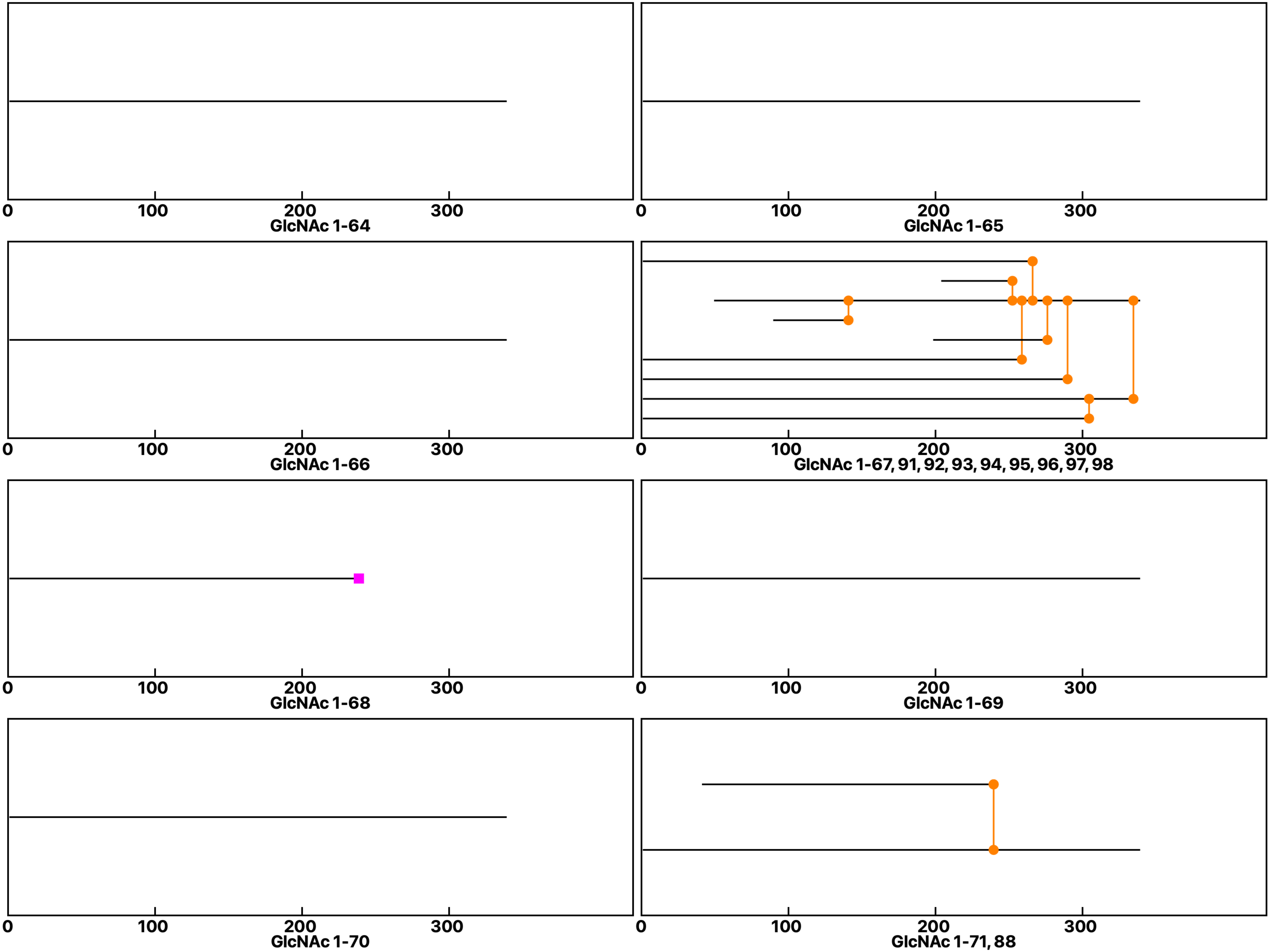

Analysis: Wild type-Dose, Treat.: GlcNAc 1, Cell: Myoblasts

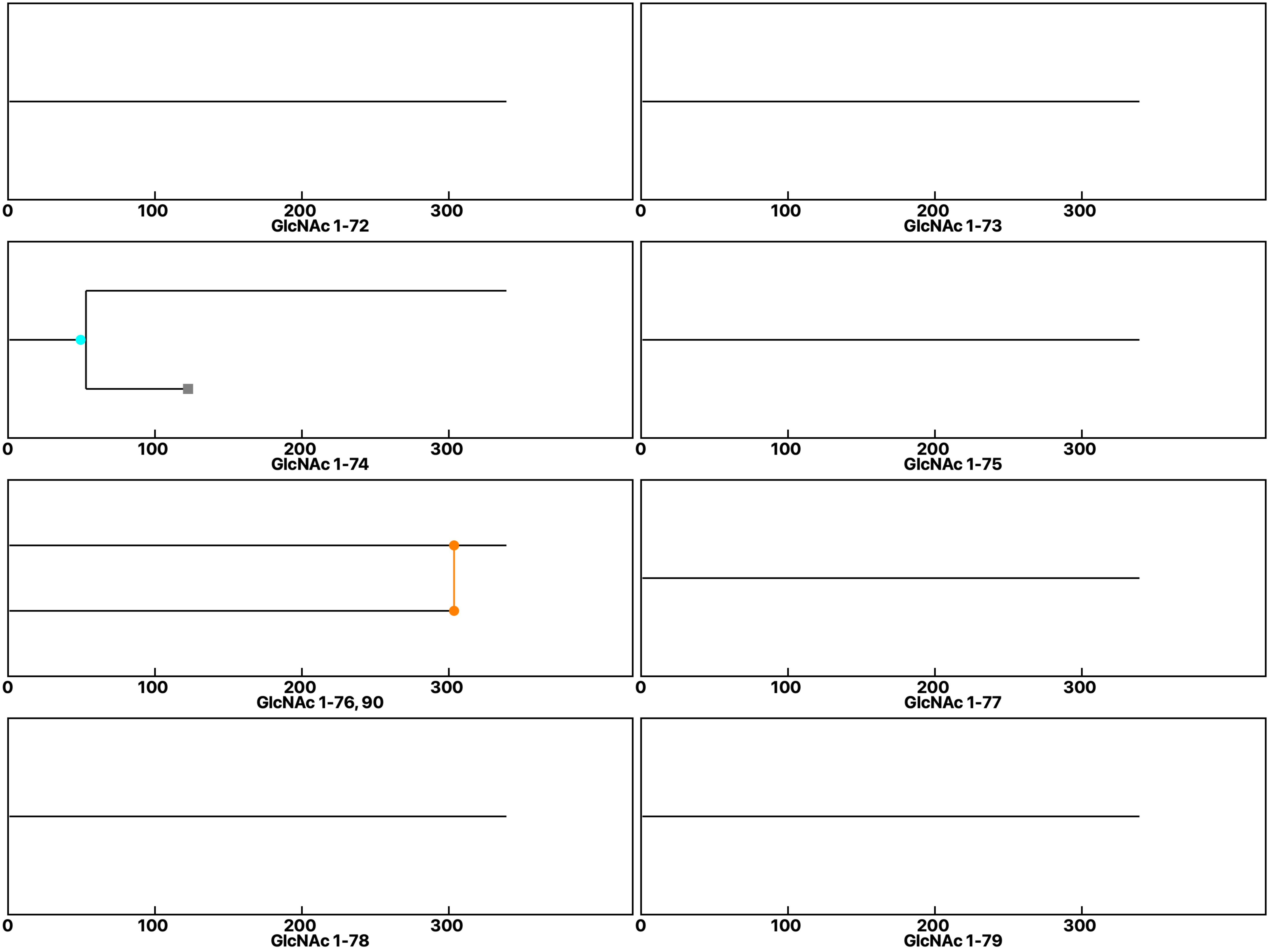
