## Supplemental data 3 for "N-Acetylglucosamine Facilitates Coordinated Myoblast Flow, Forming the Foundation for Efficient Myogenesis"

Analysis: Wild type-Dose, Treat.: GlcNAc 5, Cell: Myoblasts

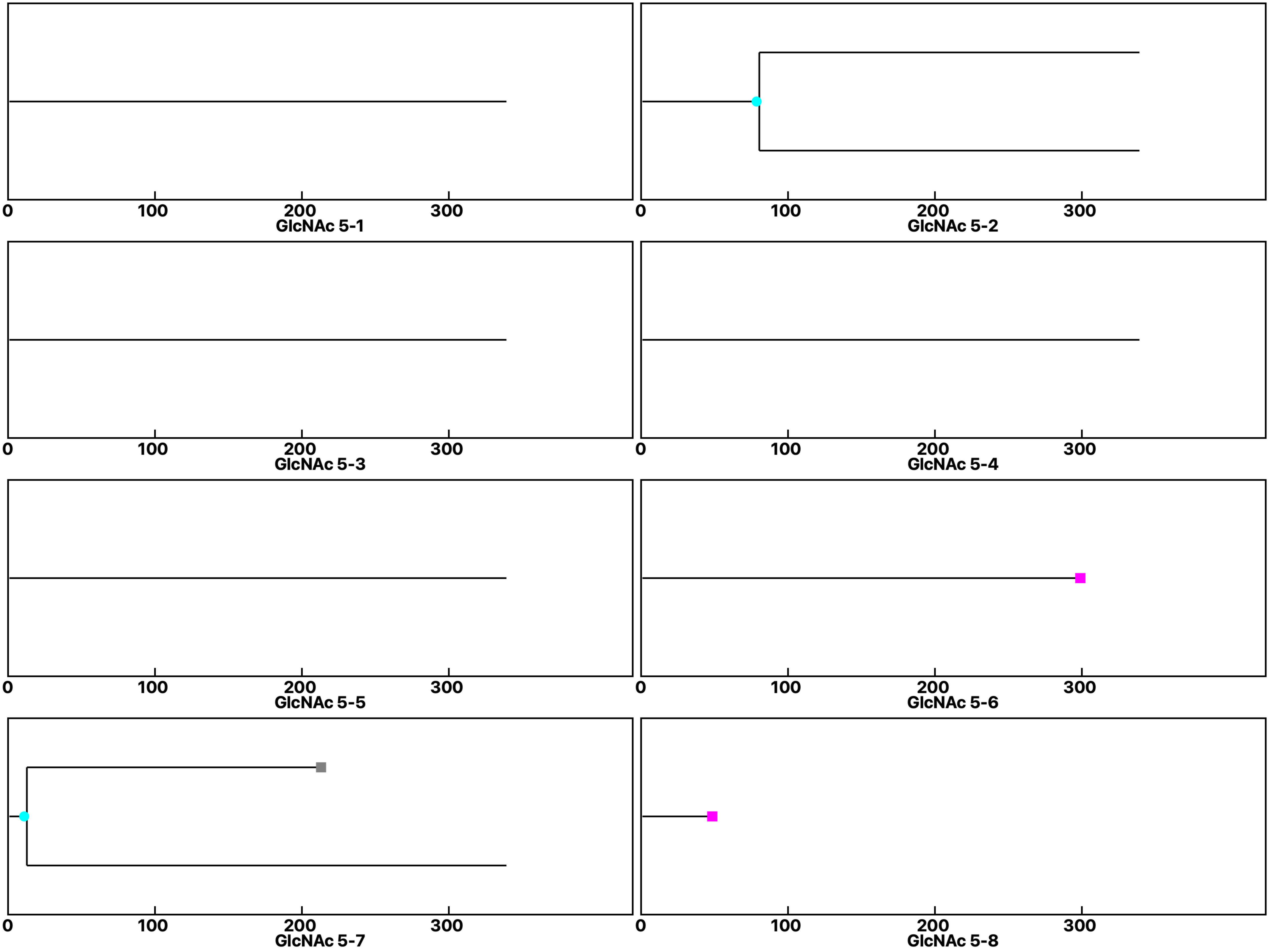

Analysis: Wild type-Dose, Treat.: GlcNAc 5, Cell: Myoblasts

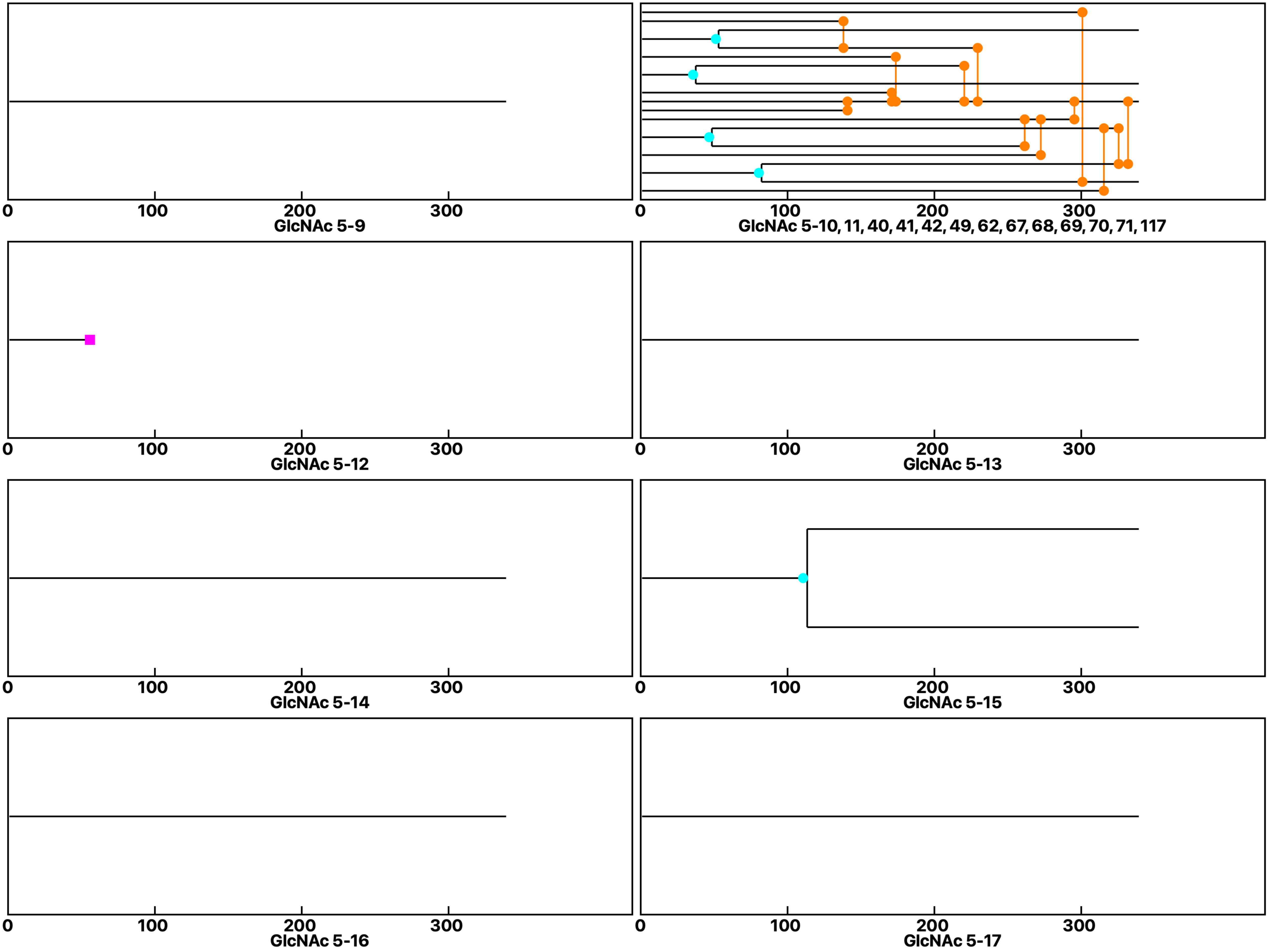

Analysis: Wild type-Dose, Treat.: GlcNAc 5, Cell: Myoblasts

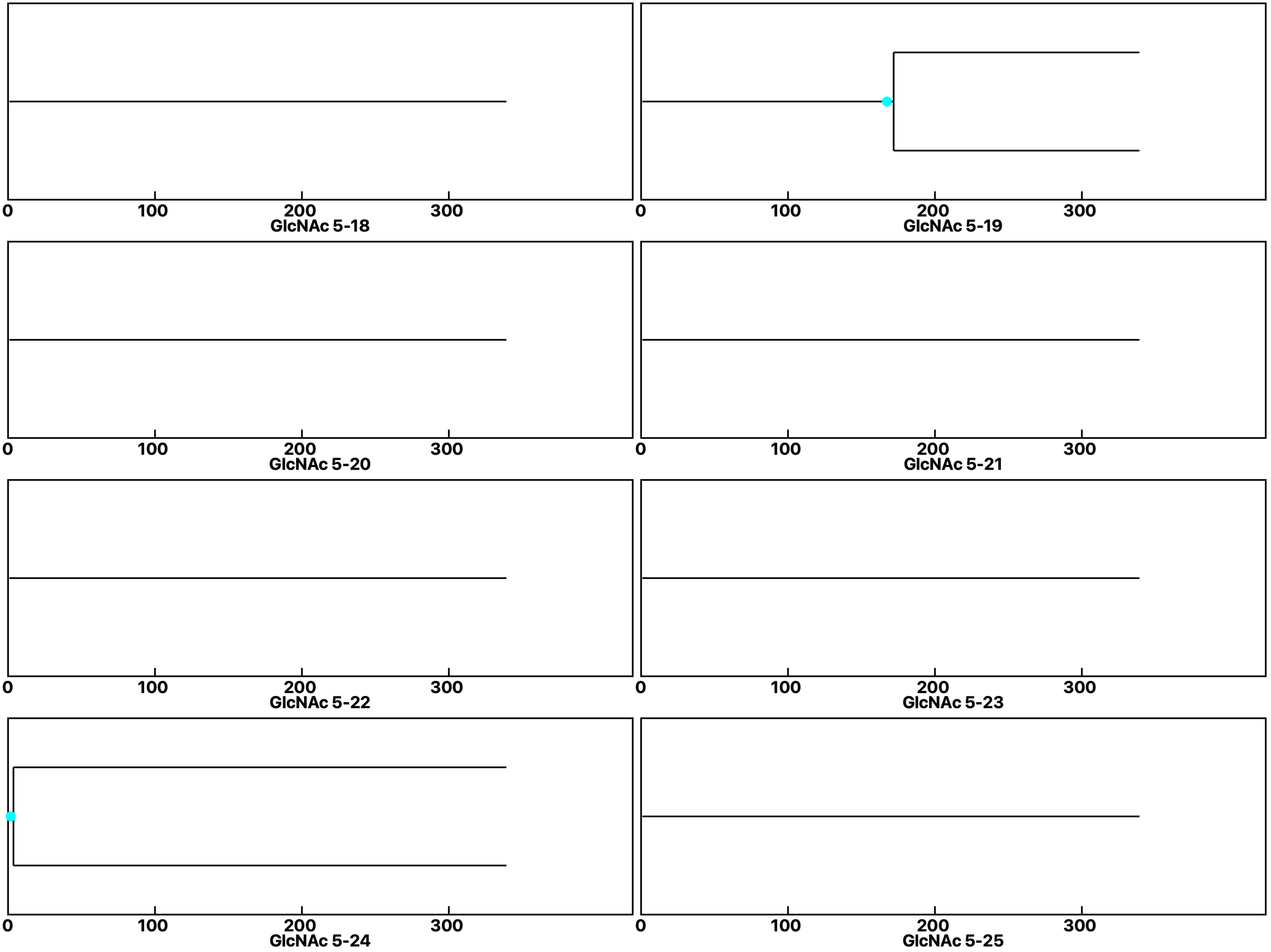

Analysis: Wild type-Dose, Treat.: GlcNAc 5, Cell: Myoblasts

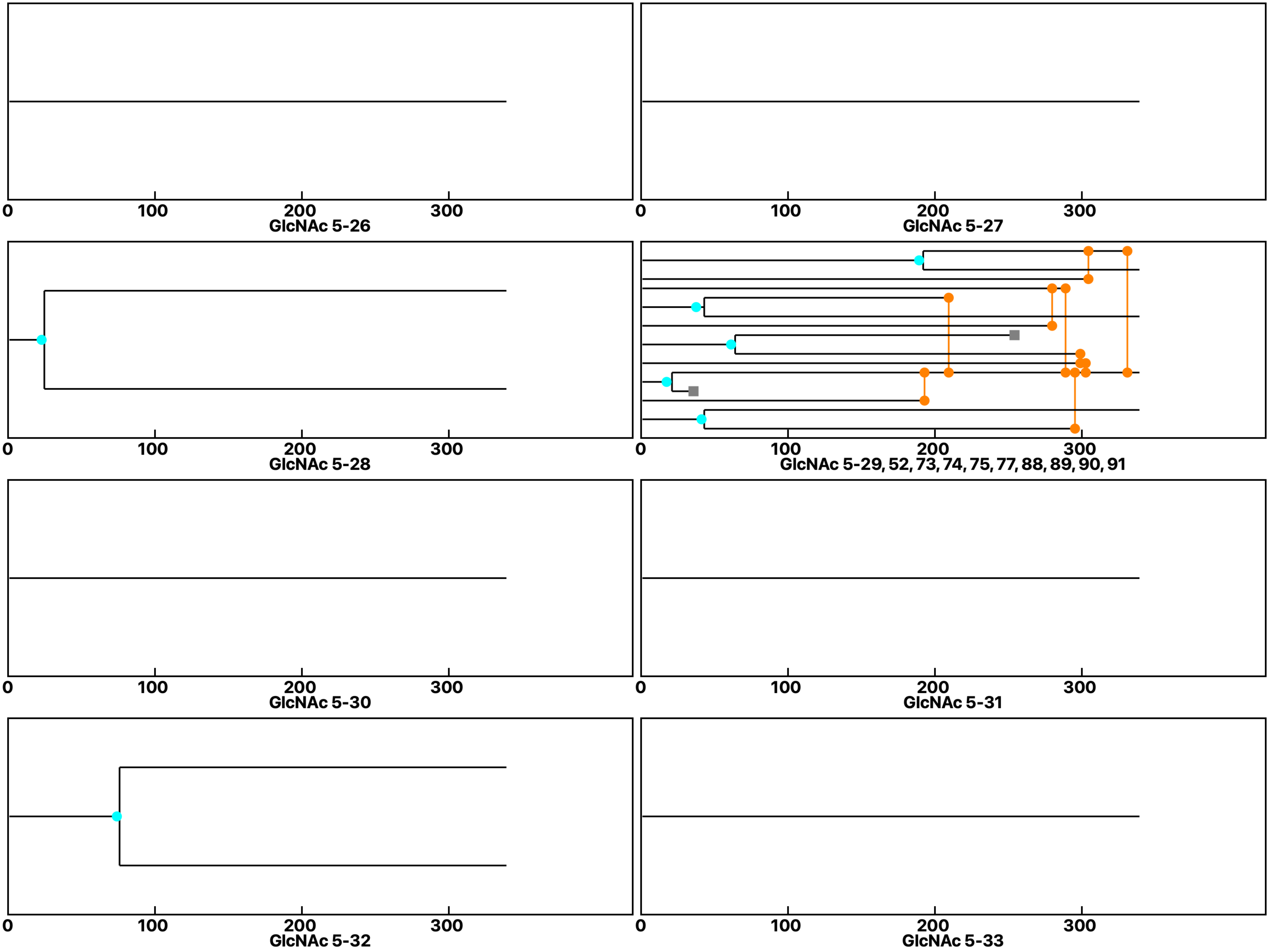

Analysis: Wild type-Dose, Treat.: GlcNAc 5, Cell: Myoblasts

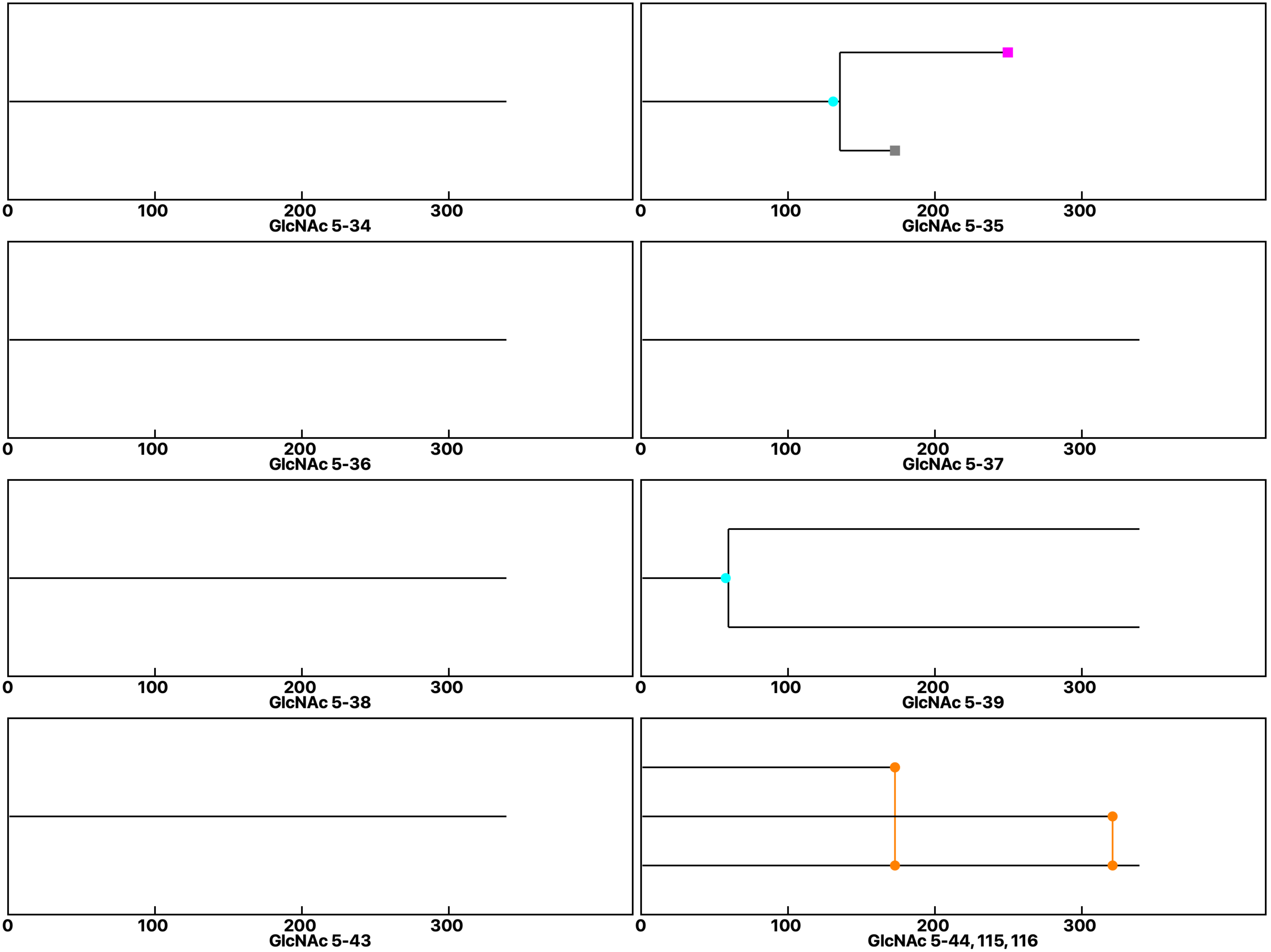

Analysis: Wild type-Dose, Treat.: GlcNAc 5, Cell: Myoblasts

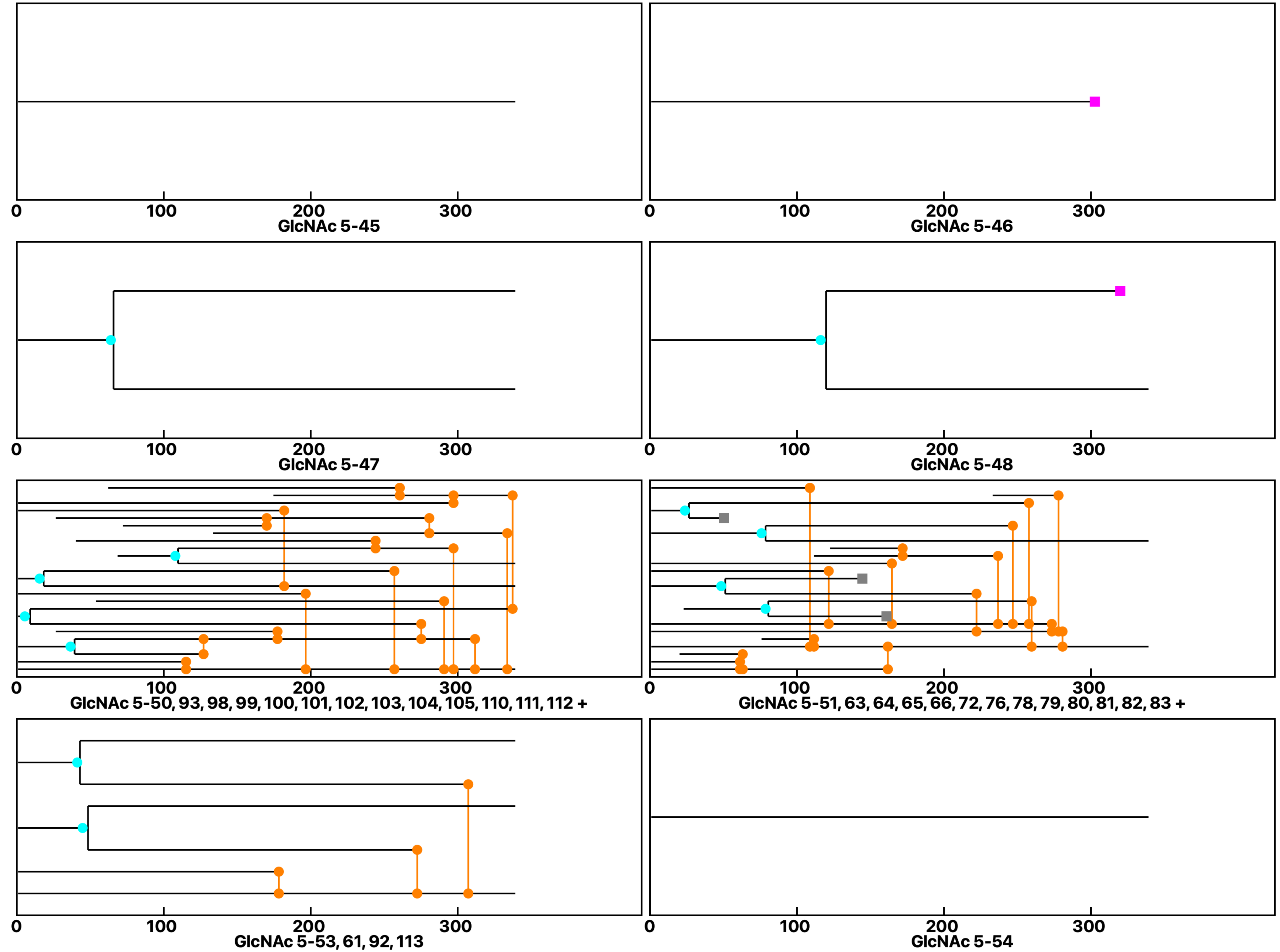

Analysis: Wild type-Dose, Treat.: GlcNAc 5, Cell: Myoblasts

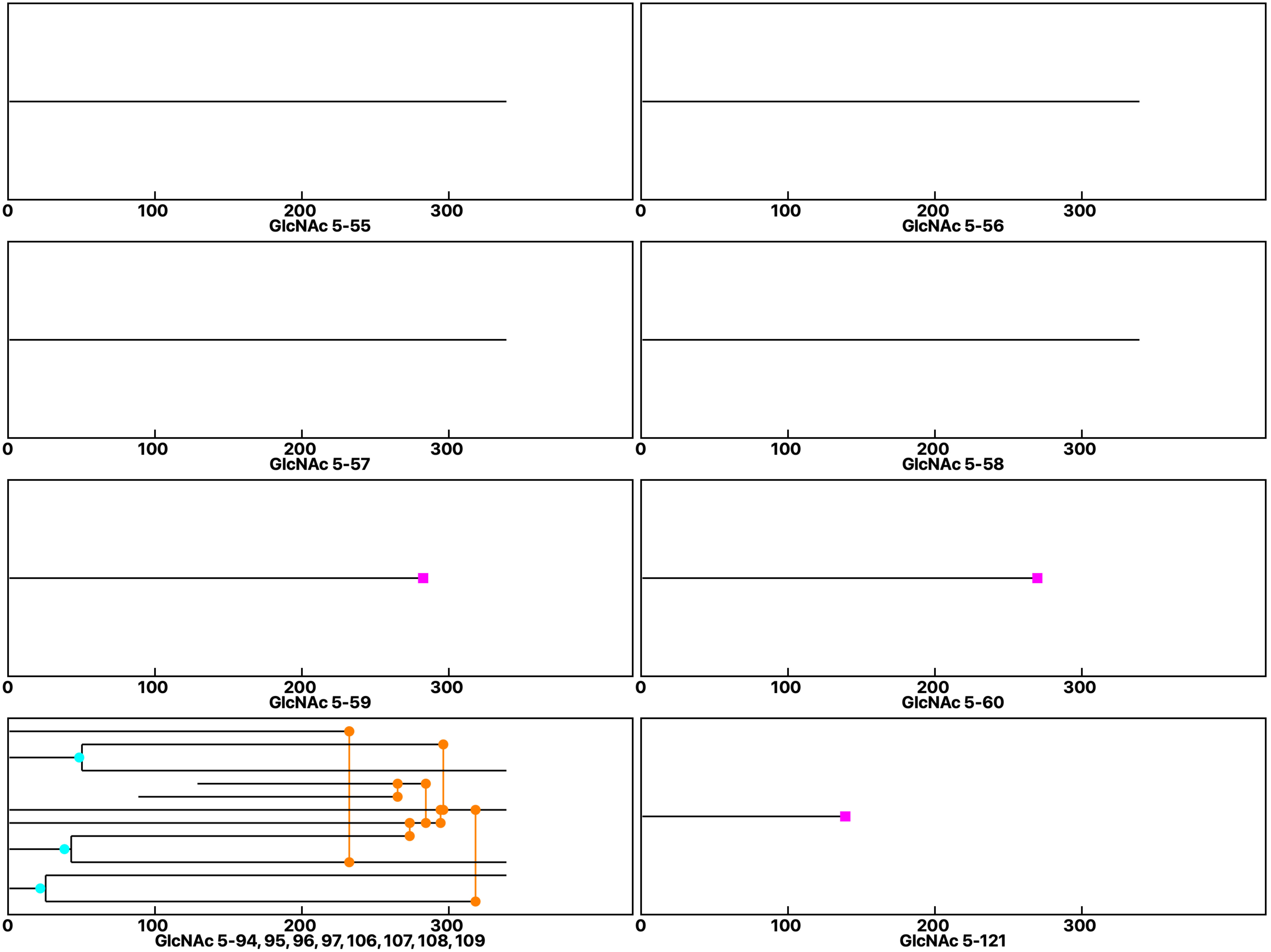

**Analysis: Wild type-Dose, Treat.: GlcNAc 5, Cell: Myoblasts**

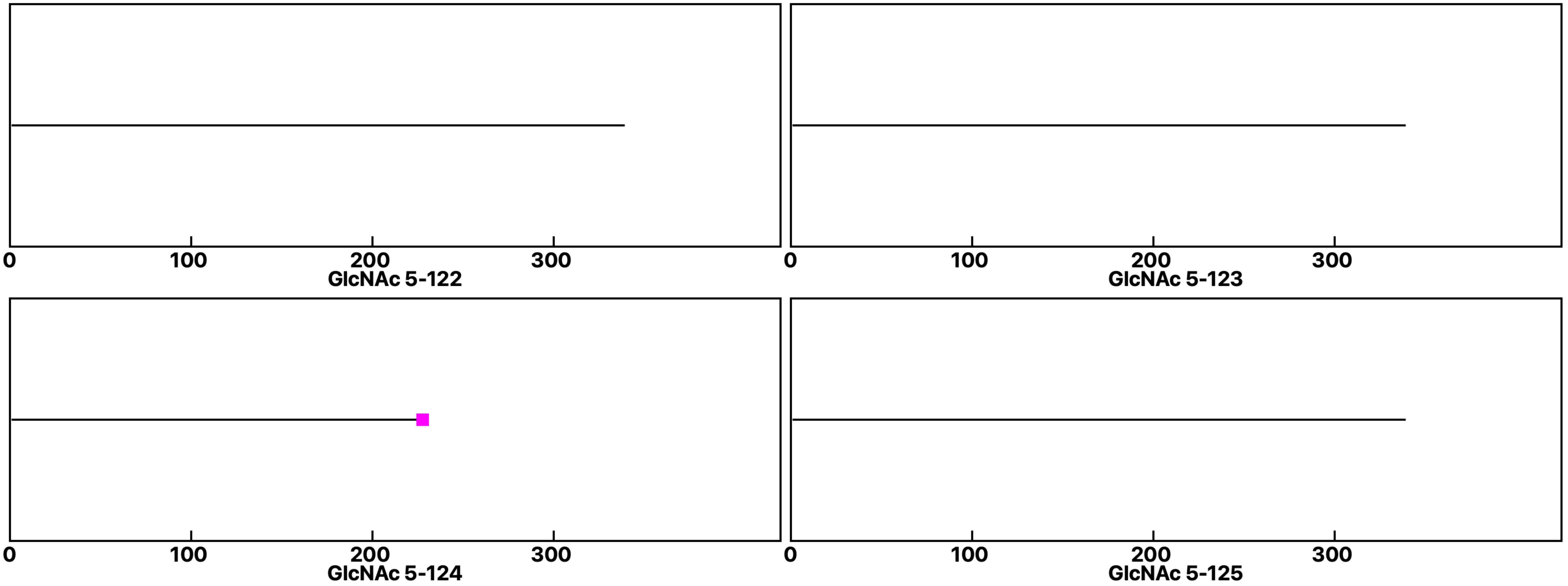
