## Supplemental data 4 for "N-Acetylglucosamine Facilitates Coordinated Myoblast Flow, Forming the Foundation for Efficient Myogenesis"

**Analysis: Wild type-Dose, Treat.: GlcNAc 10, Cell: Myoblasts**

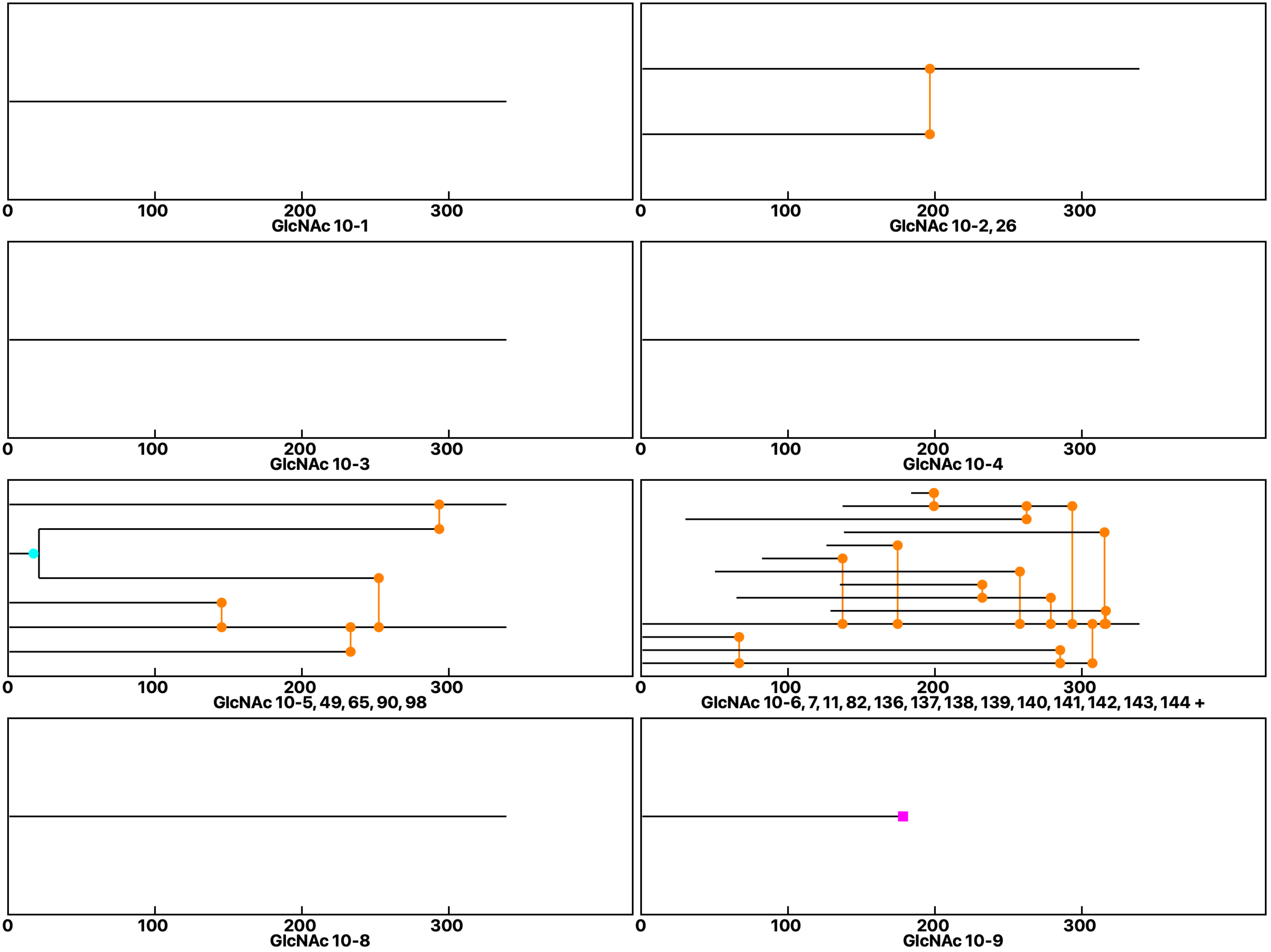

**Analysis: Wild type-Dose, Treat.: GlcNAc 10, Cell: Myoblasts**

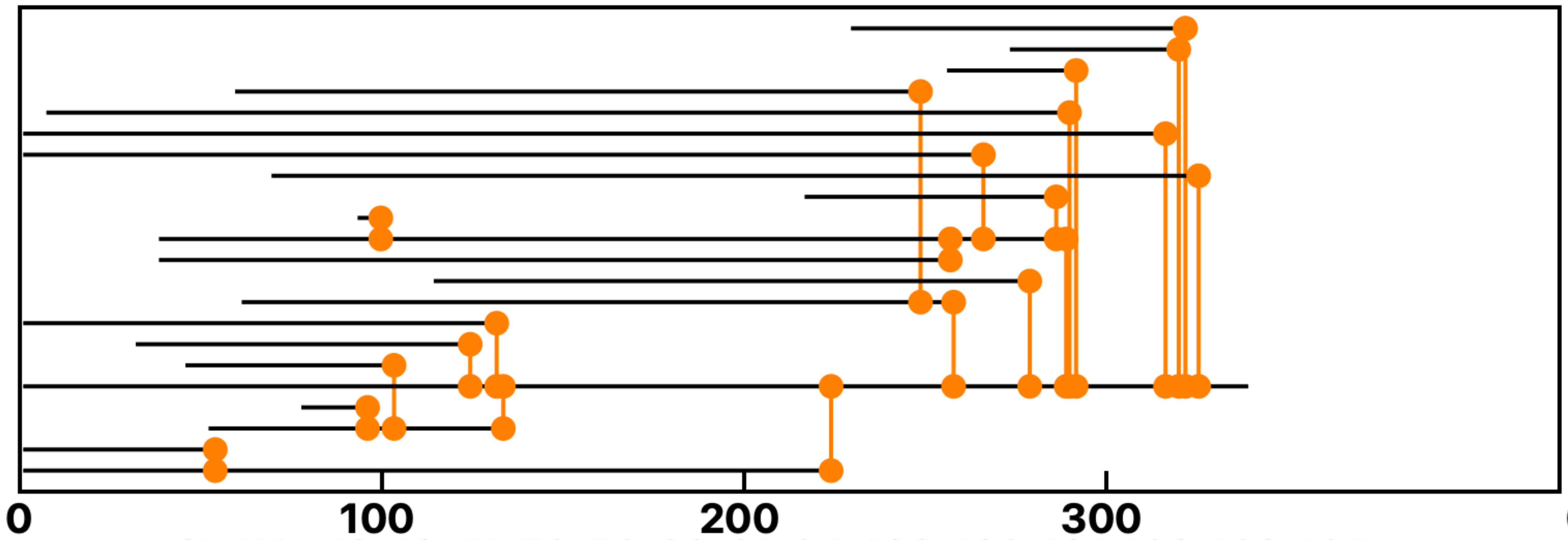

GlcNAc 10-10, 41, 58, 59, 60, 61, 64, 108, 120, 121, 122, 123, 124 +

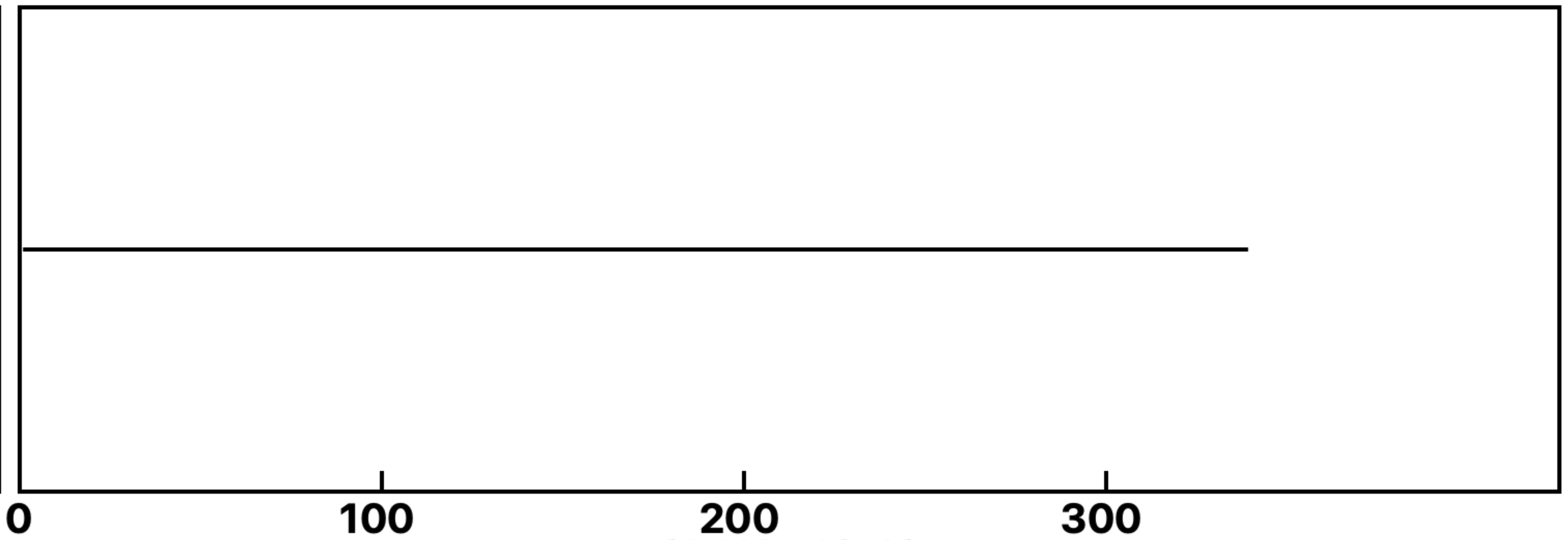

GlcNAc 10-12

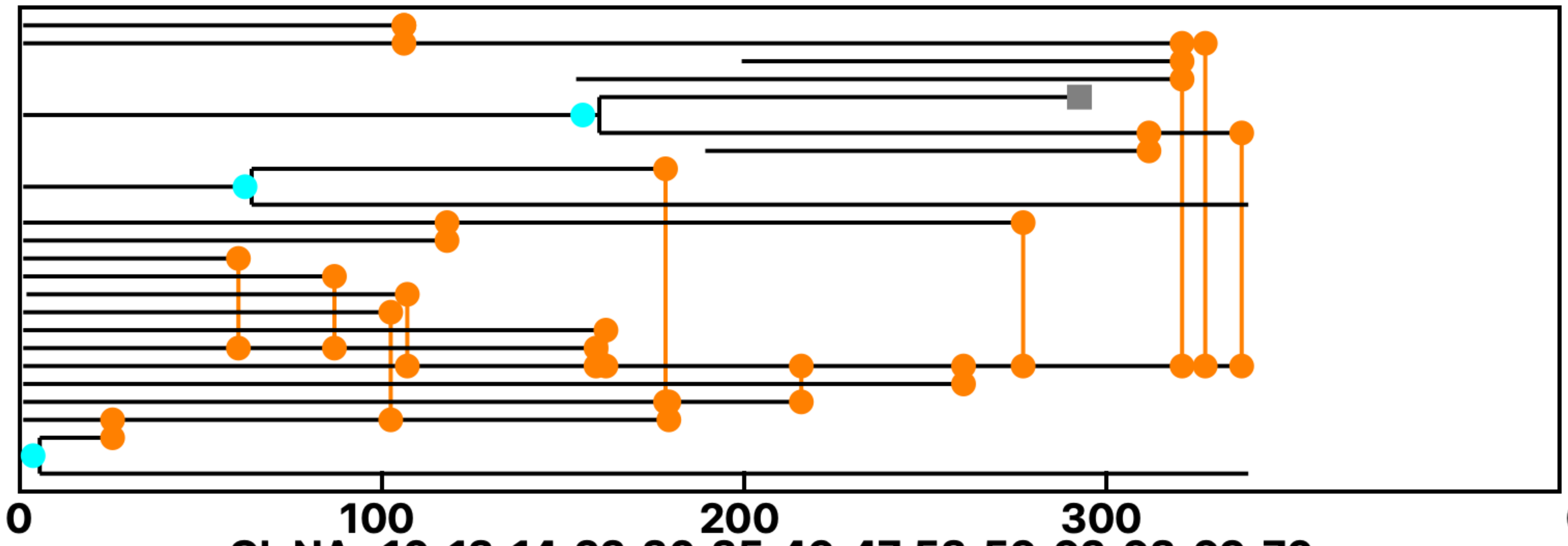

GlcNAc 10-13, 14, 22, 30, 35, 46, 47, 53, 56, 62, 63, 69, 70 +

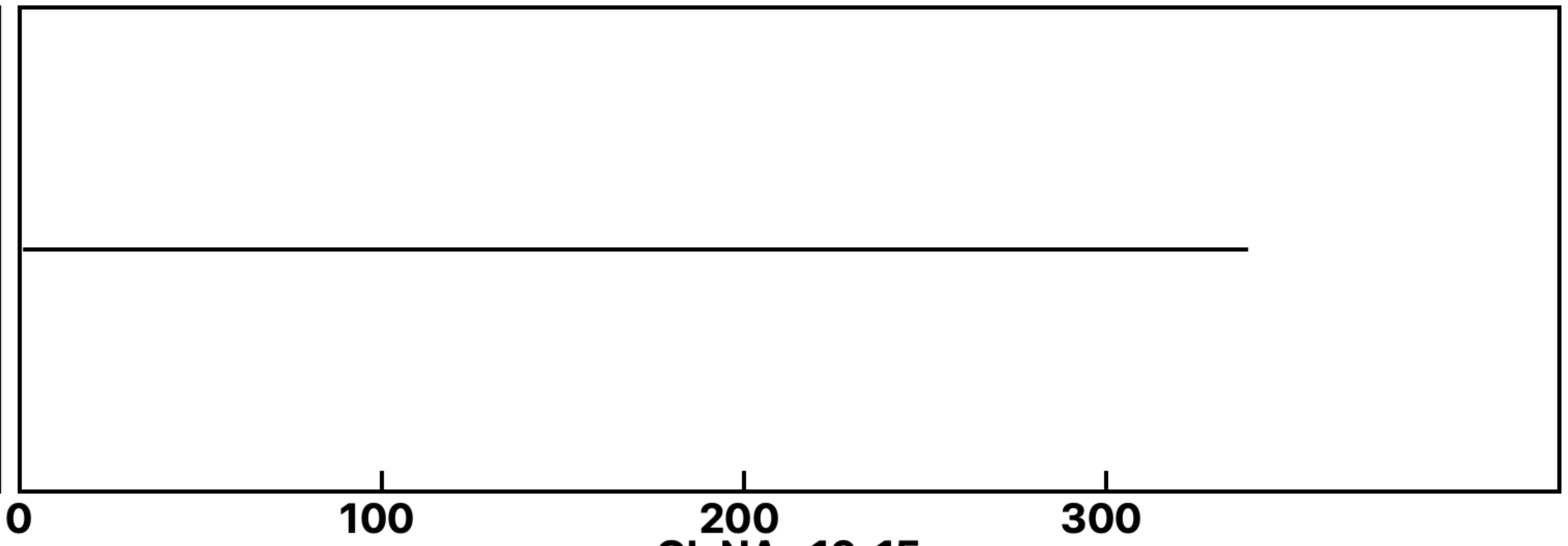

GlcNAc 10-15

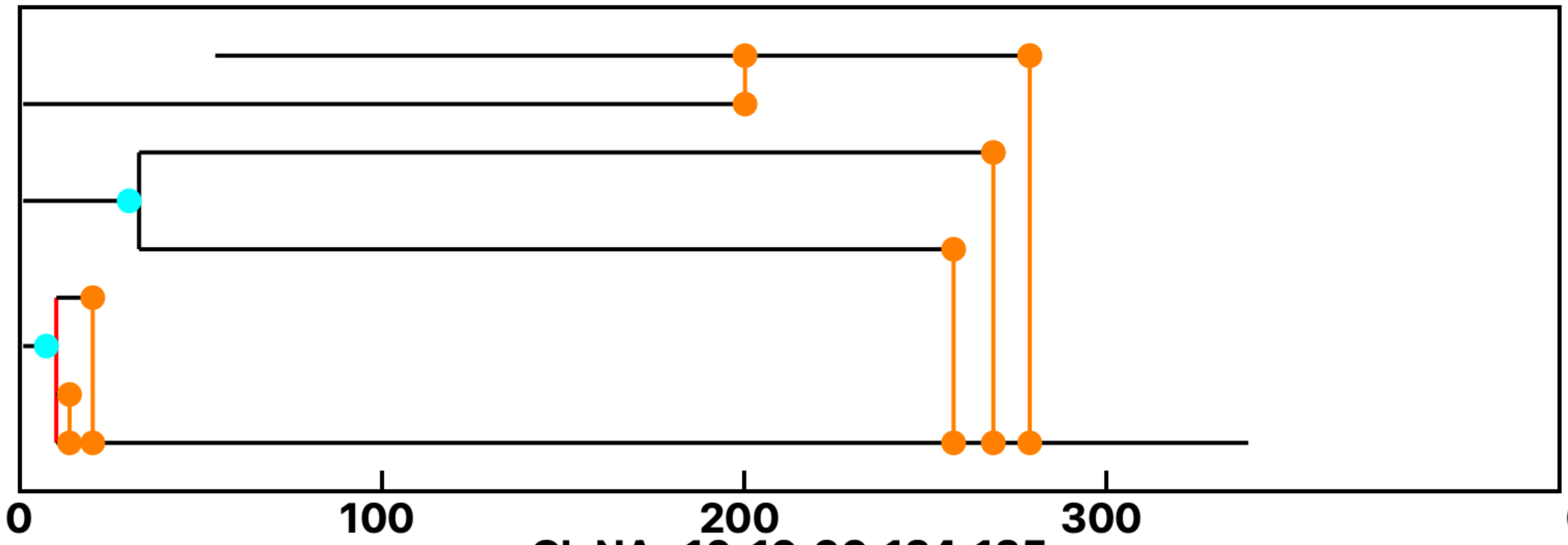

GlcNAc 10-16, 99, 134, 135

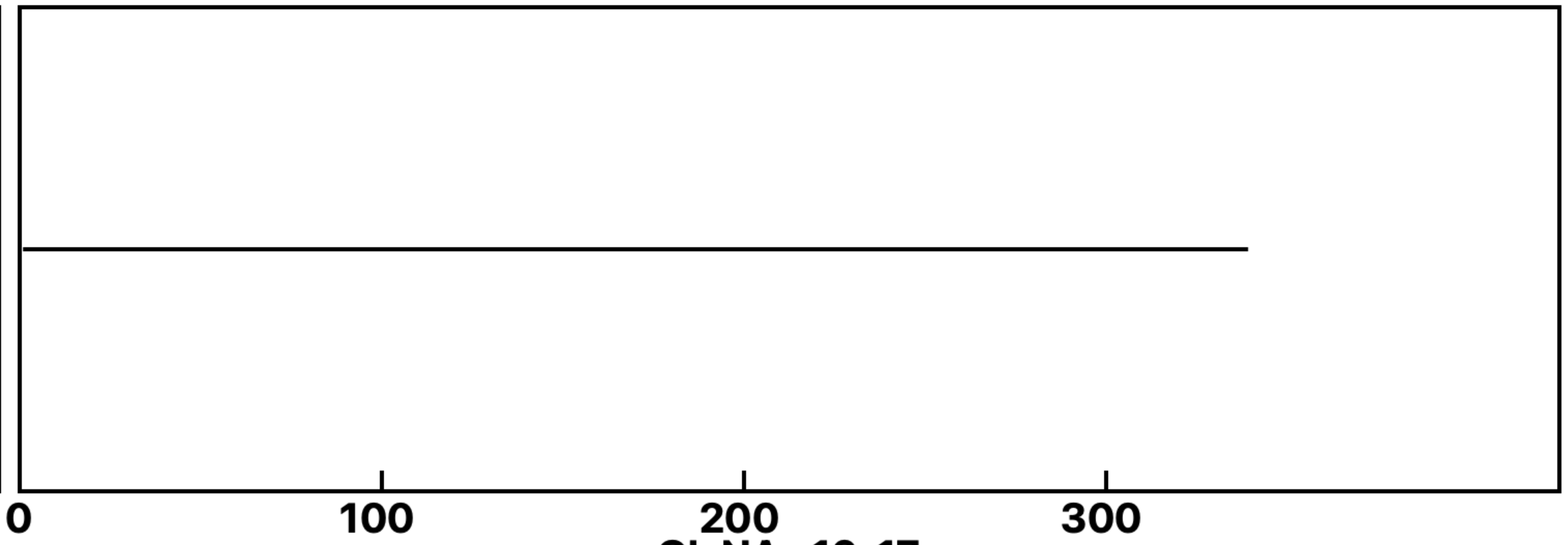

GlcNAc 10-17

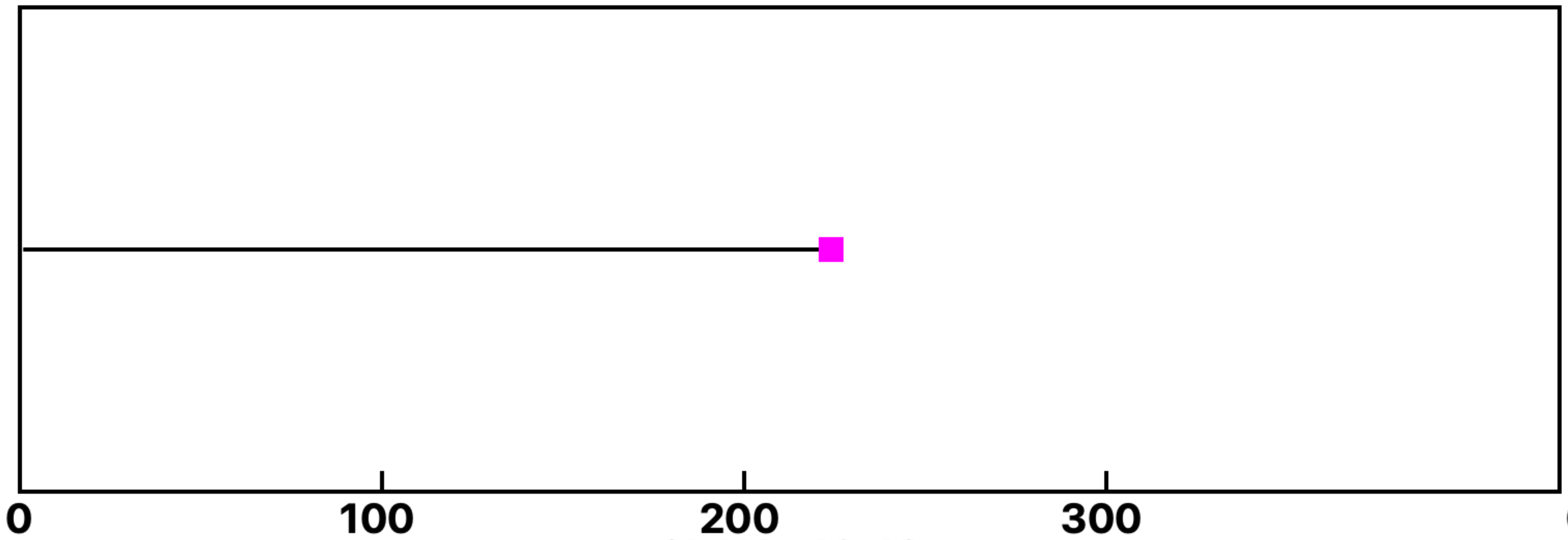

GlcNAc 10-18

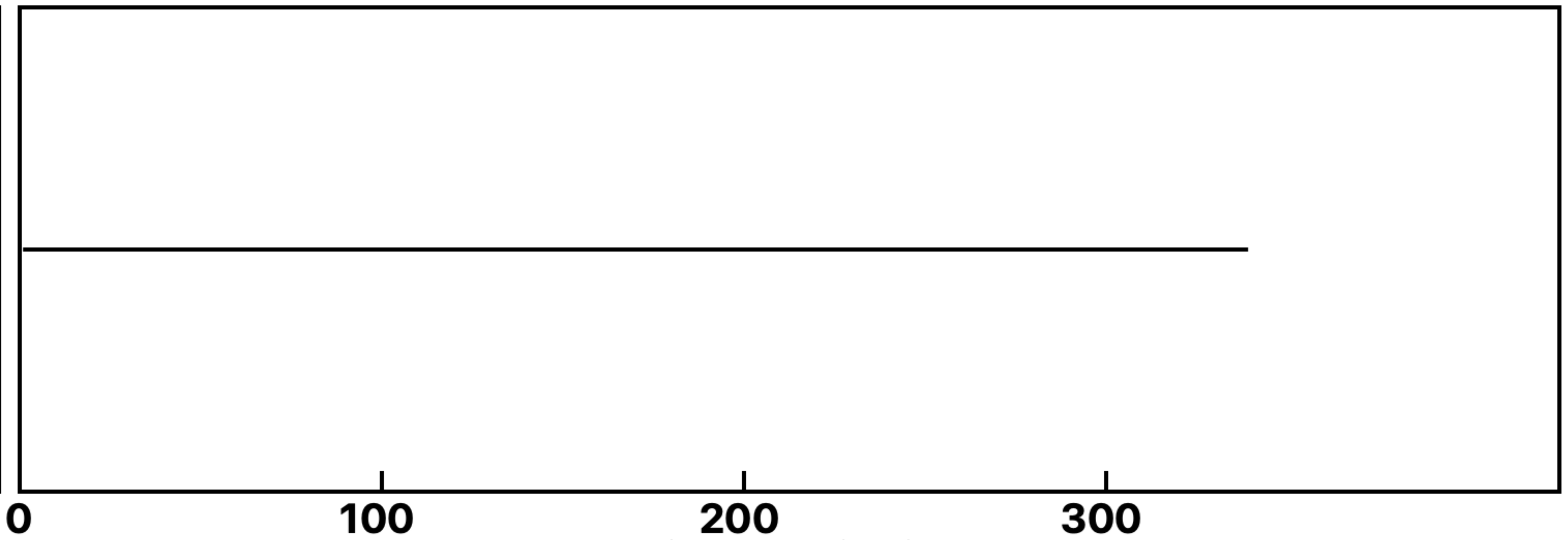

GlcNAc 10-19

**Analysis: Wild type-Dose, Treat.: GlcNAc 10, Cell: Myoblasts**

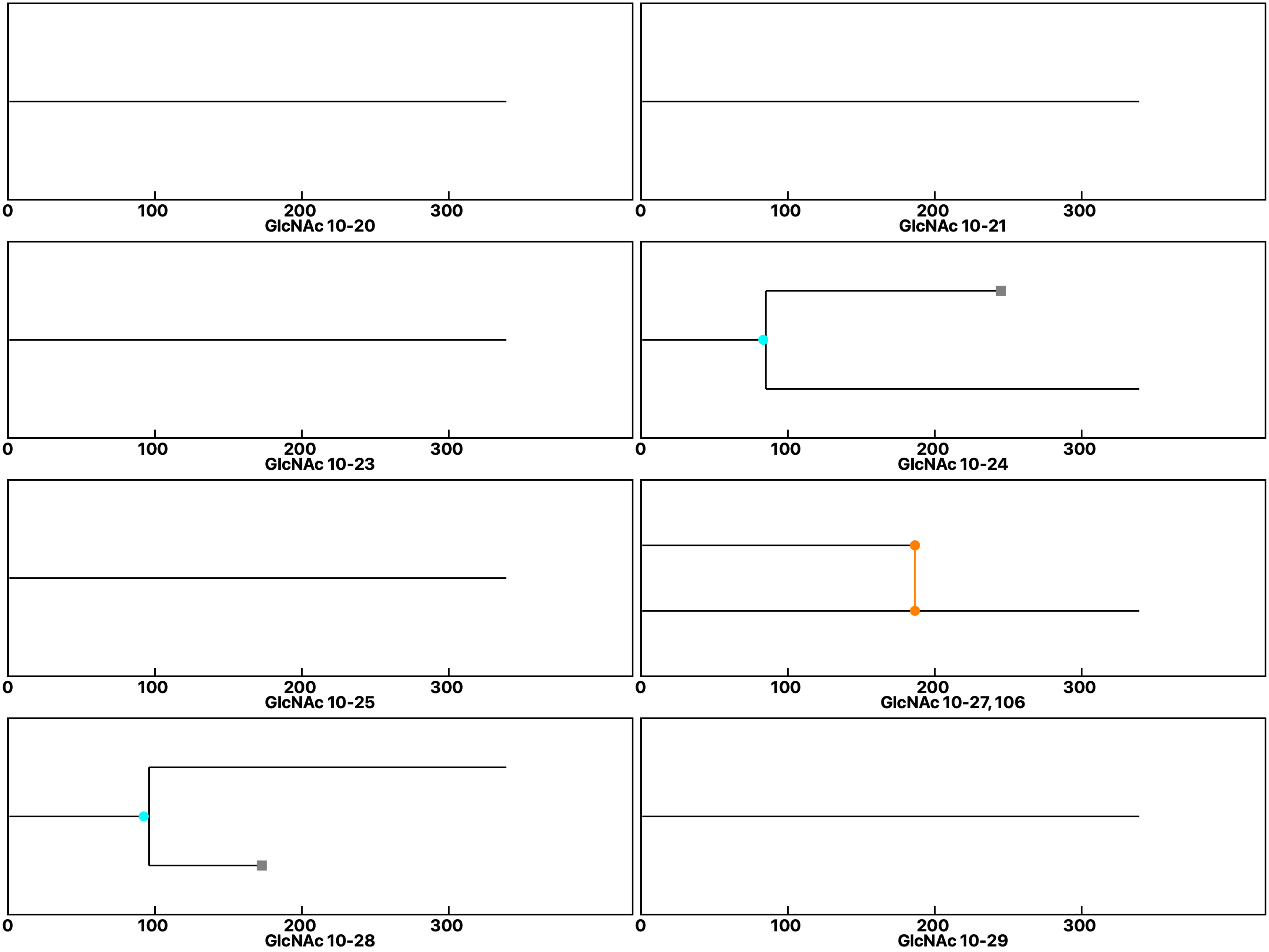

**Analysis: Wild type-Dose, Treat.: GlcNAc 10, Cell: Myoblasts**

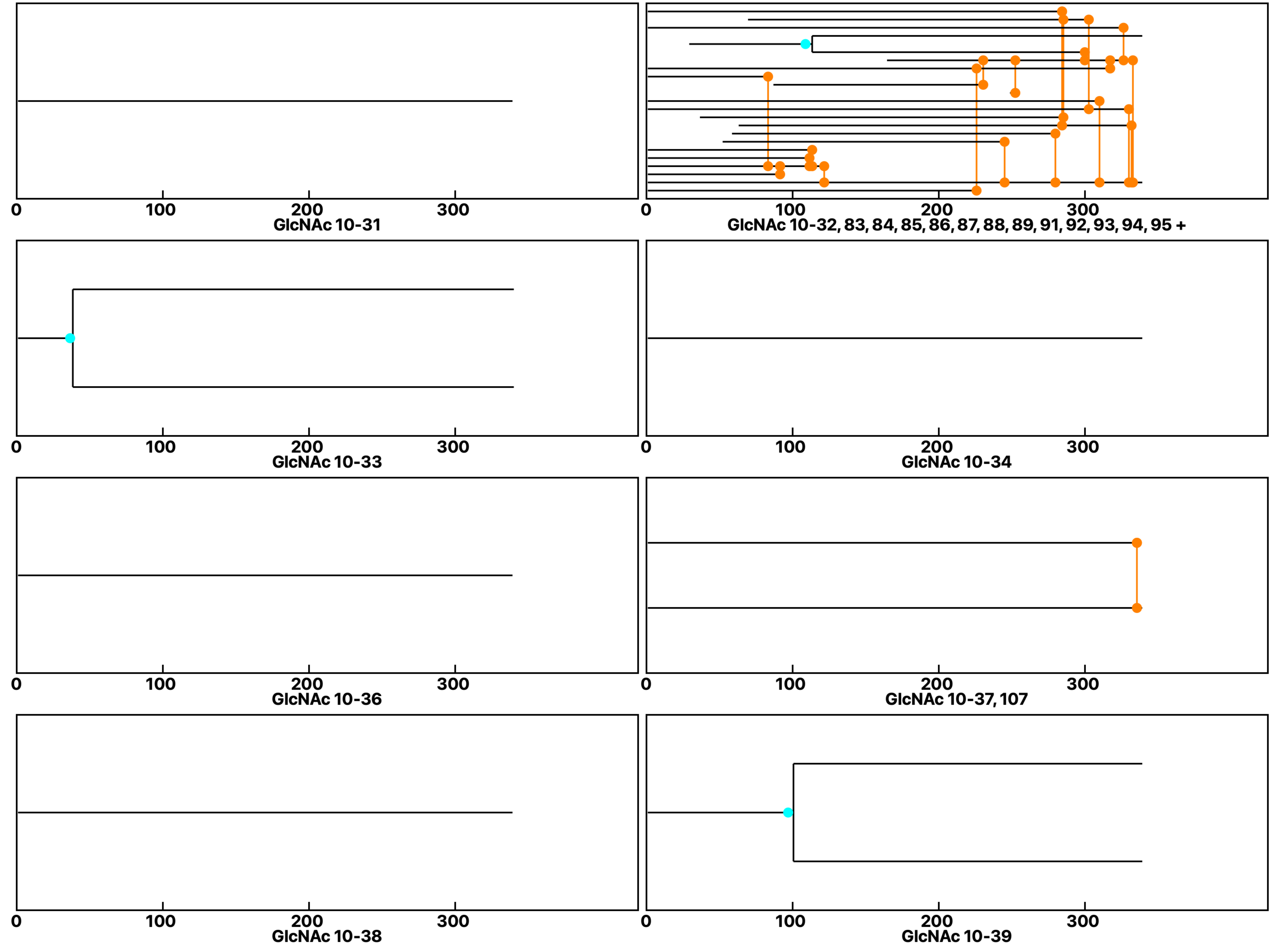

Analysis: Wild type-Dose, Treat.: GlcNAc 10, Cell: Myoblasts

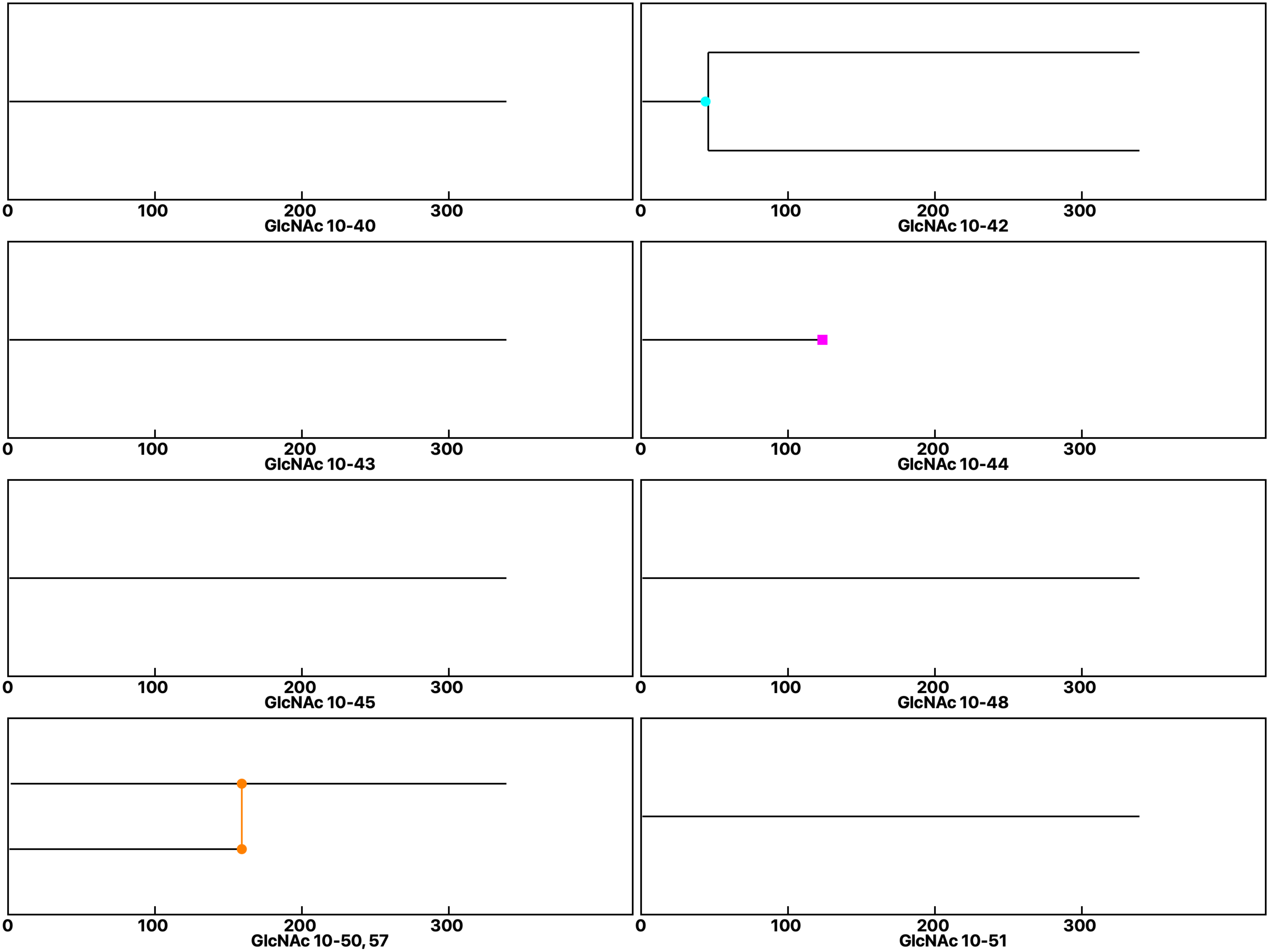

**Analysis: Wild type-Dose, Treat.: GlcNAc 10, Cell: Myoblasts**

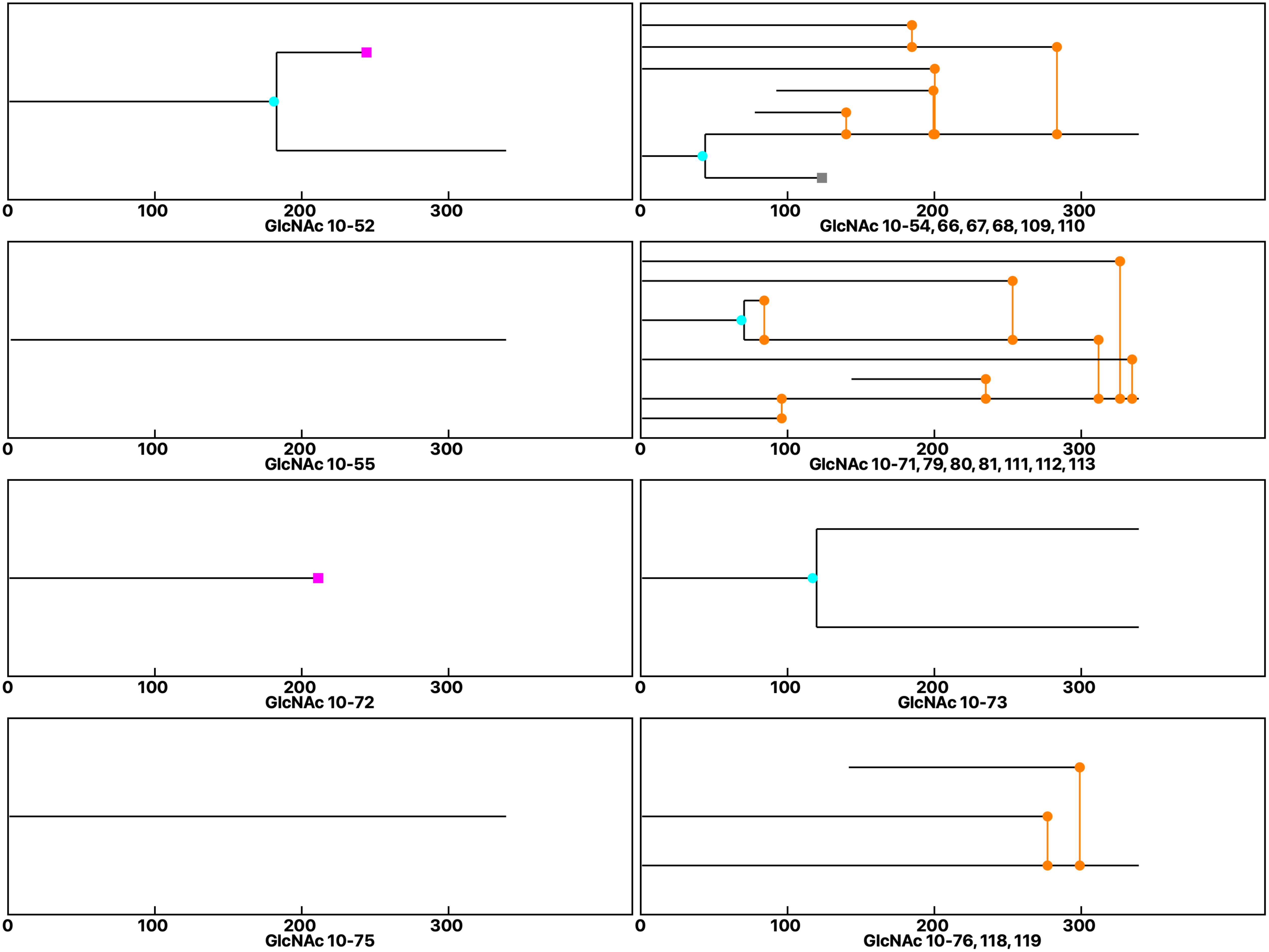
