## Supplemental data 6 for "N-Acetylglucosamine Facilitates Coordinated Myoblast Flow, Forming the Foundation for Efficient Myogenesis"

**Analysis: Wild type Gal-3-GlcNAc, Treat.: +Gal-3, Cell: Myoblasts**

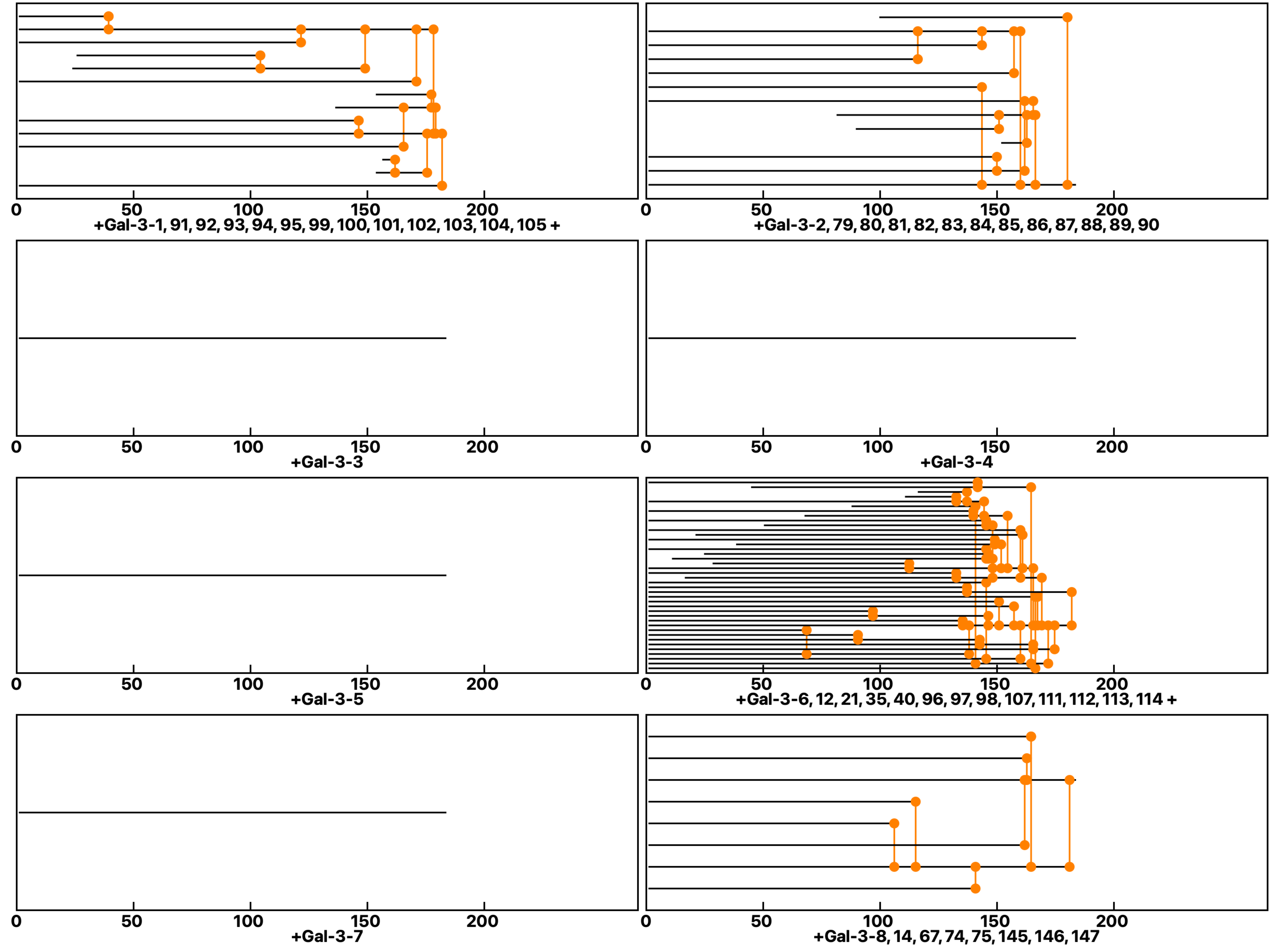

Analysis: Wild type Gal-3-GlcNAc, Treat.: +Gal-3, Cell: Myoblasts

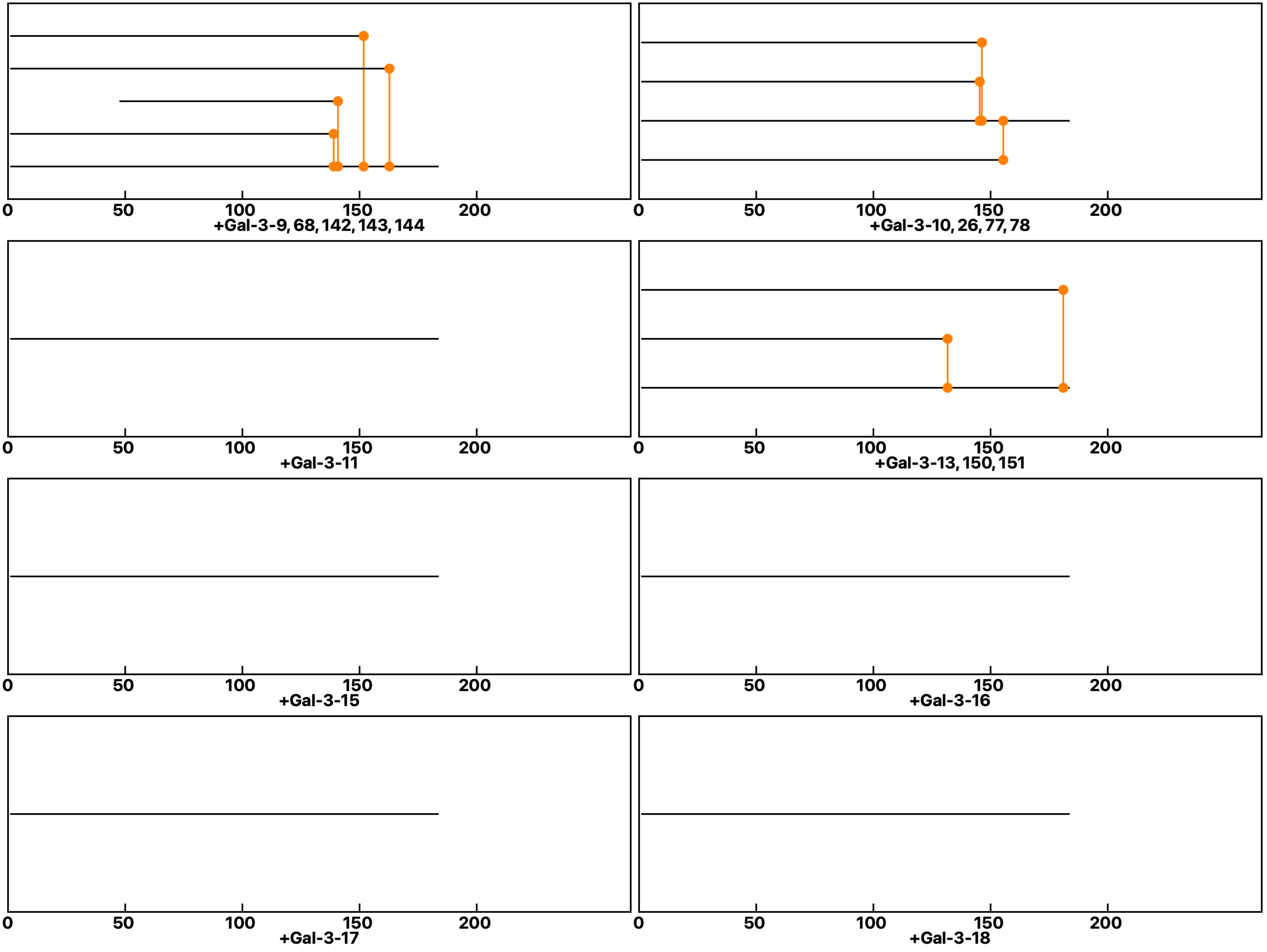

Analysis: Wild type Gal-3-GlcNAc, Treat.: +Gal-3, Cell: Myoblasts

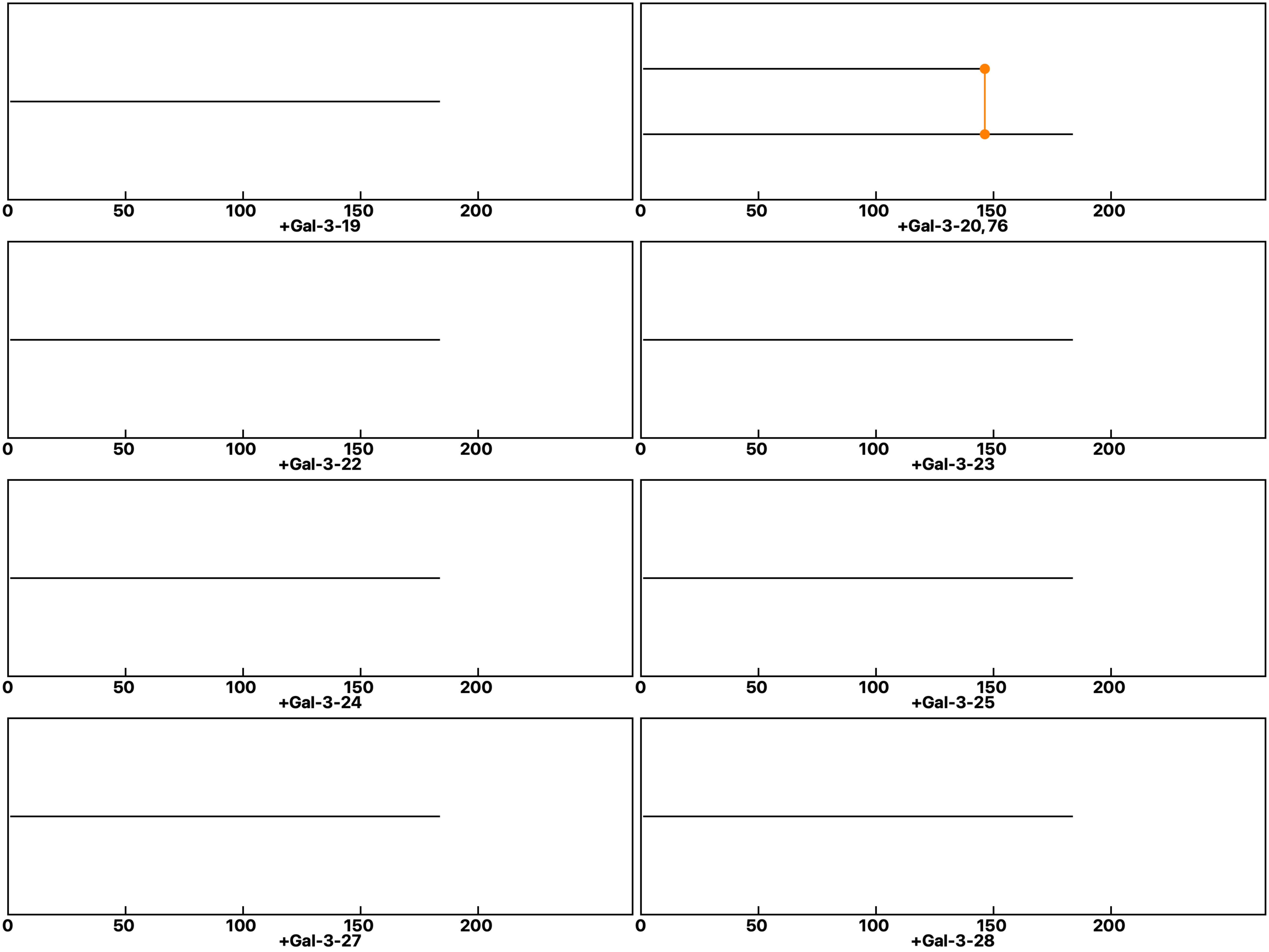

**Analysis: Wild type Gal-3-GlcNAc, Treat.: +Gal-3, Cell: Myoblasts**

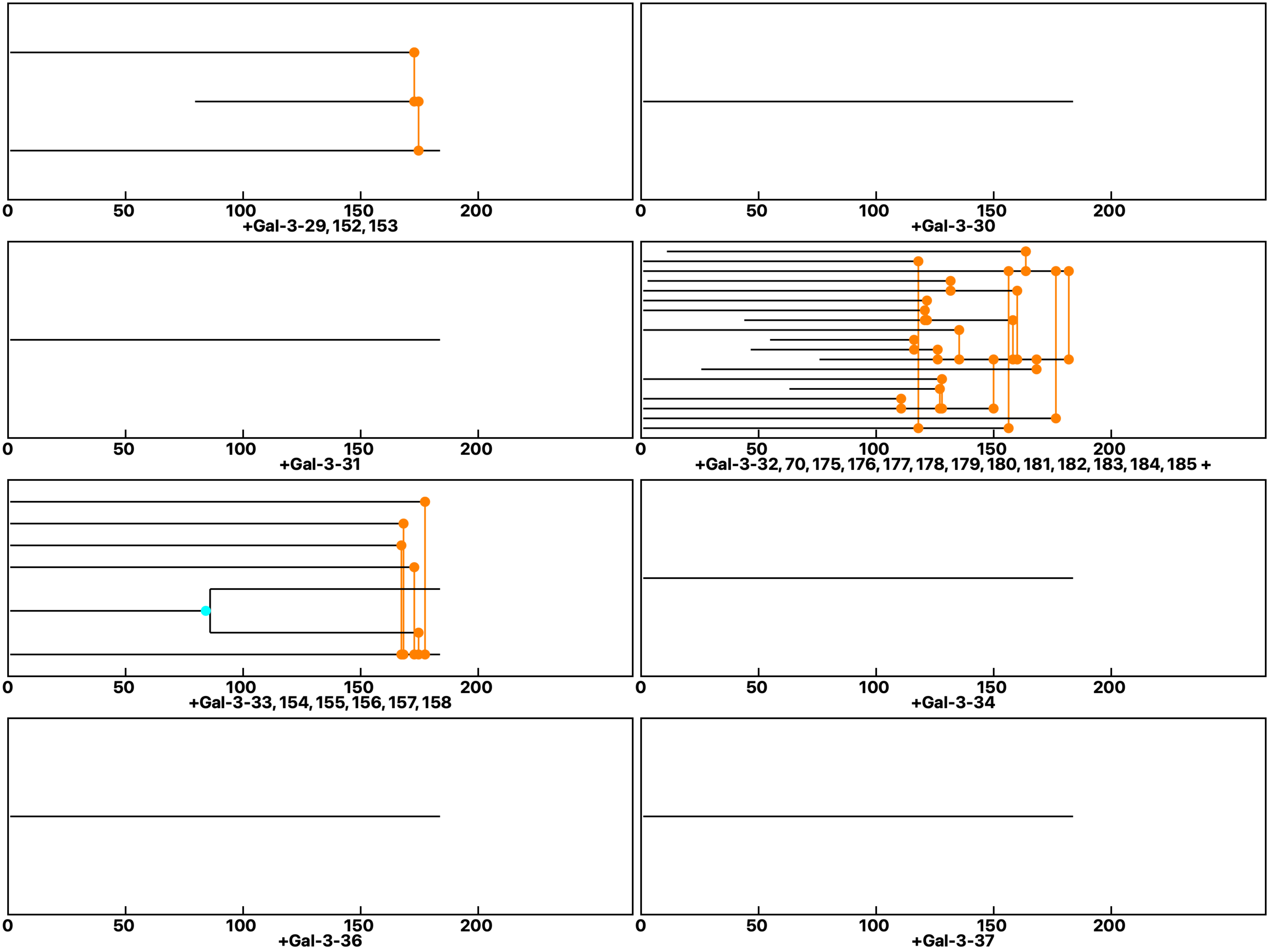

**Analysis: Wild type Gal-3-GlcNAc, Treat.: +Gal-3, Cell: Myoblasts**

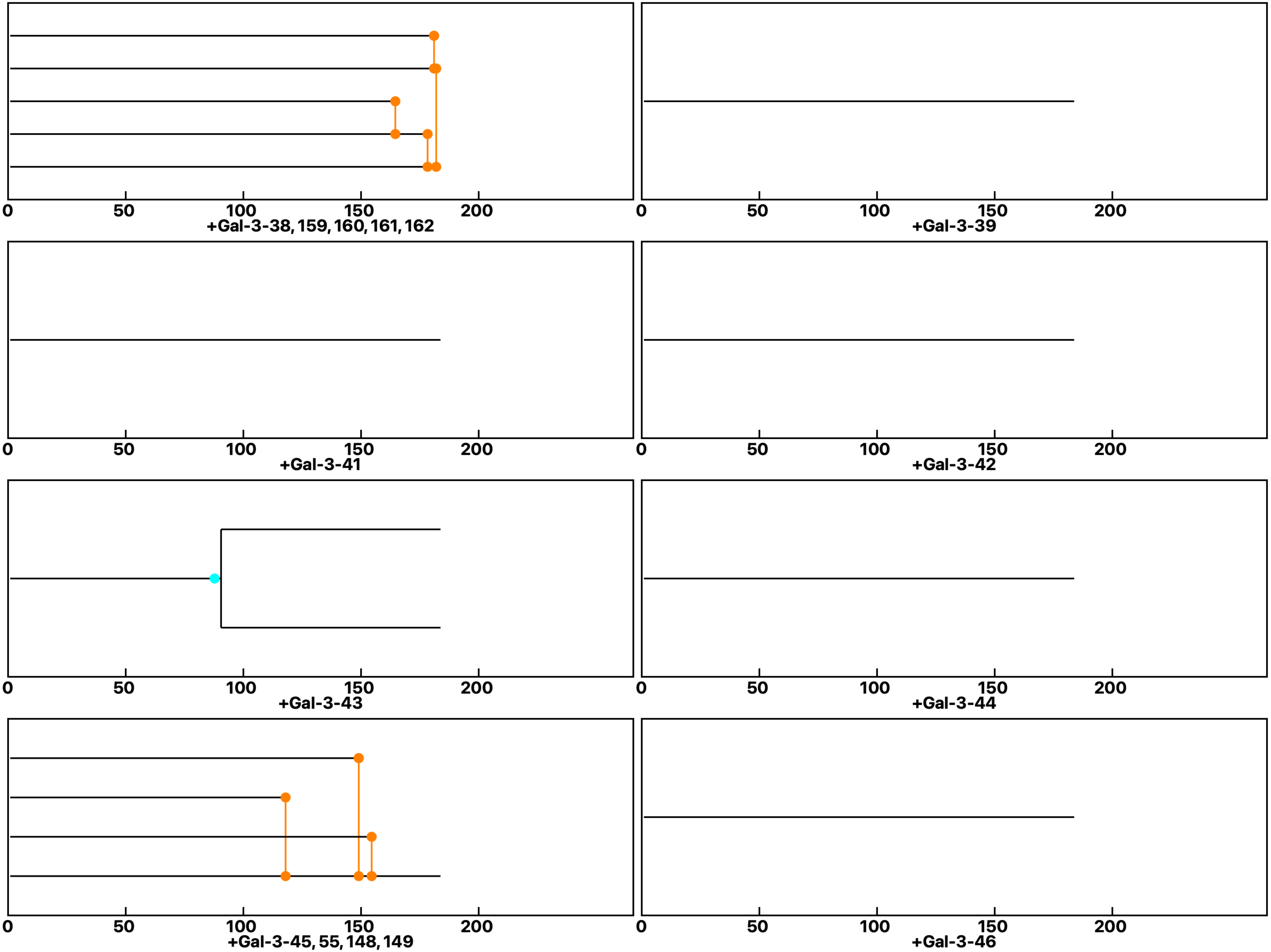

**Analysis: Wild type Gal-3-GlcNAc, Treat.: +Gal-3, Cell: Myoblasts**

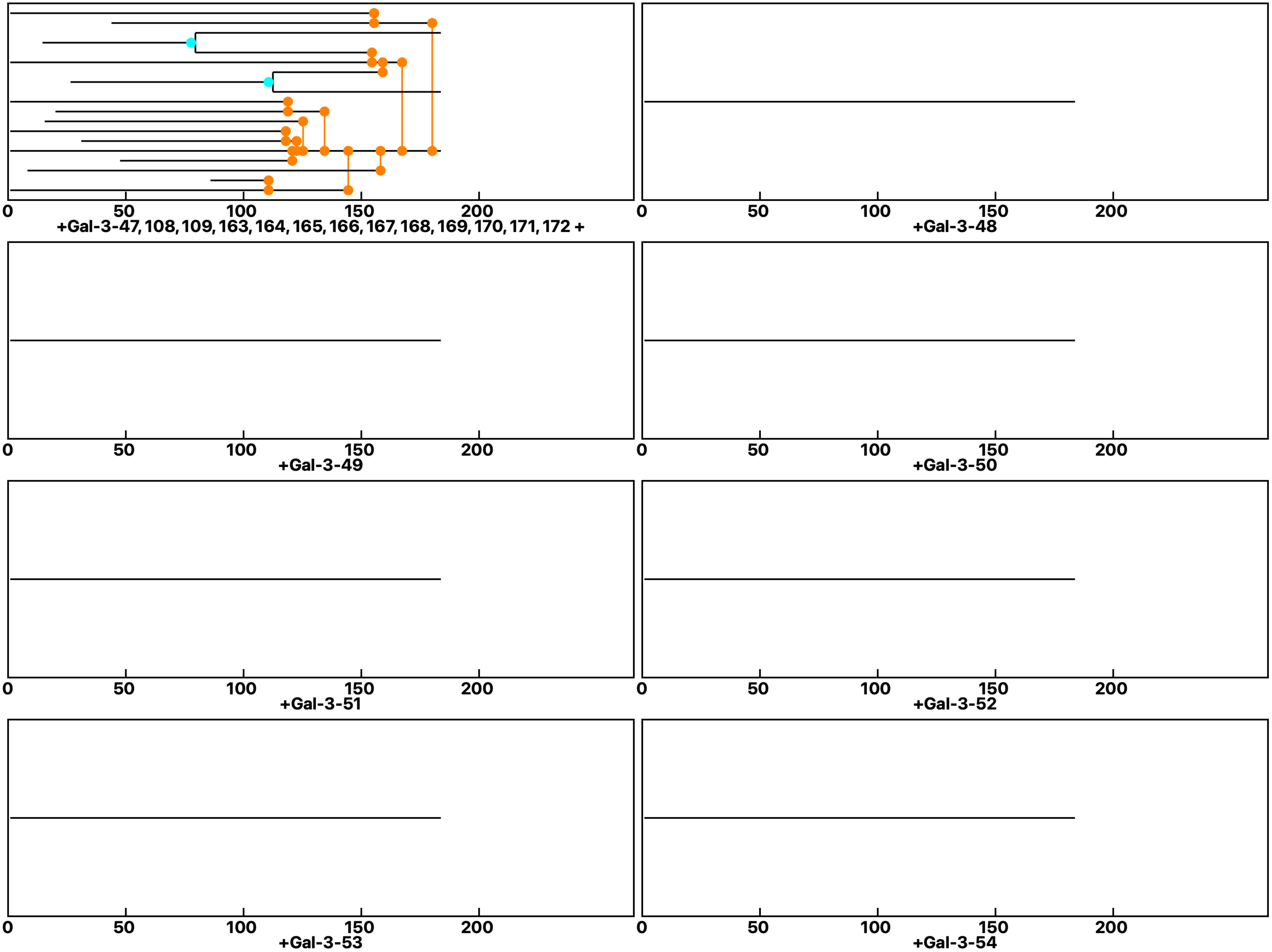

**Analysis: Wild type Gal-3-GlcNAc, Treat.: +Gal-3, Cell: Myoblasts**

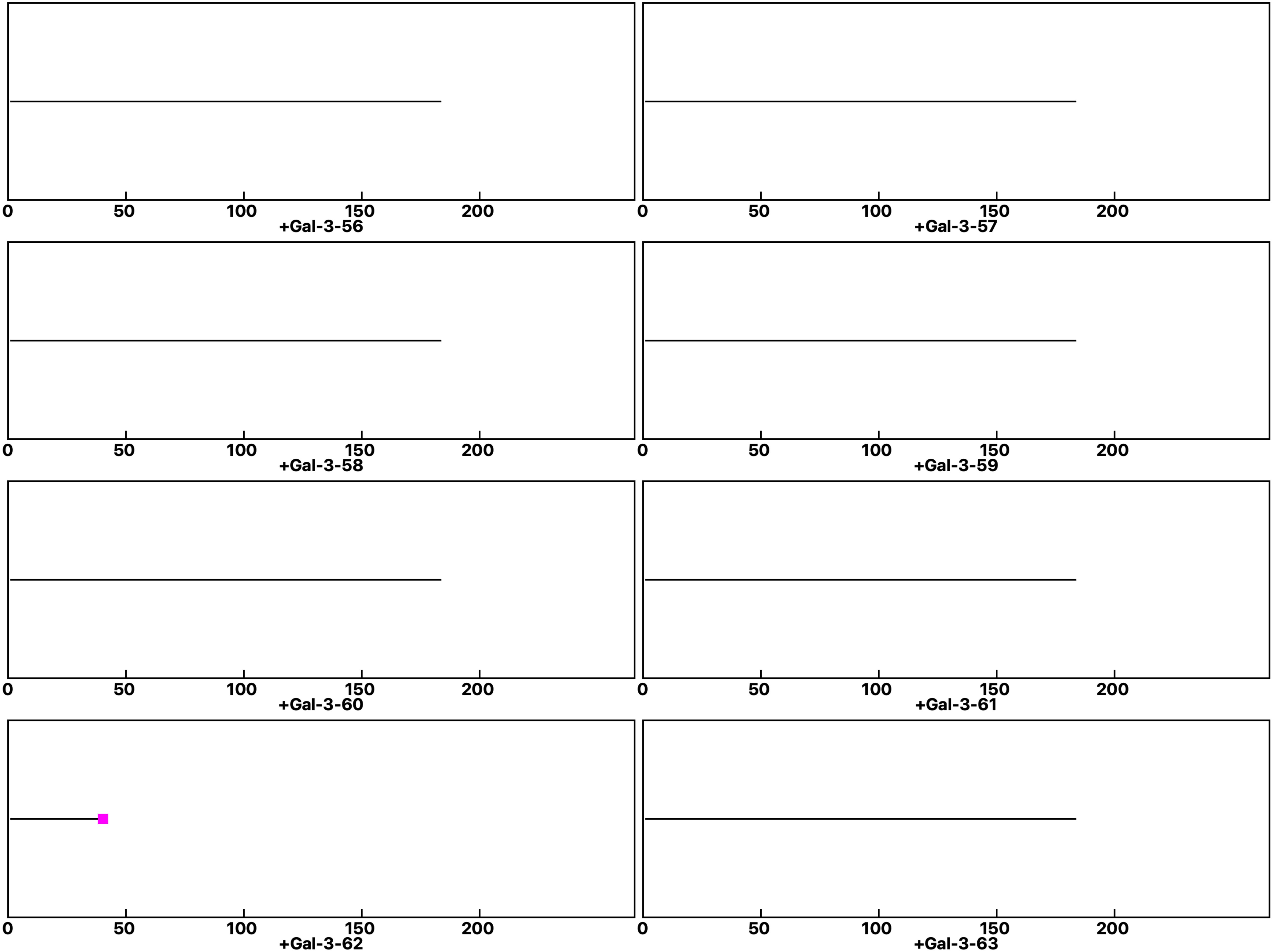

**Analysis: Wild type Gal-3-GlcNAc, Treat.: +Gal-3, Cell: Myoblasts**

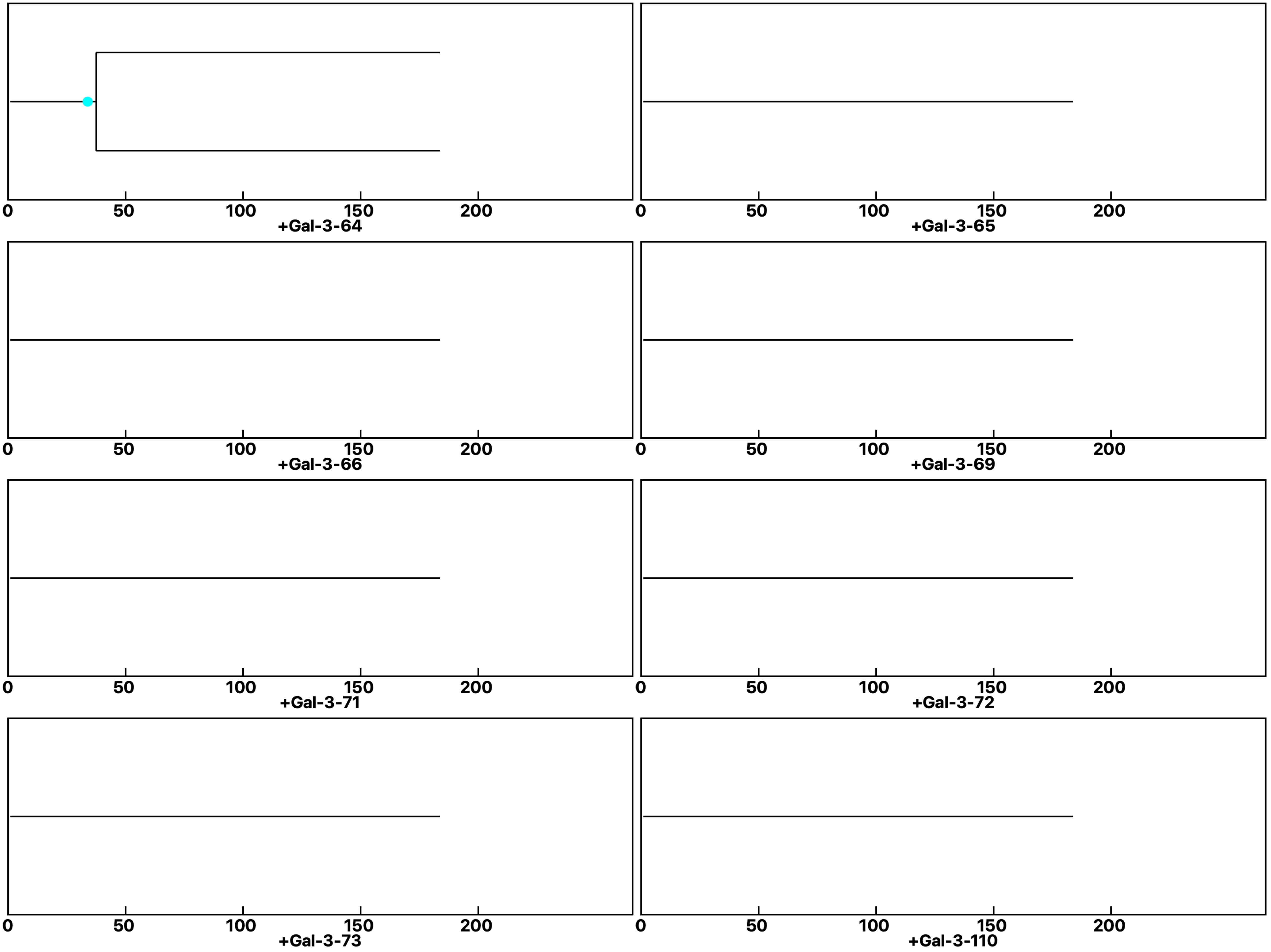
