## Supplemental data 7 for "N-Acetylglucosamine Facilitates Coordinated Myoblast Flow, Forming the Foundation for Efficient Myogenesis"

### Analysis: Wild type Gal-3-GlcNAc, Treat.: +GlcNAc, Cell: Myoblasts

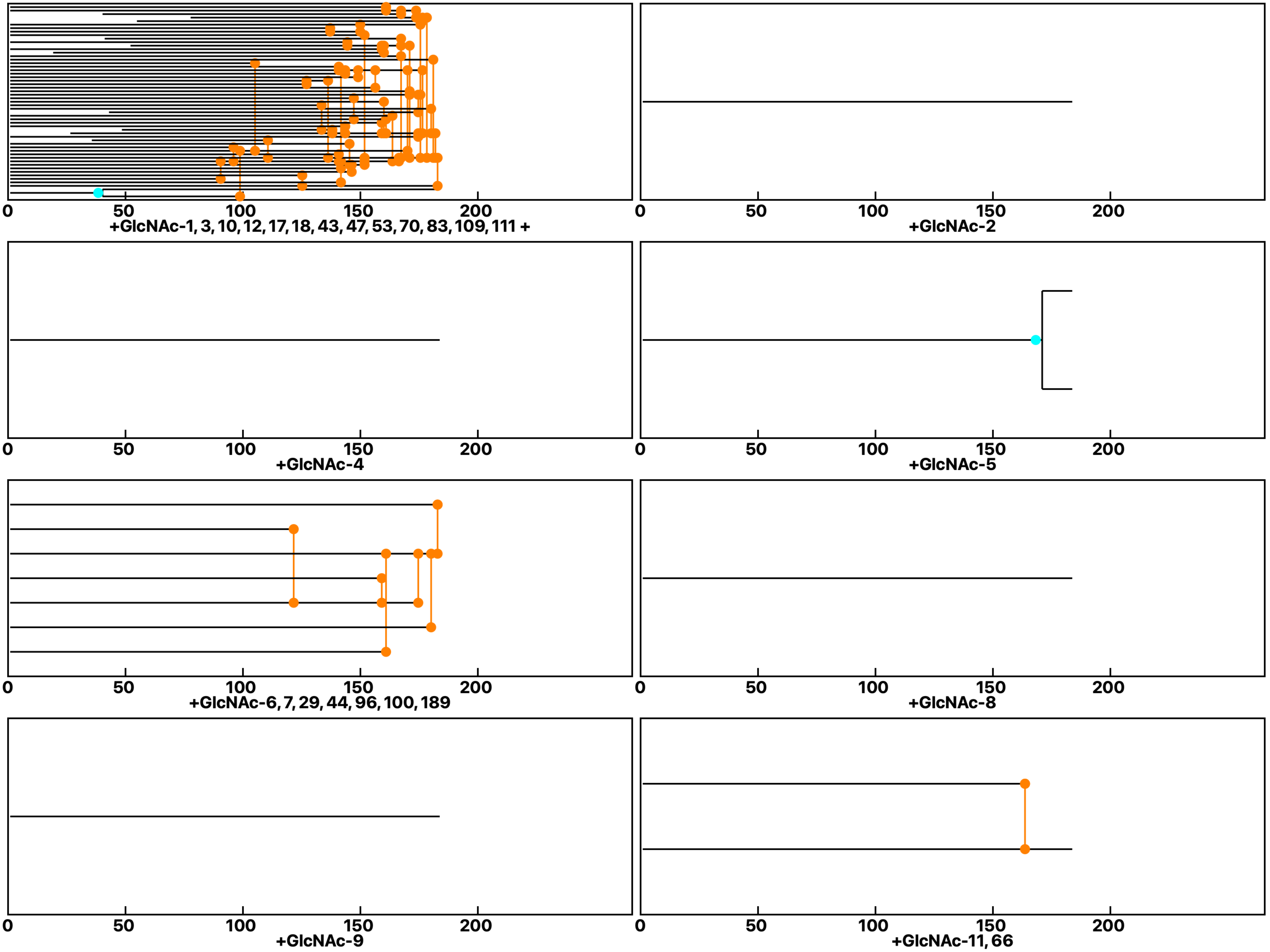

**Analysis: Wild type Gal-3-GlcNAc, Treat.: +GlcNAc, Cell: Myoblasts**

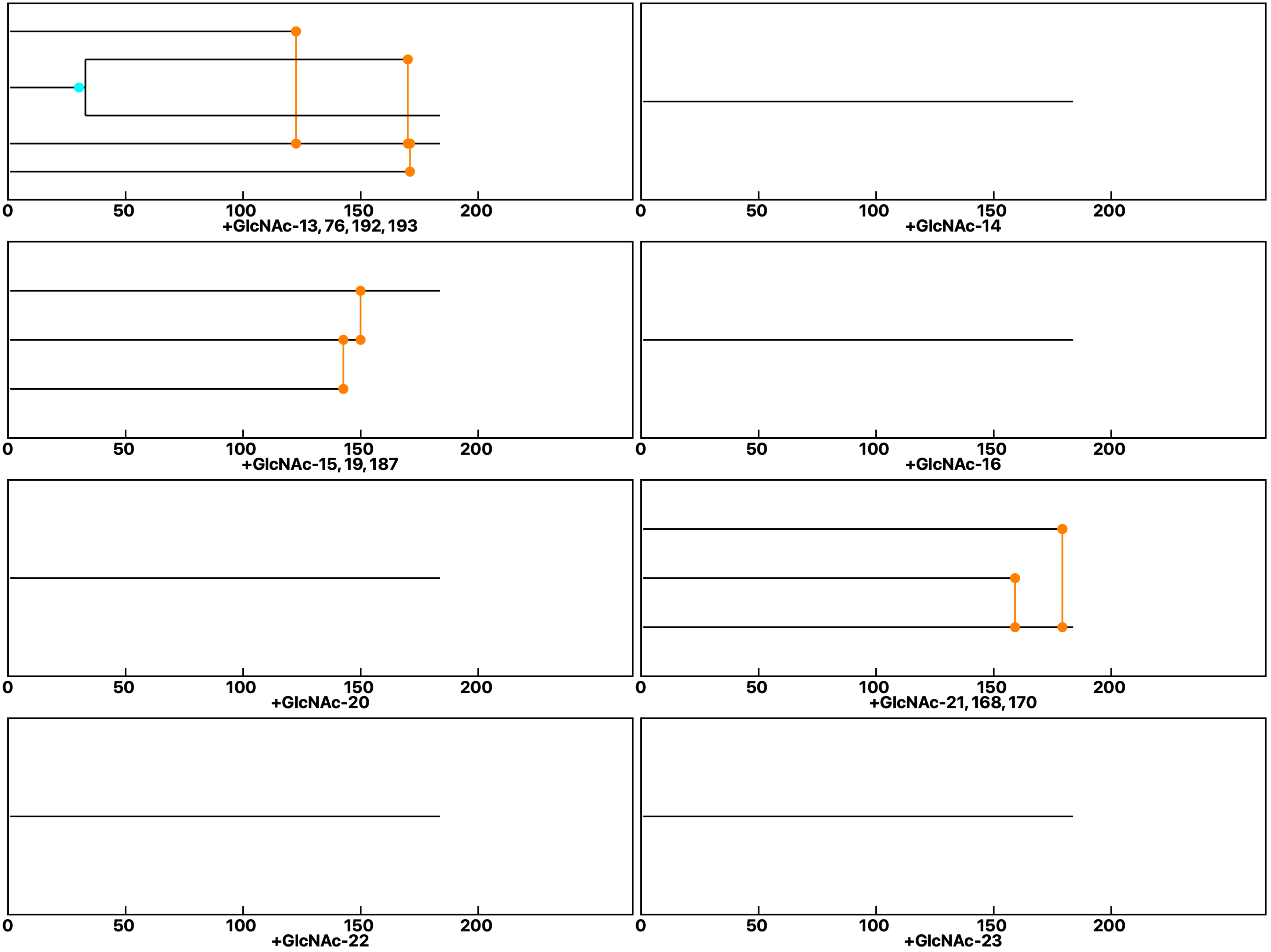

**Analysis: Wild type Gal-3-GlcNAc, Treat.: +GlcNAc, Cell: Myoblasts**

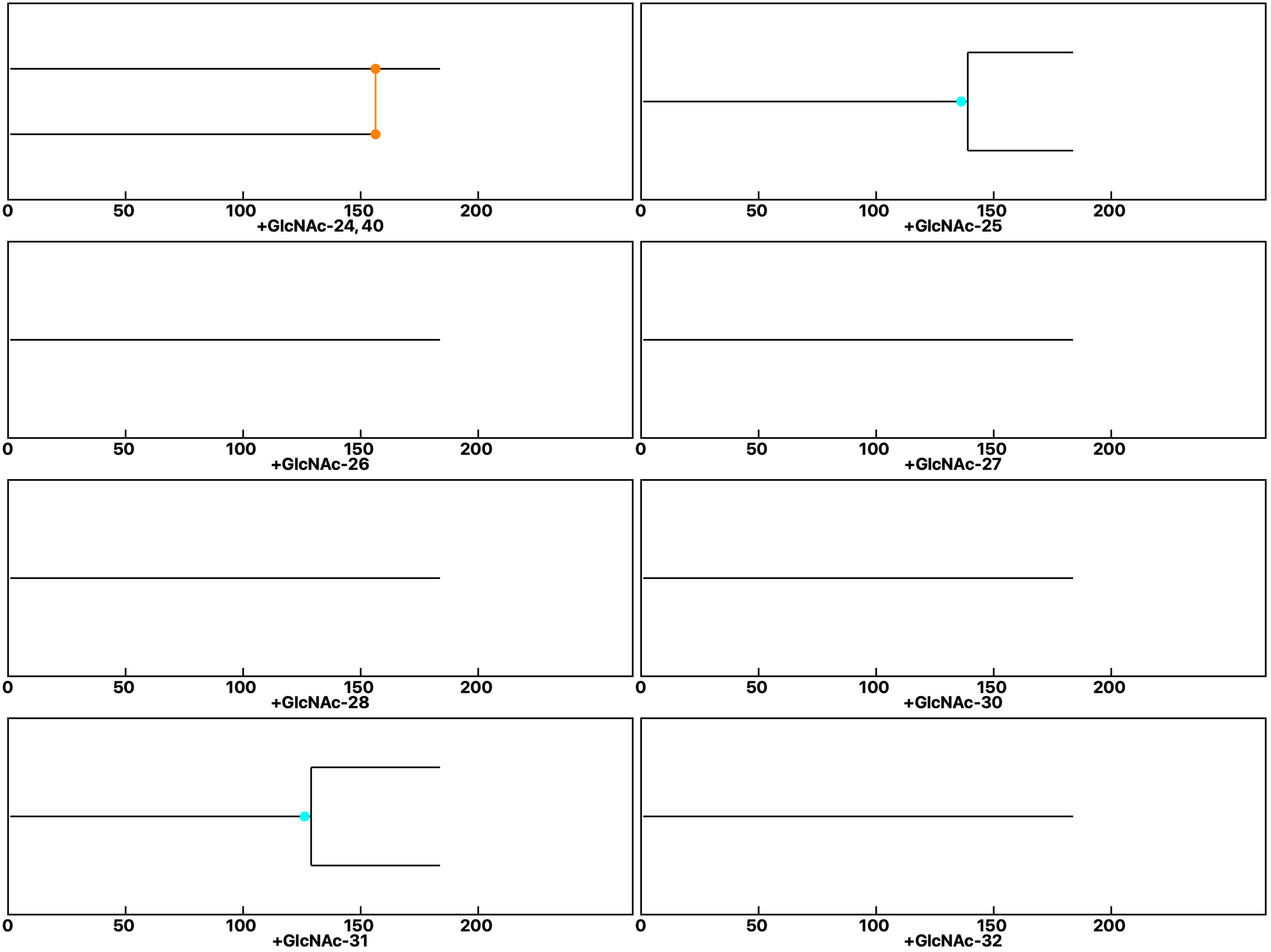

**Analysis: Wild type Gal-3-GlcNAc, Treat.: +GlcNAc, Cell: Myoblasts**

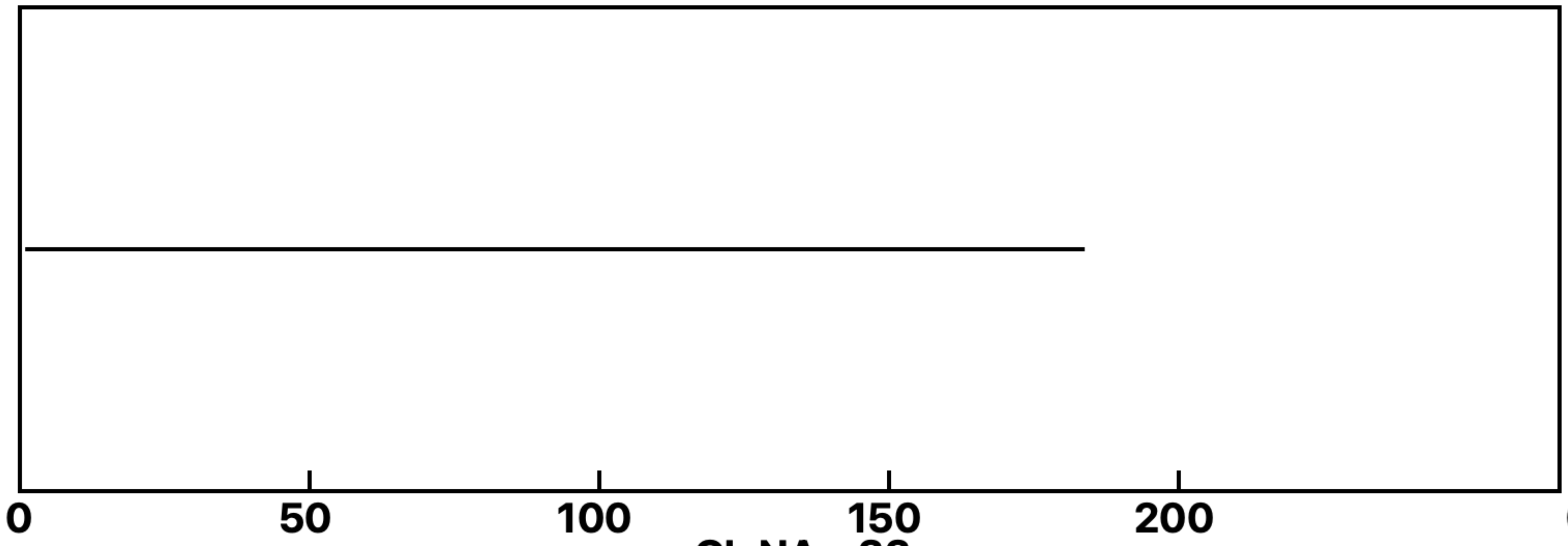

+GlcNAc-33

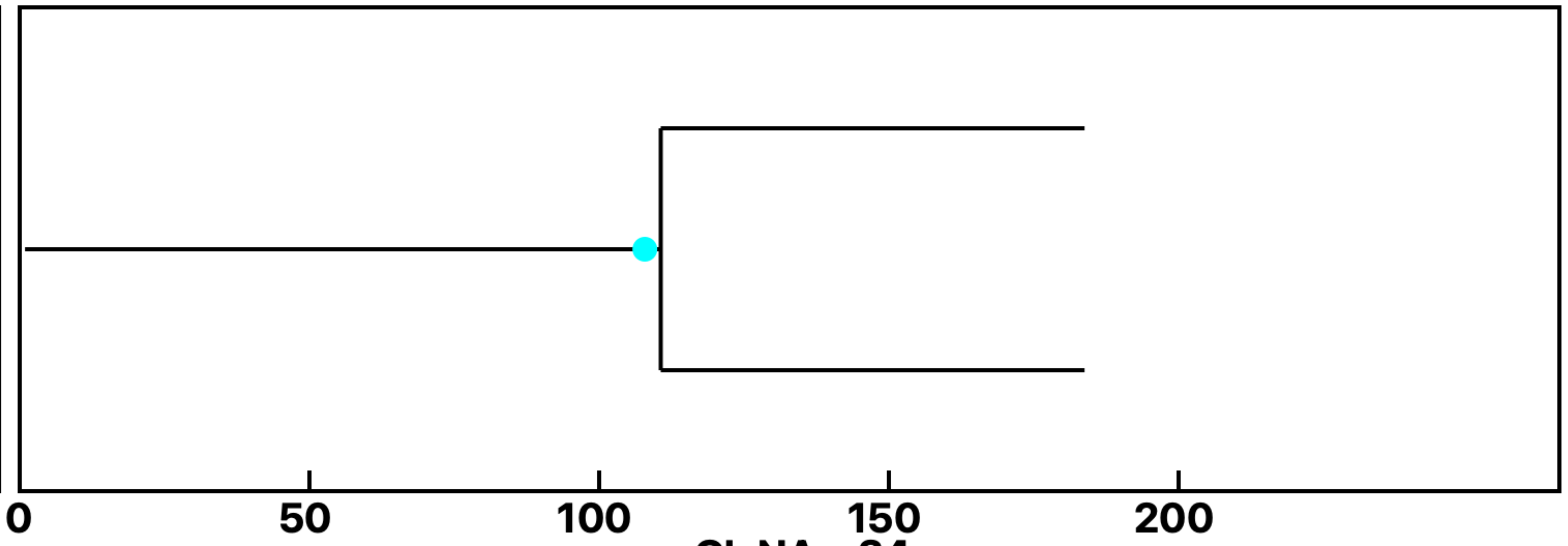

+GlcNAc-34

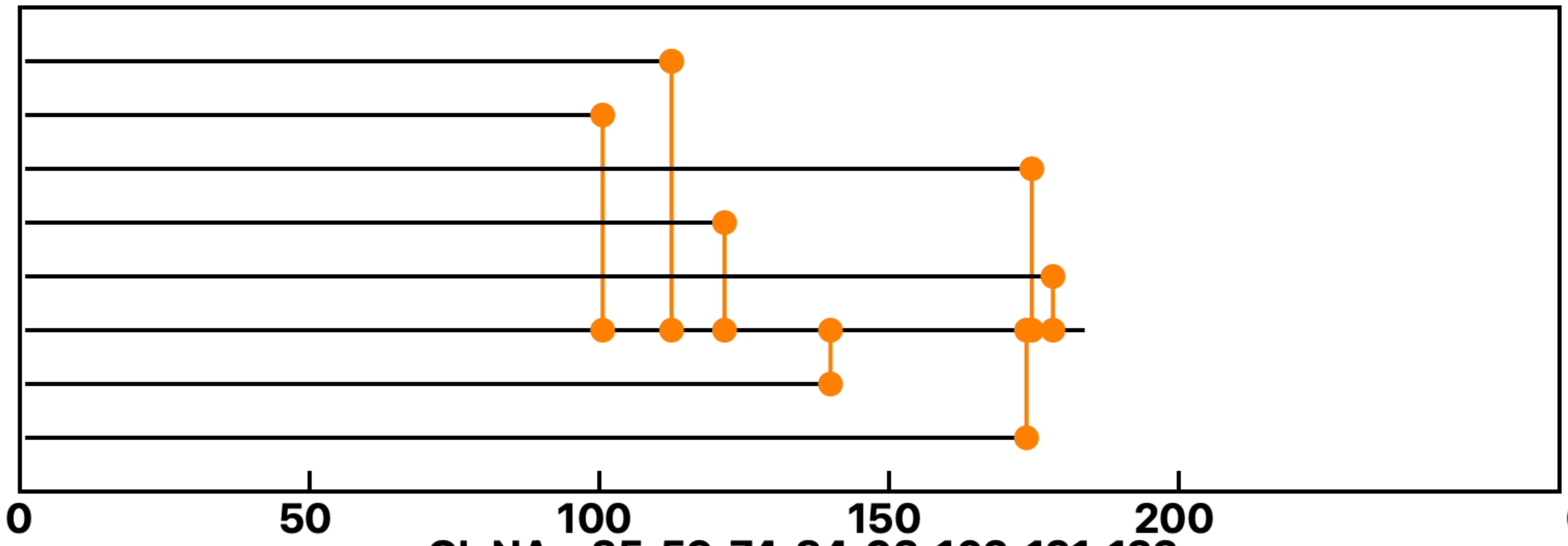

+GlcNAc-35, 59, 74, 84, 93, 102, 121, 122

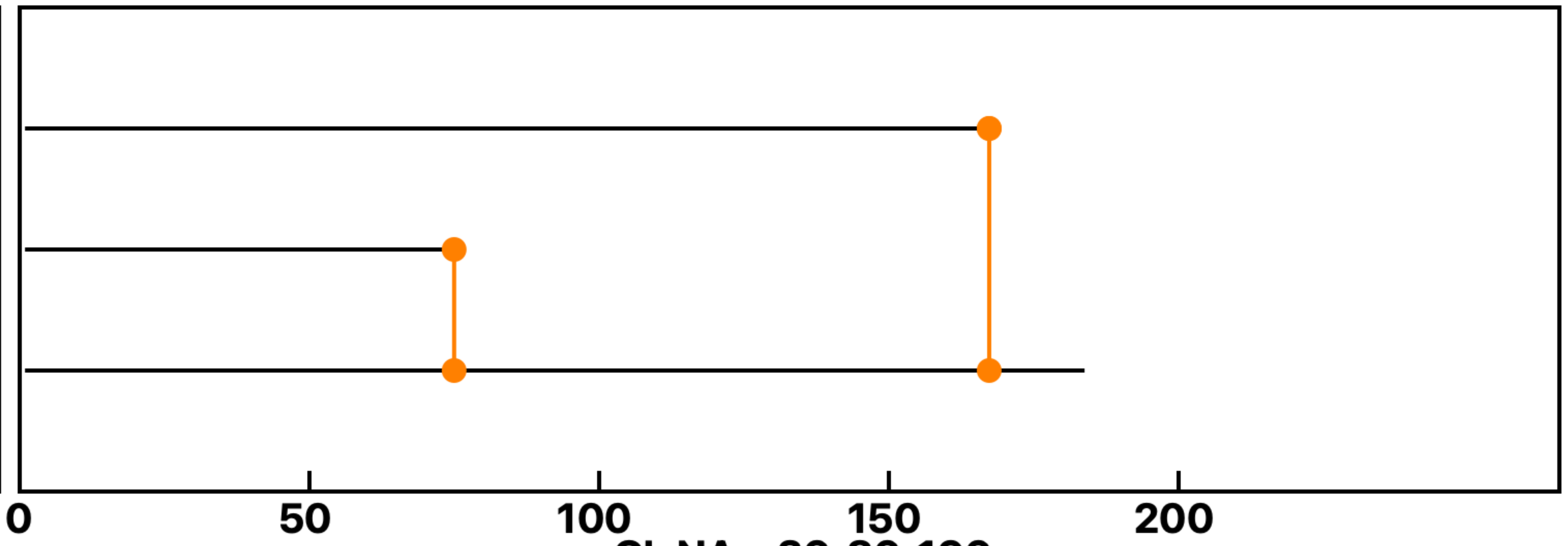

+GlcNAc-36, 86, 196

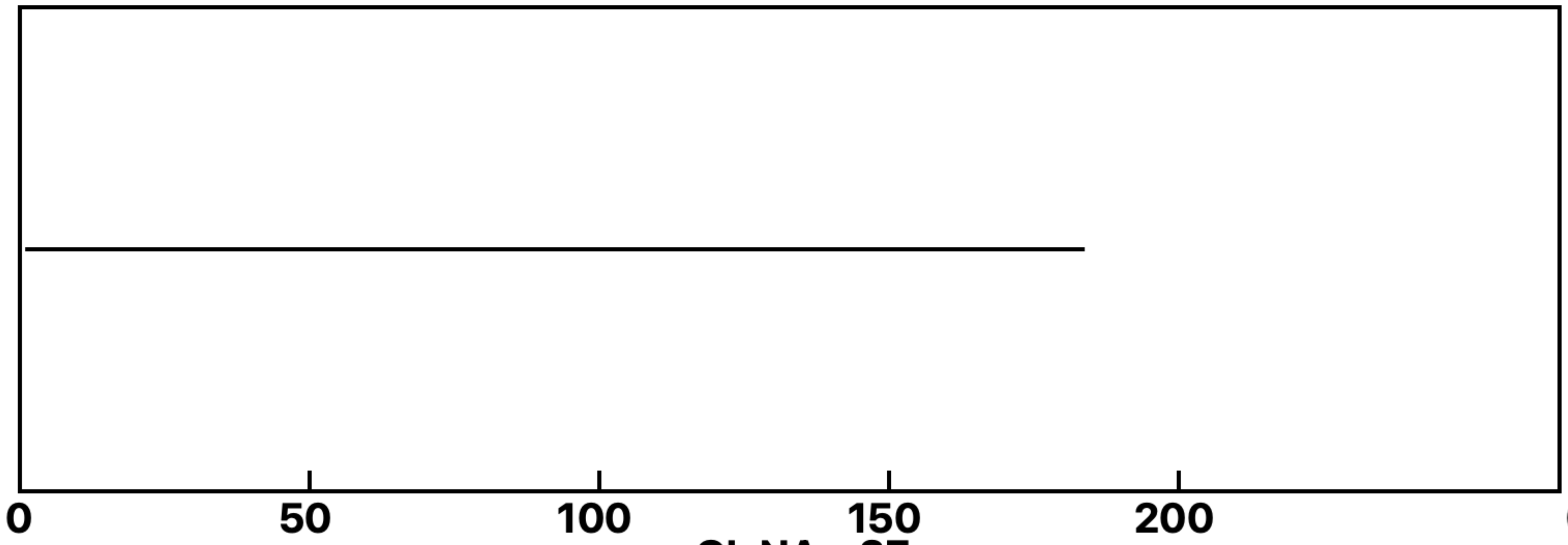

+GlcNAc-37

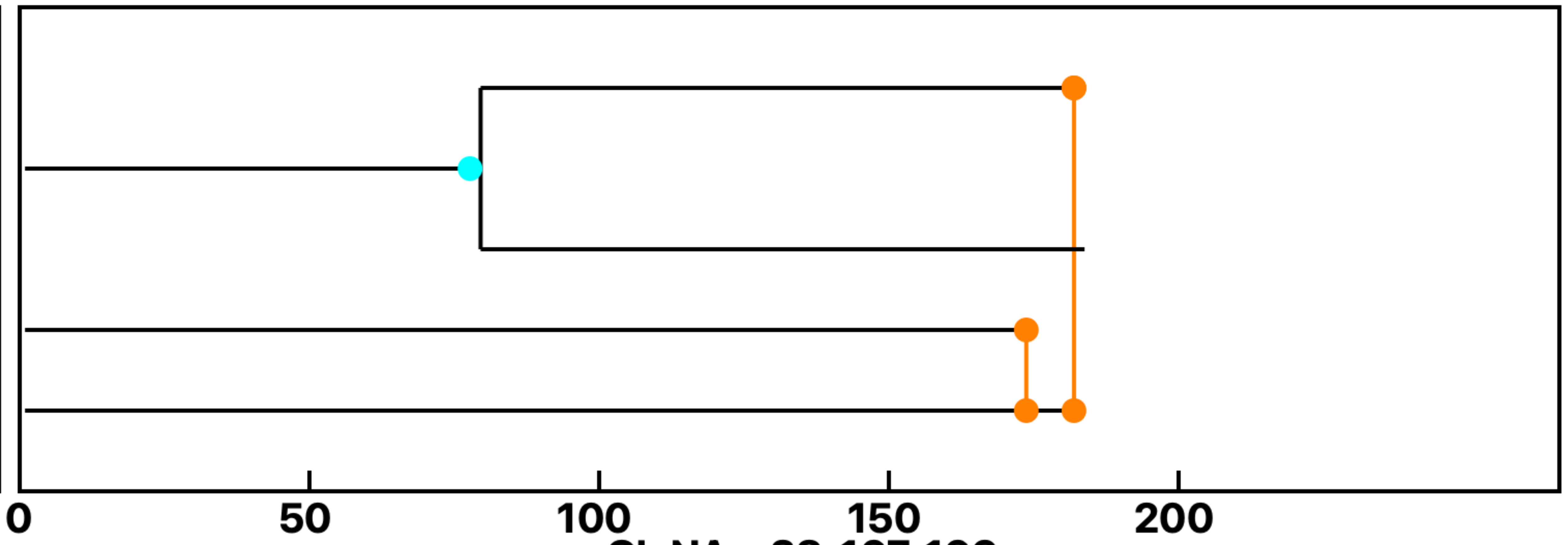

+GlcNAc-38, 167, 169

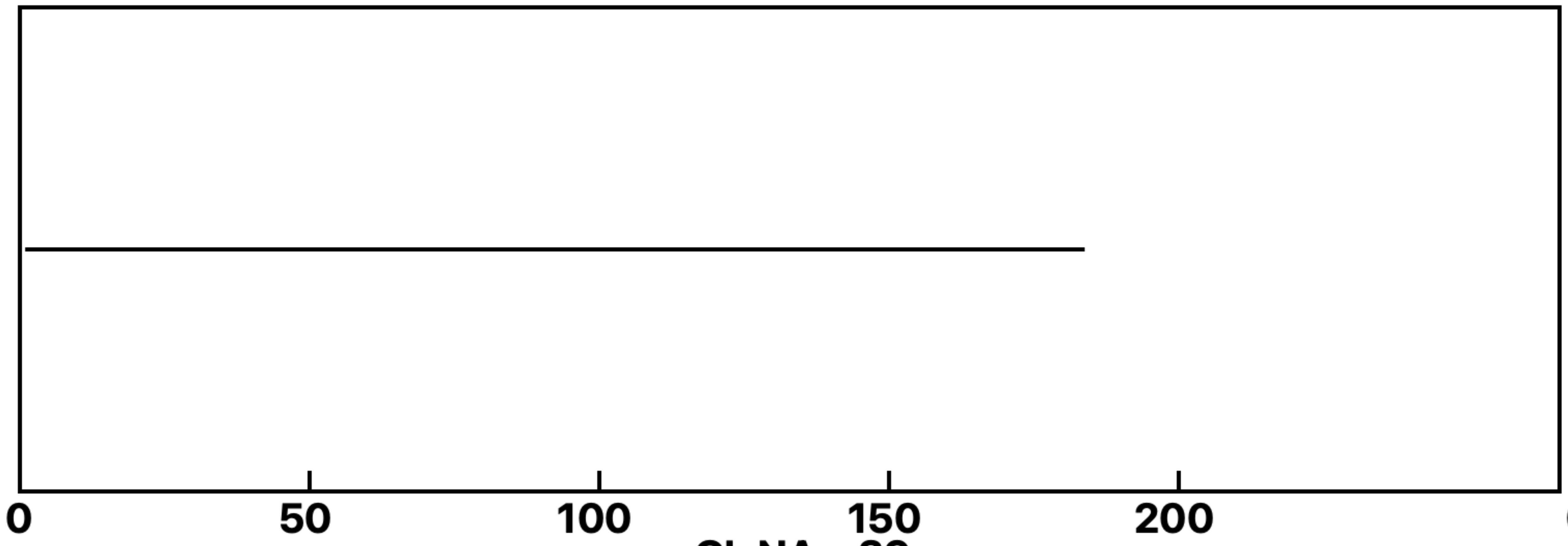

+GlcNAc-39

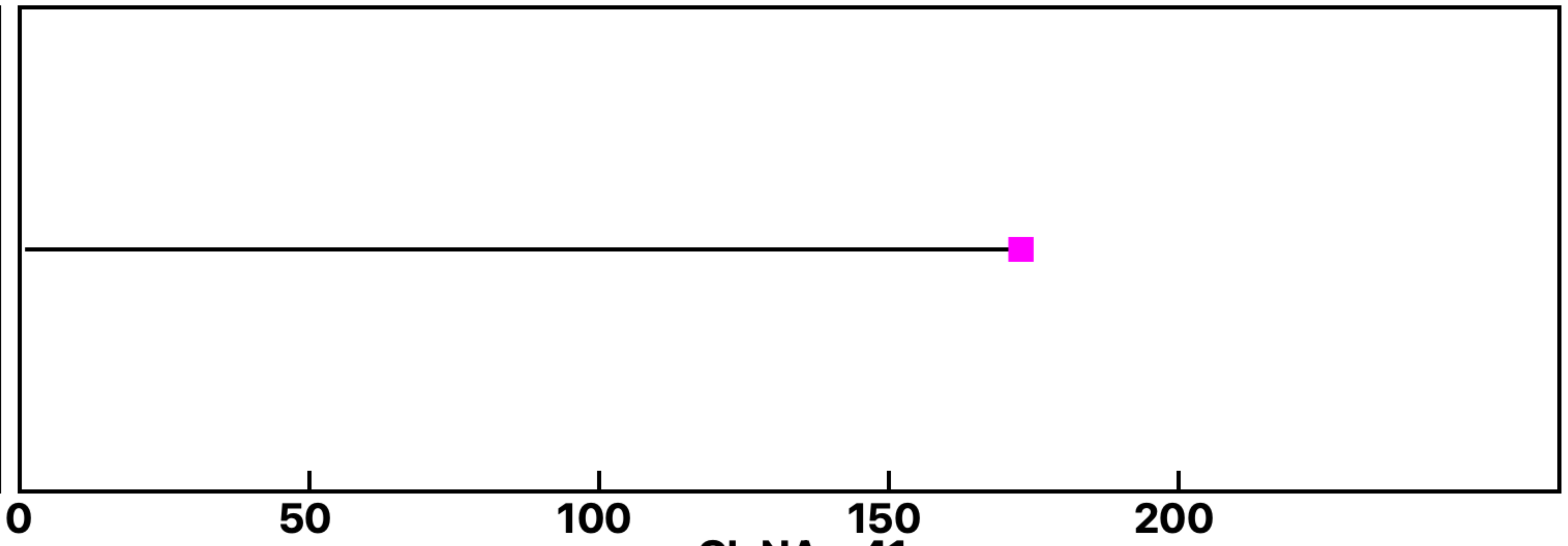

+GlcNAc-41

Analysis: Wild type Gal-3-GlcNAc, Treat.: +GlcNAc, Cell: Myoblasts

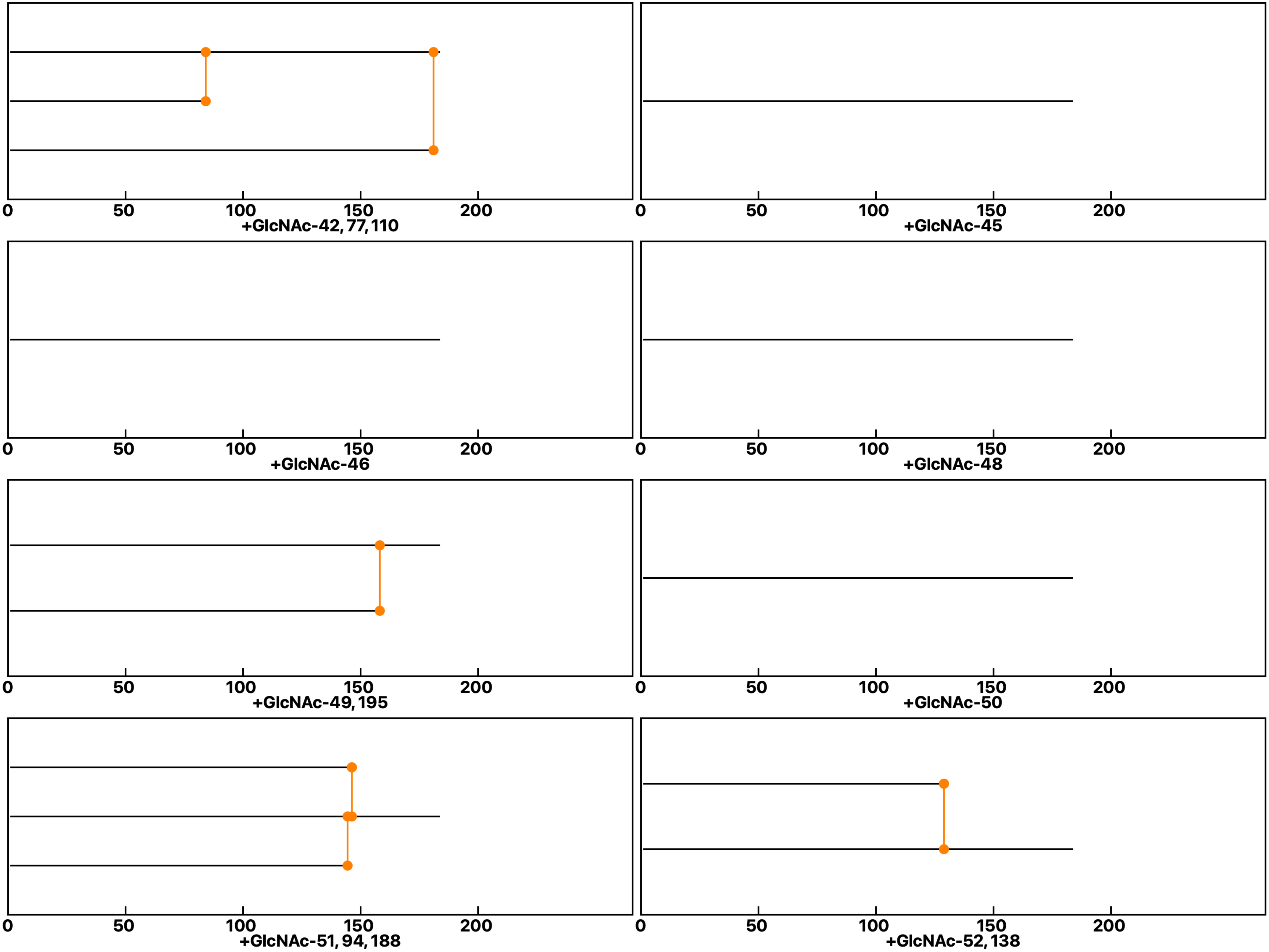

**Analysis: Wild type Gal-3-GlcNAc, Treat.: +GlcNAc, Cell: Myoblasts**

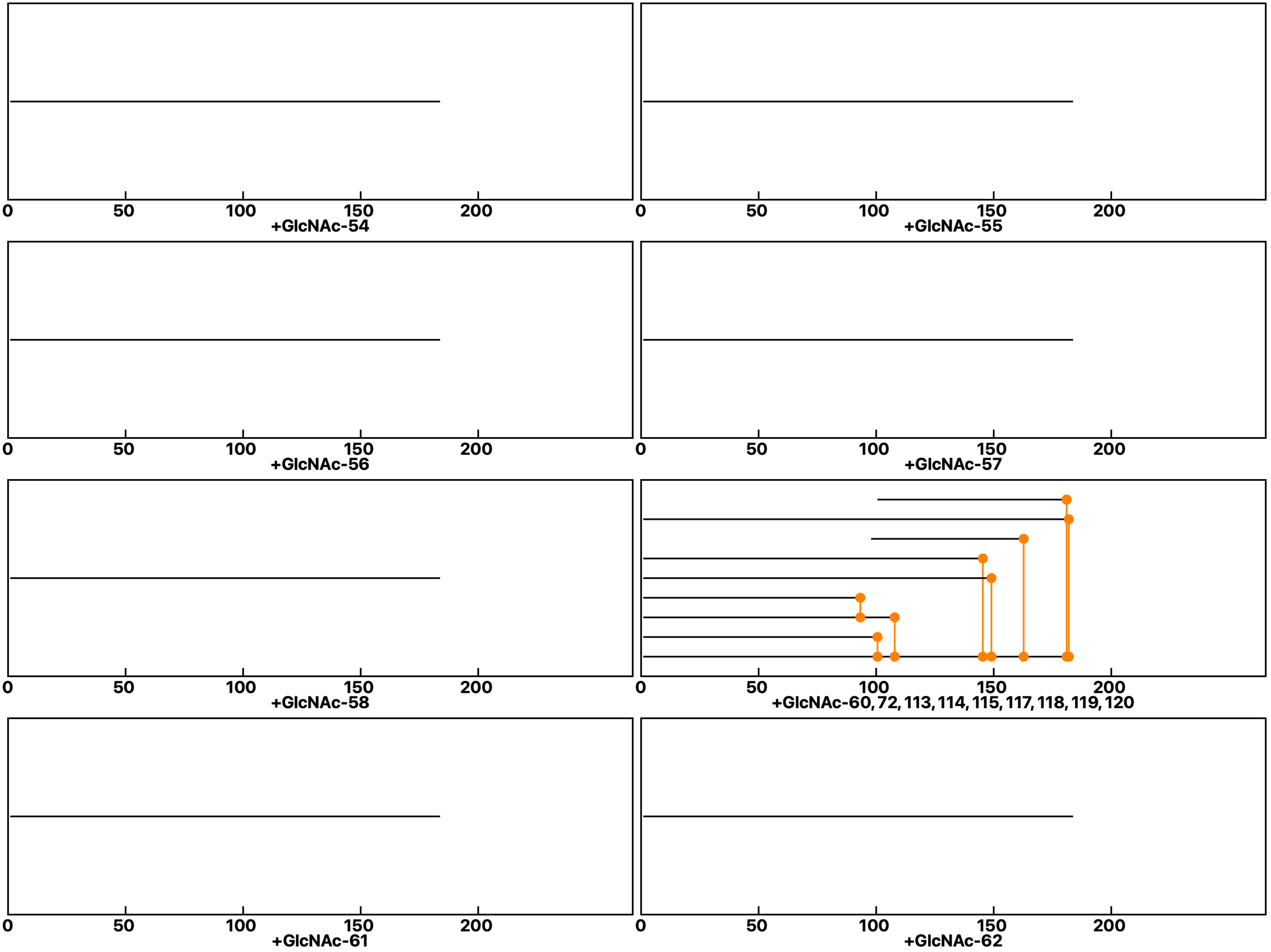

**Analysis: Wild type Gal-3-GlcNAc, Treat.: +GlcNAc, Cell: Myoblasts**

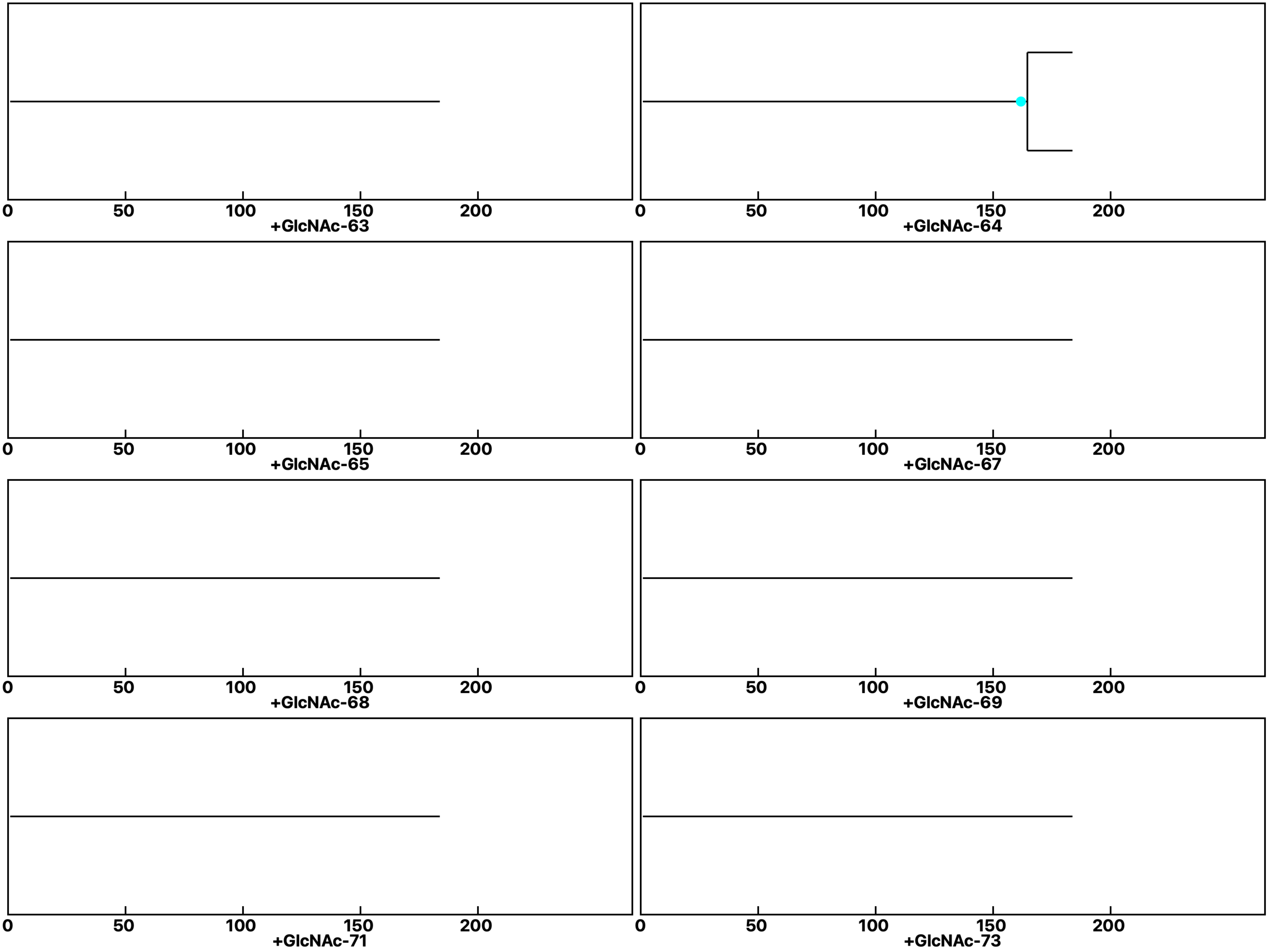

Analysis: Wild type Gal-3-GlcNAc, Treat.: +GlcNAc, Cell: Myoblasts

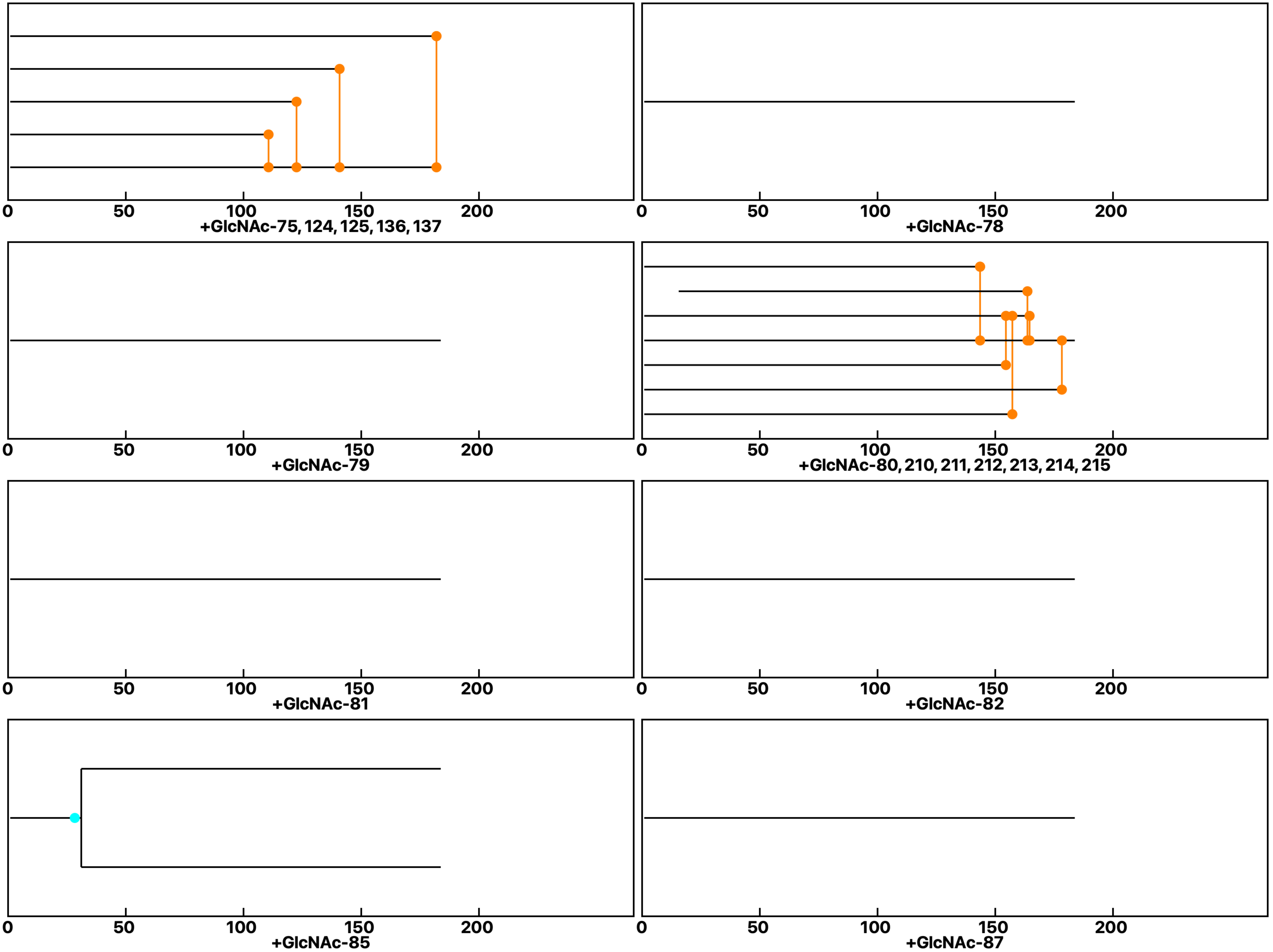

Analysis: Wild type Gal-3-GlcNAc, Treat.: +GlcNAc, Cell: Myoblasts

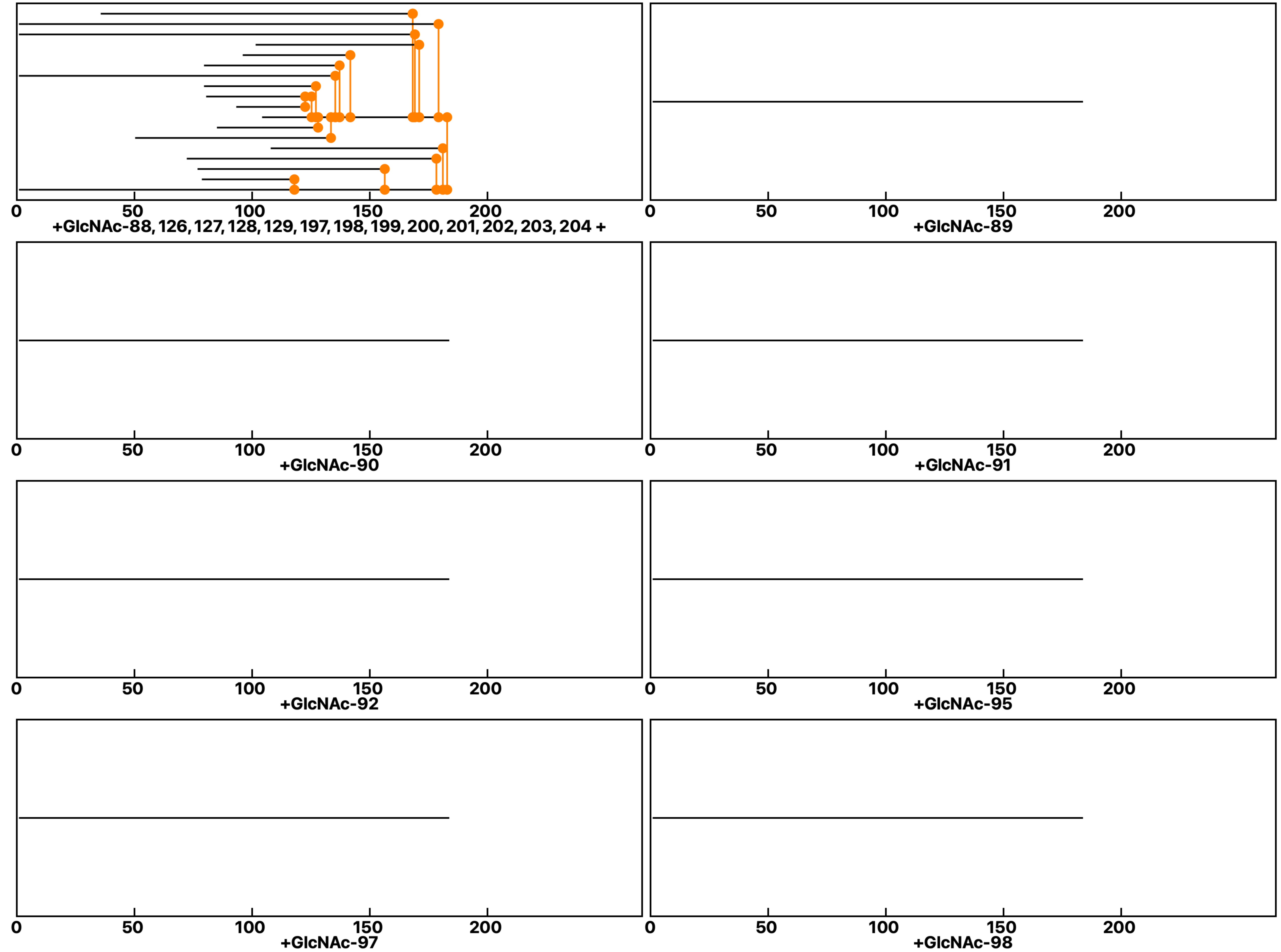

**Analysis: Wild type Gal-3-GlcNAc, Treat.: +GlcNAc, Cell: Myoblasts**

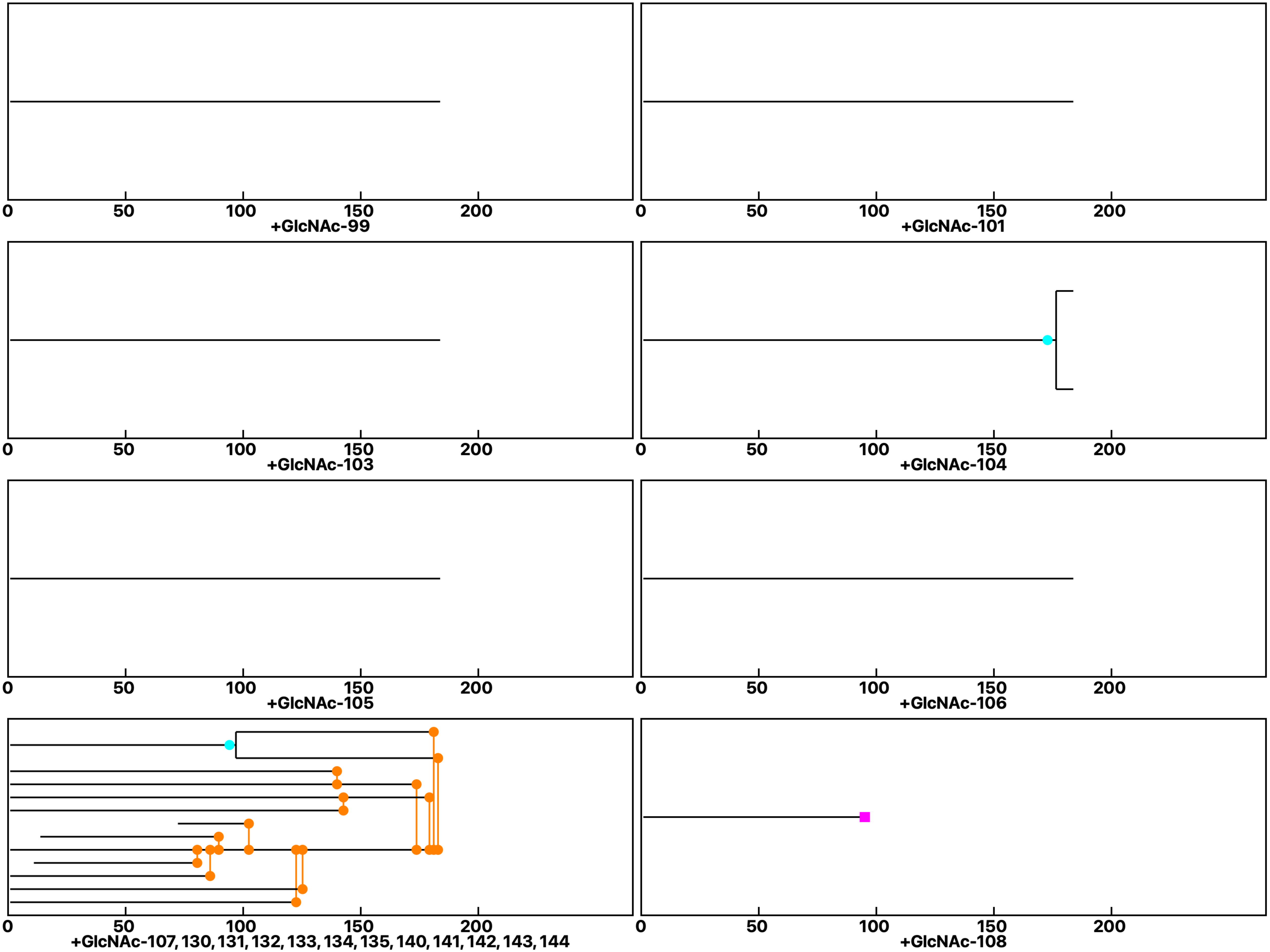

**Analysis: Wild type Gal-3-GlcNAc, Treat.: +GlcNAc, Cell: Myoblasts**

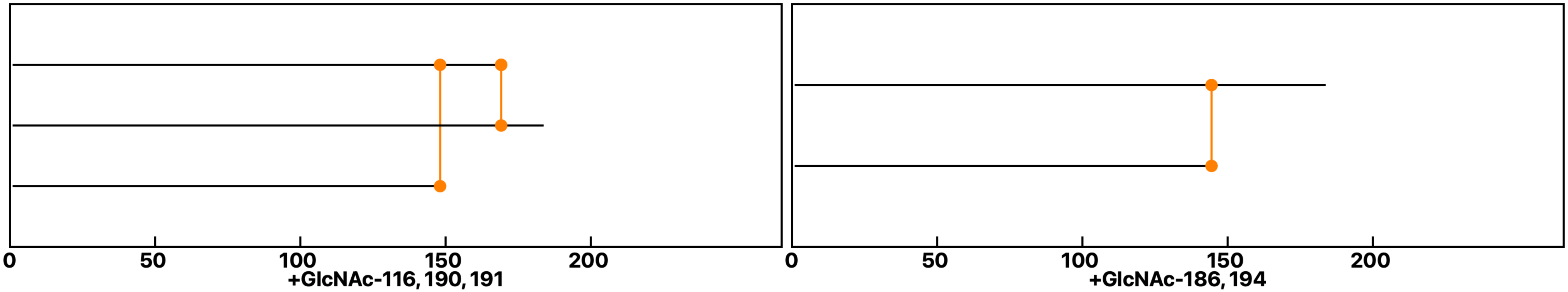
