## Supplemental data 8 for "N-Acetylglucosamine Facilitates Coordinated Myoblast Flow, Forming the Foundation for Efficient Myogenesis"

### Analysis: Wild type Gal-3-GlcNAc, Treat.: +Gal-3+GlcNAc, Cell: Myoblasts

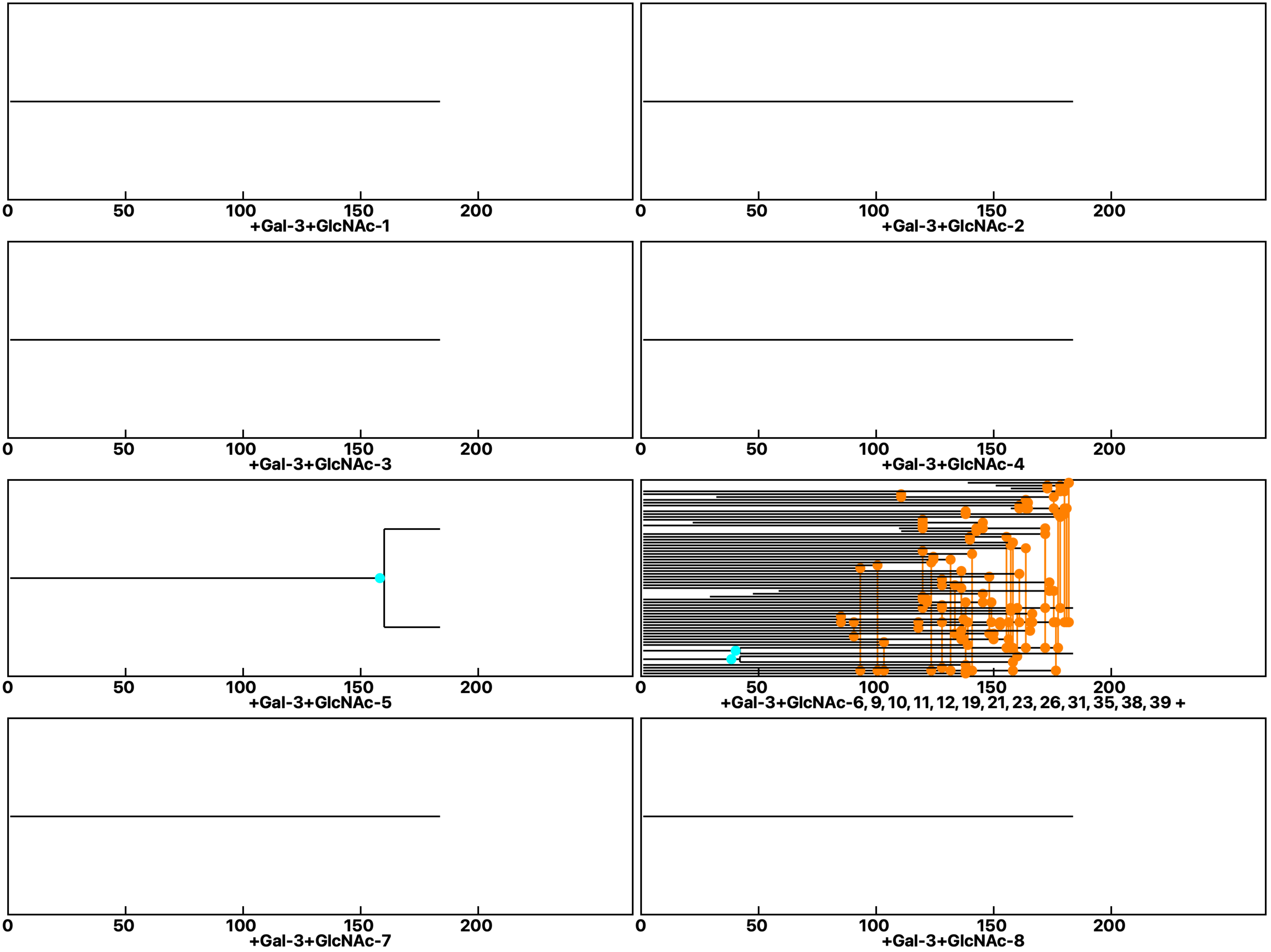

Analysis: Wild type Gal-3-GlcNAc, Treat.: +Gal-3+GlcNAc, Cell: Myoblasts

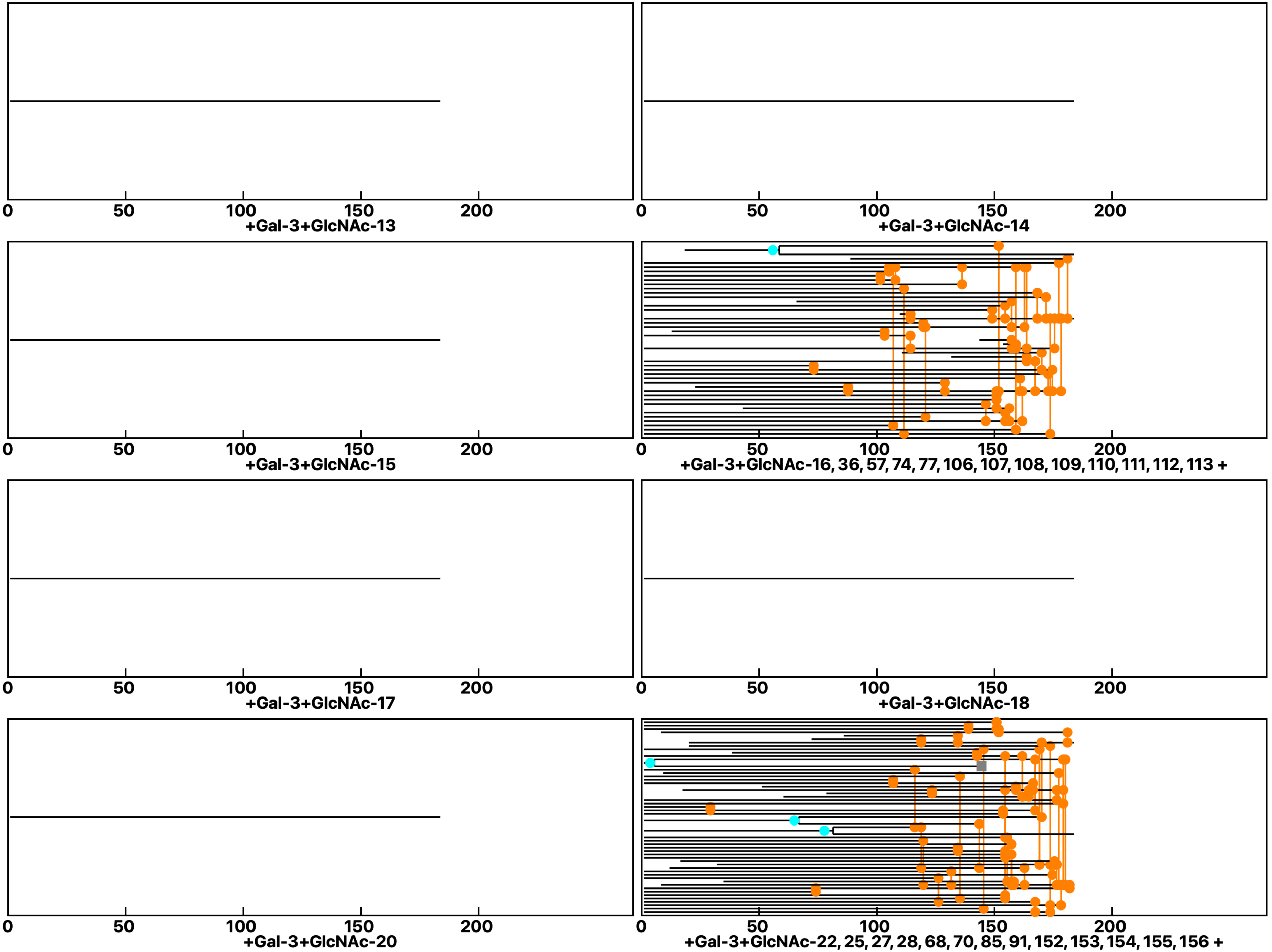

**Analysis: Wild type Gal-3-GlcNAc, Treat.: +Gal-3+GlcNAc, Cell: Myoblasts**

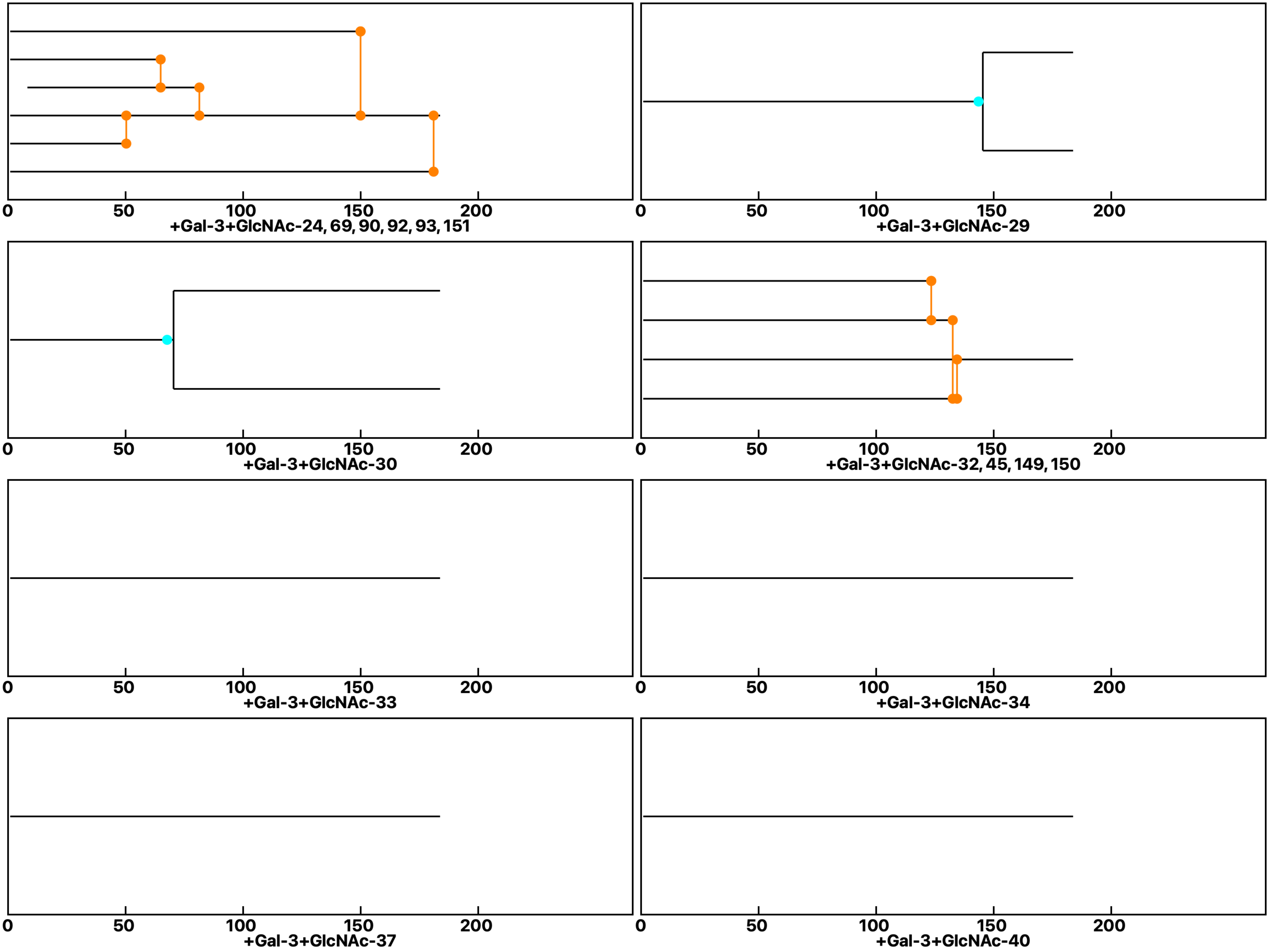

**Analysis: Wild type Gal-3-GlcNAc, Treat.: +Gal-3+GlcNAc, Cell: Myoblasts**

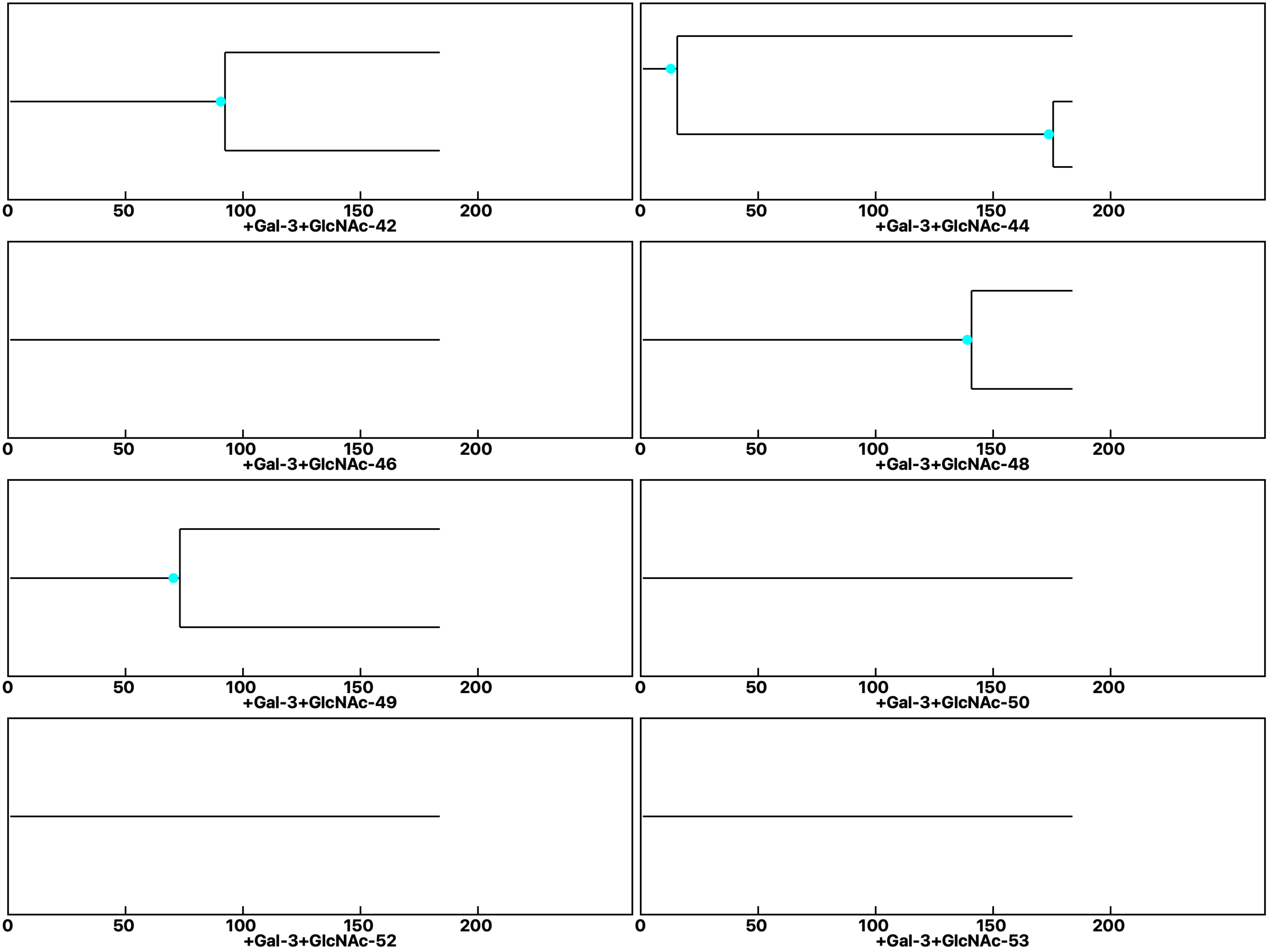

Analysis: Wild type Gal-3-GlcNAc, Treat.: +Gal-3+GlcNAc, Cell: Myoblasts

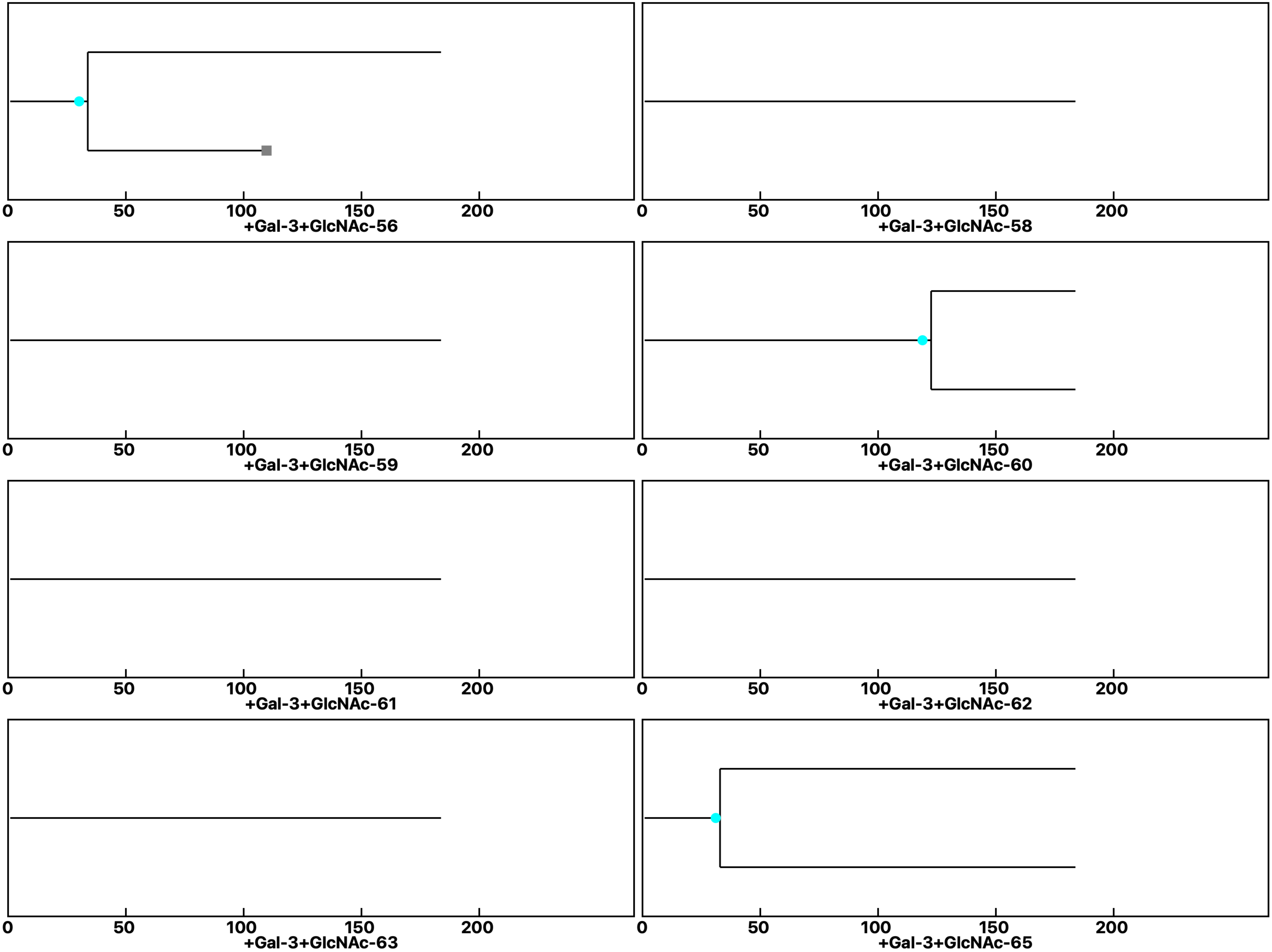

Analysis: Wild type Gal-3-GlcNAc, Treat.: +Gal-3+GlcNAc, Cell: Myoblasts

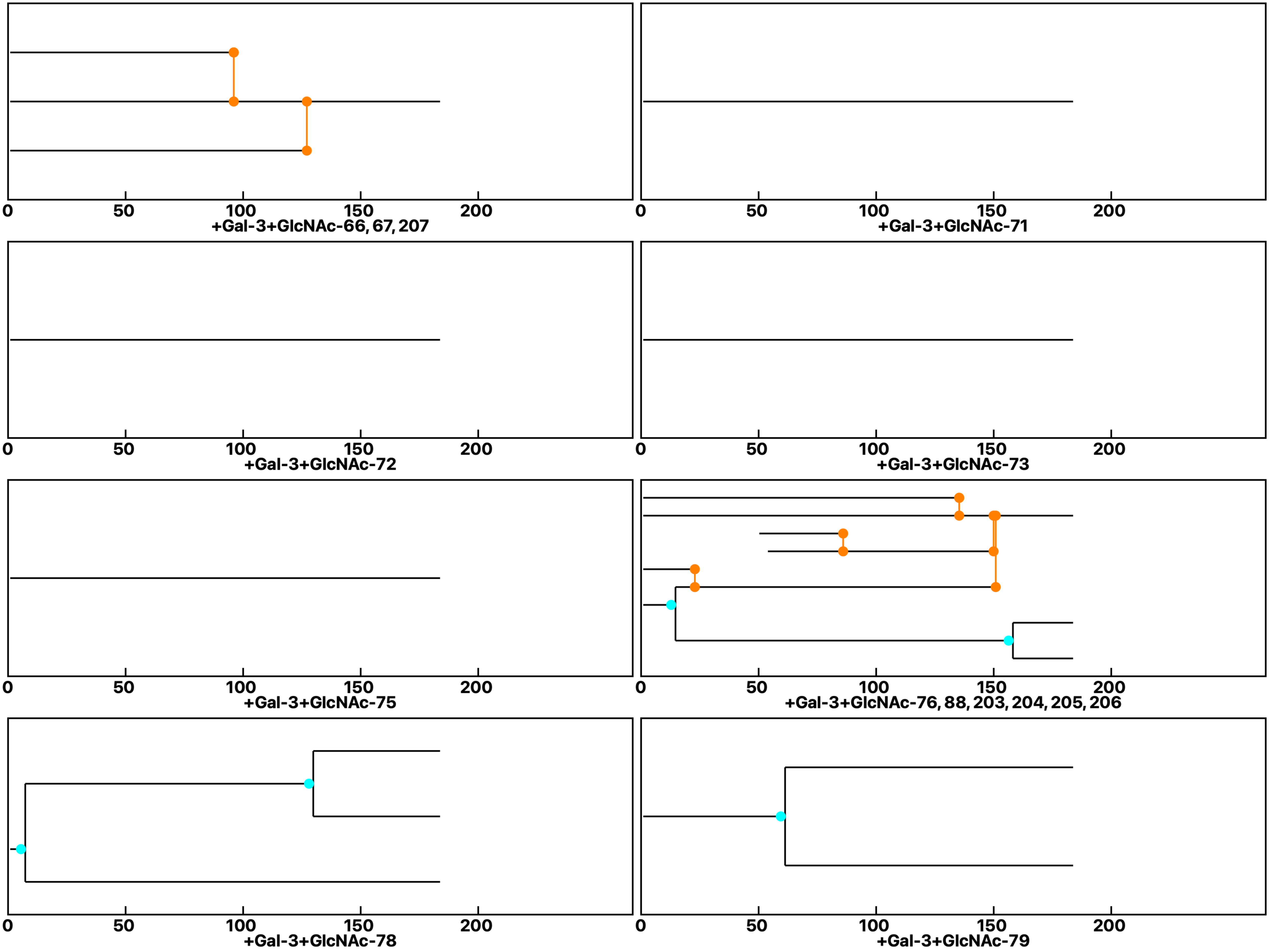

Analysis: Wild type Gal-3-GlcNAc, Treat.: +Gal-3+GlcNAc, Cell: Myoblasts

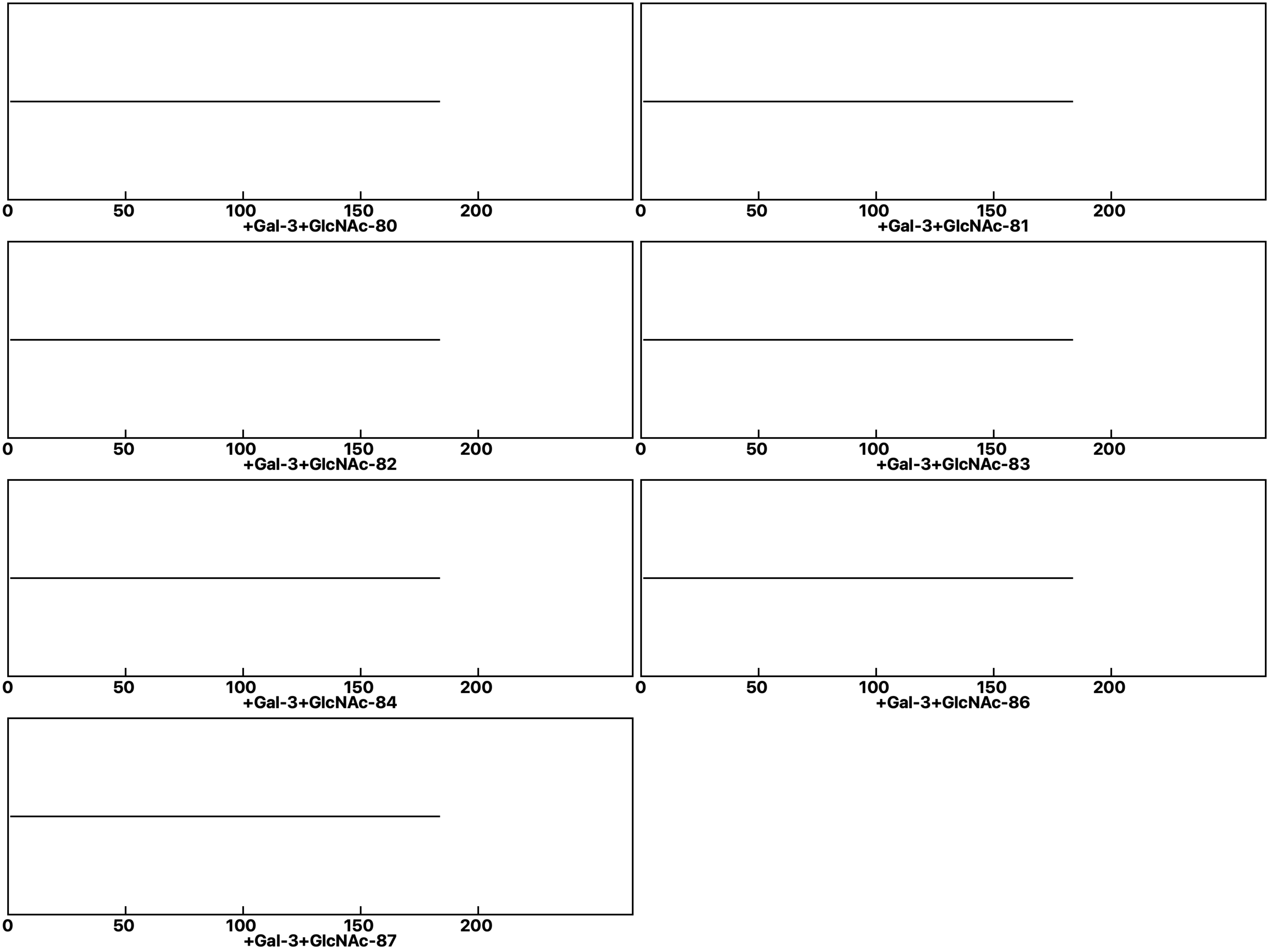
