## Supplemental data 9 for "N-Acetylglucosamine Facilitates Coordinated Myoblast Flow, Forming the Foundation for Efficient Myogenesis"

Analysis: Gal-3KO Gal-3-GlcNAc, Treat.: Control, Cell: Myoblasts

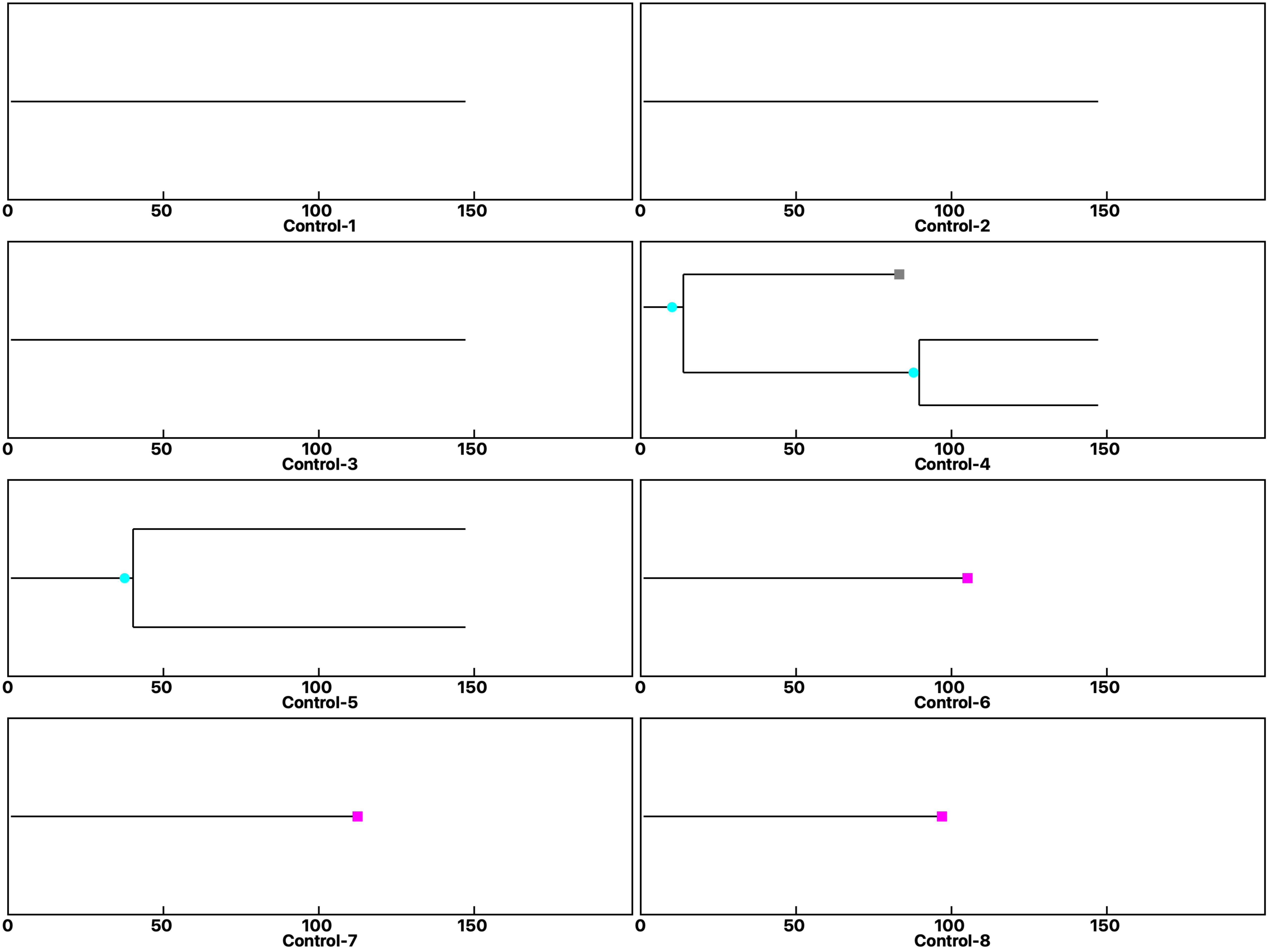

Analysis: Gal-3KO Gal-3-GlcNAc, Treat.: Control, Cell: Myoblasts

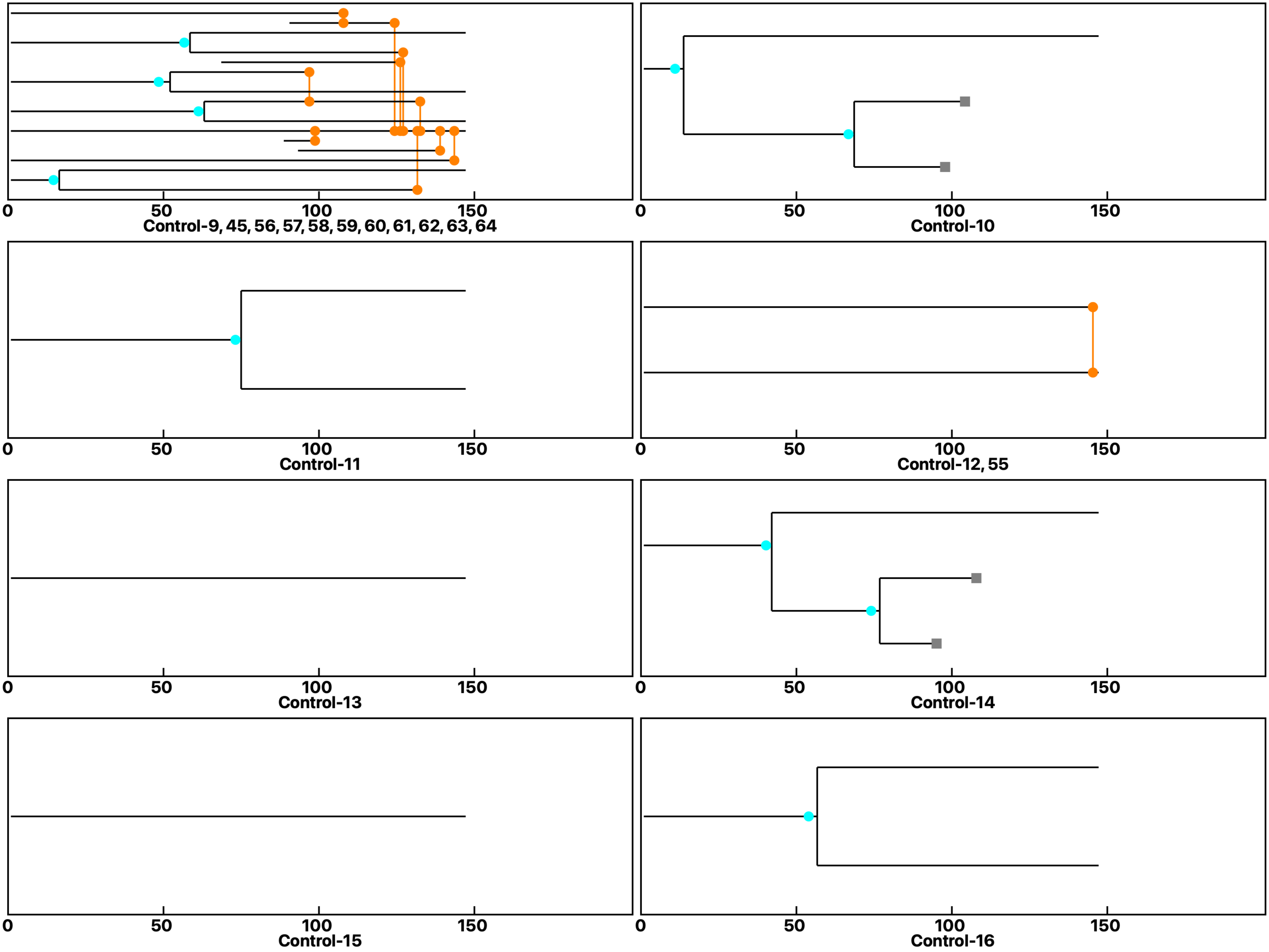

Analysis: Gal-3KO Gal-3-GlcNAc, Treat.: Control, Cell: Myoblasts

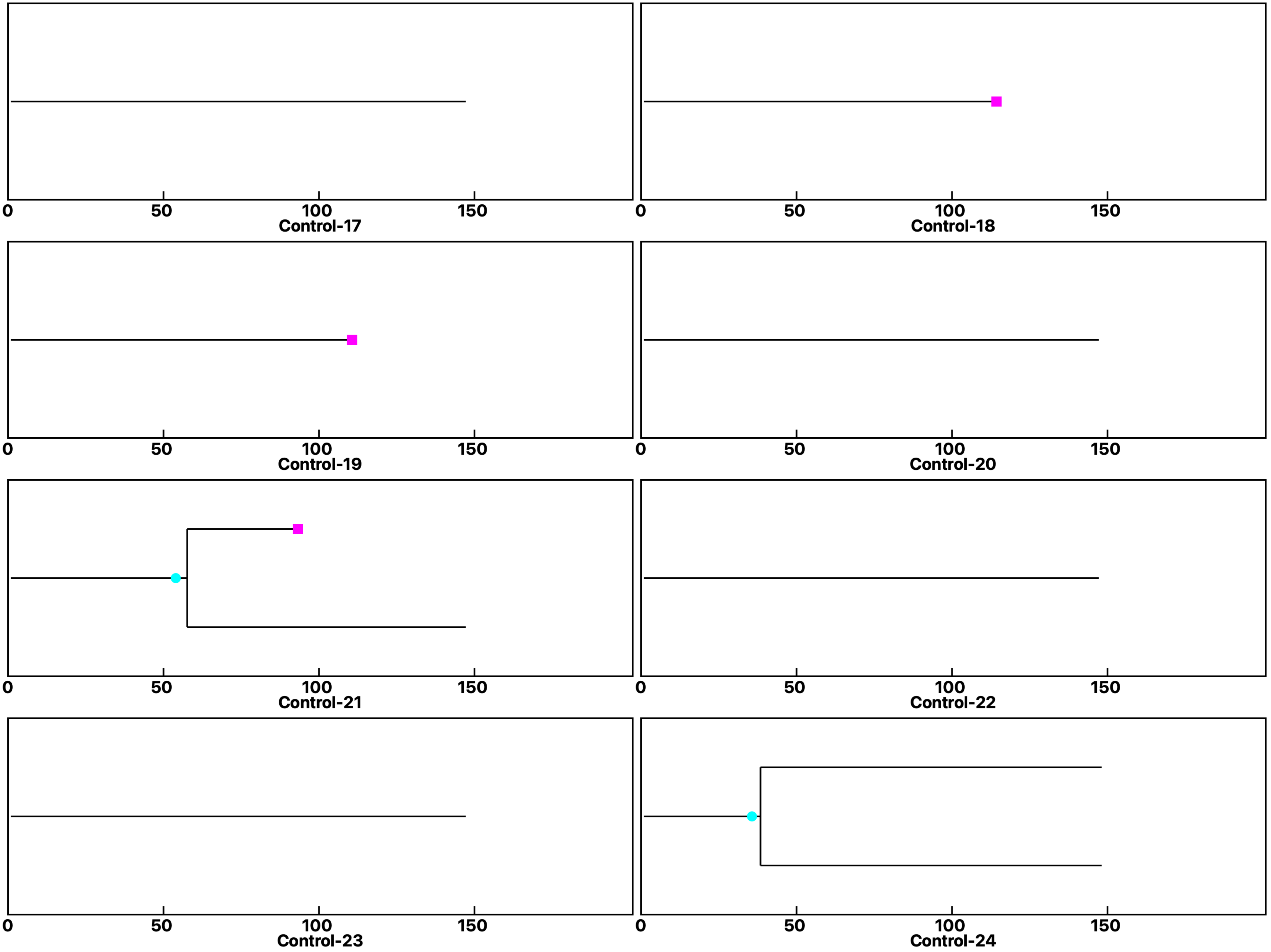

Analysis: Gal-3KO Gal-3-GlcNAc, Treat.: Control, Cell: Myoblasts

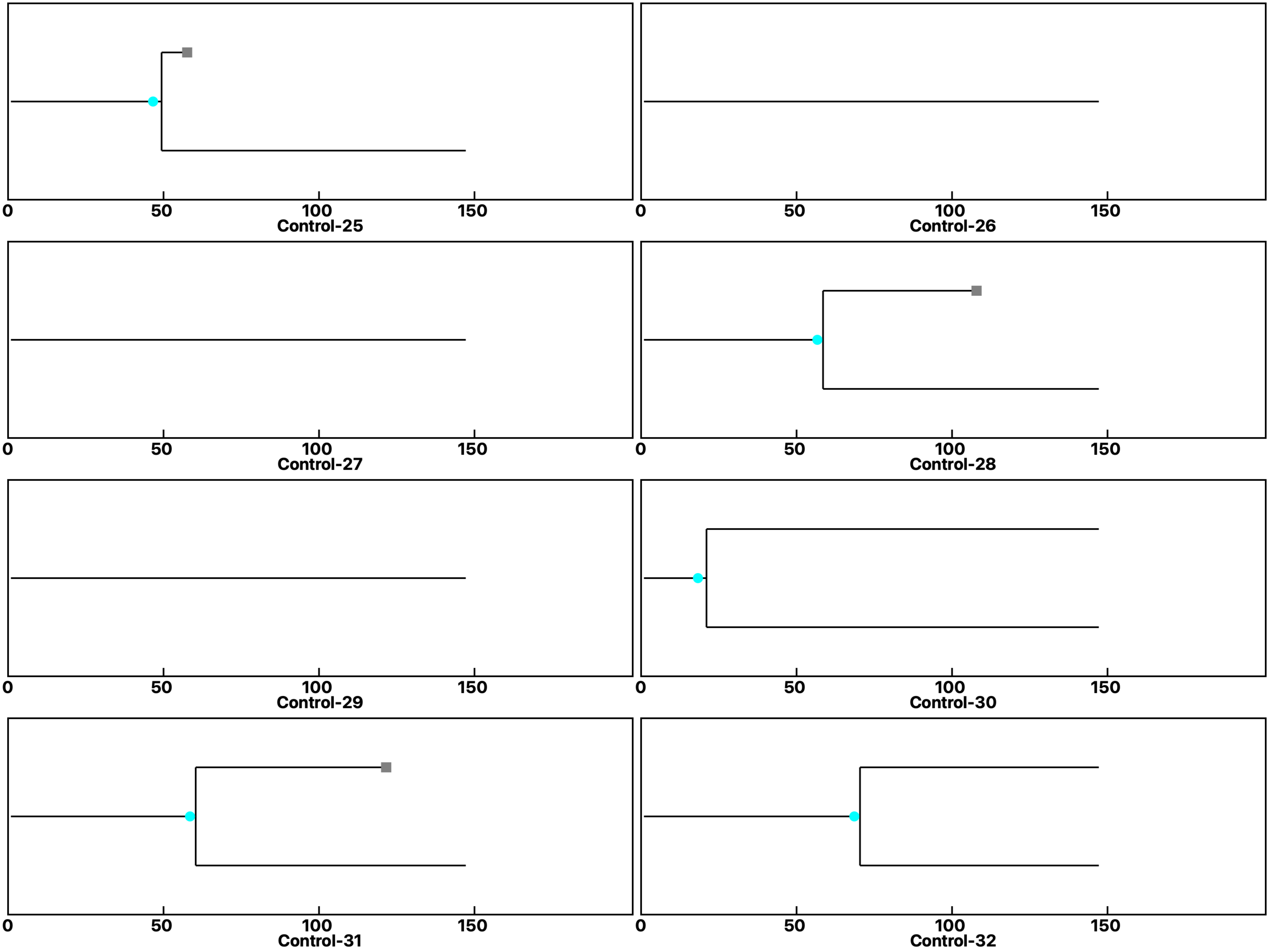

**Analysis: Gal-3KO Gal-3-GlcNAc, Treat.: Control, Cell: Myoblasts**

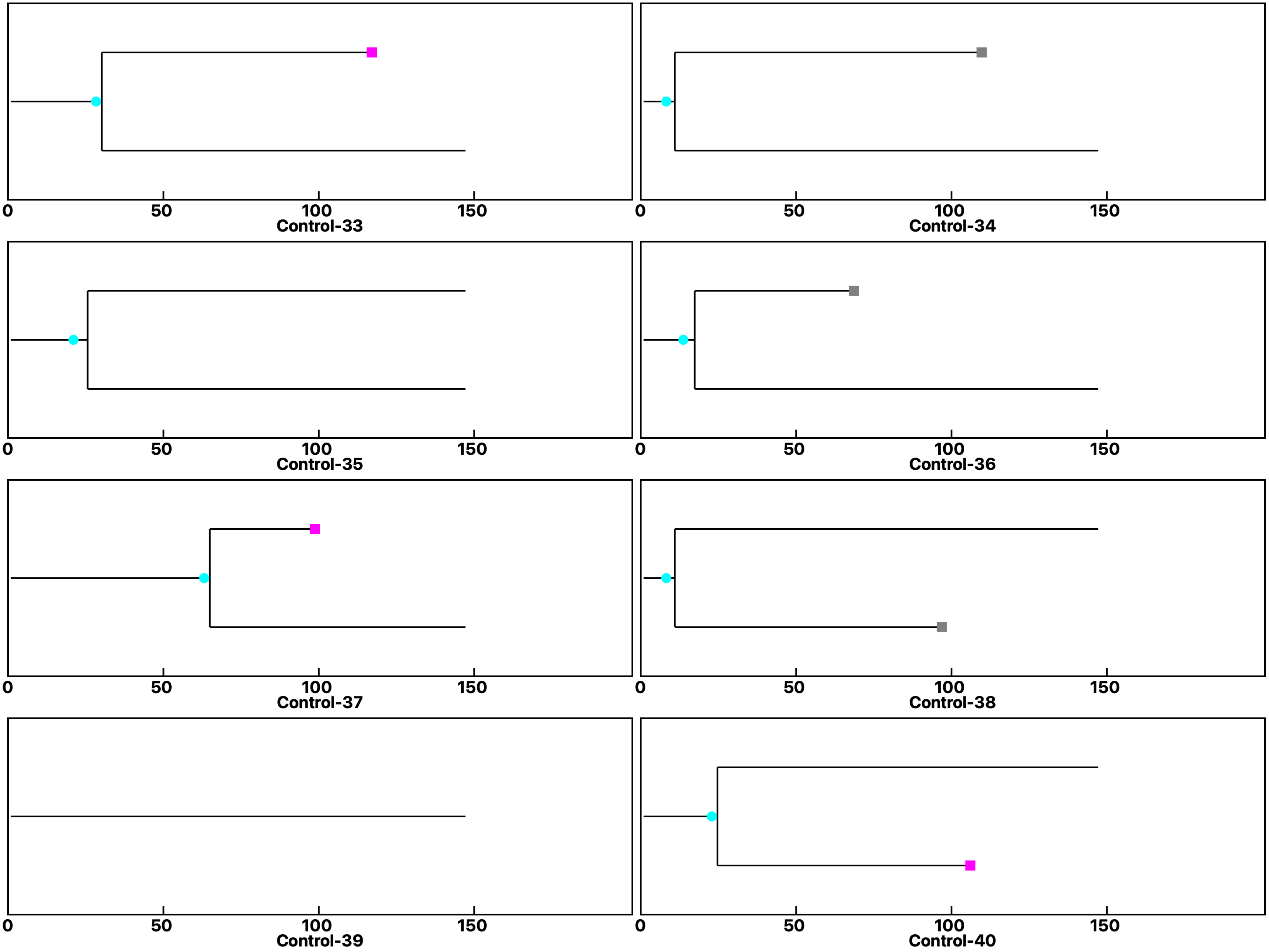

**Analysis: Gal-3KO Gal-3-GlcNAc, Treat.: Control, Cell: Myoblasts**

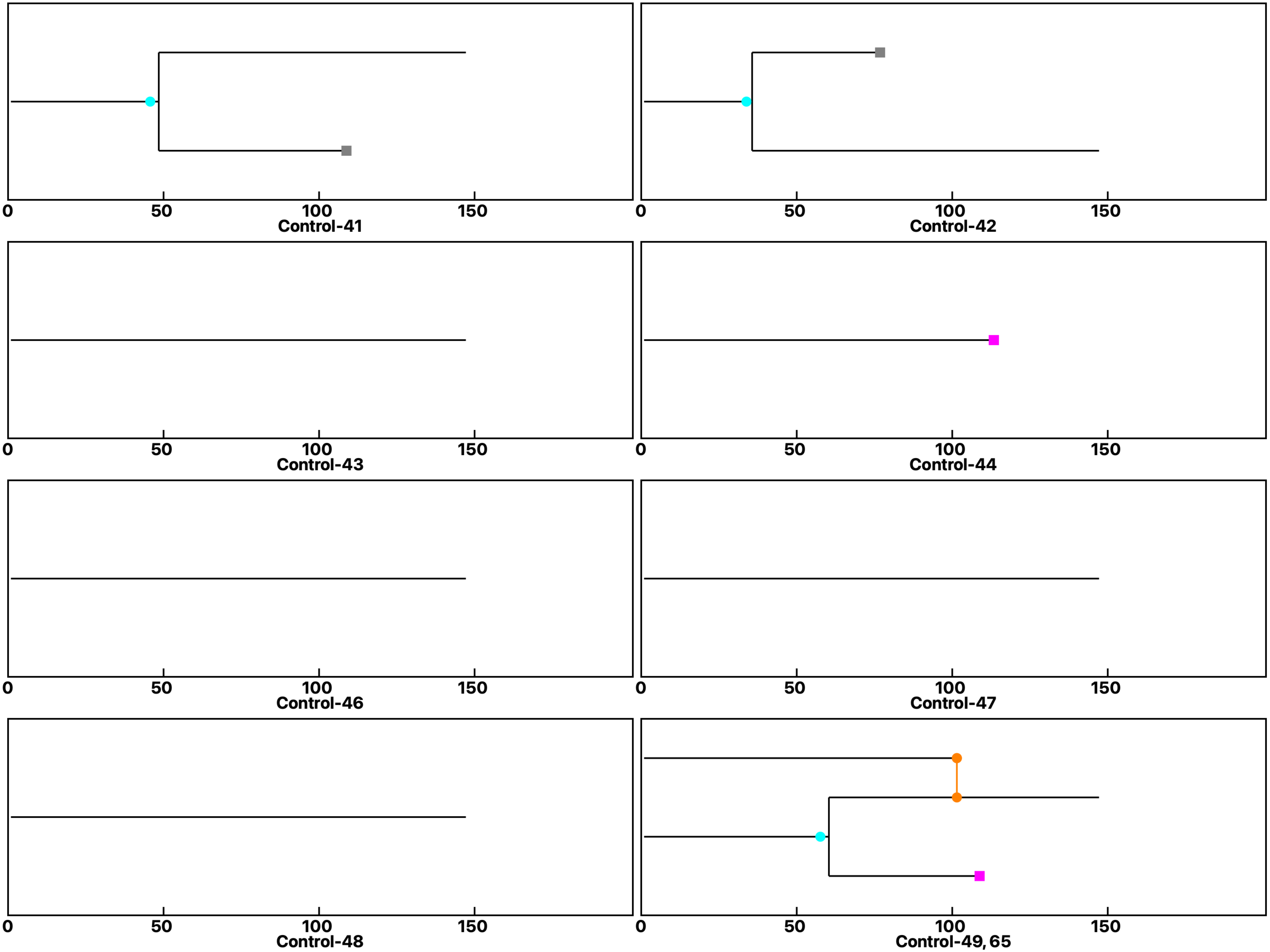

Analysis: Gal-3KO Gal-3-GlcNAc, Treat.: Control, Cell: Myoblasts

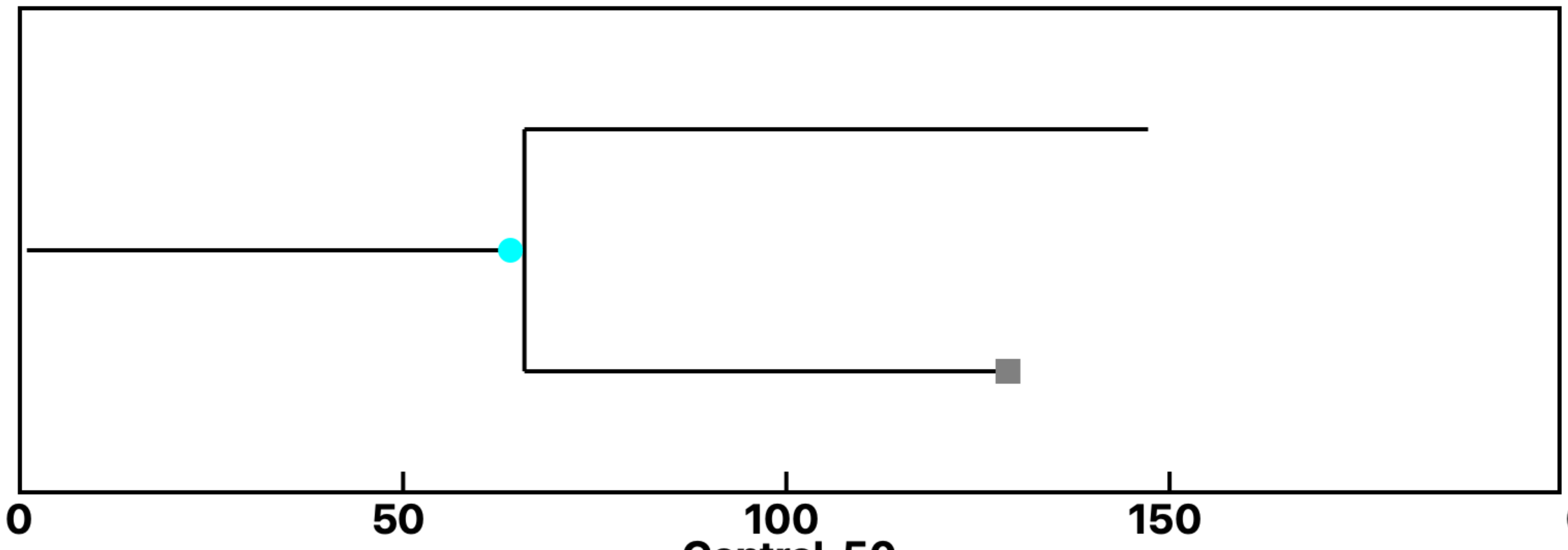

Control-50

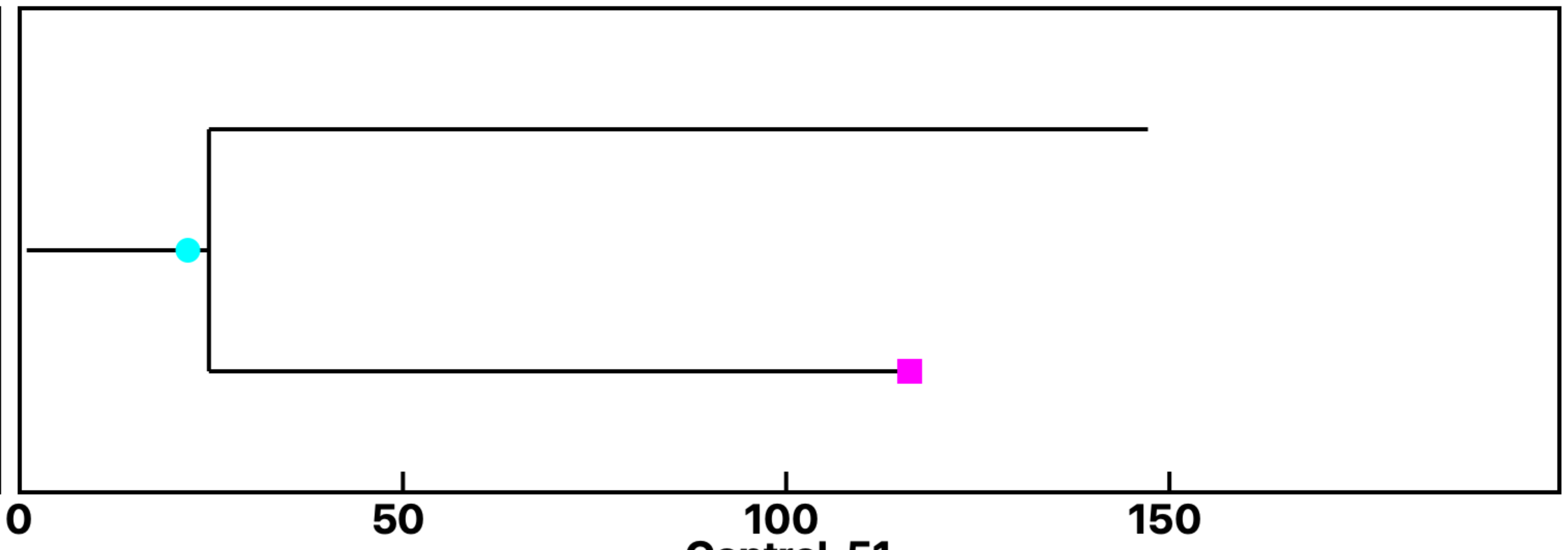

Control-51

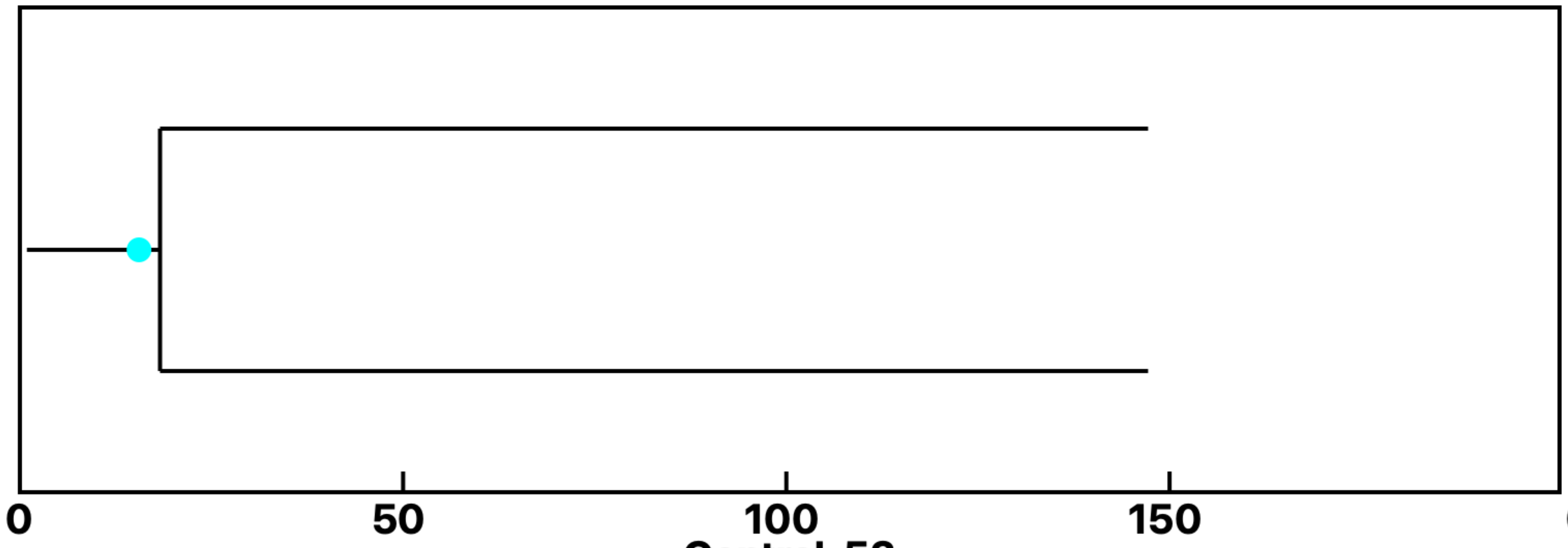

Control-52

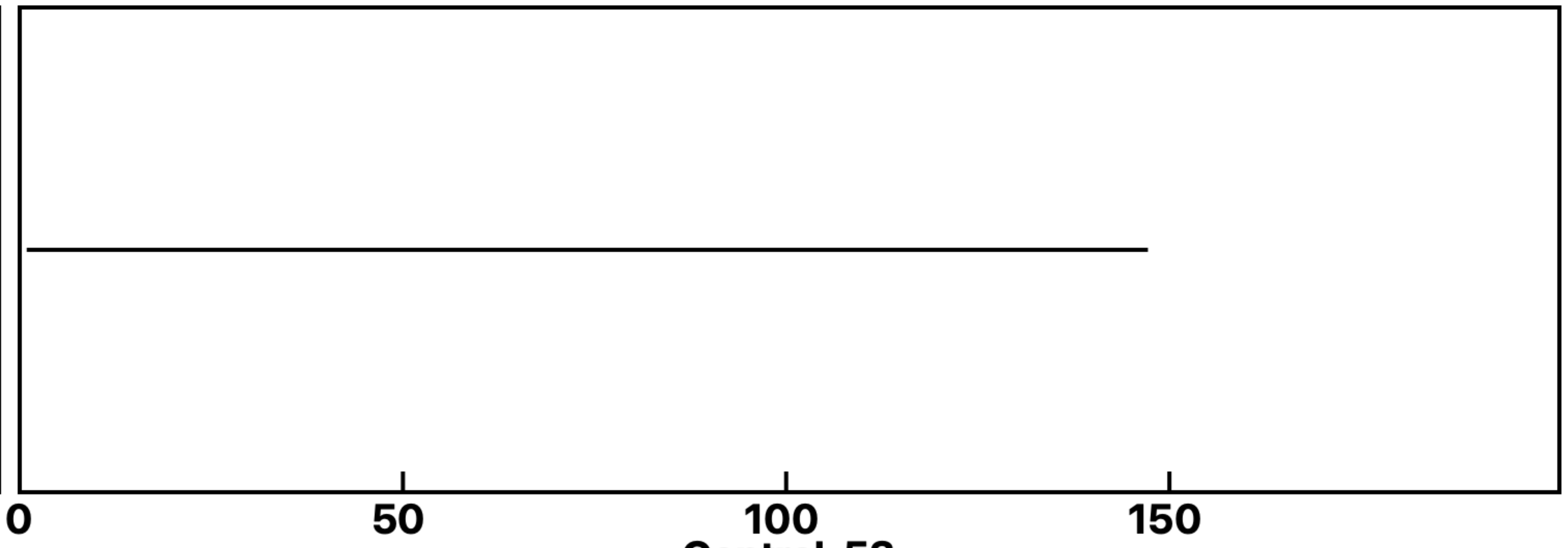

Control-53

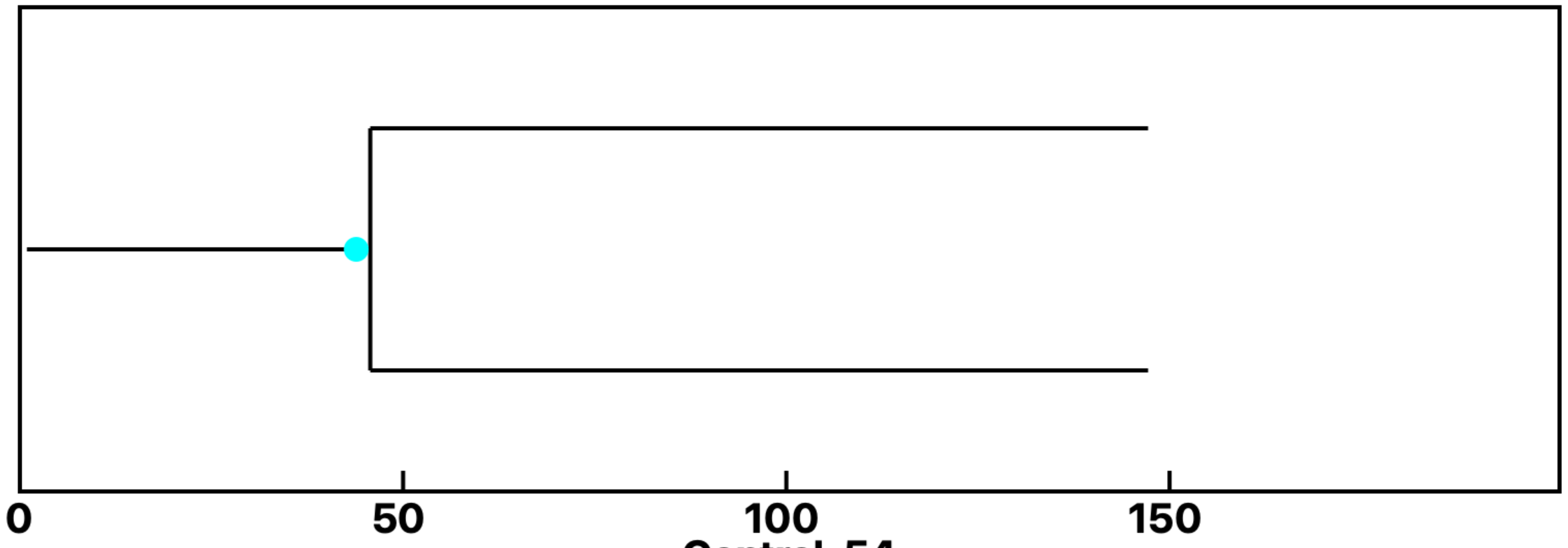

Control-54
