## Supplemental data 10 for "N-Acetylglucosamine Facilitates Coordinated Myoblast Flow, Forming the Foundation for Efficient Myogenesis"

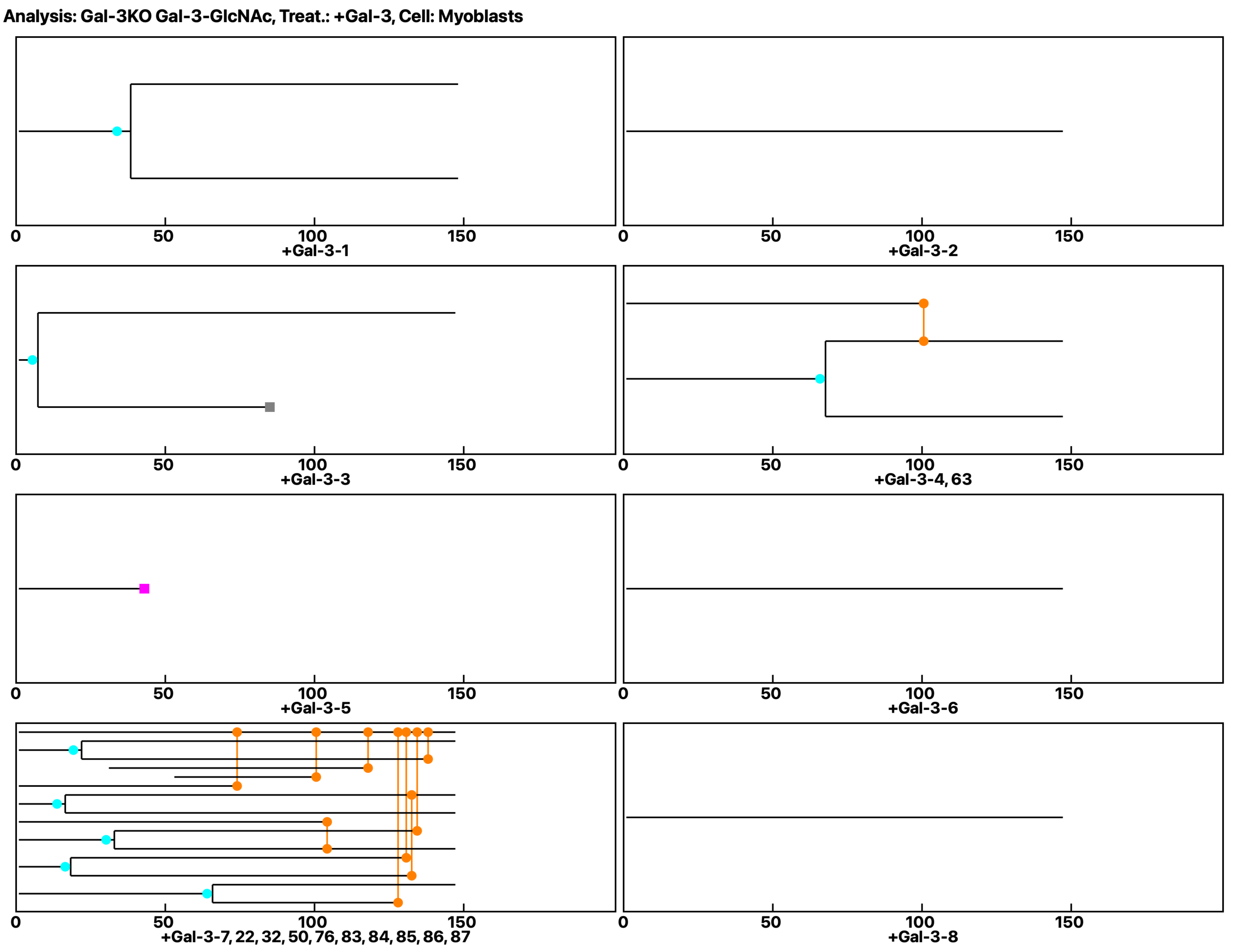
Analysis: Gal-3KO Gal-3-GlcNAc, Treat.: +Gal-3, Cell: Myoblasts
0
50
100
150
+Gal-3-1
+Gal-3-2
+Gal-3-3
+Gal-3-4, 63
+Gal-3-5
+Gal-3-6
+Gal-3-7, 22, 32, 50, 76, 83, 84, 85, 86, 87
+Gal-3-8

Analysis: Gal-3KO Gal-3-GlcNAc, Treat.: +Gal-3, Cell: Myoblasts

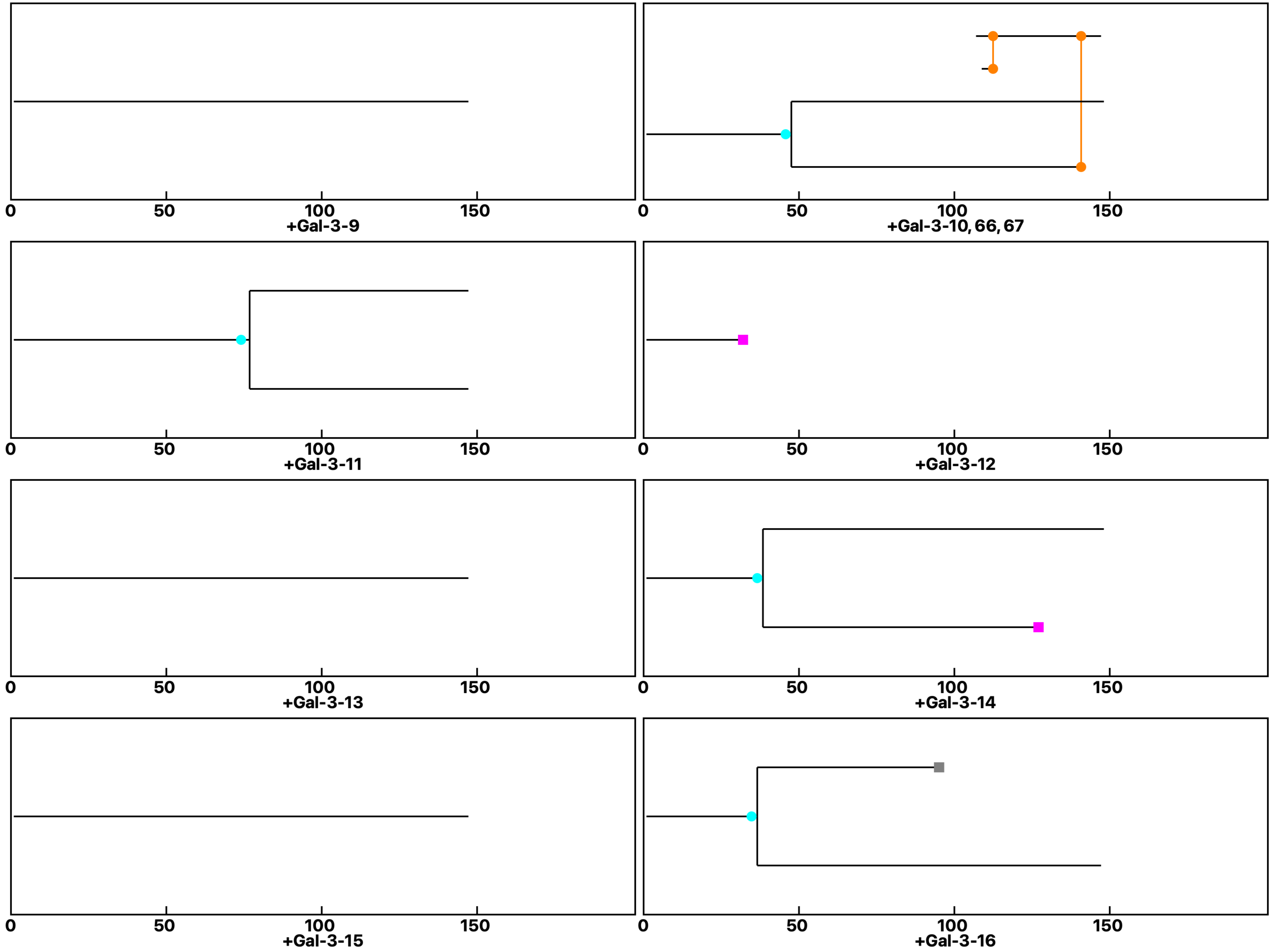

**Analysis: Gal-3KO Gal-3-GlcNAc, Treat.: +Gal-3, Cell: Myoblasts**

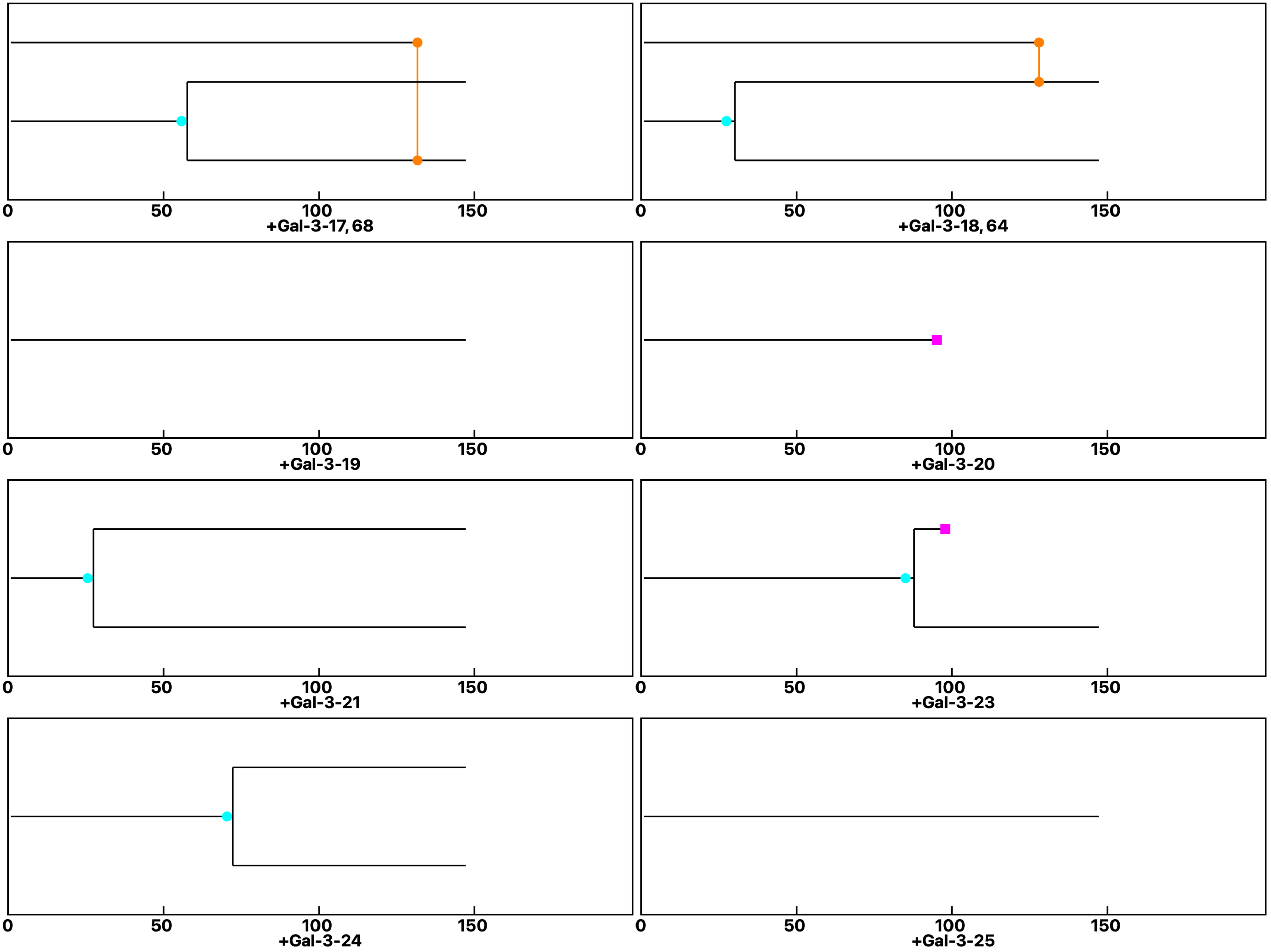

**Analysis: Gal-3KO Gal-3-GlcNAc, Treat.: +Gal-3, Cell: Myoblasts**

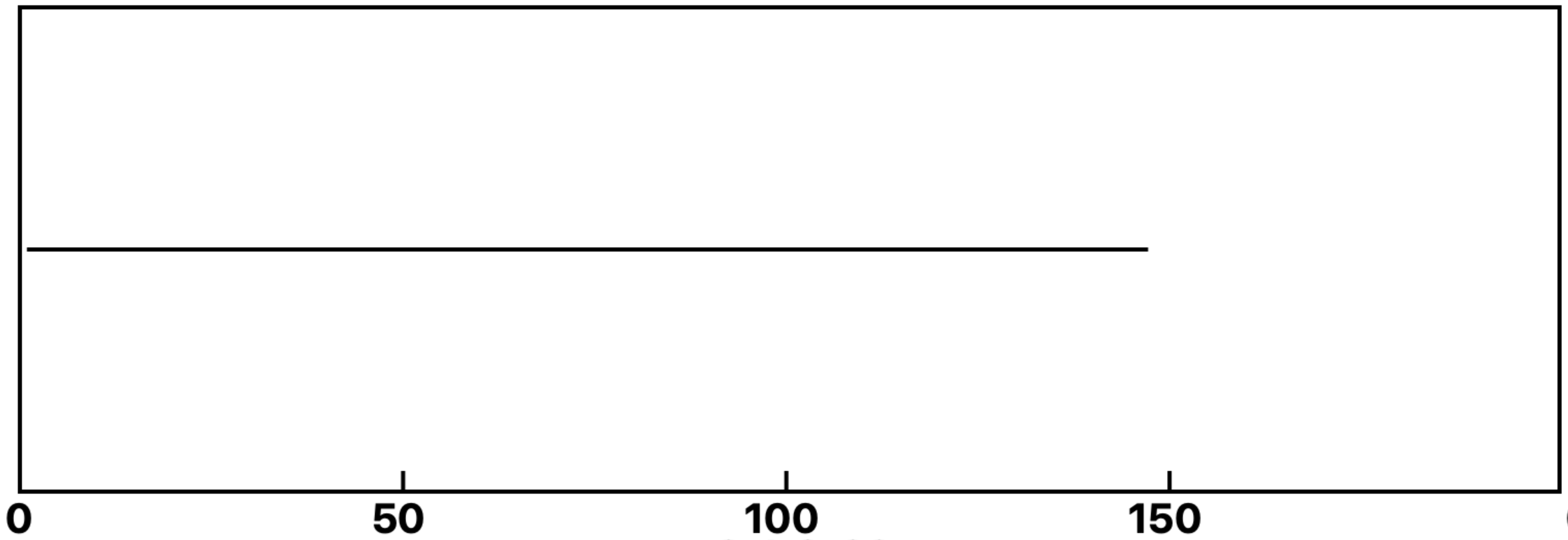

+Gal-3-26

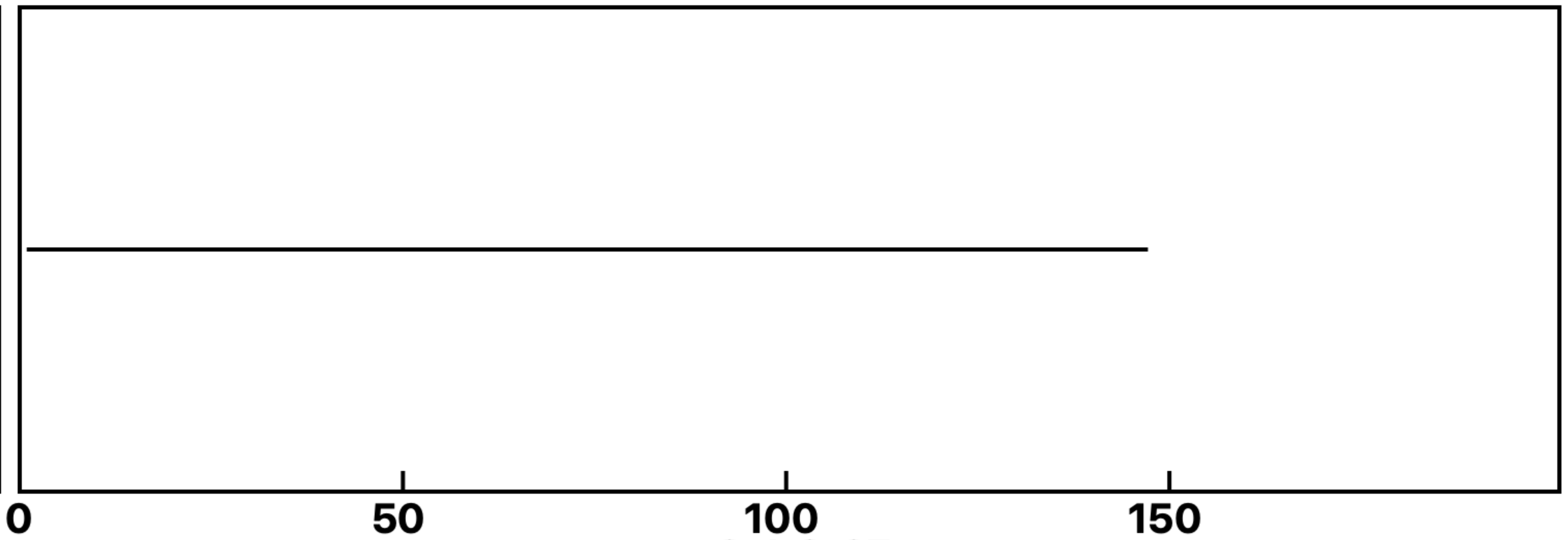

+Gal-3-27

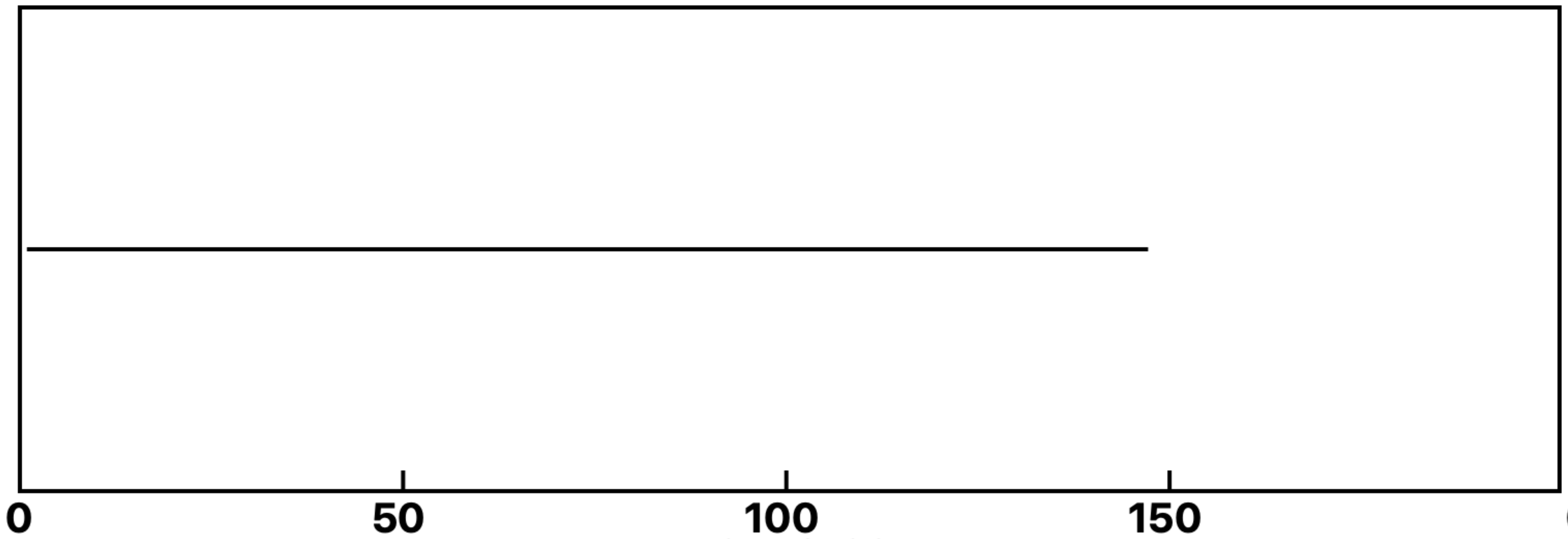

+Gal-3-28

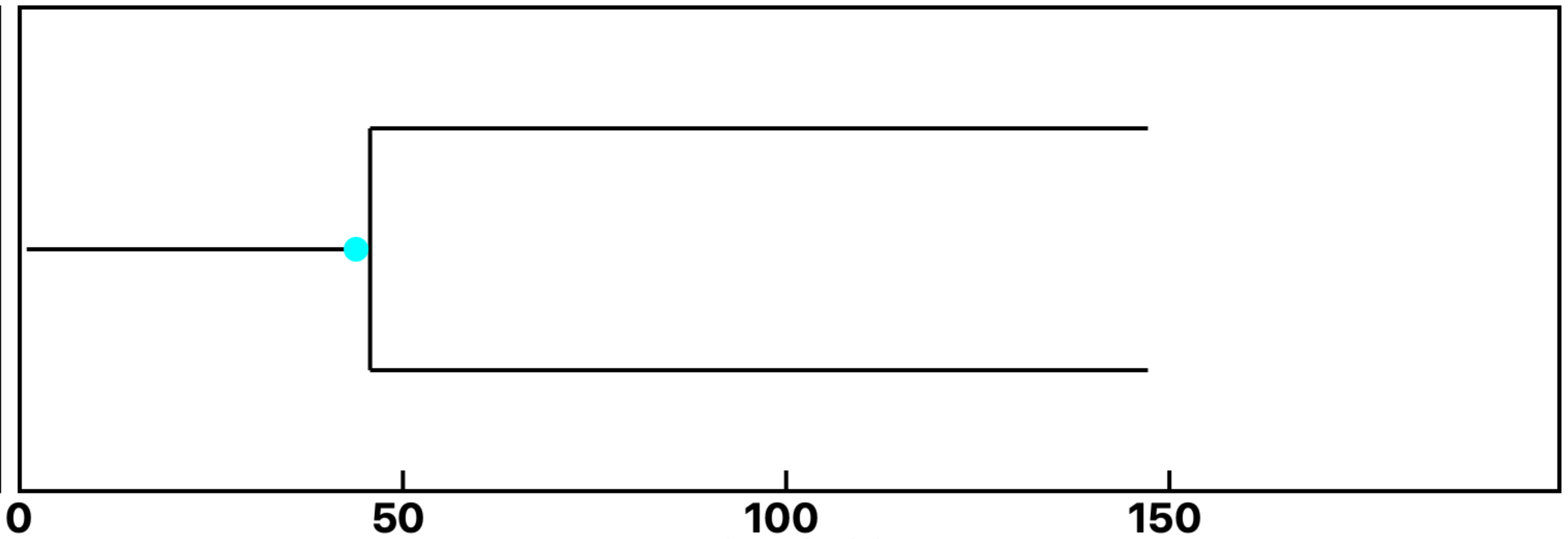

+Gal-3-29

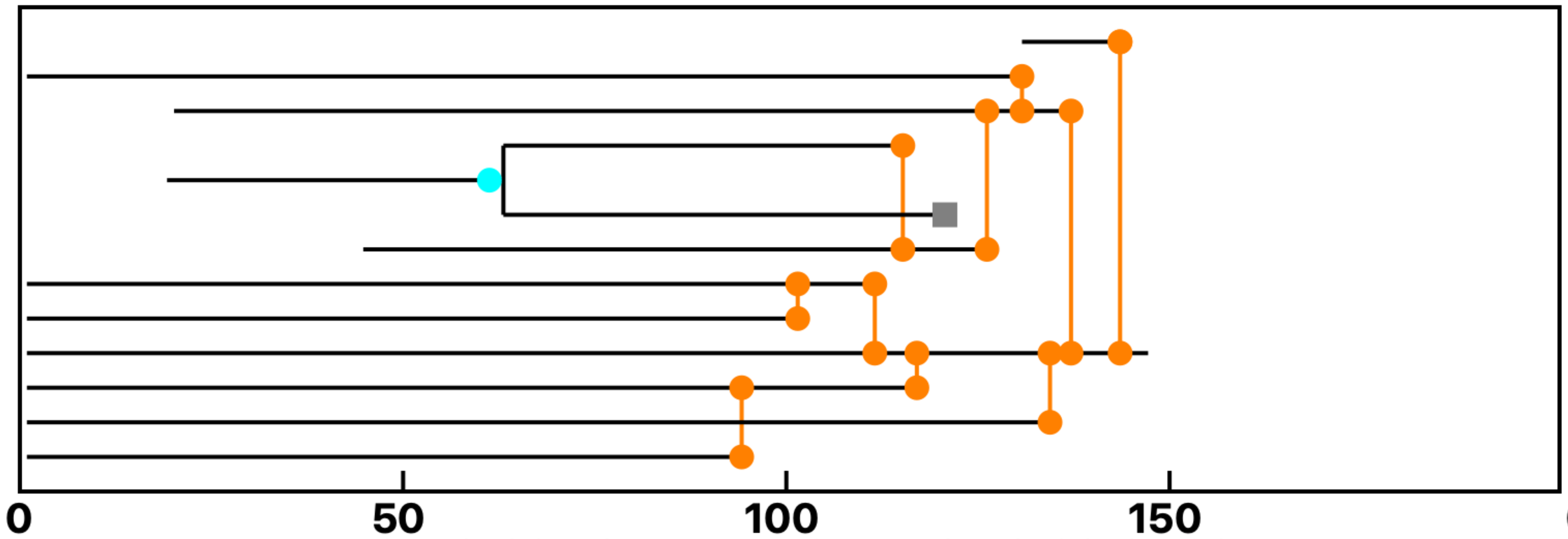

+Gal-3-30, 53, 54, 55, 56, 57, 58, 59, 60, 61, 62

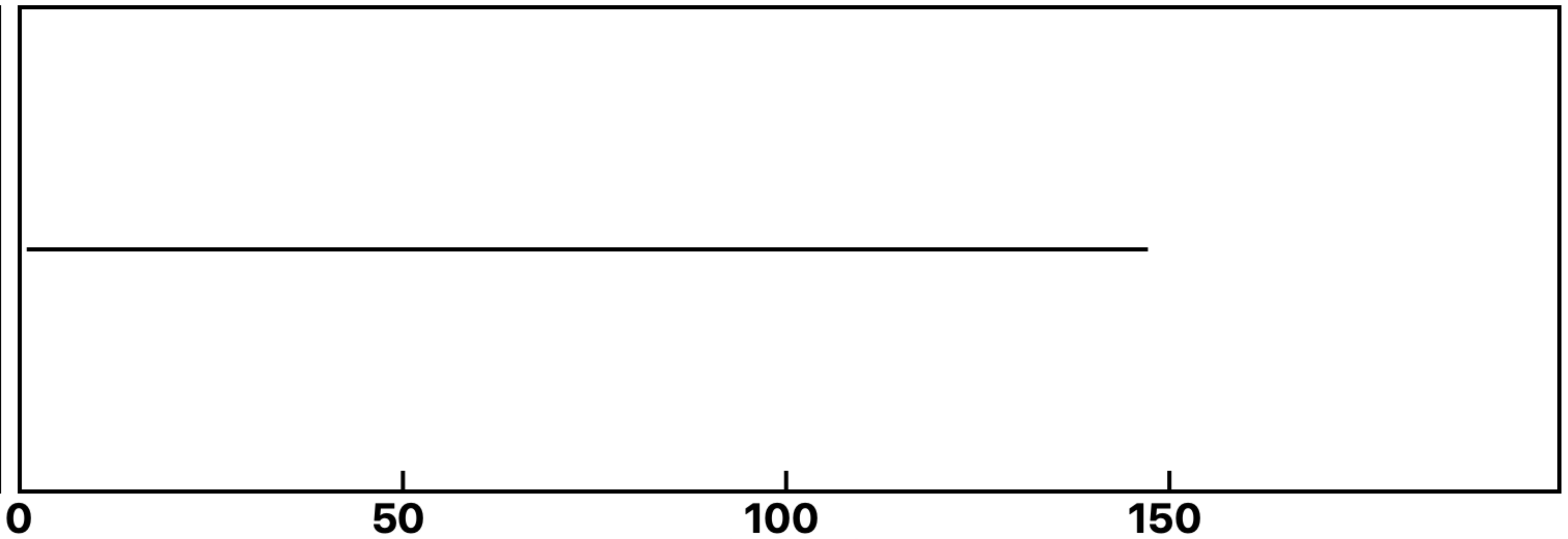

+Gal-3-31

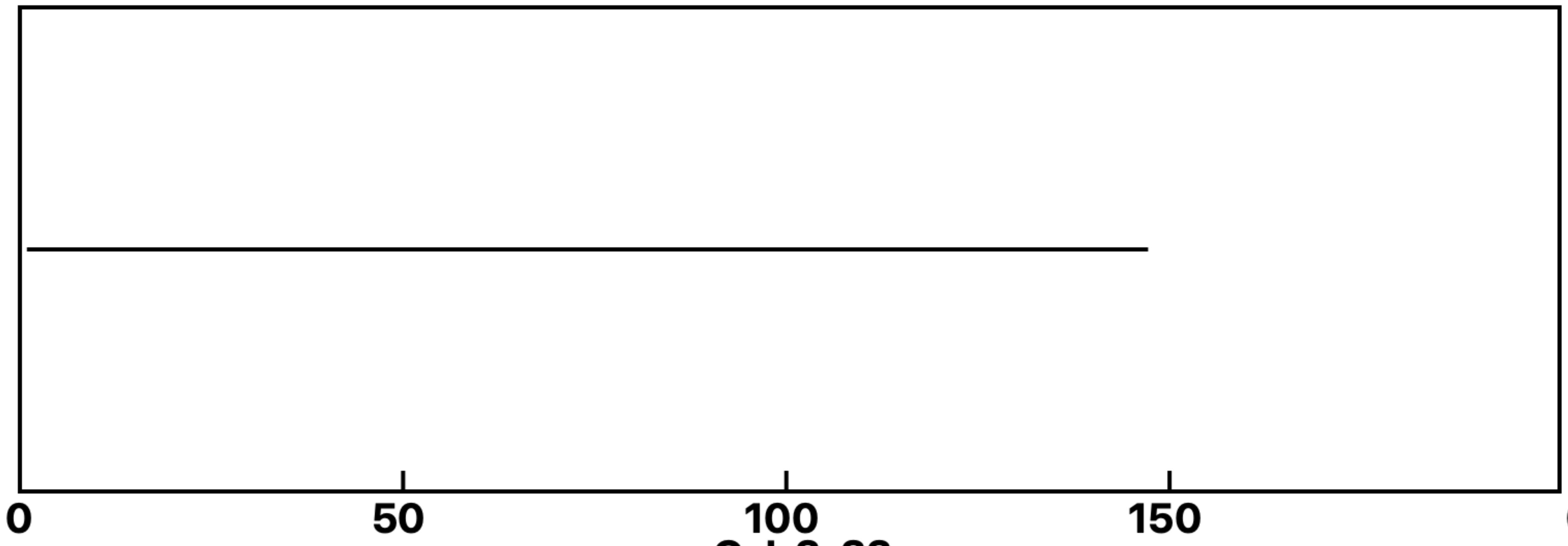

+Gal-3-33

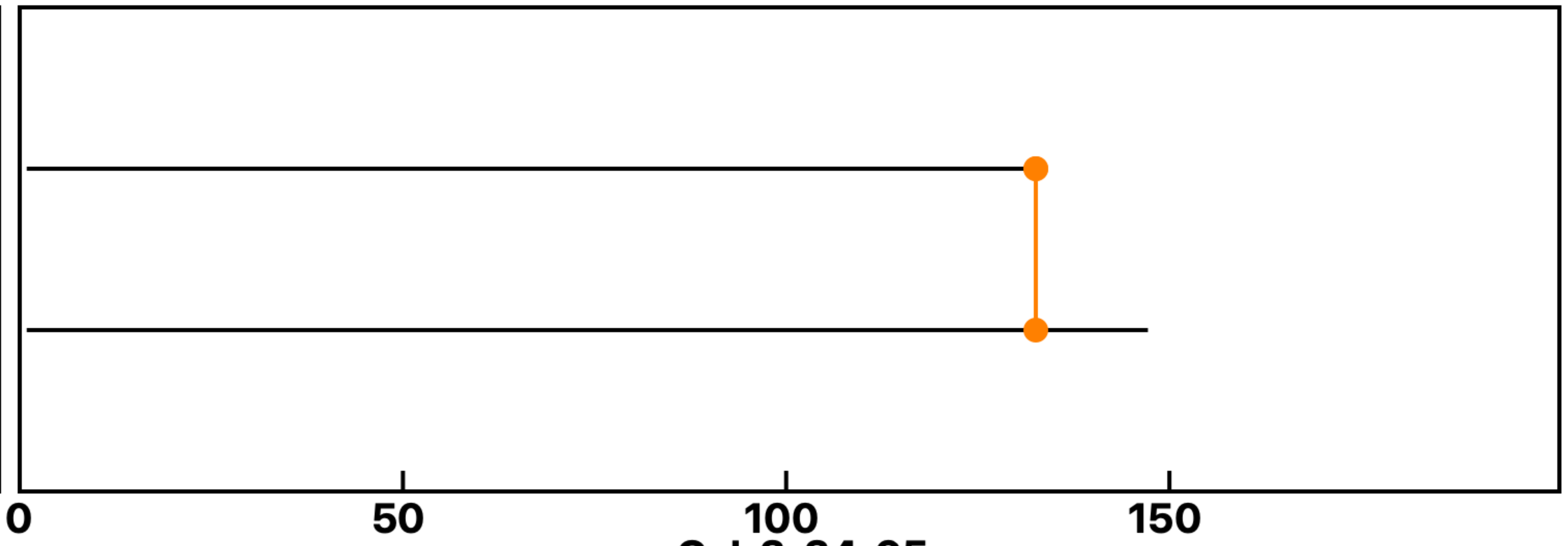

+Gal-3-34, 65

**Analysis: Gal-3KO Gal-3-GlcNAc, Treat.: +Gal-3, Cell: Myoblasts**

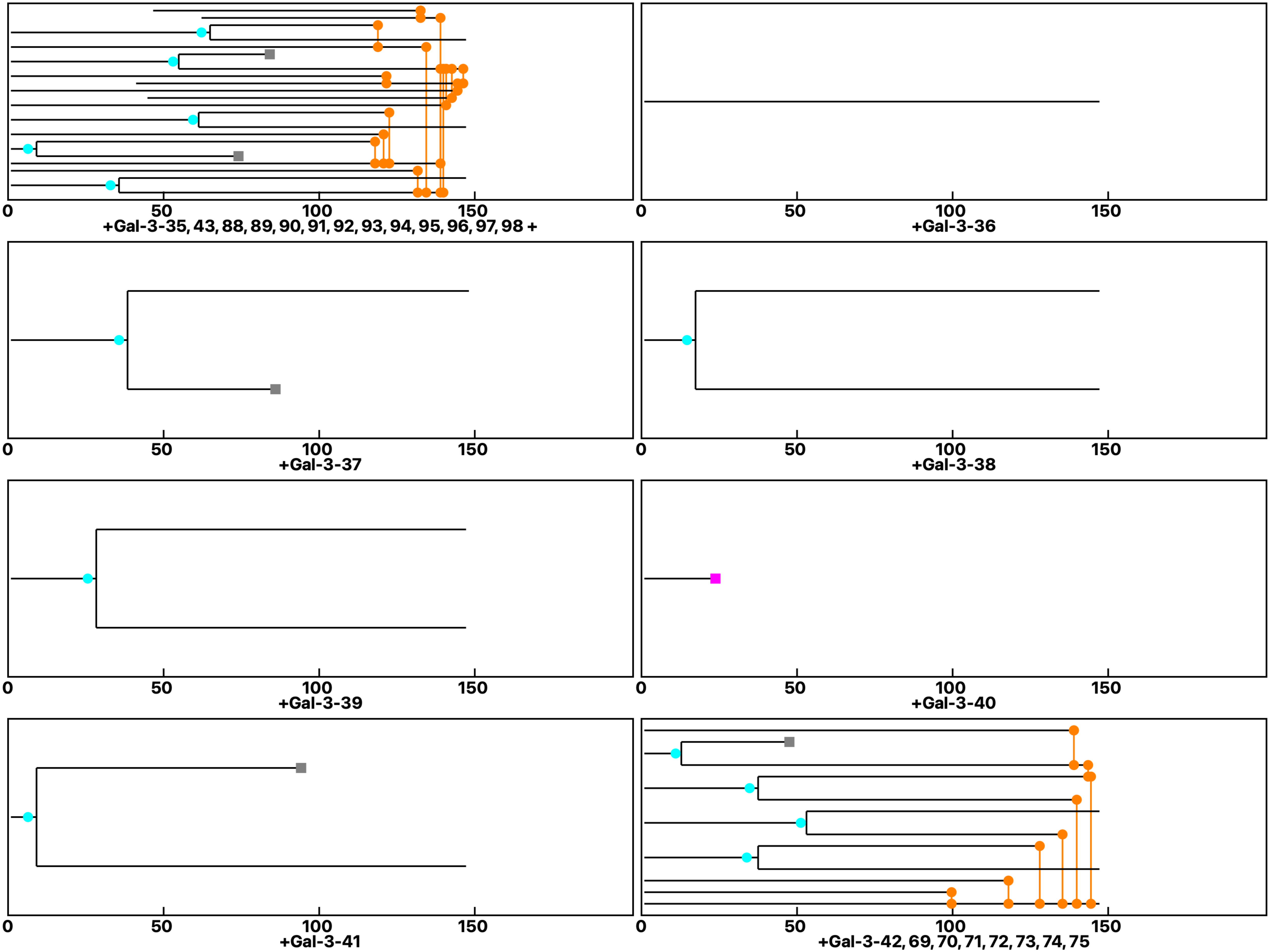

Analysis: Gal-3KO Gal-3-GlcNAc, Treat.: +Gal-3, Cell: Myoblasts

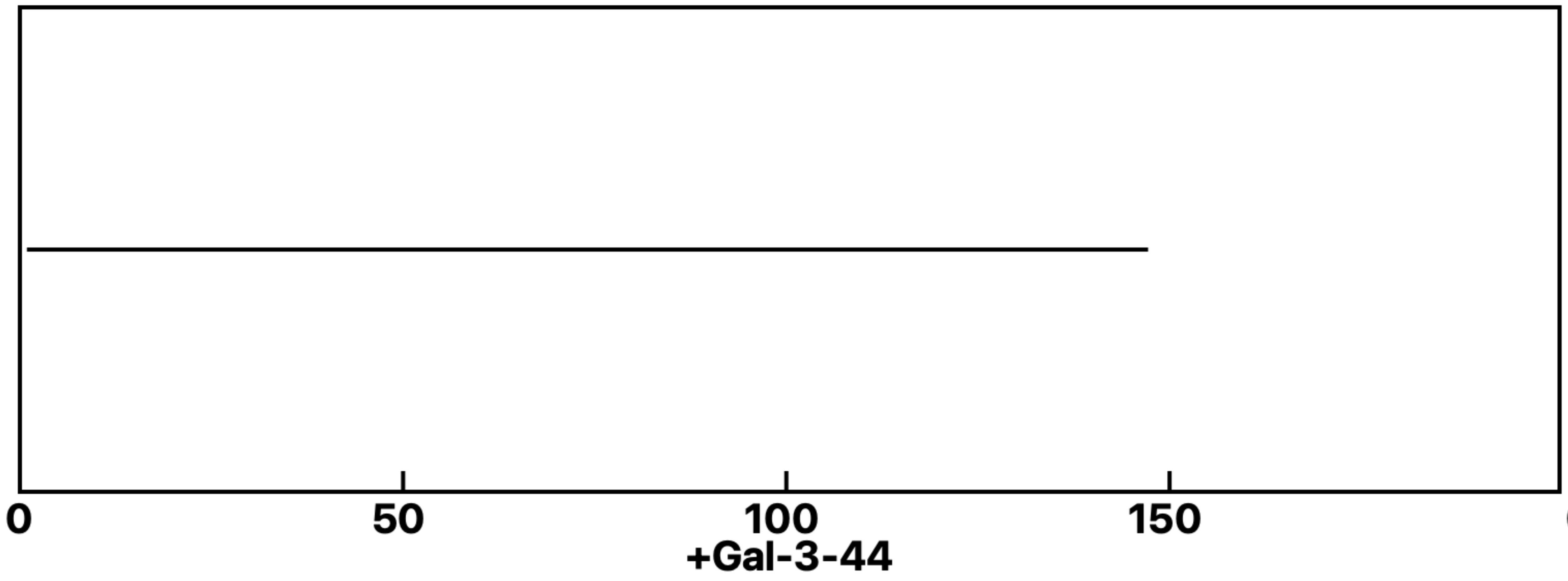


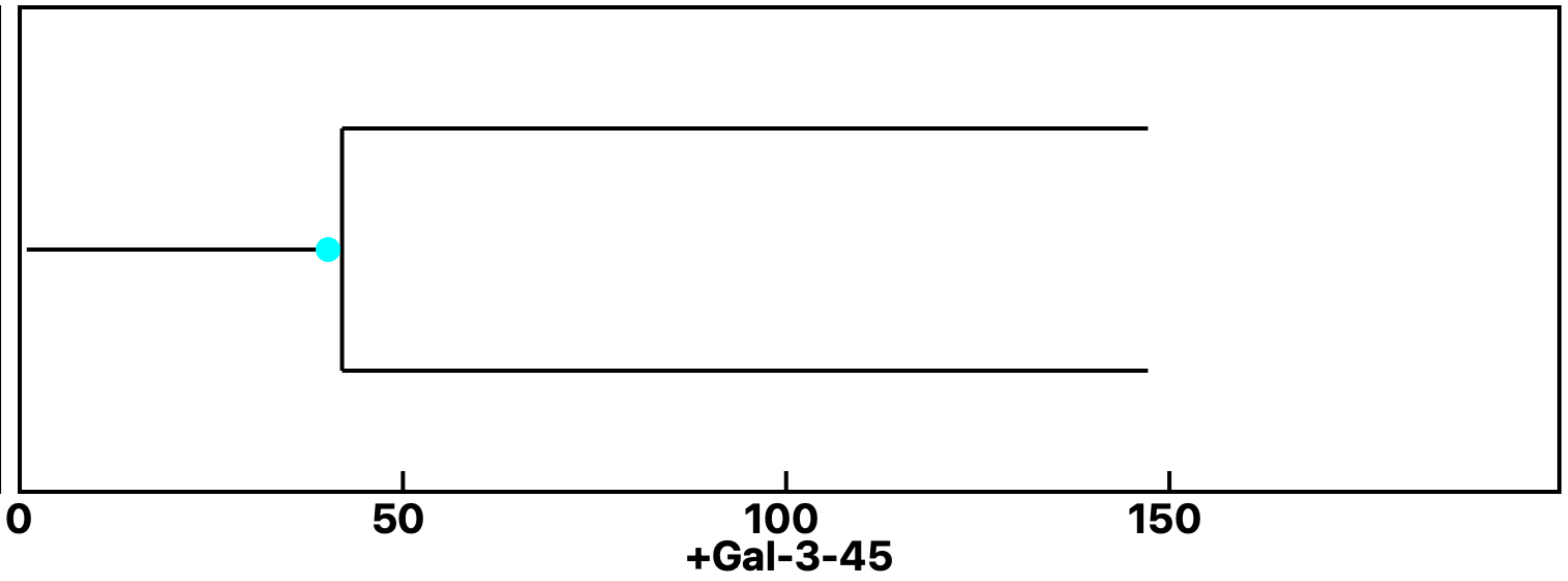


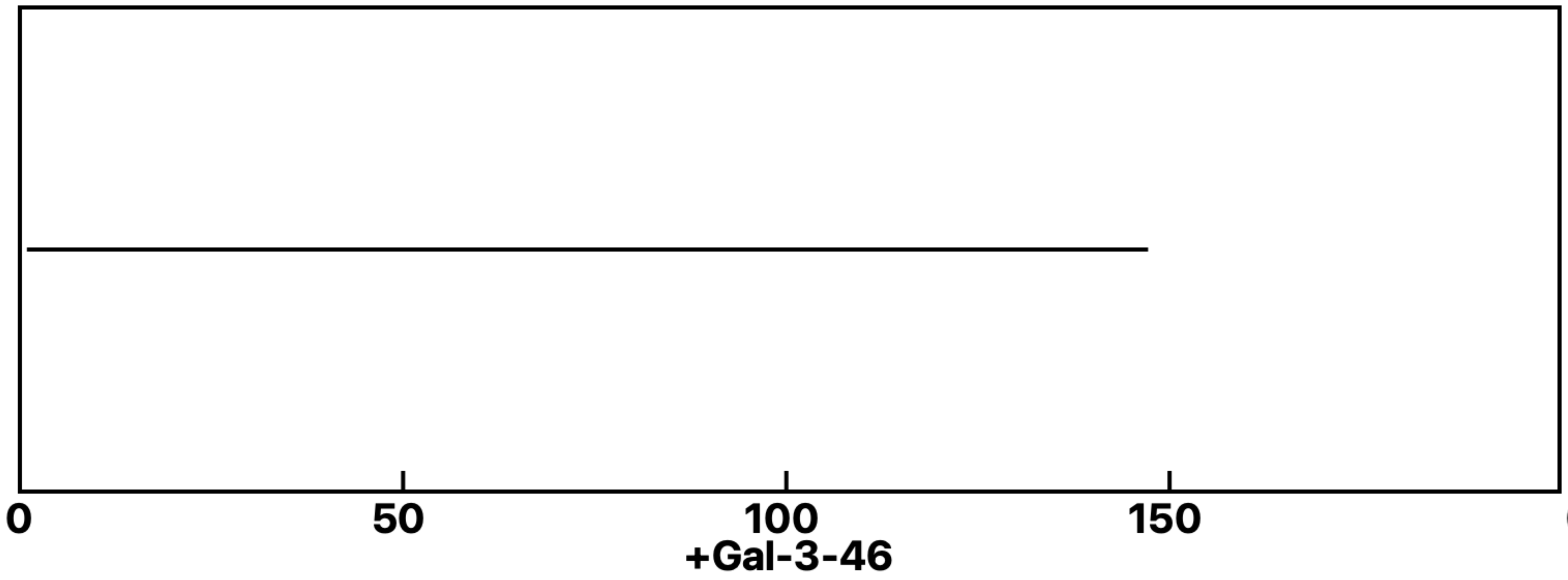


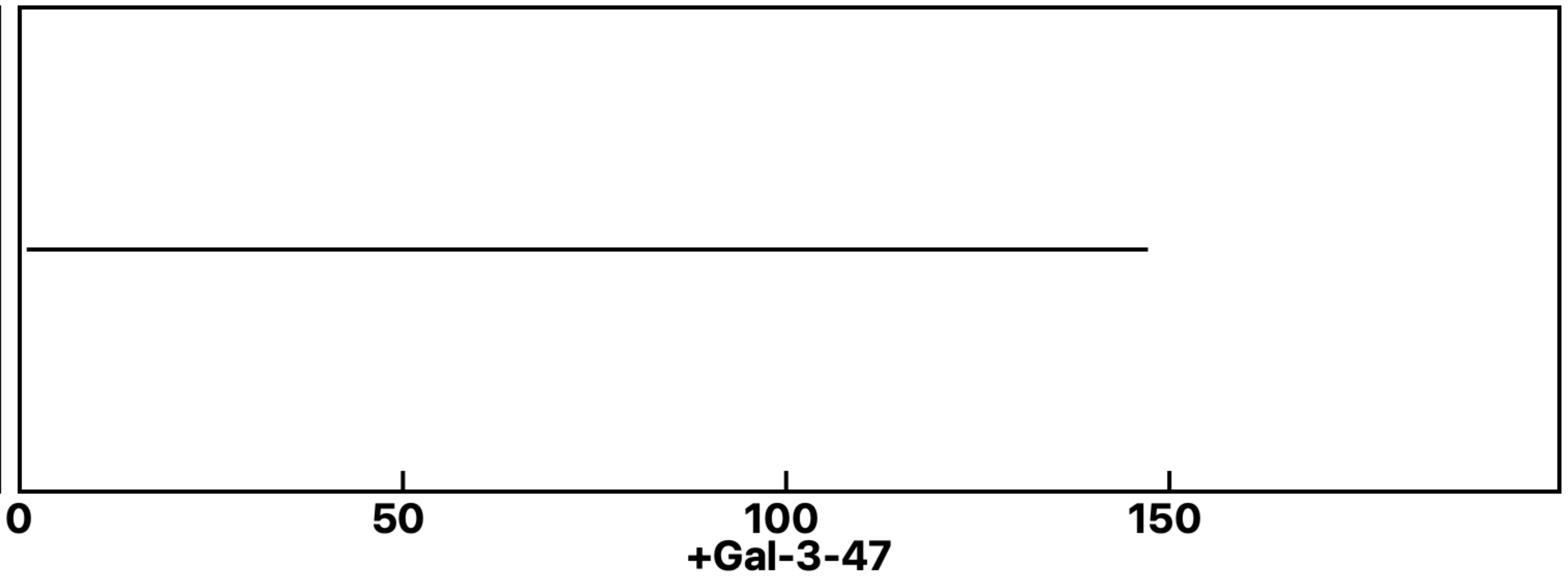


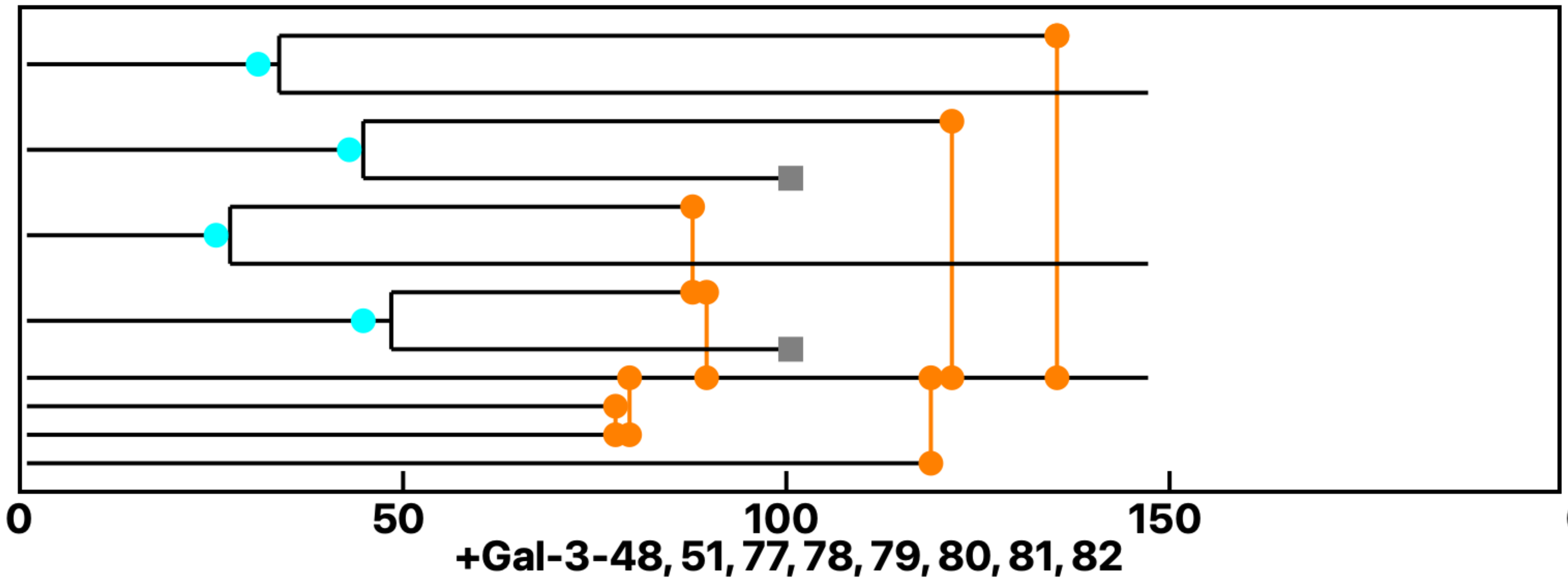


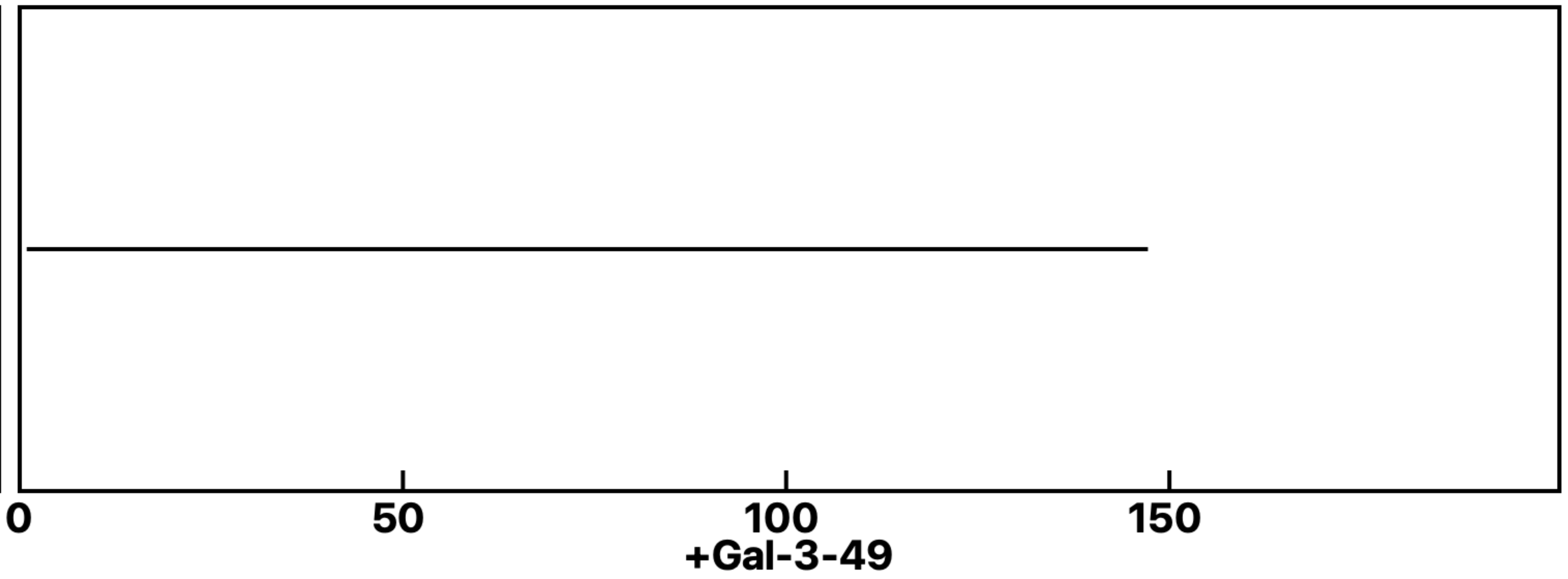


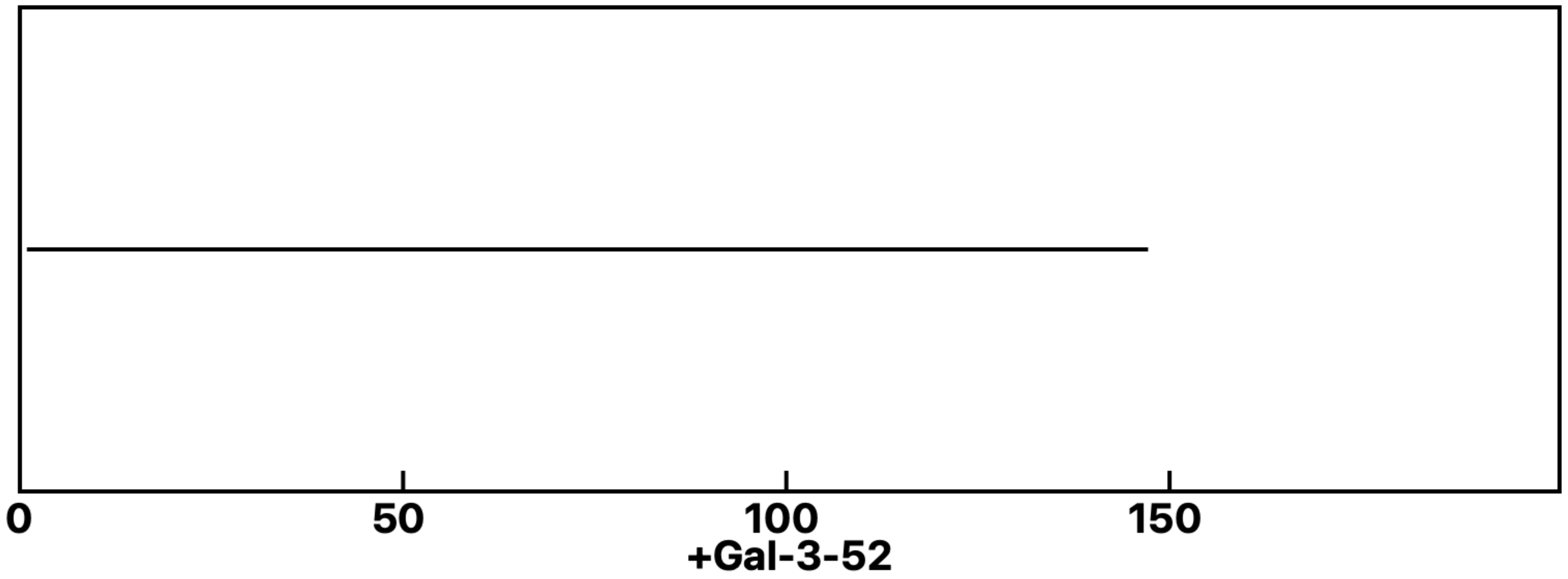
