## Supplemental data 11 for "N-Acetylglucosamine Facilitates Coordinated Myoblast Flow, Forming the Foundation for Efficient Myogenesis"

**Analysis: Gal-3KO Gal-3-GlcNAc, Treat.: +GlcNAc, Cell: Myoblasts**

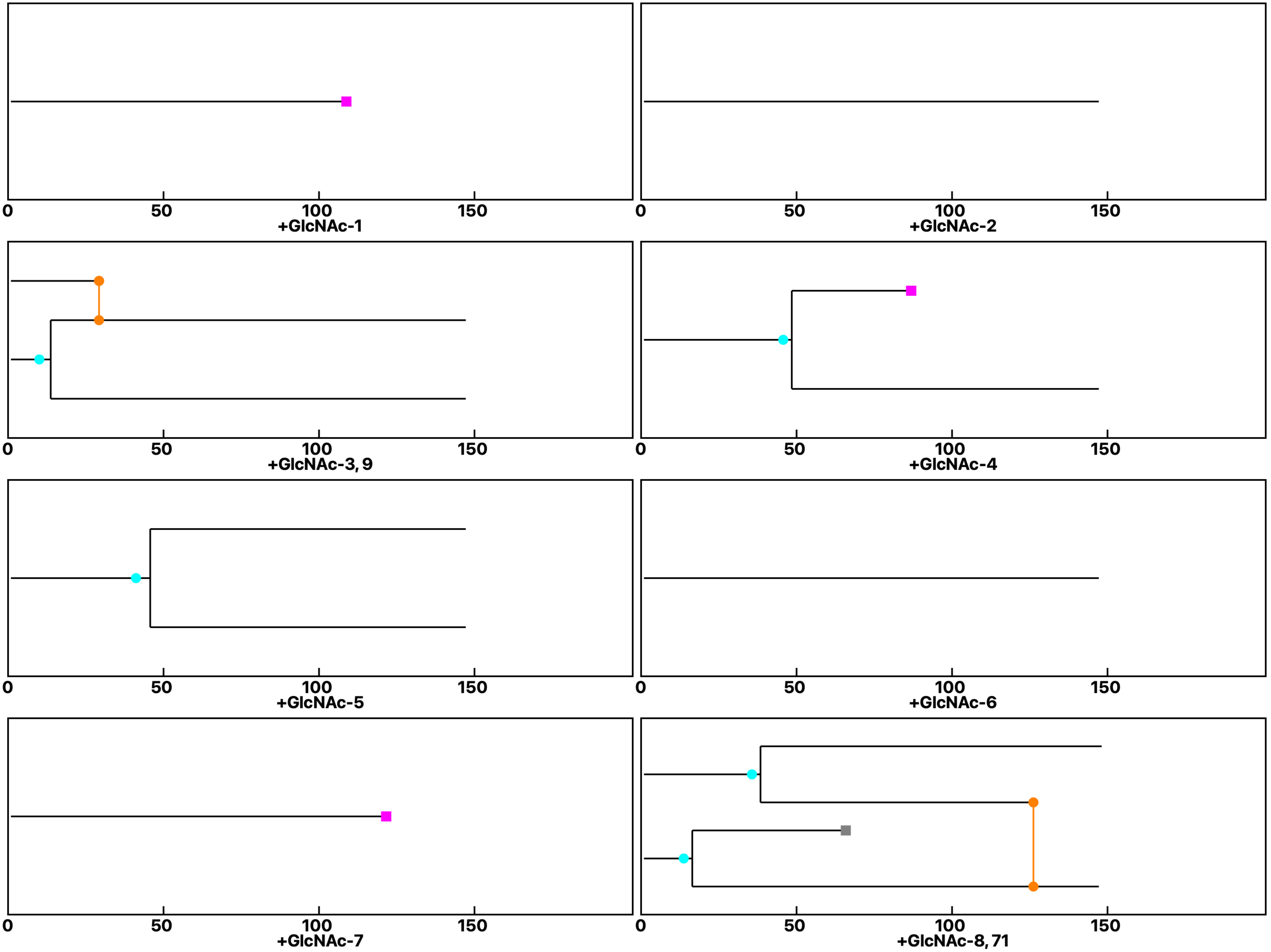

Analysis: Gal-3KO Gal-3-GlcNAc, Treat.: +GlcNAc, Cell: Myoblasts

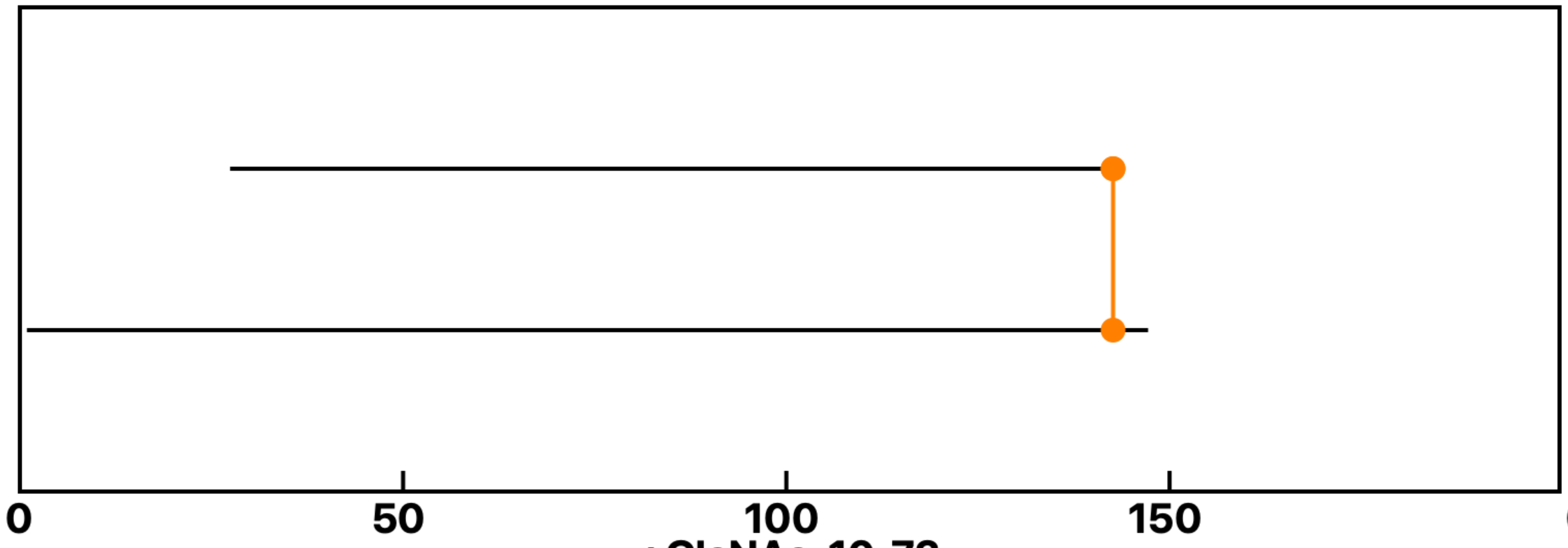


+GlcNAc-10, 72

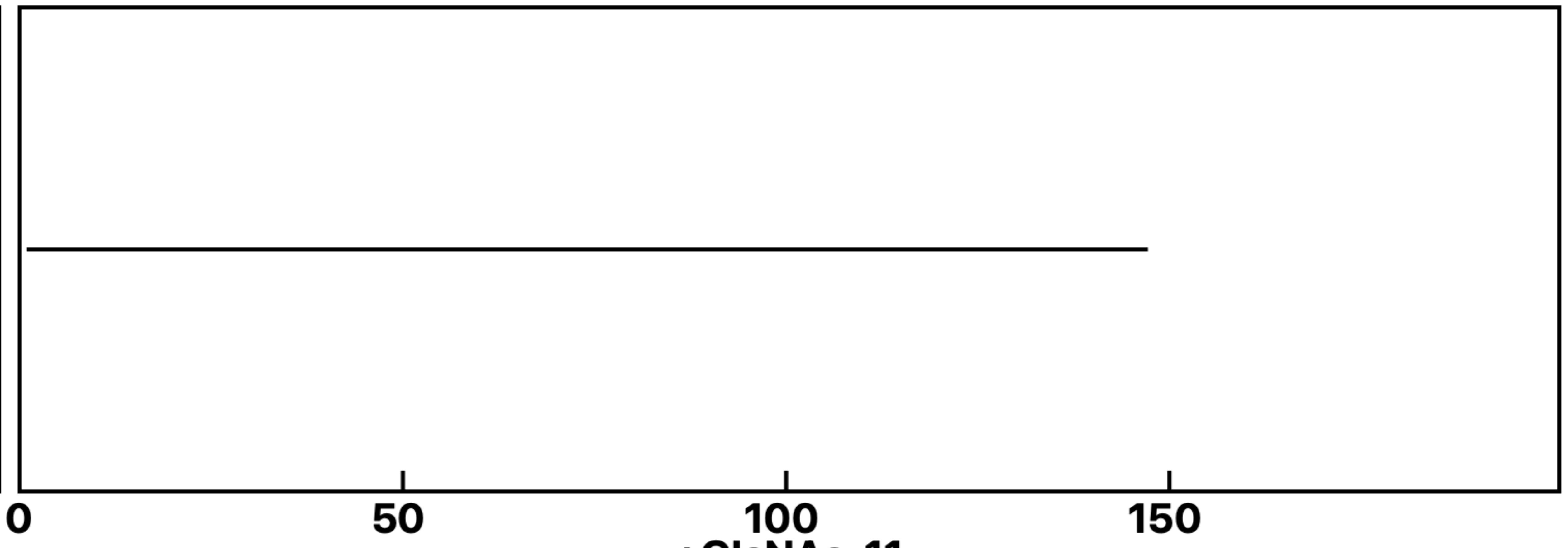


+GlcNAc-11

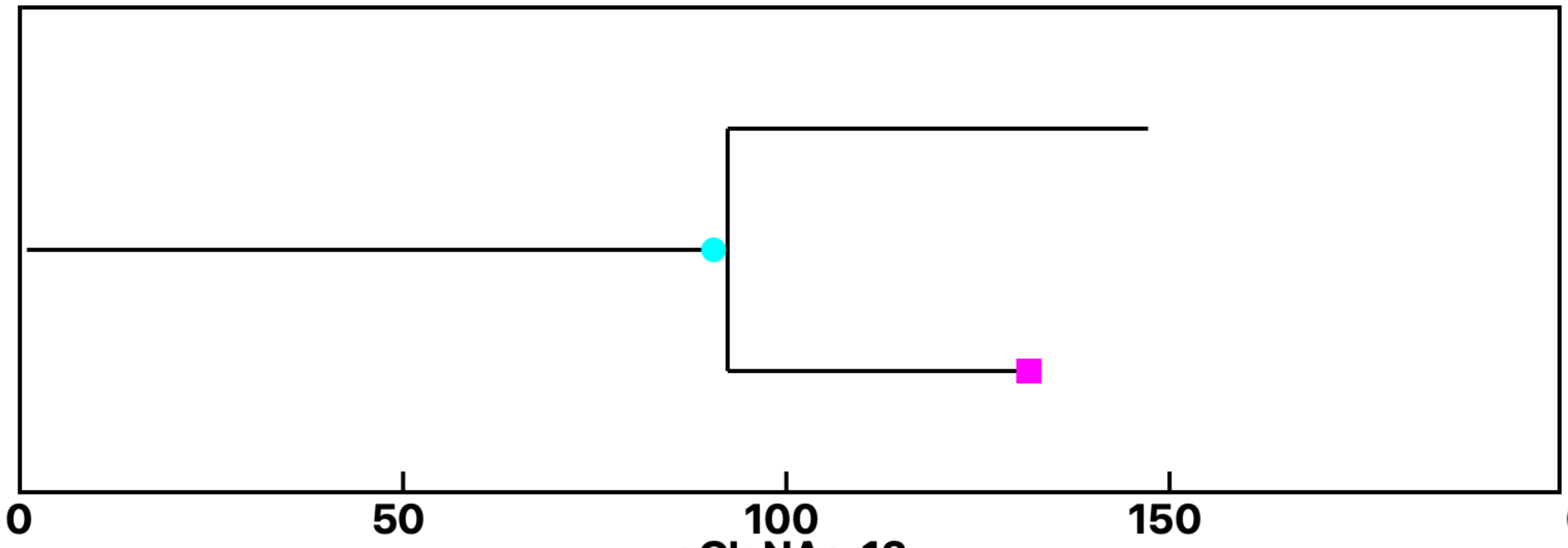


+GlcNAc-12

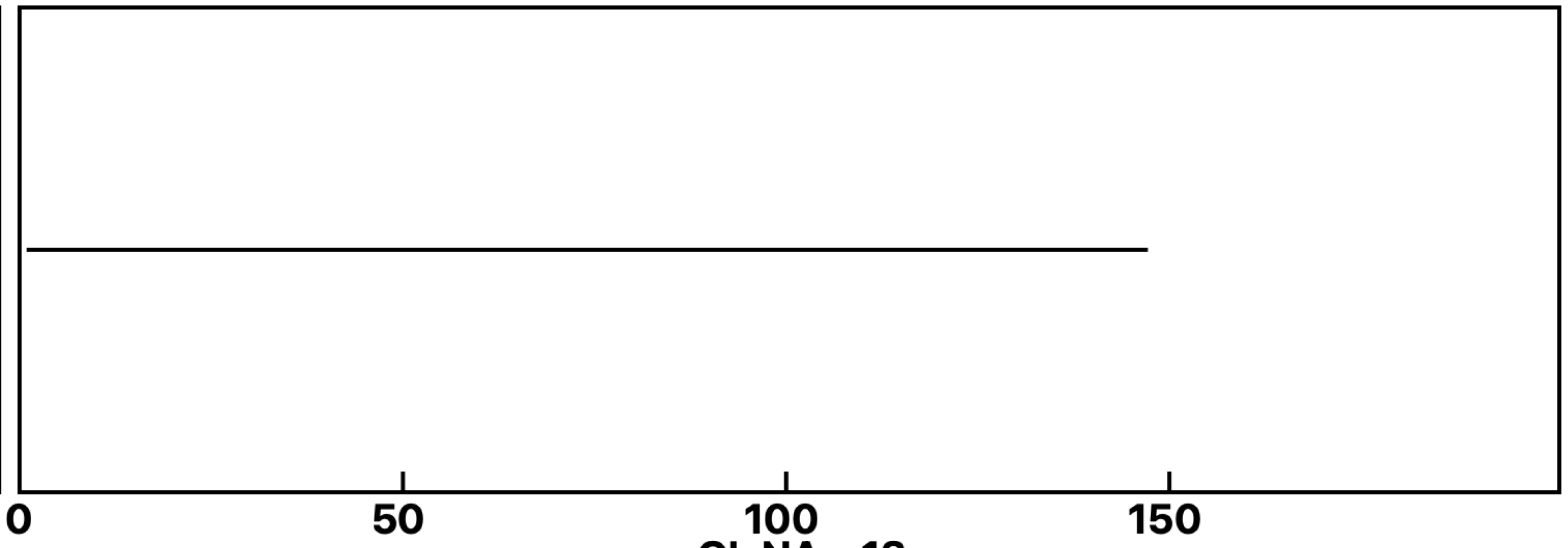


+GlcNAc-13

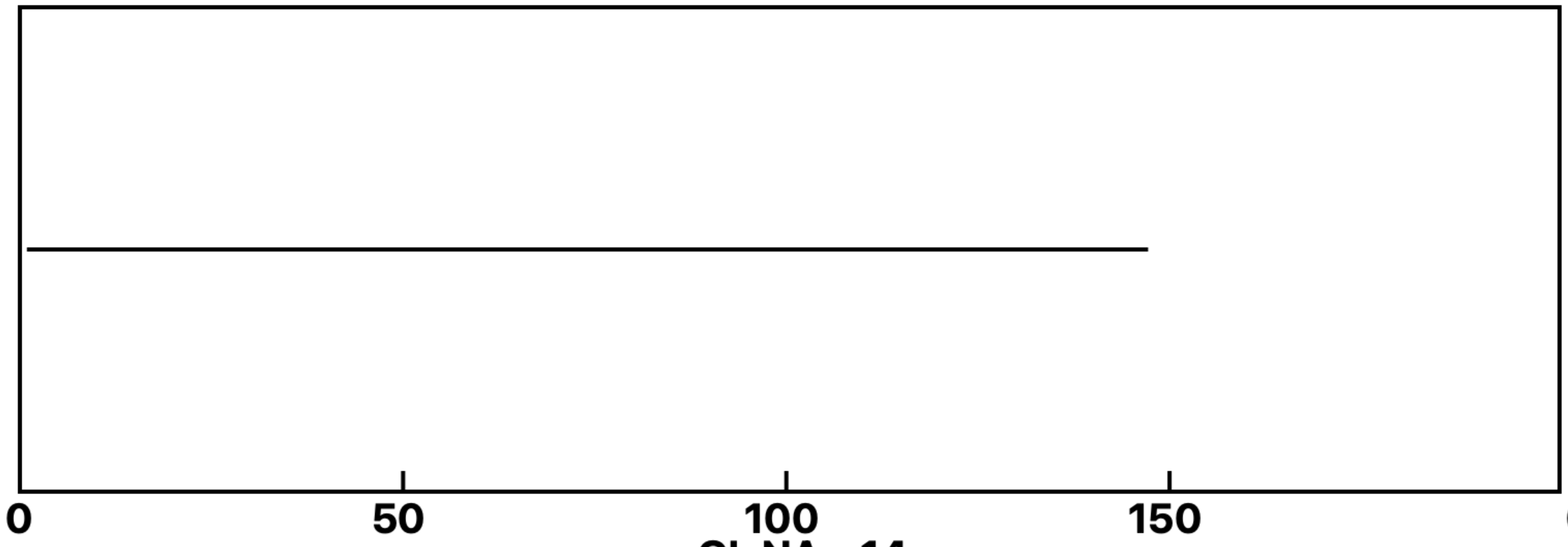


+GlcNAc-14

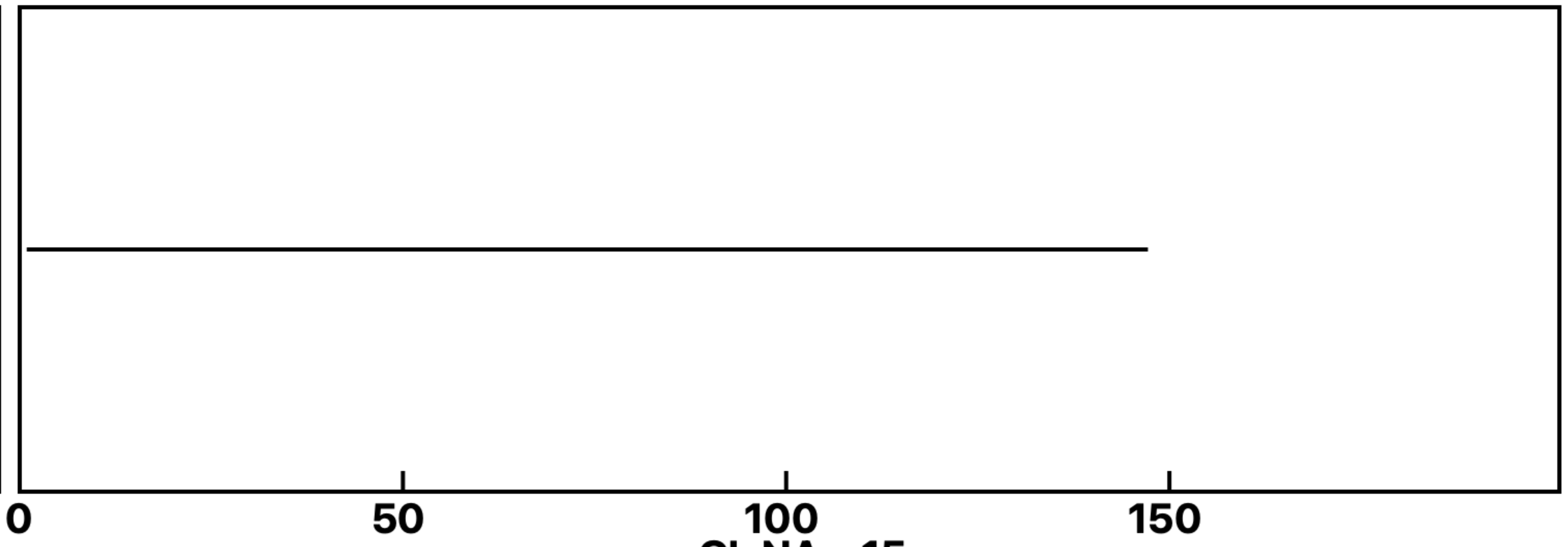


+GlcNAc-15

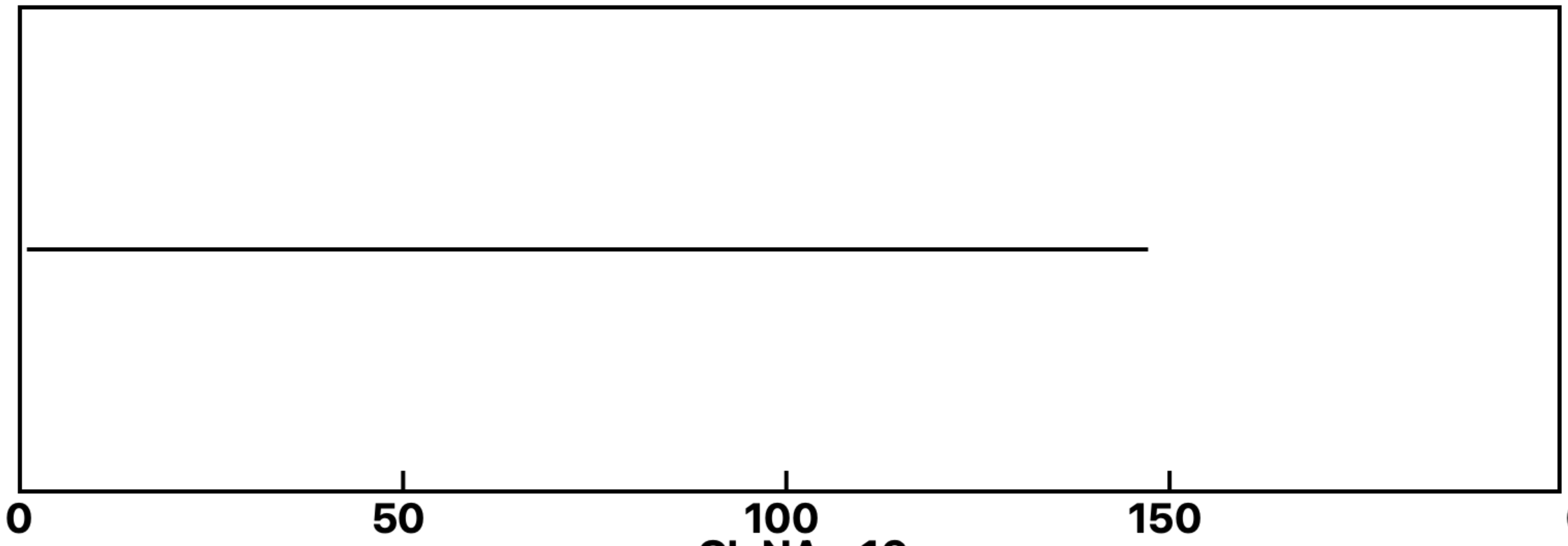


+GlcNAc-16

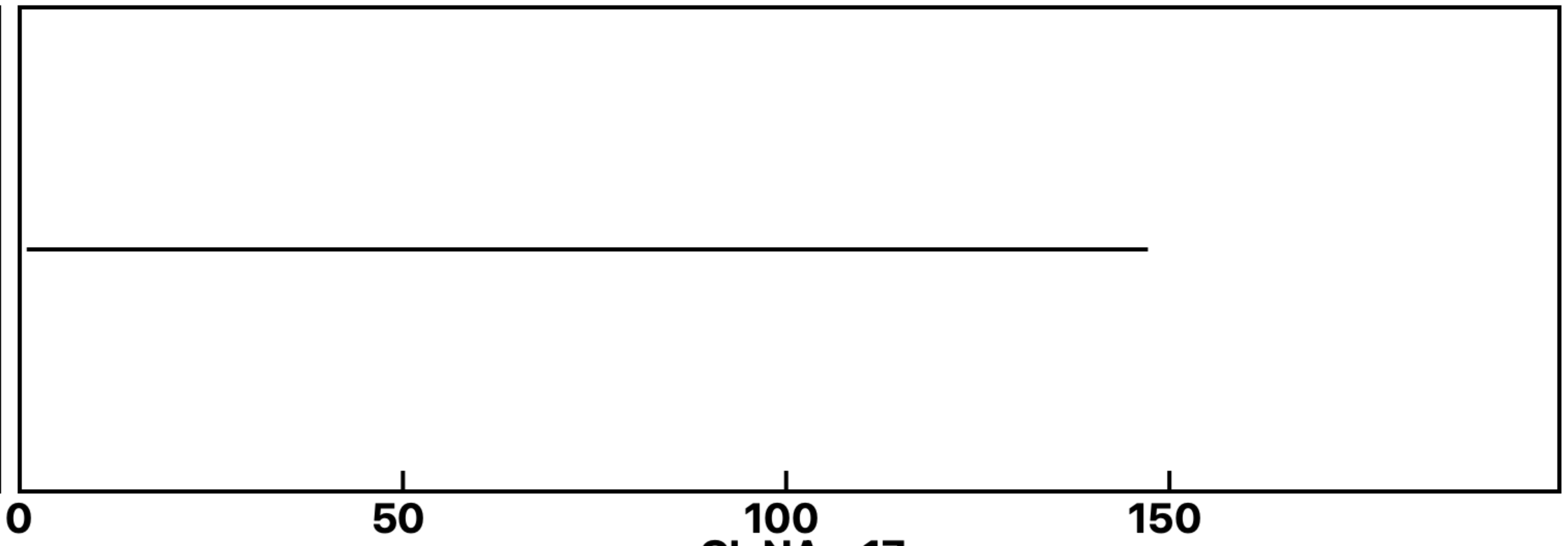


+GlcNAc-17

**Analysis: Gal-3KO Gal-3-GlcNAc, Treat.: +GlcNAc, Cell: Myoblasts**

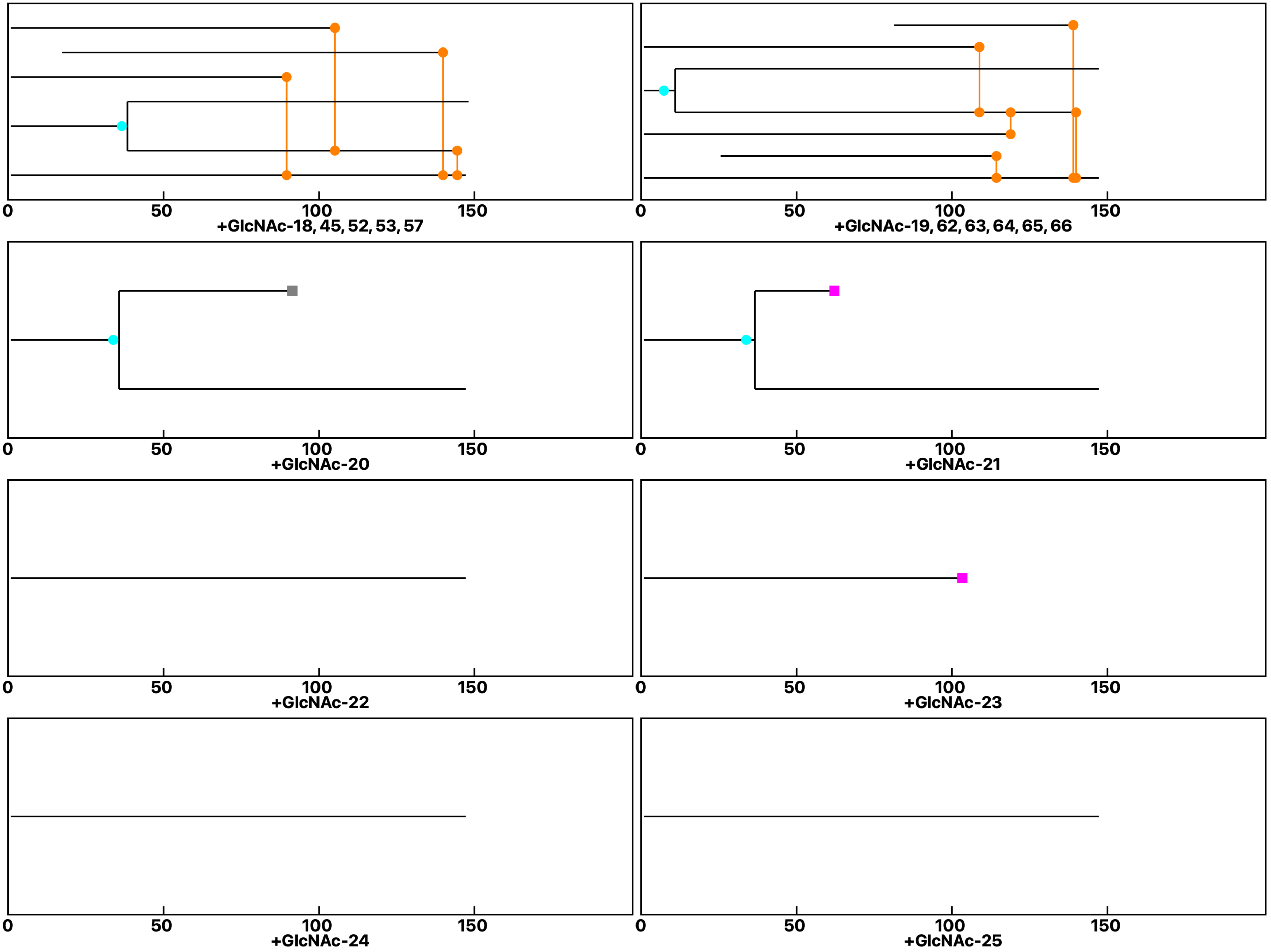

**Analysis: Gal-3KO Gal-3-GlcNAc, Treat.: +GlcNAc, Cell: Myoblasts**

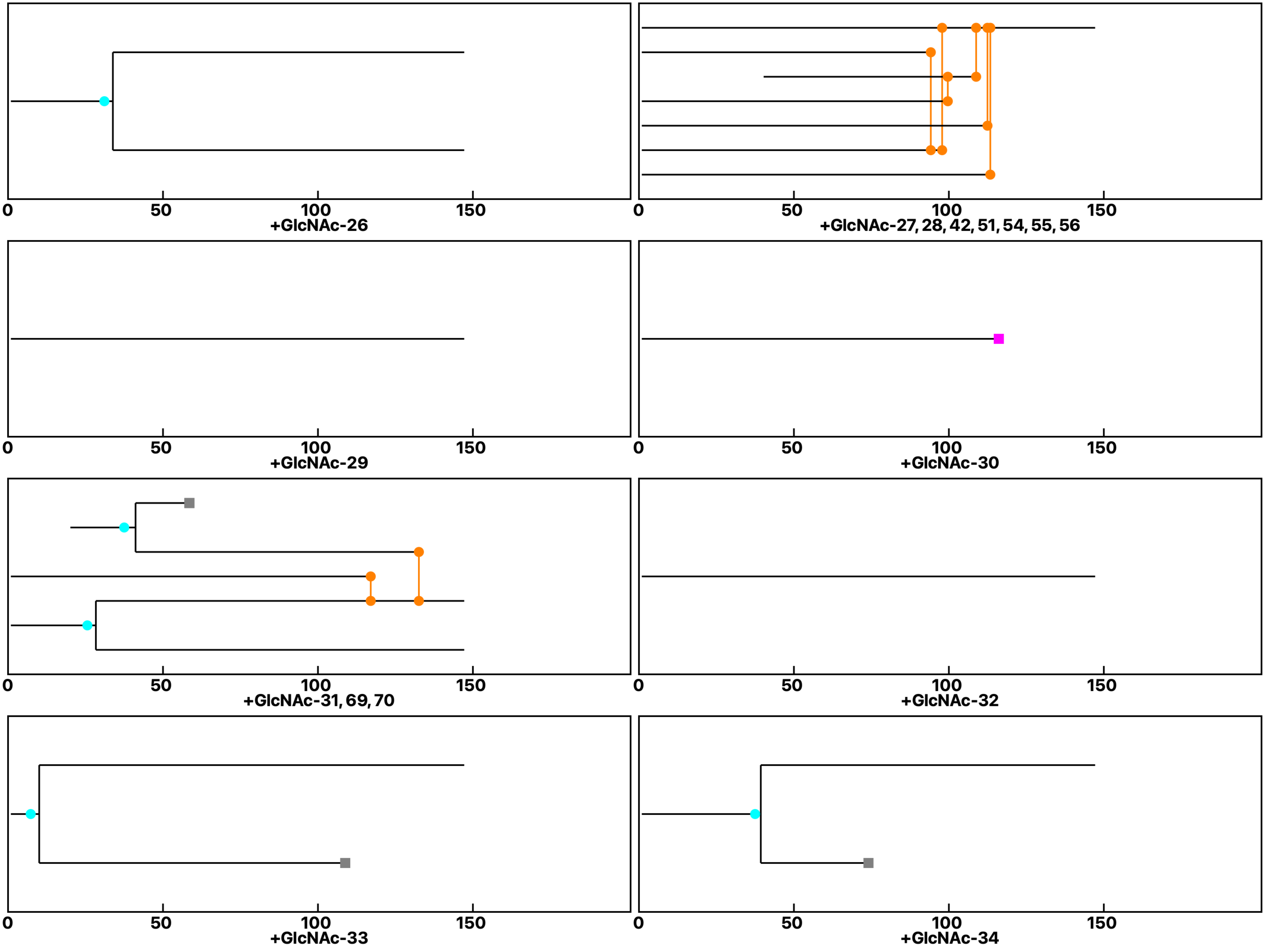

**Analysis: Gal-3KO Gal-3-GlcNAc, Treat.: +GlcNAc, Cell: Myoblasts**

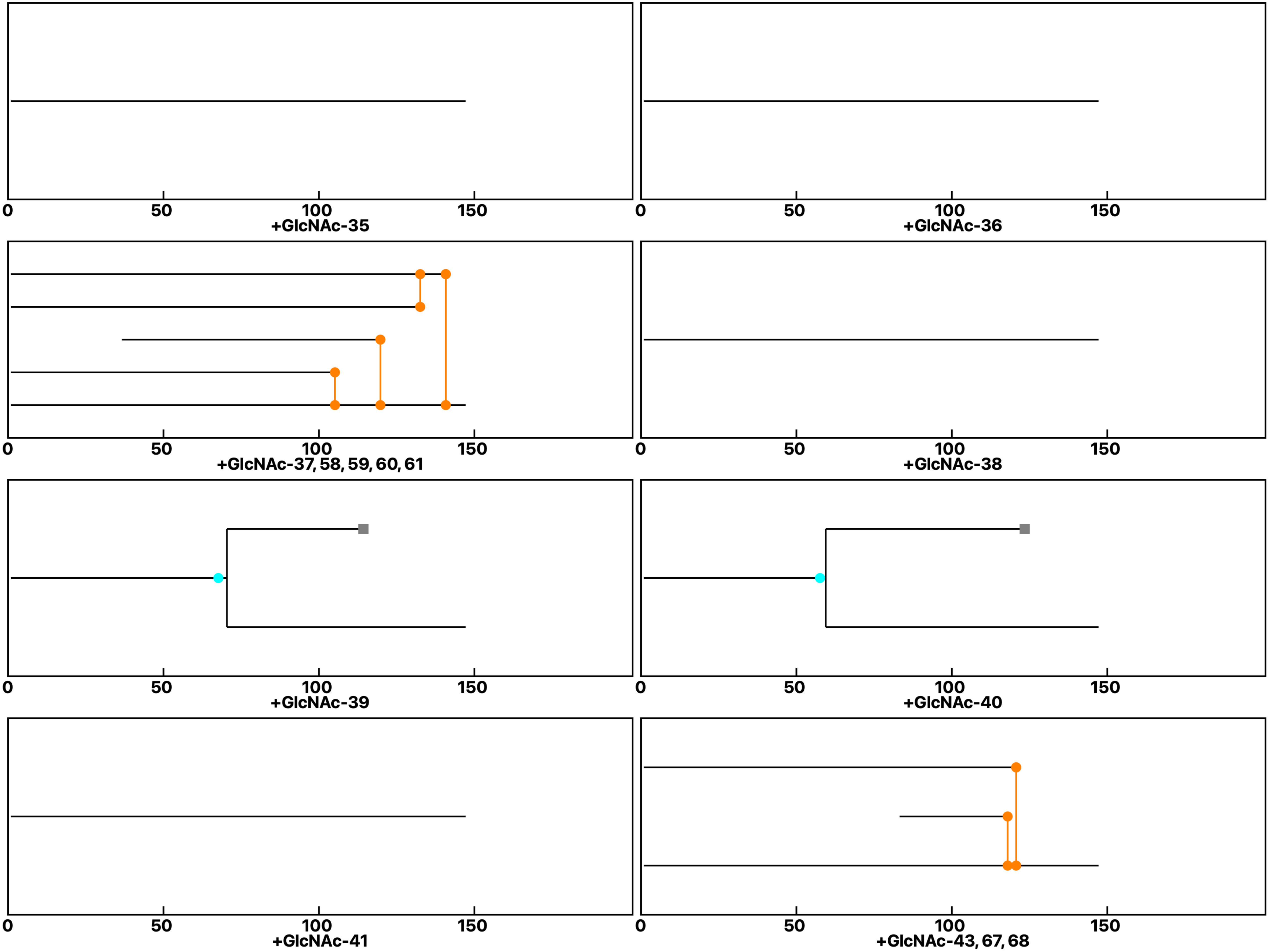

**Analysis: Gal-3KO Gal-3-GlcNAc, Treat.: +GlcNAc, Cell: Myoblasts**

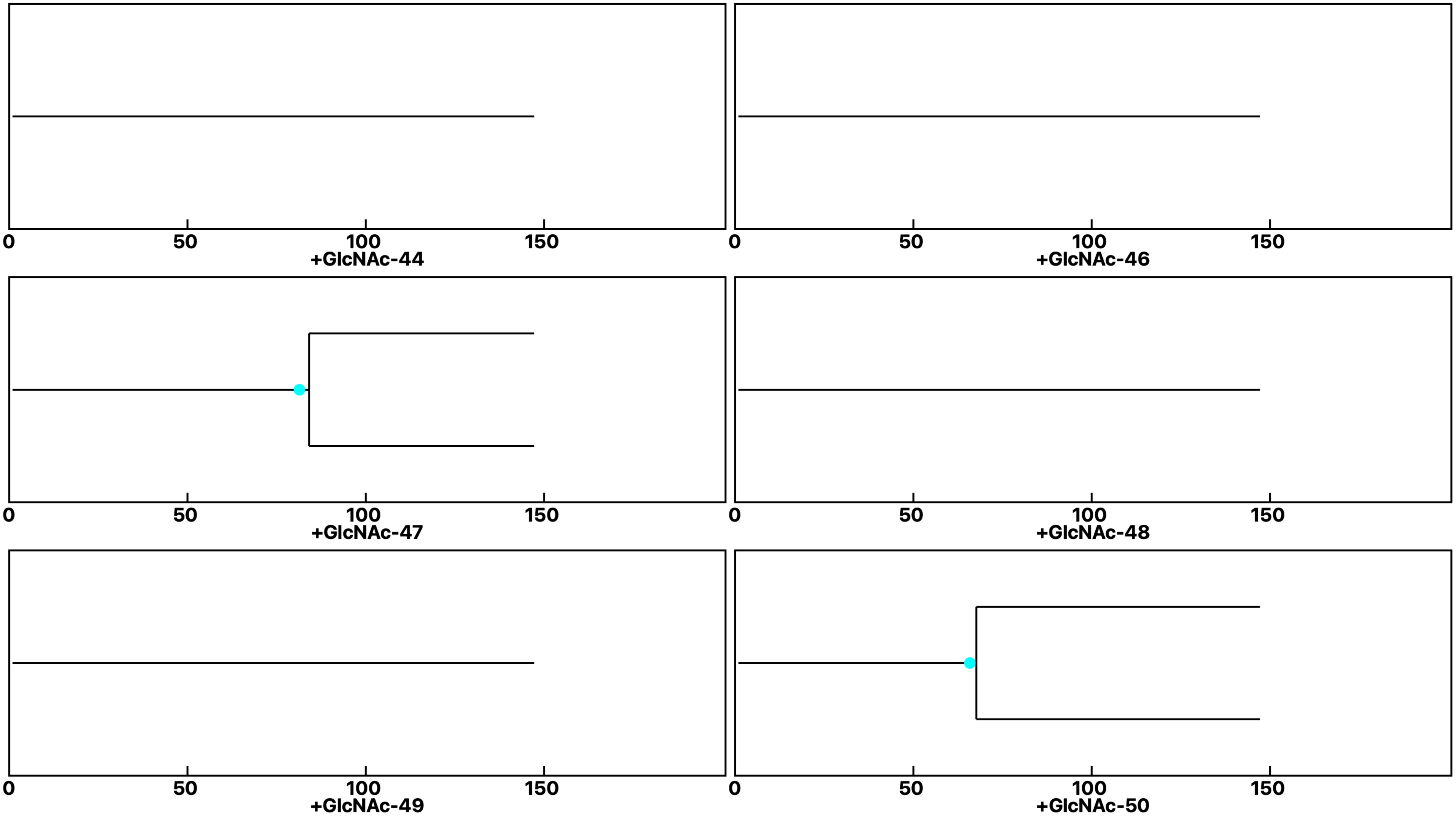
