## Supplemental data 12 for "N-Acetylglucosamine Facilitates Coordinated Myoblast Flow, Forming the Foundation for Efficient Myogenesis"

### Analysis: Gal-3KO Gal-3-GlcNAc, Treat.: +Gal-3+GlcNAc, Cell: Myoblasts

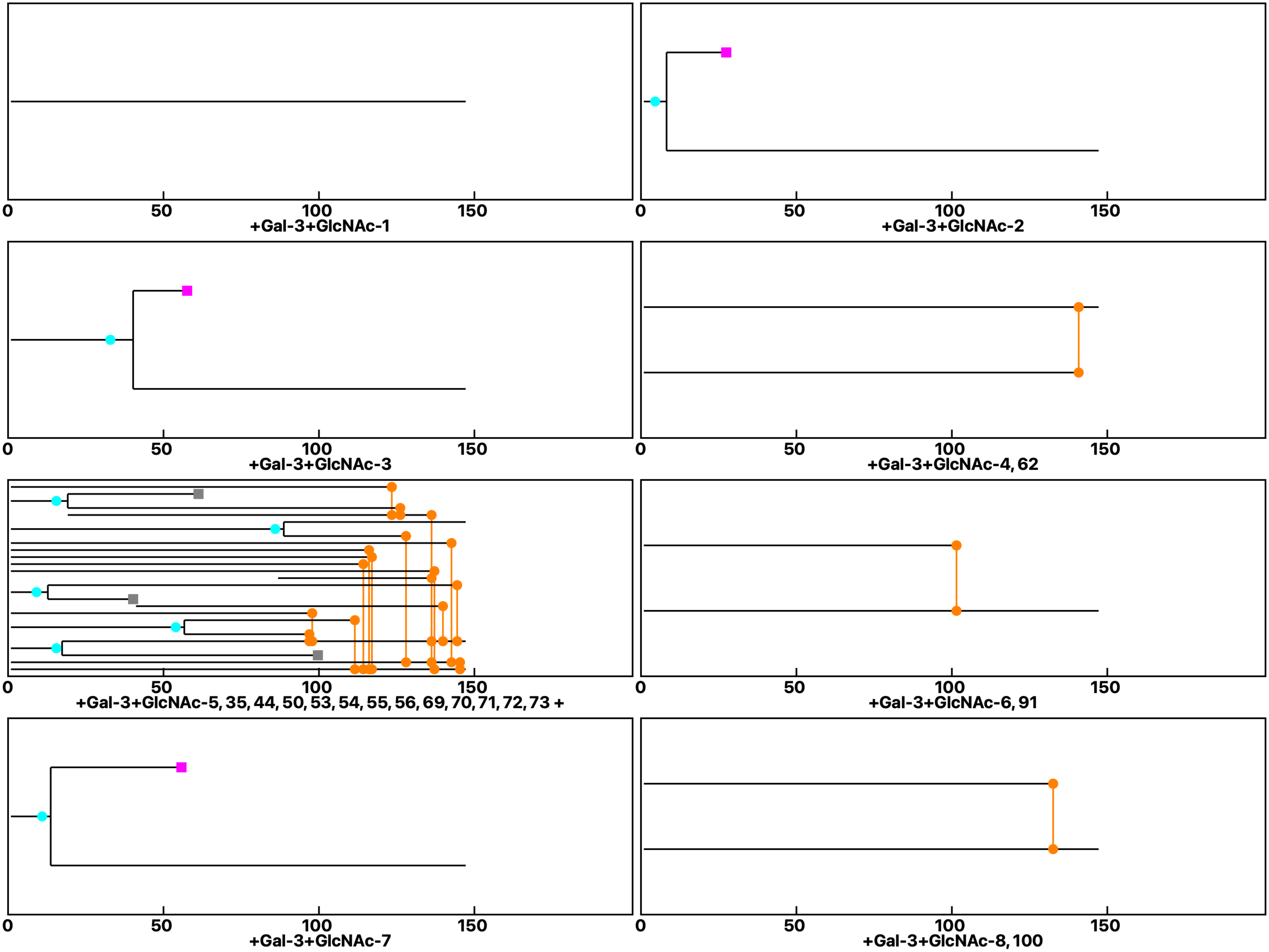

**Analysis: Gal-3KO Gal-3-GlcNAc, Treat.: +Gal-3+GlcNAc, Cell: Myoblasts**

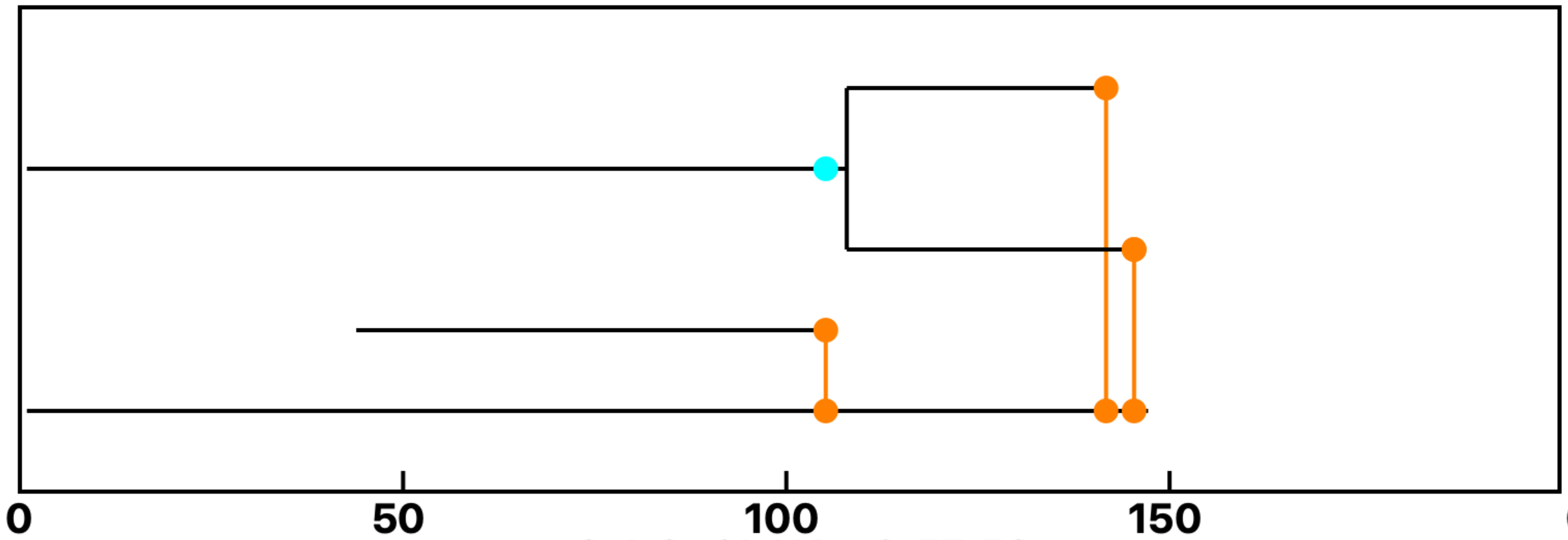


+Gal-3+GlcNAc-9, 57, 58

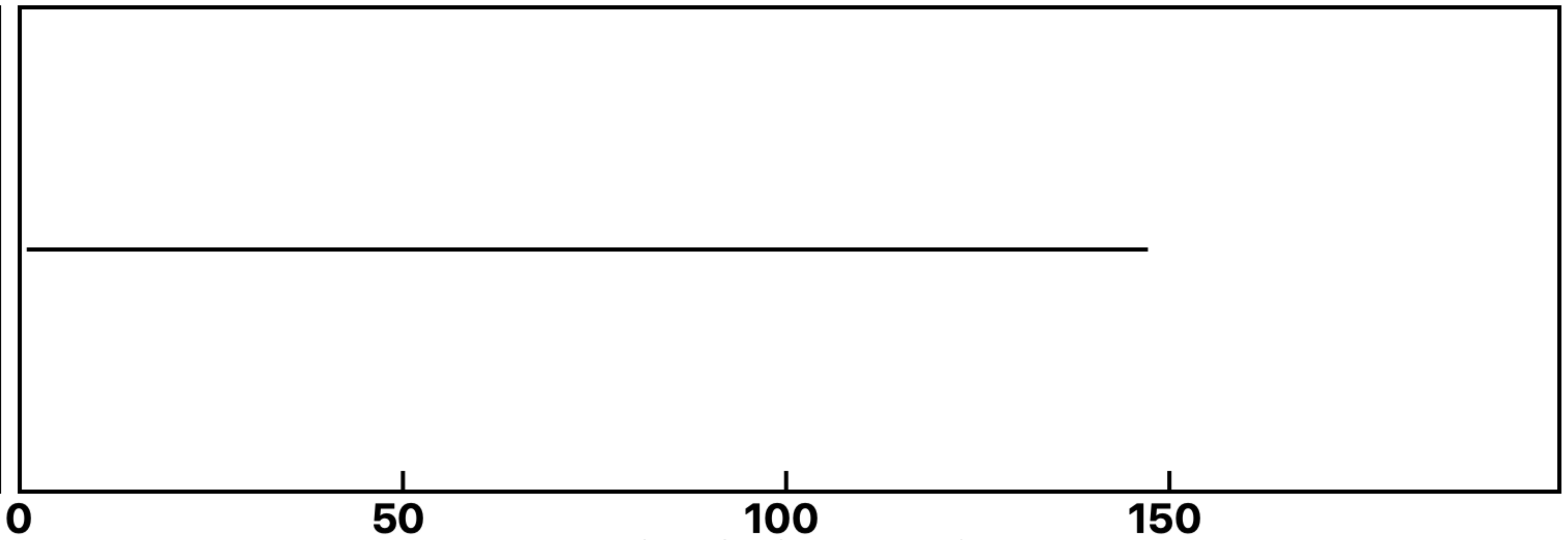


+Gal-3+GlcNAc-10

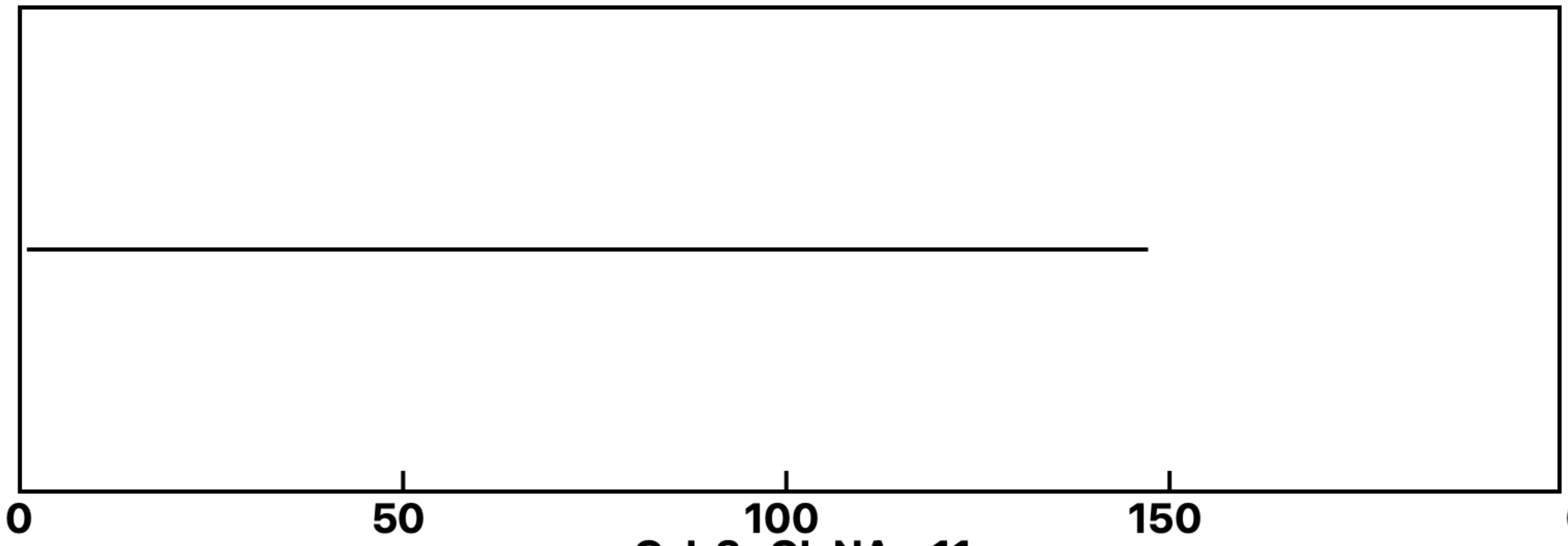


+Gal-3+GlcNAc-11

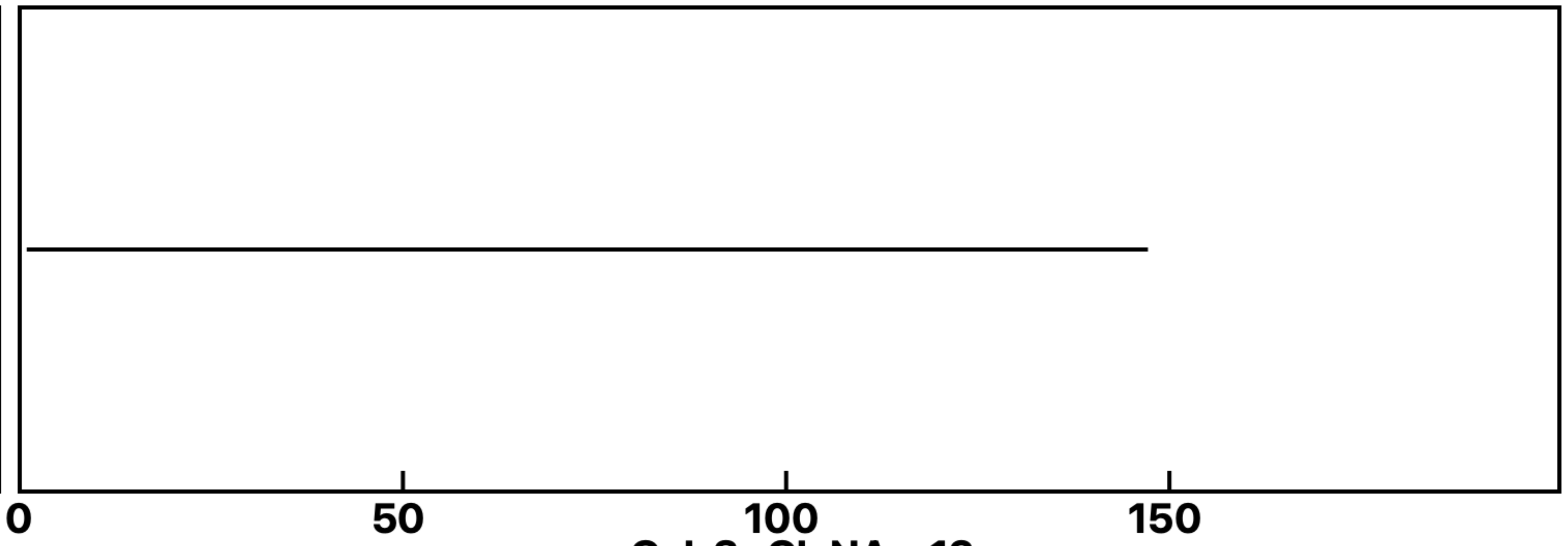


+Gal-3+GlcNAc-12

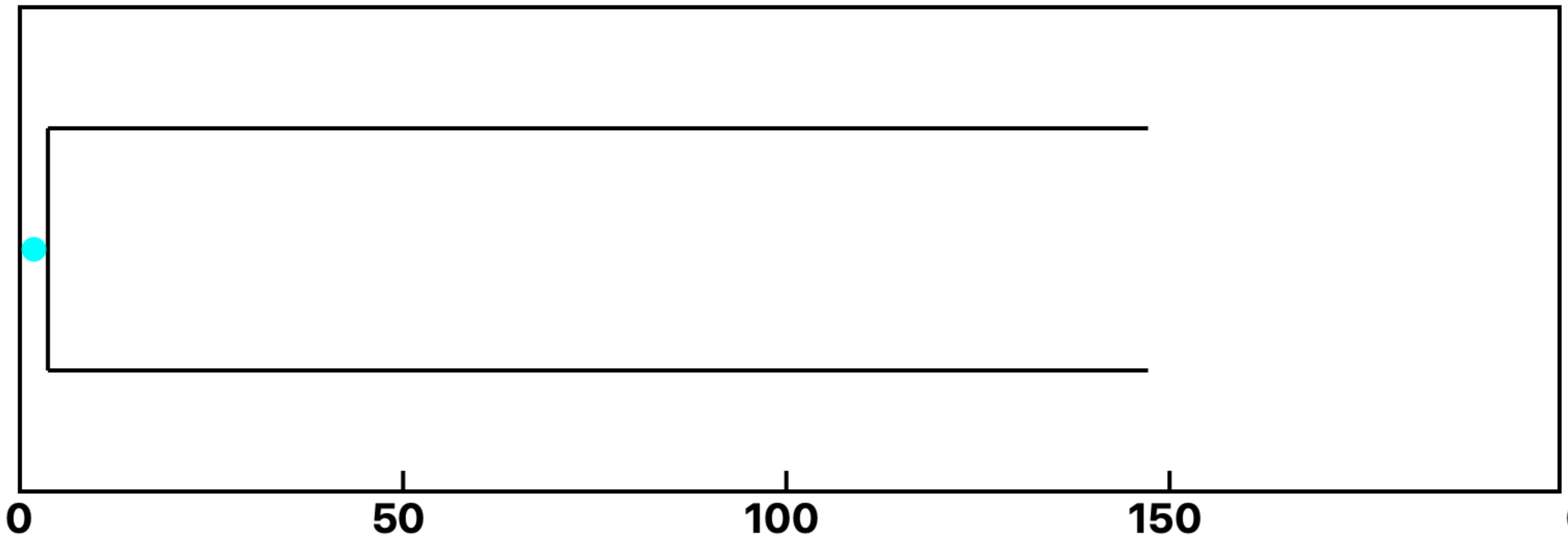


+Gal-3+GlcNAc-13

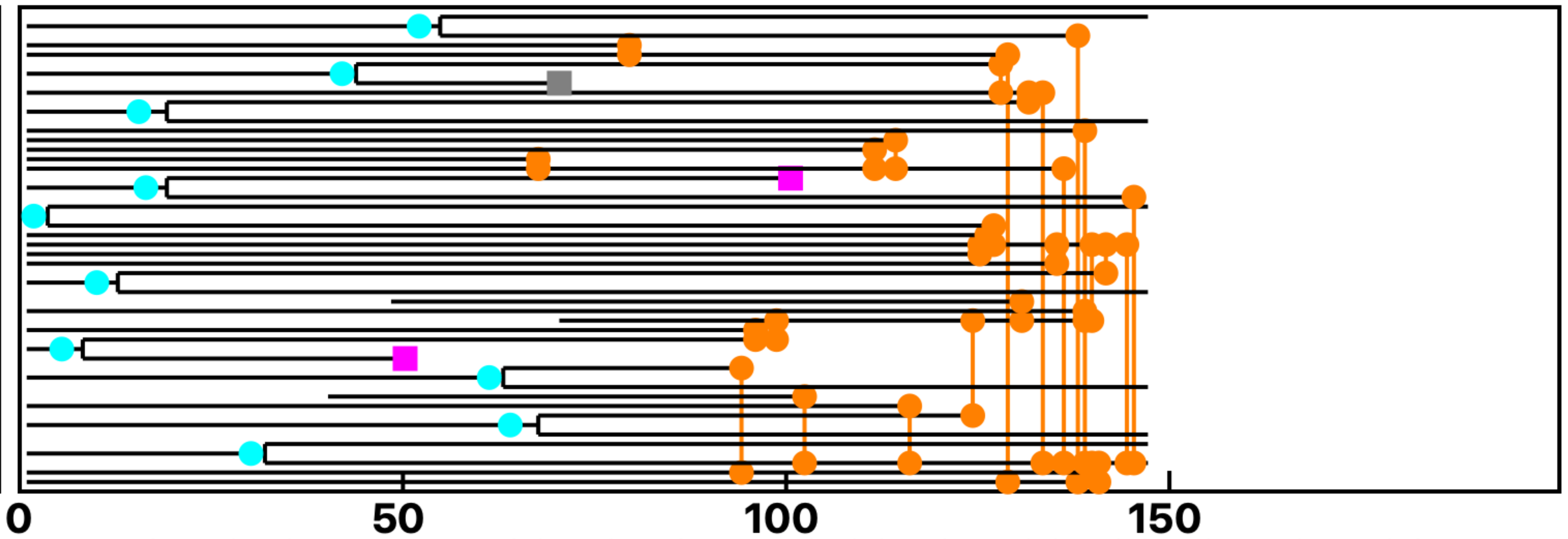


+Gal-3+GlcNAc-14, 30, 46, 49, 51, 52, 90, 101, 102, 103, 104, 105, 106 +

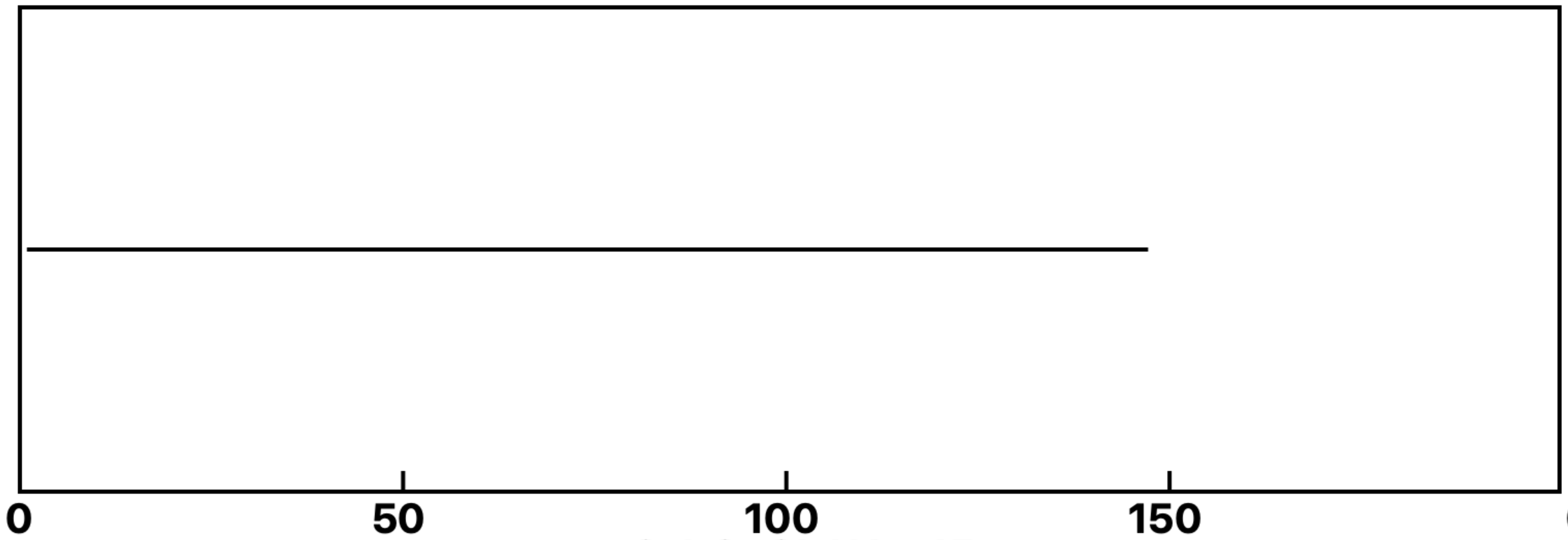


+Gal-3+GlcNAc-15

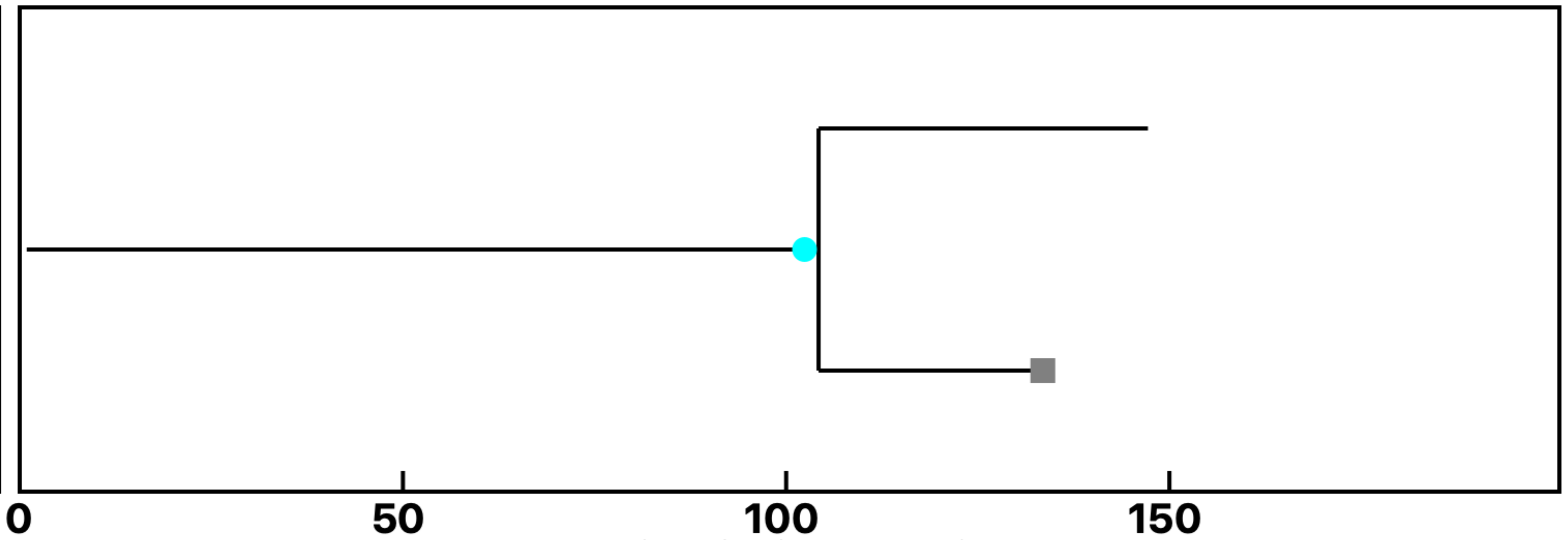


+Gal-3+GlcNAc-16

**Analysis: Gal-3KO Gal-3-GlcNAc, Treat.: +Gal-3+GlcNAc, Cell: Myoblasts**

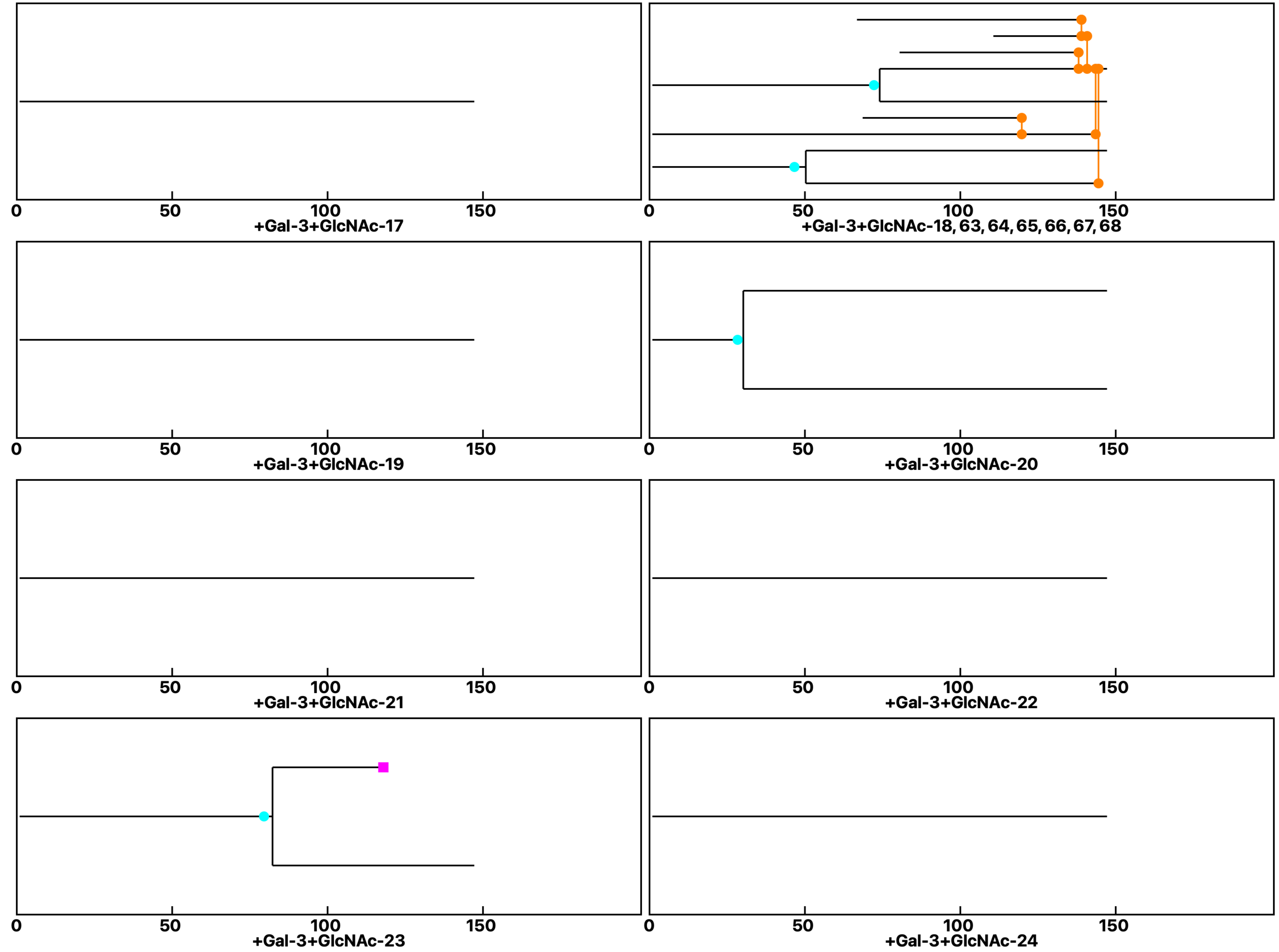

Analysis: Gal-3KO Gal-3-GlcNAc, Treat.: +Gal-3+GlcNAc, Cell: Myoblasts

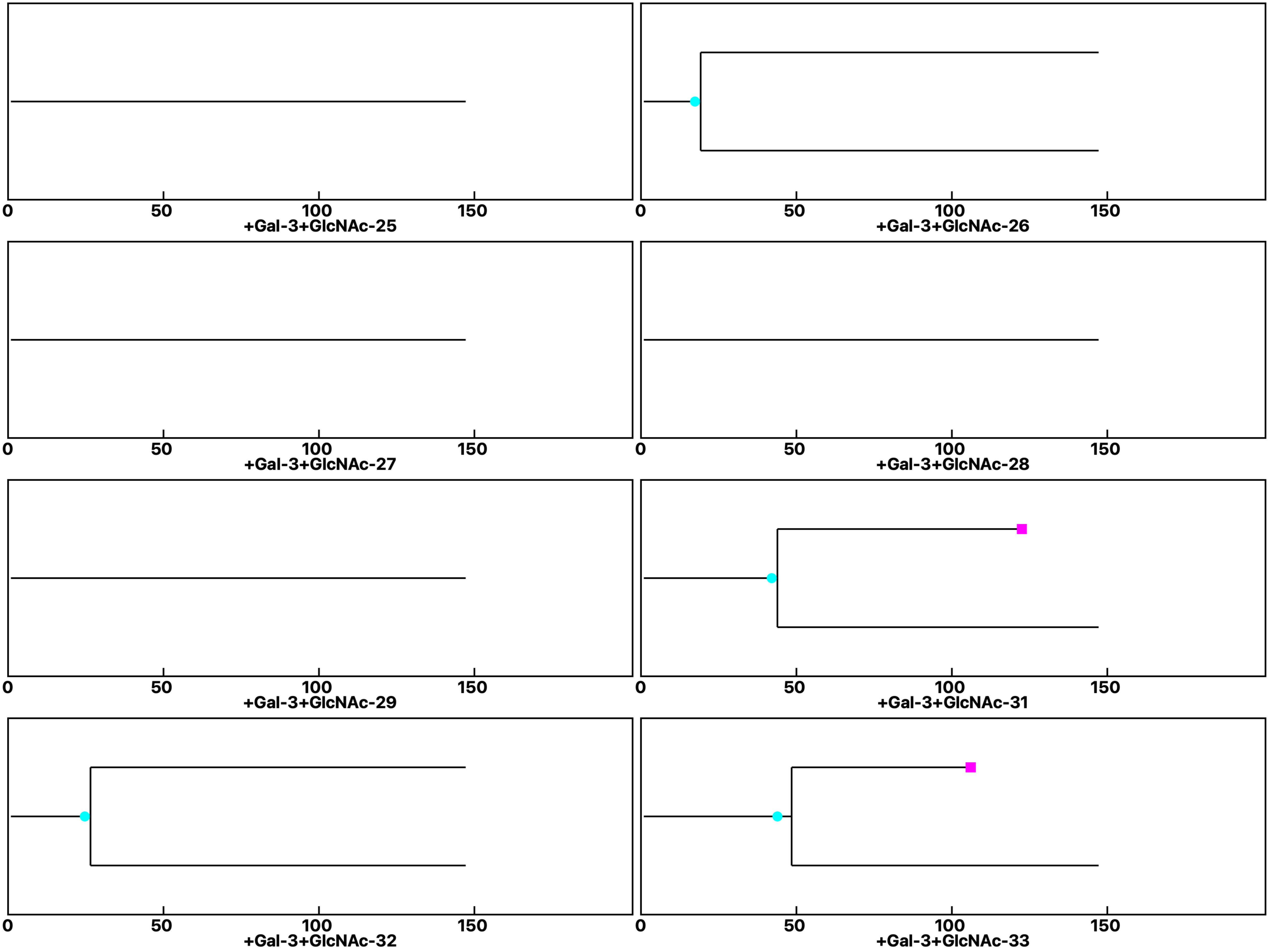

**Analysis: Gal-3KO Gal-3-GlcNAc, Treat.: +Gal-3+GlcNAc, Cell: Myoblasts**

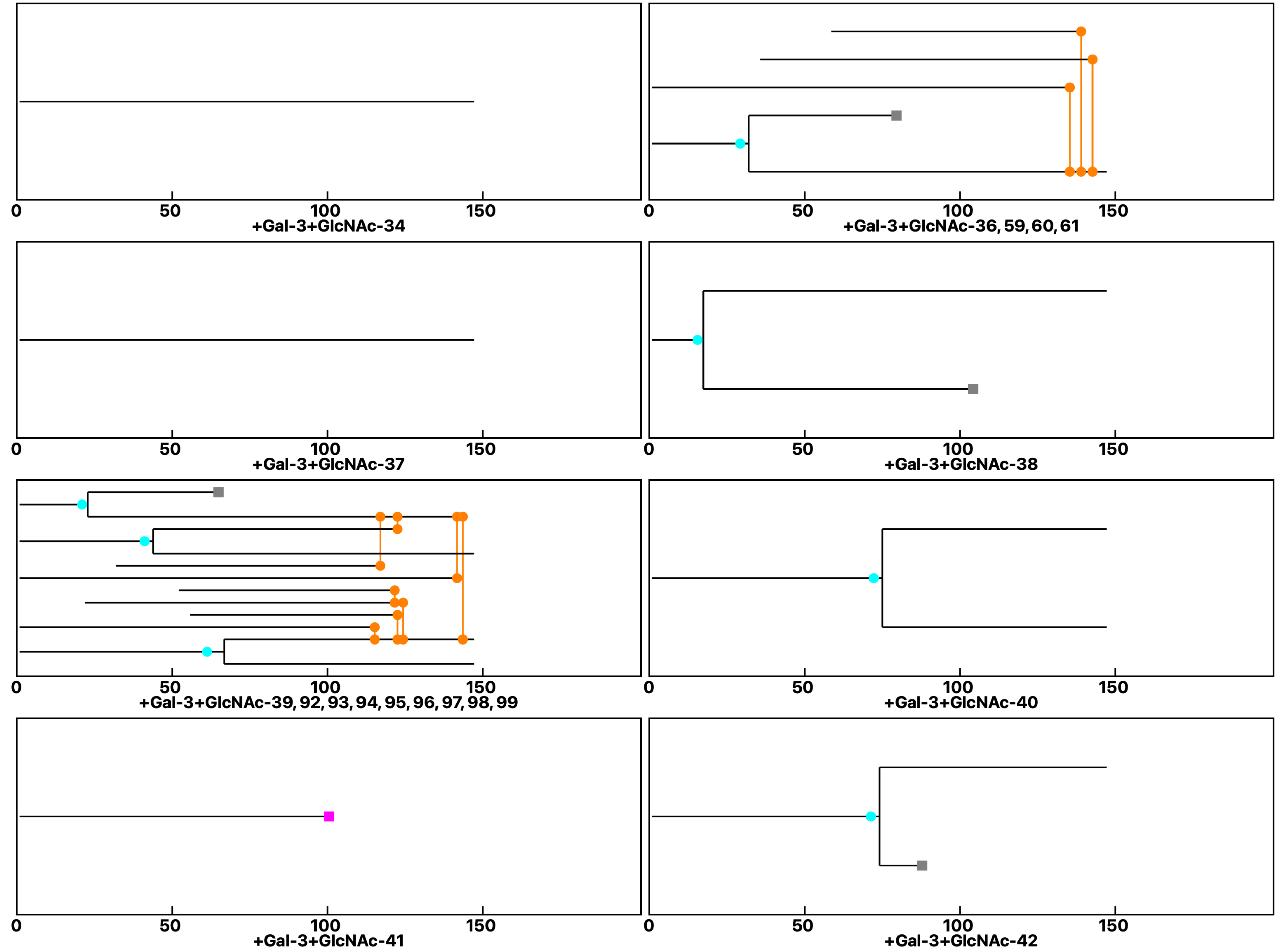

Analysis: Gal-3KO Gal-3-GlcNAc, Treat.: +Gal-3+GlcNAc, Cell: Myoblasts

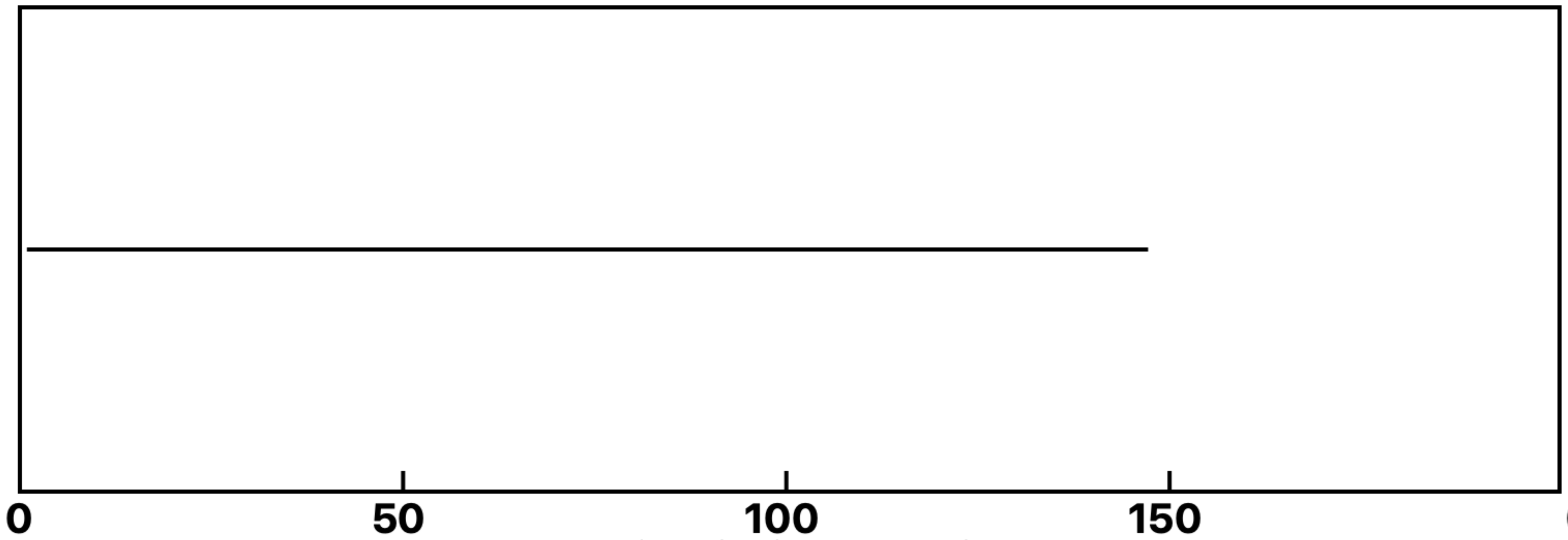


+Gal-3+GlcNAc-43

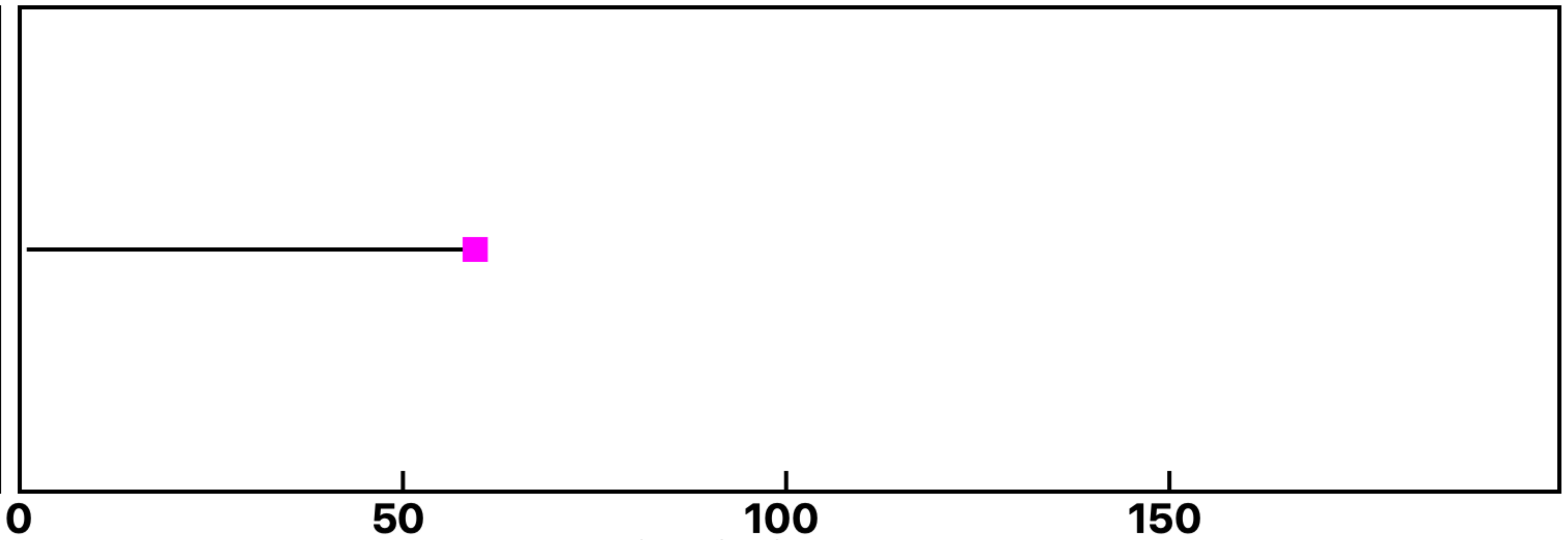


+Gal-3+GlcNAc-45

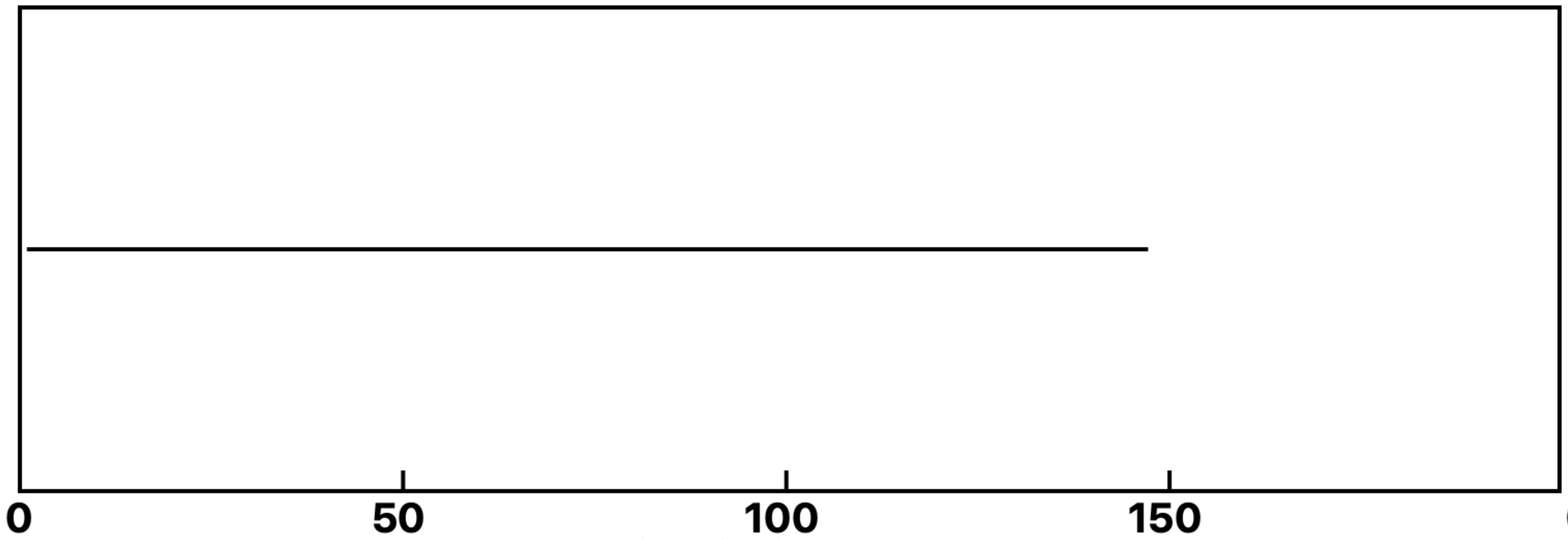


+Gal-3+GlcNAc-47

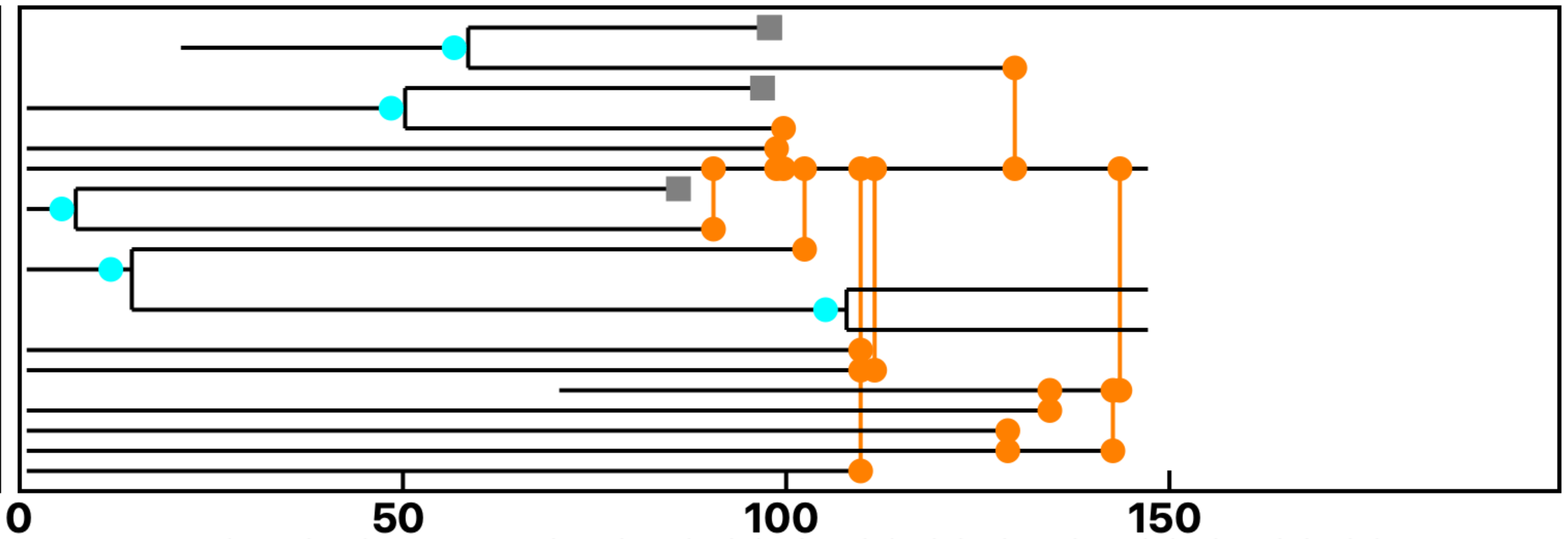


+Gal-3+GlcNAc-48, 78, 79, 80, 81, 82, 83, 84, 85, 86, 87, 88, 89
